# Network-based Virus-Host Interaction Prediction with Application to SARS-CoV-2

**DOI:** 10.1101/2020.11.09.375394

**Authors:** Hangyu Du, Feng Chen, Hongfu Liu, Pengyu Hong

## Abstract

COVID-19, caused by Severe Acute Respiratory Syndrome Coronavirus 2 (SARS-CoV-2), has quickly become a global health crisis since the first report of infection in December of 2019. However, the infection spectrum of SARS-CoV-2 and its comprehensive protein-level interactions with hosts remain unclear. There is a massive amount of under-utilized data and knowledge about RNA viruses highly relevant to SARS-CoV-2 and their hosts’ proteins. More in-depth and more comprehensive analyses of that knowledge and data can shed new insight into the molecular mechanisms underlying the COVID-19 pandemic and reveal potential risks. In this work, we constructed a multi-layer virus-host interaction network to incorporate these data and knowledge. A machine learning-based method, termed Infection Mechanism and Spectrum Prediction (IMSP), was developed to predict virus-host interactions at both protein and organism levels. Our approach revealed five potential infection targets of SARS-CoV-2, which deserved public health attention, and eight highly possible interactions between SARS-CoV-2 proteins and human proteins. Given a new virus, IMSP can utilize existing knowledge and data about other highly relevant viruses to predict multi-scale interactions between the new virus and potential hosts.

## Introduction

Severe Acute Respiratory Syndrome Coronavirus-2 (SARS-CoV-2), a novel virus causing the COVID-19 disease, was first reported in Wuhan, China, in December of 2019. Since then, it has quickly become a global health crisis (1) with over 50 million people infected and 1,250,000 death over 200 countries as of November of 2020 (2). The impact of SARS-CoV-2 has significantly surpassed previous out-breakings of coronaviruses, such as Severe Acute Respiratory Syndrome Coronavirus (SARS-CoV) in 2003 and the Middle East Respiratory Syndrome Coronavirus (MERS-CoV) in 2012. Besides humans, SARS-CoV-2 has also been confirmed to infect several other mammals closely related to human activities, including dogs (3), cats (4), tigers (5), and golden Syrian hamsters (6). It is important to identify a comprehensive set of such mammals because they can potentially serve as covert means to exacerbate the spread of COVID-19. Moreover, identifying interactions between SARS-CoV-2 proteins and host proteins can deepen our understanding of the viral invasion processes and may help design treatments and vaccines. In general, we want to promptly achieve the above two goals for new zoonotic viruses, which we believe can be done by leveraging the knowledge and data about known viruses highly relevant to the new ones.

The research community has accumulated a great deal of knowledge about several other human coronaviruses (including SARS-CoV (7–15), HCoV-HKU1 (15), HCoV-OC43 (16, 17), HCoV-NL63 (18), MERS-CoV (19–23)) and has collected a large amount of data about them. For example, it was shown that human Angiotensin-Converting Enzyme 2 (ACE2) was the primary host receptor used by the S protein (S-protein) of SARS-CoV-2 for the virus to gain entry into human cells (24) (Figure S1). ACE2 is also the host receptor used by SARS-CoV (14) and HCoV-NL63 (18). The S-protein of SARS-CoV-2 binds significantly tighter to ACE2 than its counterpart in SARS-CoV (25). After the virus enters host cells, interferon-stimulated genes (ISGs) are essential for a host to defend against viral infection (Figure S2). This knowledge and data can be utilized to investigate the infection spectrum of SARS-CoV-2 and its interactions with hosts at the protein level. Finally, we built a Virus-Host Interaction Network (VHIN) of seven viruses and seventeen hosts that summarizes the existing protein-protein interaction (PPI) and infection relationships among them (Figure 1A, more details in Figures S3, S4 and S5).

**Fig. 1.**
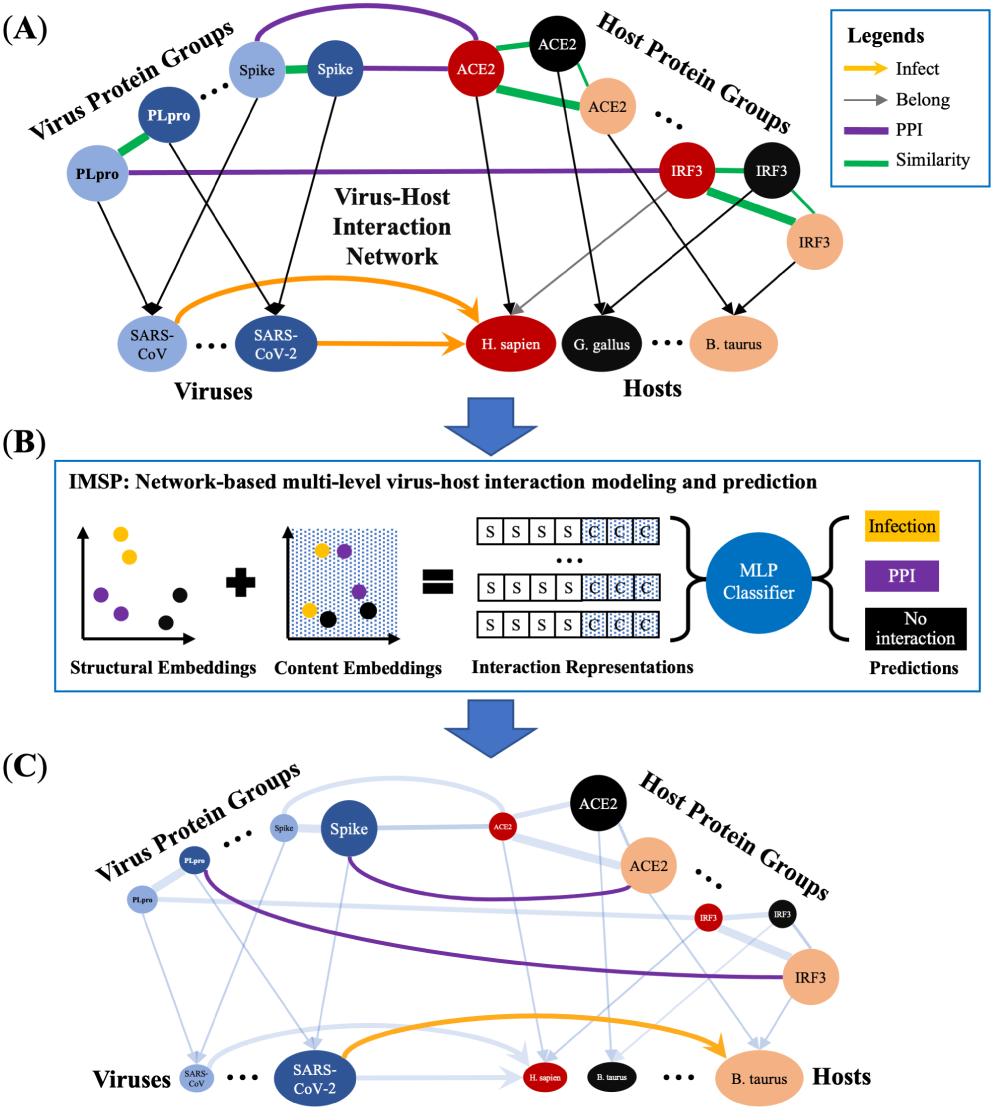
Infection Mechanism and Spectrum Prediction. **(A)** The virus-host interaction network. Nodes represent proteins, viruses, and hosts; interactions represent relationships (i.e., PPI, infect, similarity, belong). The color of a node indicates its organism. The thickness of a similarity relationship indicates its level of similarity. For the full network, refer to the viral entry graph (Figure S3), interferon signaling pathway graph (Figure S4), and infection graph (Figure S5). **(B)** IMSP learns a representation for each potential interaction, which contains a structural embedding and a content embedding. The structural embedding captures the local structural features of an interaction. The content embedding captures the attributes that reveal biological aspects of an interaction. The representation of each interaction is derived by concatenating its structural and content embeddings, where S stands for a structural embedding element, and C stands for a content embedding element. A Multi-layer Perceptron (MLP) is trained to classify the relationship between two nodes into PPI, infection, or no interaction. See the Methods section for calculating the structural and content embeddings. **(C)** Exemplar predicted interactions are highlighted and colored accordingly to their types. Existing interactions are dimmed.

We have developed a network-based multi-level virus-host interaction modeling and prediction, termed Infection Mechanism and Spectrum Prediction (IMSP) (Figure 1B, details in the Method section), which uses machine learning technique to learn from the constructed VHIN and then predict novel virus-host interactions at both the protein (i.e., Mechanism) and organism (i.e., Spectrum) levels. IMSP predicts that the SARS-CoV-2 S-protein can bind well with ACE2 receptors in seven mammalian hosts, which have not been reported. Among those hosts, five of them are predicted to have high risks of being infected by SARS-CoV-2. More-over, IMSP identifies eight new interactions between SARS-CoV-2 proteins and human proteins. To our best knowledge, our work is the first to apply machine learning techniques for predicting virus-host interactions at both protein and organism levels. Previous works (26, 27) on predicting virus-host interactions focus on the relationships between SARS-CoV-2 proteins and human proteins, which ignore other hosts that might be infected by SARS-CoV-2.

## Results

Here, we explain the construction of our virus-host interaction network in detail, highlight the predicted interactions by SARS-CoV-2, and present the overall performance evaluation of our interaction prediction model IMSP.

### The Virus–Host Interaction Network

The Virus-Host Interaction Network consists of two layers (an organism layer and a protein layer). The organism layer contains a set of viruses (including SARS-CoV-2, SARS-CoV, HCoV-229E, HCoV-HKU1, HCoV-OC43, HCoV-NL63, and MERS-CoV) and a set of hosts (including human, mouse, rat, dog, cat, camel, squirrel, cattle, chimpanzee, red junglefowl, rabbit, horse, monkey, rat, sheep, pig, and golden Syrian hamster). At the protein layer, we focus on proteins that are known to be involved in viral invasion as well as immune system response and suppression. The network contains 17 host protein homolog groups obtained from NCBI: ACE2, Dipeptidyl Peptidase 4 (DPP4), IRF3, IRF7, IRF9, Mitochondrial Anti-Viral-signaling protein (MAVS), Melanoma Differentiation-Associated protein 5 (MDA5), Nuclear Factor Kappa-light-chain-enhancer of activated B (NF-*κ*B), Protein Kinase interferon-inducible double-stranded RNA dependent Activator (PRKRA), TANK Binding Kinase 1 (TBK1), Retinoic Acid-Inducible Gene I (RIG-I), Signal Transducer and Activator of Transcription 1 (STAT1) and Signal Transducer and Activator of Transcription 2 (STAT2)). The viral proteins include homologs of Sprotein, Membrane (M) protein, Nucleocapsid (N) protein, nsp1, nsp15, ORF3b, ORF4a, ORF4b, ORF6, and Papainlike protease (PLpro). There are four types of interactions between nodes: Protein-Protein Interactions, infection, belonging, and similarity between protein homologs. Similarities between homologs were calculated using the Protein BLAST algorithm provided by NCBI. PPIs and infection relationships are gathered from academic publications (7–23).

### SARS-CoV-2 – Host Interaction Predictions

We applied IMSP on SARS-CoV-2 and other six human coronaviruses to get high-confidence predictions of PPIs and infections. Figure S1 shows the interactions between S proteins and ACE2. Figure S2 shows the interactions between virus proteins and IFN proteins in hosts. Figure S3 shows the network between the host group and the virus group in the organism layer. The virus-host network’s complete information, built with two layers of data containing 246 nodes and 1,766 interactions, is displayed in Node Table S4 and Interaction Table S5. All 65 infections and 457 PPIs predictions are shown in Table S6 and S7.

#### SARS-CoV-2 S-protein Binding Predictions

The ability of SARS-CoV-2’s S-protein to bind to the host ACE2 receptors is a key factor deciding the infection capability of SARS-CoV-2. IMSP predicts that the S-protein of SARS-CoV-2 has a high probability of binding well with the ACE2 receptors in mice, rats, sheep, chimpanzees, cattle, squirrels, and horses (Figure 2a). Mice and rats have been recognized to be susceptible to several other human coronaviruses, such as, SARS-CoV (7), MERS-CoV (28), HCoVOC43 (17) and HCoV-HKU1 (29, 30). The chimpanzee has the ACE2 with 99.38% similarity with human ACE2, which has a high possibility to bind with the same S-protein. The overall similarity of ACE2 for the horse, squirrel, sheep, and cattle is 93.45%, 91.82%, 90.81%, and 90.56% compared to human ACE2, respectively. These predictions still require more practical research is required to determine the binding affinity of these mammals’ ACE2 with SARS-CoV-2’s S-protein. It has been shown that ACE2 can tolerate up to seven amino acids changes out of 20 critical ones that contact with the S-protein, without losing the functionality as SARS-CoV-2’s receptor (31). This means that similarity might not be the only factor that influences the binding affinity between ACE2 and SARS-CoV-2’s S-protein.

**Fig. 2.**
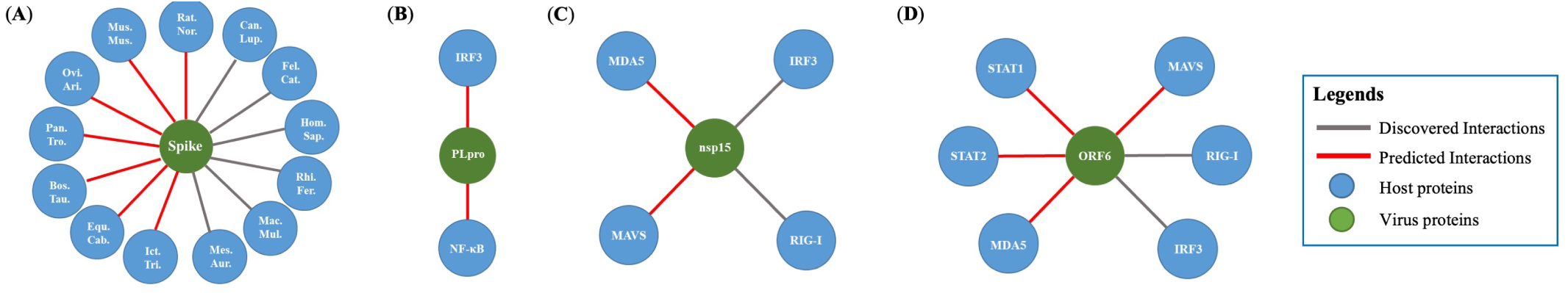
Interaction prediction for SARS-CoV-2. (A) shows the known and predicted bindings between the S-protein in SARS-CoV-2 and ACE2 in mammalian hosts. Host names are displayed in their abbreviation form: Hom.Sap. is Homo sapiens; Mus.Mus. is Mus musculus; Fel.Cat. is Felis catus; Can.Lup. is Canis lupus familiaris; Ovi.Ari. is Ovis aries; Rat.Nor. is Rattus norvegicus; Mac.Mul. is Macaca mulatta; Rhi.Fer. is Rhinolophus ferrumequinum; Mes.Aur. is Mesocricetus auratus; Bos.Tau. is Bos taurus; Ict.Tri. is Ictidomys tridecemlineatus; Equ.Cab. is Equus caballus; Pan.Tro. is Pan troglodytes. (B)-(D) show the known and predicted interactions of PLpro, nsp15, ORF6, and N protein in SARS-CoV-2 with proteins in the human IFN signaling pathway that contribute to the IFN signaling pathway suppression.

#### SARS-CoV-2 and human interferon pathway interactome prediction

The Interferon (IFN) pathway plays a critical role in the human immune response. After the virus infection is detected, the innate immune system will induce the IFN signaling, and the expression of IFN genes will increase the cellular resistance to viral invasion. Viruses have developed various strategies to inhibit IFN signaling to facilitate a successful viral invasion (32). SARS-CoV and MERS-CoV have been studied quite comprehensively in counteracting the IFN signaling responses compared with SARS-CoV-2. From IMSP, eight interactions between three SARS-CoV-2 proteins and human proteins in the innate immune pathway have been identified, shown in Figure 2(b)-(d). These interactions have a high probability of playing crucial roles in the SARS-CoV-2’s suppression of the host’s innate immune system.

PLpro was predicted to interact with the nuclear factor kappa-light-chain-enhancer of activated B (NF-*κ*B) in humans, shown in Figure 2(b). In SARS-CoV (33) and MERS-CoV (34), PLpro have been identified as being capable of reducing the NF-*κ*B reporter activity and further suppressing IRF3. From our prediction, it might also be highly possible for SARS-CoV-2’s PLpro to have similar functions.

ORF6 and nsp15 in SARS-CoV-2 have been discovered as crucial viral interferon antagonists of SARS-CoV-2. These two proteins inhibit the localization of IRF3 by interacting with RIG-I (35), as a similar function is found for SARS-CoV ORF6 (36), shown in Figure 2(c)&(d). ORF6 in SARS-CoV has also been shown to interact with STAT1 and STAT2 (37). From IMSP’s prediction, ORF6 and nsp15 in SARS-CoV-2 were suggested to have potential interactions with MDA5 and MAVS. Since MAVS works as the adaptor molecule for MDA5 (38), it is possible that a viral protein interacts with either one of these two would also have the interaction with the other. Besides, MAVS and MDA5, ORF6 was also predicted to interact with STAT1 and STAT2, the same function as in the SARS-CoV. As nsp15 and ORF6 both function in nuclear transport machinery after the viral entry (26), it is reasonable that, for these two proteins, similar interactions with innate immune pathways are predicted. Careful experiments should be conducted to identify nsp15 and ORF6’s impact on the innate immune system.

#### SARS-CoV-2 infection prediction

Based on the protein-level interaction prediction, we concluded five predicted infections for SARS-CoV-2 that is highly possible, as these mammals are not only predicted to be infected but also to have a successful SARS-CoV-2 virus binding with the host receptors. As shown in Figure 3, these animals include mice, rats, sheep, cattle, and squirrels.

**Fig. 3.**
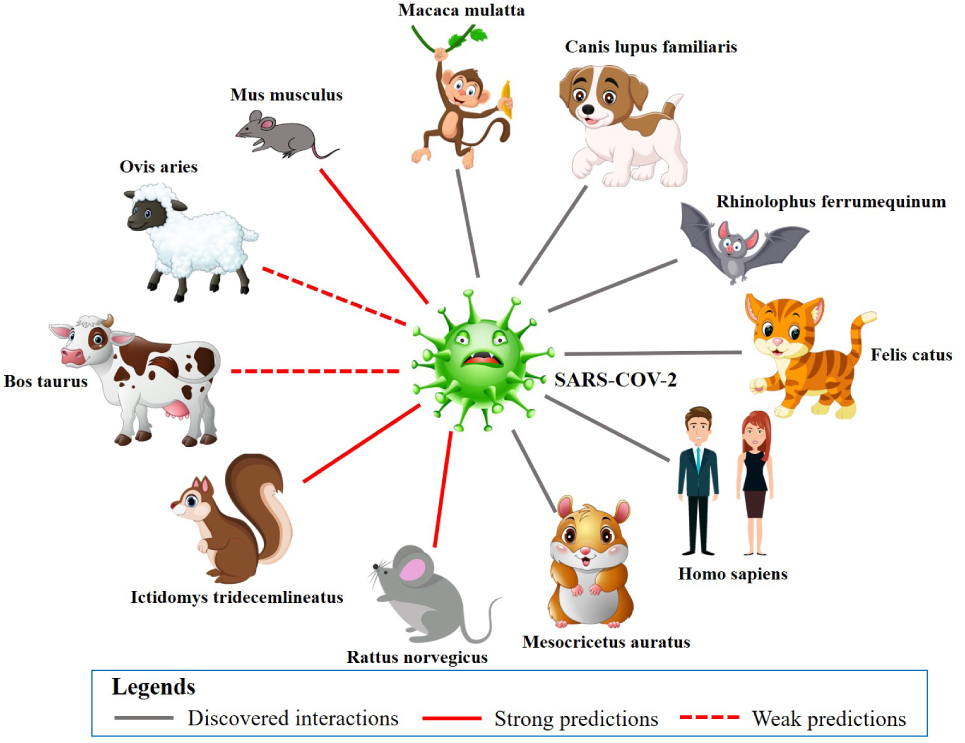
Infection prediction for SARS-CoV-2. This figure shows SARS-CoV-2’s known and predicted infected mammalian hosts. The predicted infected mammalian hosts without a predicted or proven corresponding receptor binding relationship with SARS-CoV-2’s S-protein were excluded.

Mice and rats are similar in every protein that we took into consideration in IMSP: all of the proteins have above 85% similarity, and the receptor ACE2 even has 94% similarity. Mouse has been identified as a host for all beta-coronaviruses: SASR-CoV (7), MERS-CoV (28), HCoV-OC43 (17) and HCoV-HKU1 (29, 30). SARS-CoV-2 also falls into the category of beta-coronavirus (39), which has a high possibility to infect mice and rats.

Cattle are hosts for both MERS-CoV (21) and a bovine coronavirus which is exceptionally similar to HCoV-OC43 (over 94% similarity) (16). This means that cattle can be the host for other coronaviruses. Cattle, along with sheep and squirrels, are closely related to the human living environment or daily diet. They could be potential mammalian hosts that again transmit the virus back to human society. The investigation of these highly possible infections could help identify the virus’s path and further control the transmission SARSCoV-2 from mammalian hosts. Further research on these potential hosts might be crucial to social health and safety.

### Interaction Prediction Evaluation

We compared IMSP to five other representative baseline models on our dataset in a cross-validation experiment. We conducted 30 runs in total for the experiment. The baseline models include two famous random-walk based models (Deepwalk (41) and Node2vec (43)), two neural-network-based models (Large-scale Information Network Embedding (LINE) (42) and Structural Deep Network Embedding (SDNE) (44)), and a classical matrix-based model, Graph Factorization (GF) (40). In each cross-validation run, we reserved 20% of each interaction type as the positive test set while ensuring the remaining network to be a connected one. To ensure input balance, we collected negative interactions (unconnected node pairs) to match the number of positive interactions in both training and testing sets. Firstly, we extracted known negative interactions as true negatives, which consisted of spike-receptor interactions that were demonstrated as not existed, such as between SARS-CoV-2’s S-protein and the host receptor DPP4 (the target host receptor of MERS-CoV). We split known negative interactions by a ratio of 8:2 to add to the training and testing sets. Then, since we still lack some number of negative interactions in both sets, we randomly selected unconnected node pairs as negative interactions to add to the training and testing sets. In this way, we confirmed that the training set’s composition was the same as the test set’s composition in terms of interaction type. We then applied IMSP and other models on the training set to perform network learning on the same training set and measure the test set’s interaction prediction performances. We repeated the entire measurement process for 30 runs to minimize the measurement randomness. Finally, we performed two-sample heteroscedastic T-test at the 0.01 significance level to test the significance of our model’s performance improvement compared to other models.

Table 1 shows the performance comparison, measured in several commonly used evaluation metrics, between IMSP and other existing tools. IMSP achieved an overall interaction prediction accuracy of 94.9% with a standard deviation of 0.004, which demonstrated a 6.0% gain compared to the second-best model. Our model also excelled in its weighted F1-score, achieving 0.944 with a standard deviation of 0.008, which transcended the second-best model by 7.3%. The pvalues for these two metrics were all smaller than 0.01, which indicated significant improvement for our model. In terms of the performance on specific types of interactions, i.e., infection and PPI predictions (See Fig. 4), IMSP achieved an F1-score of 0.808 with 0.119 standard deviation for infection predictions, a 14.9% increase compared to the second-best model. The p-value was also smaller than 0.01, indicating a significant improvement in our model. For PPI predictions, our model achieved an F1-score of 0.909 with 0.029 standard deviation, a 6.6% increment compared to the second-best tool. The p-value also demonstrated a significant improvement for IMSP. In conclusion, our model showed statistically significant improvements compared to all existing models in all 12 evaluation metrics.

**Table 1.**
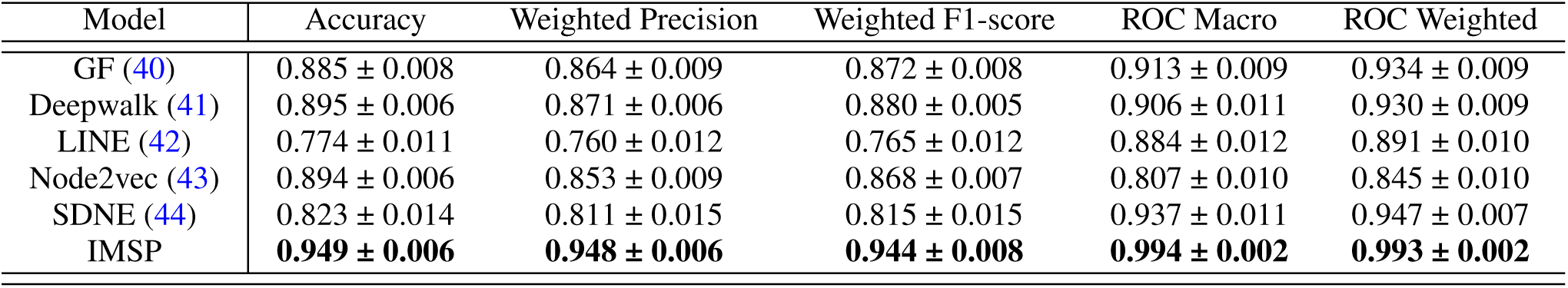
Interaction Prediction Overall Performance Evaluation and Comparison. This table gives six evaluation metrics regarding interaction prediction. During performance measurement, we generated two independent sets, training and testing sets, from the network. We set the ratio between the training and the test set to be 8:2. We ensured that the composition of positive interactions in the training set was the same as the test set. To guarantee input balance, we gathered negative interactions (unconnected node pairs) to correspondingly match the number of positive interactions in both training and testing sets. Part of negative samples was extracted from known negative interactions (true negatives), which consisted of spike-receptor interactions that were demonstrated as not existed. The remaining part of negative samples was randomly selected from other unconnected node pairs which were assumed as not existed. To eliminate performance randomness, we executed the comparison algorithm by 30 runs and recorded each run’s performance. We generated new training and test sets during each run and evaluated all models on the same test set. Then, we performed two-sample heteroscedastic T-tests for these six overall performance evaluation metrics to test the significance of IMSP’s improvement. Lastly, we reported the average with standard deviation.

| Model | Accuracy | Weighted Precision | Weighted F1-score | ROC Macro | ROC Weighted |
| --- | --- | --- | --- | --- | --- |
| GF (40) | 0.885 ± 0.008 | 0.864 ± 0.009 | 0.872 ± 0.008 | 0.913 ± 0.009 | 0.934 ± 0.009 |
| Deepwalk (41) | 0.895 ± 0.006 | 0.871 ± 0.006 | 0.880 ± 0.005 | 0.906 ± 0.011 | 0.930 ± 0.009 |
| LINE (42) | 0.774 ± 0.011 | 0.760 ± 0.012 | 0.765 ± 0.012 | 0.884 ± 0.012 | 0.891 ± 0.010 |
| Node2vec (43) | 0.894 ± 0.006 | 0.853 ± 0.009 | 0.868 ± 0.007 | 0.807 ± 0.010 | 0.845 ± 0.010 |
| SDNE (44) | 0.823 ± 0.014 | 0.811 ± 0.015 | 0.815 ± 0.015 | 0.937 ± 0.011 | 0.947 ± 0.007 |
| IMSP | **0.949 ± 0.006** | **0.948 ± 0.006** | **0.944 ± 0.008** | **0.994 ± 0.002** | **0.993 ± 0.002** |

**Table 2.**
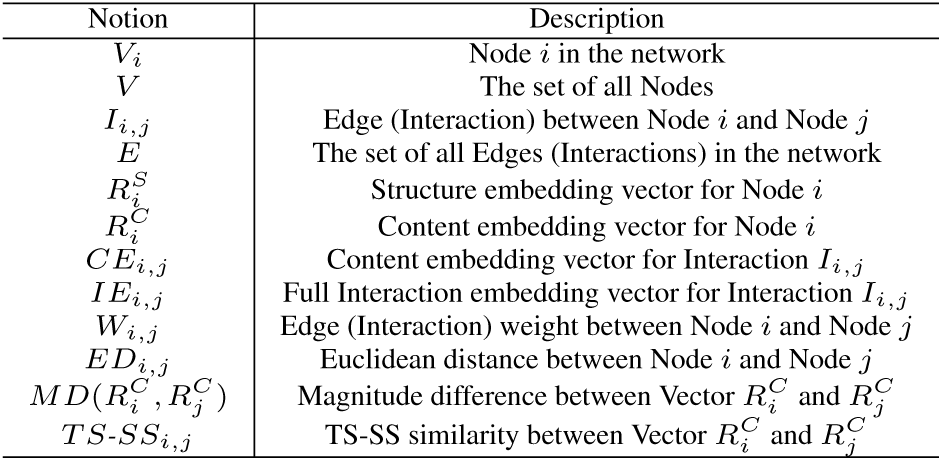
Notations.

| Notion | Description |
| --- | --- |
| $V_i$ | Node $i$ in the network |
| $V$ | The set of all Nodes |
| $I_{i,j}$ | Edge (Interaction) between Node $i$ and Node $j$ |
| $E$ | The set of all Edges (Interactions) in the network |
| $R_i^S$ | Structure embedding vector for Node $i$ |
| $R_i^C$ | Content embedding vector for Node $i$ |
| $CE_{i,j}$ | Content embedding vector for Interaction $I_{i,j}$ |
| $IE_{i,j}$ | Full Interaction embedding vector for Interaction $I_{i,j}$ |
| $W_{i,j}$ | Edge (Interaction) weight between Node $i$ and Node $j$ |
| $ED_{i,j}$ | Euclidean distance between Node $i$ and Node $j$ |
| $MD(R_i^C, R_j^C)$ | Magnitude difference between Vector $R_i^C$ and $R_j^C$ |
| $TS-SS_{i,j}$ | TS-SS similarity between Vector $R_i^C$ and $R_j^C$ |

**Fig. 4.**
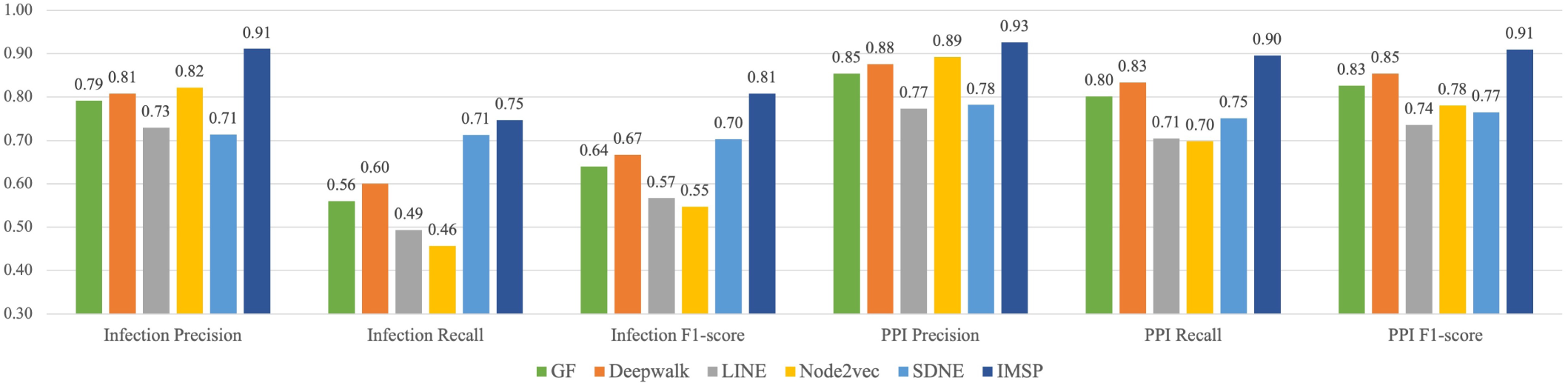
Performance on PPI and Infection Predictions. This figure demonstrates IMSP’s performance on PPI and infection predictions in comparison with five other models. The ocean-blue columns represent the performance of our IMSP model derived from the average of 30 independent runs. As a result, the IMSP model achieved 0.808 for the Infection F1-score and 0.910 for the PPI F1-score. Compared to other models, our model outperformed other models in all the evaluation metrics. Specifically, in terms of infection F1-score, our model outperformed the second-best model SDNE (44) by 14.9%. In terms of PPI F1-score. Our model also surpassed the second-best model Deepwalk (41) by 6.6%.

IMSP’s high performance might result from its ability to take full advantage of well-studied knowledge and data from previous biology research with protein-level variations being considered. Thanks to our virus-host interaction network’s novel design, such cross-organism information and multi-class linkage information can be well-preserved. Another reason behind IMSP’s performance improvement is that it factors essential biological metadata as textualized features for nodes. In this way, two connected nodes’ vectorized content substantially helps the classifier output a correct predicted class when formulating interaction representations. However, around 10% PPI predictions were unlikely by our definition, such as PPIs between S-protein and nonreceptor proteins. To minimize unlikely predictions, we incorporated the node’s function and existing biological metadata as part of the node feature. This change significantly reduced unlikely PPI predictions to around 2%. We further pushed the boundaries by utilizing known negative interactions in the protein layer to constitute part of the training and testing sets—this finally reduced the unlikely PPI predictions to less than 1%.

In conclusion, IMSP exhibited robust and stable performance in both top-level and detailed evaluation metrics, which is substantially improved compared to existing tools. When analyzing newly emerged viruses with limited available information, namely SARS-CoV-2, IMSP can provide reasonable and reliable predictions.

## Discussion

This study assembled 246 nodes (viruses, virus proteins, hosts, and host proteins) and 1,766 known interactions among these nodes, including four types of interactions: similarities, infections, interactions, and belongings. Two main parts for virus-host protein-level interactions were taken into consideration: viral entry and innate immune pathway interactions. Accompanied by these two parts, we predicted the potential host for viruses and undiscovered novel PPIs. Among seven human coronaviruses, SARS-CoV and MERS-CoV were well studied about their binding with the receptors and interacting with the innate immune pathway, whereas HCoVOC43, HCoV-NL63, HCov-HKU1, HCoV-229E, and newly emerged SARS-CoV-2 remained relatively unknown. Based on protein similarity and known interactions, 322 PPIs were predicted with this model. The actual PPIs need to be investigated further based on the prediction result.

Established discoveries about SARS-CoV-2’s viral interaction with host proteins are scarce. SARS-CoV-2 has been highly suspected of suppressing the innate immune response and reducing the production of IFN. In this way, IMSP’s findings could help discover the relations between virus invasion and host response and therefore provide clues to developing therapeutic strategies for the treatment of this disease.

More broadly, IMSP could be applied to any other analysis of the virus-host interaction network predictions. This network would be built based on the information of the protein-protein relations, protein-host relations, and host-host relations. Based on these attributes in the network and currently known relationships between some of the proteins, IMSP could predict the high-possibility interactions. We hope to use this pipeline as a tool to build a guideline for investigating various similar viruses and their mechanisms with hosts.

## Methods

### Infection Mechanism and Spectrum Prediction

Our IMSP model requests three different inputs, including pairwise similarity matrices for all host proteins and virus proteins, a set of known PPIs and infection relationships, and proteins’ biological metadata. For instance, appropriate protein metadata can include a protein’s name, layer, host, and function. Given these three inputs, the model constructs a virus-host interaction network. Then, in the network learning phase, IMSP learns and combines the structural information and the content information of interactions to represent the heterogeneous network comprehensively. Lastly, in the prediction phase, IMSP trains a neural-network-based classifier, Multi-layer Perceptron classifier, along with strict filtering procedures, to output high-possibility undiscovered interactions. Specifically, our model generated PPI and infection predictions with each prediction’s likelihood. In the following, we elaborated on the details of our IMSP model’s two main steps in terms of network construction and learning, and interaction prediction with result validation.

### Virus-host interaction network construction and learning

We utilized layers and groups to mark and separate different nodes during virus-host interaction network construction, and used interactions to represent PPI/infection/similarity/belonging relationship between nodes. For modeling the network, we constructed a weighted and undirected heterogeneous network using NetworkX (45). The network contained four groups of nodes: host, host proteins, virus proteins, and virus. We organize the virus group and the host group into the same layer, i.e., the organism layer, and we put two groups of proteins into the protein layer. By nature, the network contained four types of interactions: PPI (between virus proteins groups and host proteins groups), infection (between viruses group and hosts group), similarity (between homologs of proteins) and belonging (between organism layer and protein layer). Similarity and belonging relationships were innately connected. Other types of interactions were connected based on proven molecular level knowledge from existing research (7–23). The virus-host interaction network comprised of 246 nodes and 1,766 interactions. After the network structure was built, we used *W*_*i,j*_ to represent the biological closeness between *V*_*i*_ and *V*_*j*_. Specifically, if *I*_*i,j*_ denoted the relationship between two protein homologs, we used the full-sequence similarity between *V*_*i*_ and *V*_*j*_ from pairwise similarity matrices to represent *W*_*i,j*_. For other types of relationship, we obtained the embeddings of node content, denoted as 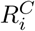 for *V*_*i*_ by passing textualized node content to the Word2vec model (46), a famous and powerful neural-network-based word embedding model. Then, we utilized the *TS-SS* similarity metric (47), a robust and reliable similarity measurement in the field of textual mining, to calculate the similarity between 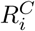 and 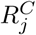 to assign to *W*_*i,j*_. Precisely, for interactions other than similarity relations, we calculated the *TS-SS*_*i,j*_ for all *i* and *j* if *I*_*i,j*_ ⊂ *E* as the following:

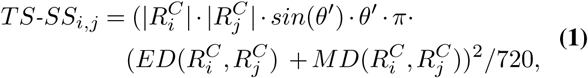

where 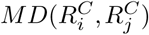 (47) is defined as the magnitude difference between 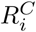 and 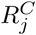, which is calculated as:

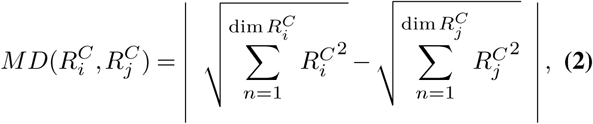

and *θ′* is defined as:

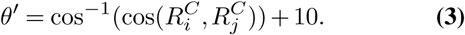

Note that *θ′* is increased by 10 degrees to overcome the problem of overlapping vectors. Then, *W*_*i,j*_ is calculated as:

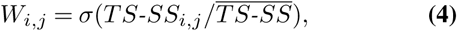

where *σ* is the sigmoid function, and 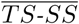 denotes the average of *TS-SS*_*i,j*_, for all *i, j*, if *i* ≠ *j* and *V*_*i*_, *V*_*j*_ ⊂*V*.

For network learning, we captured the network’s heterogeneity by adding the network’s heterogeneous content information to its structural information. Specifically, we performed network structural embedding assuming the network is homogeneous. Then, we added the content embedding on top of structural embedding to model its heterogeneity.

Firstly, for network structural embedding, we used a powerful network feature learning model, Node2vec (43), to learn the structural embedding for nodes. Node2vec is a stateof-the-art model for homogeneous network embedding. We took full advantage of Node2vec’s biased searching algorithm during our application. Precisely, the Node2vec model performed a fixed-length random walk for sampling, which takes weight into account. Let *c*_*m*_ denote the *m*-th node in walk with *c*_0_ denoting the starting node of the current random walk. Nodes *c*_*m*_ are generated by the following distribution:

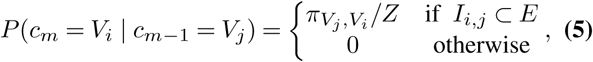

where *m* ⩾ 1, Z is the normalizing constant, and 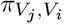 is the unnormalized transition probability between *V*_*i*_ and *V*_*j*_, which is calculated as 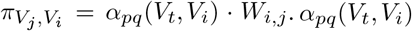, termed as search bias, is calculated as:

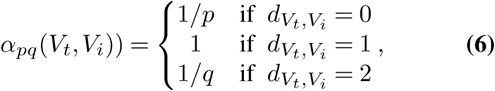

where 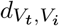 denotes the shortest path between *V*_*i*_ and *V*_*j*_, assuming that we have just traversed from *V*_*t*_ to *V*_*i*_ and are now evaluating the transition probability leaving *V*_*i*_.

In Eq. (6), p and q are Node2vec’s hyperparameters that can be adjusted to influence the probability to go back to *V*_*i*_ after visiting *V*_*j*_ and the probability to explore undiscovered components of the network. In this way, we were able to fine-tune the structural embedding model, Node2vec, to generate a conclusive embedding for our network.

Secondly, to create content representations, *CE*_*i,j*_ for all *I*_*i,j*_ in the network, we combined the textualized node content of *V*_*i*_ and *V*_*j*_ along with expected interaction type such as interacts/infects/similar/belongs, and input them into the Word2vec model (46) to generate vectorized interaction representations. Then, the full interaction representations for interaction *I*_*i,j*_ are formulated as follows:

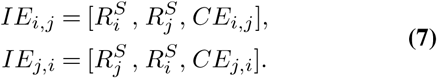

Note that by Word2vec’s nature, the input documents’ order does not affect its output, meaning that *CE*_*i,j*_ is the same as *CE*_*j,i*_. By finishing this step, we obtained all interaction representations, *IE*_*i,j*_, for all *V*_*i*_ and *V*_*j*_ ⊂ *V* and *i* ≠*j*.

### Virus-host interaction prediction

In the interaction prediction phase, we utilized a classifier to perform a multi-class classification task on the interaction representations to learn from the existing network and suggest potential undiscovered interactions. Specifically, for all *I*_*i,j*_ ⊄ *E*, we classified *IE*_*i,j*_ into three classes, i.e. PPI/infections/no-interaction. Scikit-learn’s Multi-layer Perceptron (MLP) classification model (48) was used for this multi-class interaction classification.

We conducted cross-validation experiments to fine-tune the model, where 20% of each interaction type was reserved as a positive test set. In this process, we also balanced the test set by inserting an equal number of negative interactions containing both known negatives and unknown negatives. Specifically, known negative interactions (true negatives) were generated from proven unconnected node pairs, such as the interaction between MERS-CoV’s S-protein and mammalian hosts’ ACE2 receptors. Unknown negative interactions were randomly sampled from the network, which was assumed as not existing. Then, by using the fine-tuned hyperparameters from cross-validation experiments, all positive interactions with an equal number of negative interactions were used for training the classifier. IMSP later took all other unused un-connected node pairs as input data and classified them into the class of PPI/infection/no interaction.

During prediction result correction, we defined rules from both computational and biological perspectives to mark and remove unlikely predictions. Computationally, because there exist two representations for *I*_*i,j*_, i.e., *IE*_*i,j*_ and *IE*_*j,i*_, we defined the prediction for *I*_*i,j*_ as a strong one if and only if *IE*_*i,j*_ and *IE*_*j,i*_ were all classified into the same type of interaction other than the no-interaction. Biologically, we assumed that each virus’s S-protein only interacts/binds with its target receptor. Thus, all other interactions with S-protein are considered unlikely. For example, the predicted interaction, if any, between SARS-CoV-2’s S-protein and DPP4 (the target receptor for MERS-CoV’s S-protein) would be eliminated. Thirdly, we also defined that an infection interaction was likely only when the corresponding interaction between the virus’ S-protein and an appropriate host receptor was predicted. In this way, we ensured that all the remaining predictions are both computationally and biologically meaningful and practically useful to guide further research.

## Supporting information

Latex Package

## Resource Availability

All data and codes are available at Github repositories: IMSP model, its predictions and performance evaluations can be found at https://github.com/SupremeEthan/IMSP;

Data and data parsing code can be found at https://github.com/SupremeEthan/IMSP-Parser.

## Acknowledgement

This work was partially supported NSF OAC 1920147.

## Authors’ Contributions

Conceptualization, P.H. and H.L.; Methodology, H.D., F.C., and H.L.; Software, H.D.; Formal Analysis: H.D.; Investigation, F.C. and H.D.; Writing – Original Draft, F.C. and H.D.; Writing – Review & Editing, all authors; Visualization: F.C. and H.D.; Supervision, H.L. and P.H.

## Lead Contact

Further information and requests for code and data should be directed to and will be fulfilled by the Lead Contact, Pengyu Hong.

