## Supplementary material for "Network-based Virus-Host Interaction Prediction with Application to SARS-CoV-2": Latex Package: main.tex

\documentclass[times, twoside]{style}
% Please give the surname of the lead author for the running footer
\leadauthor{H. Du \& F. Chen}
% \usepackage{dblfloatfix}
\usepackage{nidanfloat} % figure position adjustment
\usepackage[normalem]{ulem}
\usepackage{balance}

%%% HELPER CODE FOR DEALING WITH EXTERNAL REFERENCES
\usepackage{xr}
\usepackage{graphicx}
% \usepackage{sidecap}
\makeatletter
\newcommand*{\addFileDependency}[1]{
  \typeout{(#1)}
  \@addtofilelist{#1}
  \IfFileExists{#1}{}{\typeout{No file #1.}}
}
\makeatother

\newcommand*{\myexternaldocument}[1]{
    \externaldocument{#1}
    \addFileDependency{#1.tex}
    \addFileDependency{#1.aux}
}
%%% END HELPER CODE

% put all the external documents here!
\myexternaldocument{supplementary}

\useunder{\uline}{\ul}{}
\newcommand{\td}{{\bf\color{red} FIXME~}}
\begin{document}

% -----------------------------------------------------------
% -----------------------title section-----------------------
% -----------------------------------------------------------

% Use letters for affiliations, numbers to show equal authorship (if applicable) and to indicate the corresponding author
\author[$\dag$]{Hangyu Du}
\author[$\dag$]{Feng Chen}
\author[*]{Hongfu Liu}
\author[*]{Pengyu Hong}

% Cell: department(s)/subunit(s), institution, city, state/region (postal code optional), country.
\affil{Department of Computer Science, Brandeis University, Waltham, Massachusetts, United States}
\affil[$\dag$]{These authors contribute equally}

\title{Network-based Virus-Host Interaction Prediction with Application to SARS-CoV-2}

\date{\today}

\maketitle

% -----------------------------------------------------------
% ---------------------abstract section----------------------
% -----------------------------------------------------------

\begin{abstract}
COVID-19, caused by Severe Acute Respiratory Syndrome Coronavirus 2 (\mbox{SARS-CoV-2}), has quickly become a global health crisis since the first report of infection in December of 2019. However, the infection spectrum of SARS-CoV-2 and its comprehensive protein-level interactions with hosts remain unclear. There is a massive amount of under-utilized data and knowledge about RNA viruses highly relevant to \mbox{SARS-CoV-2} and their hosts' proteins. More in-depth and more comprehensive analyses of that knowledge and data can shed new insight into the molecular mechanisms underlying the \mbox{COVID-19} pandemic and reveal potential risks. In this work, we constructed a multi-layer virus-host interaction network to incorporate these data and knowledge. A machine learning-based method, termed Infection Mechanism and Spectrum Prediction (IMSP), was developed to predict virus-host interactions at both protein and organism levels. Our approach revealed five potential infection targets of \mbox{SARS-CoV-2}, which deserved public health attention, and eight highly possible interactions between \mbox{SARS-CoV-2} proteins and human proteins. Given a new virus, IMSP can utilize existing knowledge and data about other highly relevant viruses to predict multi-scale interactions between the new virus and potential hosts.
\end{abstract}

\begin{keywords}
Keywords: Coronavirus, COVID-19, SARS-CoV-2, Machine Learning, Interaction Prediction, Protein-Protein-Interaction, virus-host interaction network
\end{keywords}

% -----------------------------------------------------------
% --------------------introduction section-------------------
% -----------------------------------------------------------

\section*{Introduction}
Severe Acute Respiratory Syndrome Coronavirus-2 (SARS-CoV-2), a novel virus causing the COVID-19 disease, was first reported in Wuhan, China, in December of 2019. Since then, it has quickly become a global health crisis~\cite{li2020early} with over 50 million people infected and 1,250,000 death over 200 countries as of November of 2020~\cite{world2020coronavirus}. The impact of \mbox{SARS-CoV-2} has significantly surpassed previous outbreakings of coronaviruses, such as Severe Acute Respiratory Syndrome Coronavirus (SARS-CoV) in 2003 and the Middle East Respiratory Syndrome Coronavirus (MERS-CoV) in 2012. Besides humans, SARS-CoV-2 has also been confirmed to infect several other mammals closely related to human activities, including dogs~\cite{sit2020infection}, cats~\cite{halfmann2020transmission}, tigers~\cite{wang2020complete}, and golden Syrian hamsters~\cite{sia2020pathogenesis}. It is important to identify a comprehensive set of such mammals because they can potentially serve as covert means to exacerbate the spread of \mbox{COVID-19}. Moreover, identifying interactions between \mbox{SARS-CoV-2} proteins and host proteins can deepen our understanding of the viral invasion processes and may help design treatments and vaccines. In general, we want to promptly achieve the above two goals for new zoonotic viruses, which we believe can be done by leveraging the knowledge and data about known viruses highly relevant to the new ones.

\ \ \ The research community has accumulated a great deal of knowledge about several other human coronaviruses (including  SARS-CoV~\cite{channappanavar2016dysregulated, frieman2008sars, matthews2014sars, totura2012sars, hu2017sars, chen2014sars, li2010host, li2003angiotensin, lei2009mavs}, HCoV-HKU1~\cite{lei2009mavs}, HCoV-OC43~\cite{szczepanski2019canine,niu2020non}, HCoV-NL63~\cite{hofmann2005human}, MERS-CoV~\cite{haagmans2014middle, mou2013receptor, kandeil2019middle, bailey2014crystal, lui2016middle}) and has collected a large amount of data about them. For example, it was shown that human Angiotensin-Converting Enzyme 2 (ACE2) was the primary host receptor used by the S protein (S-protein) of \mbox{SARS-CoV-2} for the virus to gain entry into human cells ~\cite{zhang2020angiotensin} (Figure S1). ACE2 is also the host receptor used by SARS-CoV~\cite{li2003angiotensin} and HCoV-NL63~\cite{hofmann2005human}. The S-protein of \mbox{SARS-CoV-2} binds significantly tighter to ACE2 than its counterpart in SARS-CoV~\cite{shang2020structural}. After the virus enters host cells, interferon-stimulated genes (ISGs) are essential for a host to defend against viral infection (Figure S2). This knowledge and data can be utilized to investigate the infection spectrum of SARS-CoV-2 and its interactions with hosts at the protein level. Finally, we built a Virus-Host Interaction Network (VHIN) of seven viruses and seventeen hosts that summarizes the existing protein-protein interaction (PPI) and infection relationships among them (Figure \ref{fig:model_graph}A, more details in Figures S3, S4 and S5). 

% model and conclusion
\ \ \ We have developed a network-based multi-level virus-host interaction modeling and prediction, termed Infection Mechanism and Spectrum Prediction (IMSP) (Figure \ref{fig:model_graph}B, details in the Method section), which uses machine learning technique to learn from the constructed VHIN and then predict novel virus-host interactions at both the protein (i.e., Mechanism) and organism (i.e., Spectrum) levels. IMSP predicts that the SARS-CoV-2 S-protein can bind well with ACE2 receptors in seven mammalian hosts, which have not been reported. Among those hosts, five of them are predicted to have high risks of being infected by SARS-CoV-2. Moreover, IMSP identifies eight new interactions between SARS-CoV-2 proteins and human proteins. To our best knowledge, our work is the first to apply machine learning techniques for predicting virus-host interactions at both protein and organism levels. Previous works ~\cite{gordon2020sars,messina2020covid} on predicting virus-host interactions focus on the relationships between \mbox{SARS-CoV-2} proteins and human proteins, which ignore other hosts that might be infected by \mbox{SARS-CoV-2}.

\begin{figure}[!h]
  \centering
  \includegraphics[width=8.5cm]{Figures/model.png}
  \caption{\textbf{Infection Mechanism and Spectrum Prediction.} \textbf{(A)} The virus-host interaction network. Nodes represent proteins, viruses, and hosts; interactions represent relationships (i.e., PPI, infect, similarity, belong). The color of a node indicates its organism. The thickness of a similarity relationship indicates its level of similarity. For the full network, refer to the viral entry graph (Figure S3), interferon signaling pathway graph (Figure S4), and infection graph (Figure S5). \textbf{(B)} IMSP learns a representation for each potential interaction, which contains a structural embedding and a content embedding. The structural embedding captures the local structural features of an interaction. The content embedding captures the attributes that reveal biological aspects of an interaction. The representation of each interaction is derived by concatenating its structural and content embeddings, where S stands for a structural embedding element, and C stands for a content embedding element. A Multi-layer Perceptron (MLP) is trained to classify the relationship between two nodes into PPI, infection, or no interaction. See the Methods section for calculating the structural and content embeddings. \textbf{(C)} Exemplar predicted interactions are highlighted and colored accordingly to their types. Existing interactions are dimmed. }
  \medskip
  \label{fig:model_graph}
\end{figure}
% -----------------------------------------------------------
% ----------------------results section----------------------
% -----------------------------------------------------------
\section*{Results}
Here, we explain the construction of our virus-host interaction network in detail, highlight the predicted interactions by \mbox{SARS-CoV-2}, and present the overall performance evaluation of our interaction prediction model IMSP.

\subsection*{The Virus--Host Interaction Network\\}
The Virus-Host Interaction Network consists of two layers (an organism layer and a protein layer). The organism layer contains a set of viruses (including \mbox{SARS-CoV-2}, \mbox{SARS-CoV}, \mbox{HCoV-229E}, \mbox{HCoV-HKU1}, \mbox{HCoV-OC43}, \mbox{HCoV-NL63}, and \mbox{MERS-CoV}) and a set of hosts (including human, mouse, rat, dog, cat, camel, squirrel, cattle, chimpanzee, red junglefowl, rabbit, horse, monkey, rat, sheep, pig, and golden Syrian hamster). At the protein layer, we focus on proteins that are known to be involved in viral invasion as well as immune system response and suppression. The network contains 17 host protein homolog groups obtained from NCBI: ACE2, Dipeptidyl Peptidase 4 (DPP4), IRF3, IRF7, IRF9, Mitochondrial Anti-Viral-signaling protein (MAVS), Melanoma Differentiation-Associated protein 5 (MDA5), Nuclear Factor Kappa-light-chain-enhancer of activated B (NF-$\kappa$B), Protein Kinase interferon-inducible double-stranded RNA dependent Activator (PRKRA), TANK Binding Kinase 1 (TBK1), Retinoic Acid-Inducible Gene I (RIG-I), Signal Transducer and Activator of Transcription 1 (STAT1) and Signal Transducer and Activator of Transcription 2 (STAT2)). The viral proteins include homologs of S-protein, Membrane (M) protein, Nucleocapsid (N) protein, nsp1, nsp15, ORF3b, ORF4a, ORF4b, ORF6, and Papain-like protease (PLpro). There are four types of interactions between nodes: Protein-Protein Interactions, infection, belonging, and similarity between protein homologs. Similarities between homologs were calculated using the Protein BLAST algorithm provided by NCBI. PPIs and infection relationships are gathered from academic publications~\cite{channappanavar2016dysregulated, frieman2008sars, matthews2014sars, totura2012sars, hu2017sars, chen2014sars, li2010host, li2003angiotensin, lei2009mavs,szczepanski2019canine,niu2020non,hofmann2005human,haagmans2014middle, mou2013receptor, kandeil2019middle, bailey2014crystal, lui2016middle}.

\subsection*{SARS-CoV-2 -- Host Interaction Predictions\\}
We applied IMSP on SARS-CoV-2 and other six human coronaviruses to get high-confidence predictions of PPIs and infections. Figure S1 shows the interactions between S proteins and ACE2. Figure S2 shows the interactions between virus proteins and IFN proteins in hosts. Figure S3 shows the network between the host group and the virus group in the organism layer. The virus-host network's complete information, built with two layers of data containing 246 nodes and 1,766 interactions, is displayed in Node Table S4 and Interaction Table S5. All 65 infections and 457 PPIs predictions are shown in Table S6 and S7.

\subsubsection*{SARS-CoV-2 S-protein Binding Predictions\\}
\begin{figure*}[!ht]
    \centering
    \includegraphics[width=18cm]{Figures/protein prediction SARS-CoV-2.jpg}
    \caption{\textbf{Interaction prediction for SARS-CoV-2.} \textit{(A) shows the known and predicted bindings between the S-protein in SARS-CoV-2 and ACE2 in mammalian hosts. Host names are displayed in their abbreviation form: Hom.Sap. is Homo sapiens; Mus.Mus. is Mus musculus; Fel.Cat. is Felis catus; Can.Lup. is Canis lupus familiaris; Ovi.Ari. is Ovis aries; Rat.Nor. is Rattus norvegicus; Mac.Mul. is Macaca mulatta; Rhi.Fer. is Rhinolophus ferrumequinum; Mes.Aur. is Mesocricetus auratus; Bos.Tau. is Bos taurus; Ict.Tri. is Ictidomys tridecemlineatus; Equ.Cab. is Equus caballus; Pan.Tro. is Pan troglodytes. (B)-(D) show the known and predicted interactions of PLpro, nsp15, ORF6, and N protein in SARS-CoV-2 with proteins in the human IFN signaling pathway that contribute to the IFN signaling pathway suppression.}}
    \medskip
    \label{fig:interactions}
\end{figure*}

The ability of \mbox{SARS-CoV-2}'s S-protein to bind to the host ACE2 receptors is a key factor deciding the infection capability of \mbox{SARS-CoV-2}. IMSP predicts that the S-protein of \mbox{SARS-CoV-2} has a high probability of binding well with the ACE2 receptors in mice, rats, sheep, chimpanzees, cattle, squirrels, and horses (Figure~\ref{fig:interactions}a). Mice and rats have been recognized to be susceptible to several other human coronaviruses, such as, SARS-CoV~\cite{channappanavar2016dysregulated}, MERS-CoV~\cite{cockrell2016mouse}, HCoV-OC43~\cite{niu2020non} and HCoV-HKU1~\cite{gorse2009human,lim2016human}. The chimpanzee has the ACE2 with 99.38\% similarity with human ACE2, which has a high possibility to bind with the same S-protein. The overall similarity of ACE2 for the horse, squirrel, sheep, and cattle is 93.45\%, 91.82\%, 90.81\%, and 90.56\% compared to human ACE2, respectively.  These predictions still require more practical research is required to determine the binding affinity of these mammals' ACE2 with SARS-CoV-2's S-protein. It has been shown that ACE2 can tolerate up to seven amino acids changes out of 20 critical ones that contact with the S-protein, without losing the functionality as SARS-CoV-2's receptor~\cite{zhai2020comparison}. This means that
similarity might not be the only factor that influences the binding affinity between ACE2 and SARS-CoV-2's S-protein.

\subsubsection*{SARS-CoV-2 and human interferon pathway interactome prediction\\}
The Interferon (IFN) pathway plays a critical role in the human immune response. After the virus infection is detected, the innate immune system will induce the IFN signaling, and the expression of IFN genes will increase the cellular resistance to viral invasion. Viruses have developed various strategies to inhibit IFN signaling to facilitate a successful viral invasion~\cite{schulz2016viral}. SARS-CoV and MERS-CoV have been studied quite comprehensively in counteracting the IFN signaling responses compared with SARS-CoV-2. From IMSP, eight interactions between three SARS-CoV-2 proteins and human proteins in the innate immune pathway have been identified, shown in Figure~\ref{fig:interactions}(b)-(d). These interactions have a high probability of playing crucial roles in the SARS-CoV-2's suppression of the host's innate immune system.

\ \ \ PLpro was predicted to interact with the nuclear factor kappa-light-chain-enhancer of activated B (NF-$\kappa$B) in humans, shown in Figure~\ref{fig:interactions}(b). In SARS-CoV~\cite{frieman2009severe} and MERS-CoV~\cite{mielech2014mers}, PLpro have been identified as being capable of reducing the NF-$\kappa$B reporter activity and further suppressing IRF3. From our prediction, it might also be highly possible for SARS-CoV-2's PLpro to have similar functions.

\ \ \ ORF6 and nsp15 in SARS-CoV-2 have been discovered as crucial viral interferon antagonists of SARS-CoV-2. These two proteins inhibit the localization of IRF3 by interacting with RIG-I~\cite{yuen2020sars}, as a similar function is found for SARS-CoV ORF6~\cite{kopecky2007severe}, shown in Figure~\ref{fig:interactions}(c)\&(d). ORF6 in SARS-CoV has also been shown to interact with STAT1 and STAT2~\cite{frieman2007severe}. From IMSP's prediction, ORF6 and nsp15 in SARS-CoV-2 were suggested to have potential interactions with MDA5 and MAVS. Since MAVS works as the adaptor molecule for MDA5~\cite{wu2015rig}, it is possible that a viral protein interacts with either one of these two would also have the interaction with the other. Besides, MAVS and MDA5, ORF6 was also predicted to interact with STAT1 and STAT2, the same function as in the SARS-CoV. As nsp15 and ORF6 both function in nuclear transport machinery after the viral entry~\cite{gordon2020sars}, it is reasonable that, for these two proteins, similar interactions with innate immune pathways are predicted. Careful experiments should be conducted to identify nsp15 and ORF6's impact on the innate immune system.

\begin{figure}[!ht]
    \centering
    \includegraphics[width = 8.5cm]{Figures/Cartoon infection final.jpg}
    \caption{\textbf{Infection prediction for SARS-CoV-2.}\textit{This figure shows SARS-CoV-2's known and predicted infected mammalian hosts. The predicted infected mammalian hosts without a predicted or proven corresponding receptor binding relationship with SARS-CoV-2's S-protein were excluded.}}
    \label{fig:cartoon_infection}
\end{figure}

\subsubsection*{SARS-CoV-2 infection prediction\\}
Based on the protein-level interaction prediction, we concluded five predicted infections for SARS-CoV-2 that is highly possible, as these mammals are not only predicted to be infected but also to have a successful SARS-CoV-2 virus binding with the host receptors. As shown in Figure~\ref{fig:cartoon_infection}, these animals include mice, rats, sheep, cattle, and squirrels.

\ \ \ Mice and rats are similar in every protein that we took into consideration in IMSP: all of the proteins have above 85\% similarity, and the receptor ACE2 even has 94\% similarity. Mouse has been identified as a host for all beta-coronaviruses: SASR-CoV~\cite{channappanavar2016dysregulated}, MERS-CoV~\cite{cockrell2016mouse}, HCoV-OC43~\cite{niu2020non} and HCoV-HKU1~\cite{gorse2009human,lim2016human}. SARS-CoV-2 also falls into the category of beta-coronavirus~\cite{lu2020genomic}, which has a high possibility to infect mice and rats.

\ \ \ Cattle are hosts for both MERS-CoV~\cite{kandeil2019middle} and a bovine coronavirus which is exceptionally similar to HCoV-OC43 (over 94\% similarity)~\cite{szczepanski2019canine}. This means that cattle can be the host for other coronaviruses. Cattle, along with sheep and squirrels, are closely related to the human living environment or daily diet. They could be potential mammalian hosts that again transmit the virus back to human society. The investigation of these highly possible infections could help identify the virus's path and further control the transmission SARS-CoV-2 from mammalian hosts. Further research on these potential hosts might be crucial to social health and safety.


\subsection*{Interaction Prediction Evaluation\\}
\begin{table*}[b]
\resizebox{.98\textwidth}{!}{
\scriptsize
\begin{tabular}{c|ccccc}
\hline
Model     & Accuracy       & Weighted Precision & Weighted F1-score & ROC Macro      & ROC Weighted   \\ \hline\hline
GF~\cite{ahmed2013distributed} & 0.885 ± 0.008 & 0.864 ± 0.009 & 0.872 ± 0.008 & 0.913 ± 0.009 & 0.934 ± 0.009\\
Deepwalk~\cite{Perozzi:2014:DOL:2623330.2623732} & 0.895 ± 0.006          & 0.871 ± 0.006          & 0.880 ± 0.005          & 0.906 ± 0.011          & 0.930 ± 0.009          \\
LINE~\cite{tang2015line} & 0.774 ± 0.011           & 0.760 ± 0.012            & 0.765 ± 0.012          & 0.884 ± 0.012            & 0.891 ± 0.010          \\
Node2vec~\cite{grover2016node2vec} & 0.894 ± 0.006          & 0.853 ± 0.009         & 0.868 ± 0.007          & 0.807 ± 0.010           & 0.845 ± 0.010          \\
SDNE~\cite{Wang:2016:SDN:2939672.2939753} & 0.823 ± 0.014          & 0.811 ± 0.015             & 0.815 ± 0.015           & 0.937 ± 0.011          & 0.947 ± 0.007          \\
IMSP  & \textbf{0.949 ± 0.006} & \textbf{0.948 ± 0.006} & \textbf{0.944 ± 0.008} & \textbf{0.994 ± 0.002} & \textbf{0.993 ± 0.002} \\
\hline\hline
\end{tabular}
}
\caption{\textbf{Interaction Prediction Overall Performance Evaluation and Comparison.} \textit{This table gives six evaluation metrics regarding interaction prediction. During performance measurement, we generated two independent sets, training and testing sets, from the network. We set the ratio between the training and the test set to be 8:2. We ensured that the composition of positive interactions in the training set was the same as the test set. To guarantee input balance, we gathered negative interactions (unconnected node pairs) to correspondingly match the number of positive interactions in both training and testing sets. Part of negative samples was extracted from known negative interactions (true negatives), which consisted of spike-receptor interactions that were demonstrated as not existed. The remaining part of negative samples was randomly selected from other unconnected node pairs which were assumed as not existed. To eliminate performance randomness, we executed the comparison algorithm by 30 runs and recorded each run's performance. We generated new training and test sets during each run and evaluated all models on the same test set. Then, we performed two-sample heteroscedastic T-tests for these six overall performance evaluation metrics to test the significance of IMSP's improvement. Lastly, we reported the average with standard deviation.}}
\label{tab:overall_perf}
\end{table*}

\begin{figure*}[!ht]
    \centering
    \includegraphics[width=18cm]{Figures/perf.jpg}
    \caption{\textbf{Performance on PPI and Infection Predictions.} \textit{This figure demonstrates IMSP's performance on PPI and infection predictions in comparison with five other models. The ocean-blue columns represent the performance of our IMSP model derived from the average of 30 independent runs. As a result, the IMSP model achieved 0.808 for the Infection F1-score and 0.910 for the PPI F1-score. Compared to other models, our model outperformed other models in all the evaluation metrics. Specifically, in terms of infection F1-score, our model outperformed the second-best model SDNE~\cite{Wang:2016:SDN:2939672.2939753} by 14.9\%. In terms of PPI F1-score. Our model also surpassed the second-best model Deepwalk~\cite{Perozzi:2014:DOL:2623330.2623732} by 6.6\%. }}
    \medskip
    \label{fig:detaileED_perf}
\end{figure*}

We compared IMSP to five other representative baseline models on our dataset in a cross-validation experiment. We conducted 30 runs in total for the experiment. The baseline models include two famous random-walk based models (Deepwalk~\cite{Perozzi:2014:DOL:2623330.2623732} and Node2vec~\cite{grover2016node2vec}), two neural-network-based models (Large-scale Information Network Embedding (LINE)~\cite{tang2015line} and Structural Deep Network Embedding (SDNE)~\cite{Wang:2016:SDN:2939672.2939753}), and a classical matrix-based model, Graph Factorization (GF)~\cite{ahmed2013distributed}. In each cross-validation run, we reserved 20\% of each interaction type as the positive test set while ensuring the remaining network to be a connected one. To ensure input balance, we collected negative interactions (unconnected node pairs) to match the number of positive interactions in both training and testing sets. Firstly, we extracted known negative interactions as true negatives, which consisted of spike-receptor interactions that were demonstrated as not existed, such as between SARS-CoV-2's S-protein and the host receptor DPP4 (the target host receptor of MERS-CoV). We split known negative interactions by a ratio of 8:2 to add to the training and testing sets. Then, since we still lack some number of negative interactions in both sets, we randomly selected unconnected node pairs as negative interactions to add to the training and testing sets. In this way, we confirmed that the training set's composition was the same as the test set's composition in terms of interaction type. We then applied IMSP and other models on the training set to perform network learning on the same training set and measure the test set's interaction prediction performances. We repeated the entire measurement process for 30 runs to minimize the measurement randomness. Finally, we performed two-sample heteroscedastic T-test at the 0.01 significance level to test the significance of our model's performance improvement compared to other models.

\ \ \ Table~\ref{tab:overall_perf} shows the performance comparison, measured in several commonly used evaluation metrics, between IMSP and other existing tools. IMSP achieved an overall interaction prediction accuracy of 94.9\% with a standard deviation of 0.004, which demonstrated a 6.0\% gain compared to the second-best model. Our model also excelled in its weighted F1-score, achieving 0.944 with a standard deviation of 0.008, which transcended the second-best model by 7.3\%. The p-values for these two metrics were all smaller than 0.01, which indicated significant improvement for our model. In terms of the performance on specific types of interactions, i.e., infection and PPI predictions (See Fig.~\ref{fig:detaileED_perf}), IMSP achieved an F1-score of 0.808 with 0.119 standard deviation for infection predictions, a 14.9\% increase compared to the second-best model. The p-value was also smaller than 0.01, indicating a significant improvement in our model. For PPI predictions, our model achieved an F1-score of 0.909 with 0.029 standard deviation, a 6.6\% increment compared to the second-best tool. The p-value also demonstrated a significant improvement for IMSP. In conclusion, our model showed statistically significant improvements compared to all existing models in all 12 evaluation metrics.

\ \ \ IMSP's high performance might result from its ability to take full advantage of well-studied knowledge and data from previous biology research with protein-level variations being considered. Thanks to our virus-host interaction network's novel design, such cross-organism information and multi-class linkage information can be well-preserved. Another reason behind IMSP's performance improvement is that it factors essential biological metadata as textualized features for nodes. In this way, two connected nodes' vectorized content substantially helps the classifier output a correct predicted class when formulating interaction representations. However, around 10\% PPI predictions were unlikely by our definition, such as PPIs between S-protein and non-receptor proteins. To minimize unlikely predictions, we incorporated the node's function and existing biological metadata as part of the node feature. This change significantly reduced unlikely PPI predictions to around 2\%. We further pushed the boundaries by utilizing known negative interactions in the protein layer to constitute part of the training and testing sets—this finally reduced the unlikely PPI predictions to less than 1\%.

\ \ \ In conclusion, IMSP exhibited robust and stable performance in both top-level and detailed evaluation metrics, which is substantially improved compared to existing tools. When analyzing newly emerged viruses with limited available information, namely SARS-CoV-2, IMSP can provide reasonable and reliable predictions.

% -----------------------------------------------------------
% ---------------------discussion section--------------------
% -----------------------------------------------------------

\section*{Discussion}
This study assembled 246 nodes (viruses, virus proteins, hosts, and host proteins) and 1,766 known interactions among these nodes, including four types of interactions: similarities, infections, interactions, and belongings. Two main parts for virus-host protein-level interactions were taken into consideration: viral entry and innate immune pathway interactions. Accompanied by these two parts, we predicted the potential host for viruses and undiscovered novel PPIs. Among seven human coronaviruses, SARS-CoV and MERS-CoV were well studied about their binding with the receptors and interacting with the innate immune pathway, whereas HCoV-OC43, HCoV-NL63, HCov-HKU1, HCoV-229E, and newly emerged SARS-CoV-2 remained relatively unknown. Based on protein similarity and known interactions, 322 PPIs were predicted with this model. The actual PPIs need to be investigated further based on the prediction result.

\ \ \ Established discoveries about SARS-CoV-2's viral interaction with host proteins are scarce. SARS-CoV-2 has been highly suspected of suppressing the innate immune response and reducing the production of IFN. In this way, IMSP's findings could help discover the relations between virus invasion and host response and therefore provide clues to developing therapeutic strategies for the treatment of this disease.

\ \ \ More broadly, IMSP could be applied to any other analysis of the virus-host interaction network predictions. This network would be built based on the information of the protein-protein relations, protein-host relations, and host-host relations. Based on these attributes in the network and currently known relationships between some of the proteins, IMSP could predict the high-possibility interactions. We hope to use this pipeline as a tool to build a guideline for investigating various similar viruses and their mechanisms with hosts.

% -----------------------------------------------------------
% -----------------methods/materials section-----------------
% -----------------------------------------------------------

\section*{Methods}
\subsection*{Infection Mechanism and Spectrum Prediction\\}
Our IMSP model requests three different inputs, including pairwise similarity matrices for all host proteins and virus proteins, a set of known PPIs and infection relationships, and proteins' biological metadata. For instance, appropriate protein metadata can include a protein's name, layer, host, and function. Given these three inputs, the model constructs a virus-host interaction network. Then, in the network learning phase, IMSP learns and combines the structural information and the content information of interactions to represent the heterogeneous network comprehensively. Lastly, in the prediction phase, IMSP trains a neural-network-based classifier, Multi-layer Perceptron classifier, along with strict filtering procedures, to output high-possibility undiscovered interactions. Specifically, our model generated PPI and infection predictions with each prediction's likelihood. In the following, we elaborated on the details of our IMSP model's two main steps in terms of network construction and learning, and interaction prediction with result validation.

\subsection*{Virus-host interaction network construction and learning\\}
\begin{table}[!t]
\scriptsize
\centering
\begin{tabular}{c|c}
\hline
Notion     & Description\\ \hline
$V_i$ & Node $i$ in the network\\
$V$ & The set of all Nodes\\
$I_{i,j}$ & Edge (Interaction) between Node $i$ and Node $j$\\
$E$ & The set of all Edges (Interactions) in the network\\
$R^{S}_i$ & Structure embedding vector for Node $i$\\
$R^{C}_i$ & Content embedding vector for Node $i$\\
$CE_{i,j}$ & Content embedding vector for Interaction $I_{i,j}$\\
$IE_{i, j}$ & Full Interaction embedding vector for Interaction $I_{i,j}$\\
$W_{i, j}$ & Edge (Interaction) weight between Node $i$ and Node $j$\\
$ED_{i, j}$ & Euclidean distance between Node $i$ and Node $j$\\
$MD(R^{C}_i, R^{C}_j)$ & Magnitude difference between Vector $R^{C}_i$ and $R^{C}_j$\\
$TS\textit{-}SS_{i, j}$ & TS-SS similarity between Vector $R^{C}_i$ and $R^{C}_j$\\
\hline
\end{tabular}
\label{tab:notation table}
\caption{\textbf{Notations.}}
\end{table}

We utilized layers and groups to mark and separate different nodes during virus-host interaction network construction, and used interactions to represent PPI/infection/similarity/belonging relationship between nodes. For modeling the network, we constructed a weighted and undirected heterogeneous network using NetworkX~\cite{SciPyProceedings_11}. The network contained four groups of nodes: host, host proteins, virus proteins, and virus. We organize the virus group and the host group into the same layer, i.e., the organism layer, and we put two groups of proteins into the protein layer. By nature, the network contained four types of interactions: PPI (between virus proteins groups and host proteins groups), infection (between viruses group and hosts group), similarity (between homologs of proteins) and belonging (between organism layer and protein layer). Similarity and belonging relationships were innately connected. Other types of interactions were connected based on proven molecular level knowledge from existing research~\cite{channappanavar2016dysregulated, frieman2008sars, matthews2014sars, totura2012sars, hu2017sars, chen2014sars, li2010host, li2003angiotensin, lei2009mavs,szczepanski2019canine,niu2020non,hofmann2005human,haagmans2014middle, mou2013receptor, kandeil2019middle, bailey2014crystal, lui2016middle}. The virus-host interaction network comprised of 246 nodes and 1,766 interactions. After the network structure was built, we used $W_{i, j}$ to represent the biological closeness between $V_i$ and $V_j$. Specifically, if $I_{i, j}$ denoted the relationship between two protein homologs, we used the full-sequence similarity between $V_i$ and $V_j$ from pairwise similarity matrices to represent $W_{i, j}$. For other types of relationship, we obtained the embeddings of node content, denoted as $R^{C}_i$ for $V_i$ by passing textualized node content to the Word2vec model~\cite{mikolov2013efficient}, a famous and powerful neural-network-based word embedding model. Then, we utilized the \textit{TS-SS} similarity metric~\cite{7474366}, a robust and reliable similarity measurement in the field of textual mining, to calculate the similarity between $R^{C}_i$ and $R^{C}_j$ to assign to $W_{i, j}$. Precisely, for interactions other than similarity relations, we calculated the $TS\textit{-}SS_{i, j}$ for all $i$ and $j$ if $I_{i, j} \subset E$ as the following:
\begin{equation}
\begin{split}
TS\textit{-}SS_{i, j} =&\ (|R^{C}_i|\cdot|R^{C}_j|\cdot sin(\theta')\cdot \theta'\cdot \pi \cdot \\
 & (ED(R^{C}_i, R^{C}_j) \ + MD(R^{C}_i, R^{C}_j))^2/720,
\end{split}
\end{equation}
where $MD(R^{C}_i, R^{C}_j)$~\cite{7474366} is defined as the magnitude difference between $R^{C}_i$ and $R^{C}_j$, which is calculated as:
\begin{equation}
MD(R^{C}_i, R^{C}_j) = \left |\ \ \sqrt{\sum_{n=1}^{\dim{R^{C}_i}} {R^{C}_i}^2} - \sqrt{\sum_{n=1}^{\dim{R^{C}_j}} {R^{C}_j}^2}\ \  \right   |,
\end{equation}
and $\theta'$ is defined as:
\begin{equation}
\theta' = \cos^{-1}(\cos({R^{C}_i}, {R^{C}_j})) + 10.
\end{equation}
Note that $\theta'$ is increased by 10 degrees to overcome the problem of overlapping vectors. Then, $W_{i, j}$ is calculated as:
\begin{equation}
W_{i, j} = \sigma(TS\textit{-}SS_{i, j}/\overline{TS\textit{-}SS}),
\end{equation}
where $\sigma$ is the sigmoid function, and $\overline{TS\textit{-}SS}$ denotes the average of $TS\textit{-}SS_{i, j}$, for all $i,j$, if $i \neq j$ and $V_i, V_j \subset V$.

\ \ \ For network learning, we captured the network's heterogeneity by adding the network's heterogeneous content information to its structural information. Specifically, we performed network structural embedding assuming the network is homogeneous. Then, we added the content embedding on top of structural embedding to model its heterogeneity.

\ \ \ Firstly, for network structural embedding, we used a powerful network feature learning model, Node2vec~\cite{grover2016node2vec}, to learn the structural embedding for nodes. Node2vec is a state-of-the-art model for homogeneous network embedding. We took full advantage of Node2vec's biased searching algorithm during our application. Precisely, the Node2vec model performed a fixed-length random walk for sampling, which takes weight into account. Let $c_m$ denote the \mbox{$m$-th} node in walk with $c_0$ denoting the starting node of the current random walk. Nodes $c_m$ are generated by the following distribution:
\begin{equation}
P(c_m = V_i\ |\ c_{m-1} = V_j) = \left\{\begin{matrix}
\pi_{V_j, V_i}/Z & \textup{if}\ \ I_{i, j} \subset E\\ 
0 & \textup{otherwise}
\end{matrix}\right.,
\end{equation}
where $m \geqslant 1$, Z is the normalizing constant, and $\pi_{V_j, V_i}$ is the unnormalized transition probability between $V_i$ and $V_j$, which is calculated as $\pi_{V_j, V_i} = \alpha_{pq}(V_t, V_i) \cdot W_{i, j}$. $\alpha_{pq}(V_t, V_i)$, termed as search bias, is calculated as:
\begin{equation}\label{eq:pq}
\alpha_{pq}(V_t, V_i)) = \left\{\begin{matrix}
1/p & \textup{if}\ \ d_{V_t, V_i} = 0\\ 
1 & \textup{if}\ \ d_{V_t, V_i} = 1\\ 
1/q & \textup{if}\ \ d_{V_t, V_i} = 2
\end{matrix}\right.,
\end{equation}

where $d_{V_t, V_i}$ denotes the shortest path between $V_i$ and $V_j$, assuming that we have just traversed from $V_t$ to $V_i$ and are now evaluating the transition probability leaving $V_i$. 

\ \ \ In ~\eqref{eq:pq}, p and q are Node2vec's hyperparameters that can be adjusted to influence the probability to go back to $V_i$ after visiting $V_j$ and the probability to explore undiscovered components of the network. In this way, we were able to fine-tune the structural embedding model, Node2vec, to generate a conclusive embedding for our network.

\ \ \ Secondly, to create content representations, $CE_{i, j}$ for all $I_{i, j}$ in the network, we combined the textualized node content of $V_i$ and $V_j$ along with expected interaction type such as interacts/infects/similar/belongs, and input them into the Word2vec model~\cite{mikolov2013efficient} to generate vectorized interaction representations. Then, the full interaction representations for interaction $I_{i, j}$ are formulated as follows:
\begin{equation}
\begin{split}
IE_{i, j} &= [R^{S}_i\:,\:R^{S}_j,\:CE_{i,j}], \\ 
IE_{j, i} &= [R^{S}_j\:,\:R^{S}_i,\:CE_{j, i}].
\end{split}
\end{equation}
\ \ \ Note that by Word2vec's nature, the input documents' order does not affect its output, meaning that $CE_{i, j}$ is the same as $CE_{j, i}$. By finishing this step, we obtained all interaction representations, $IE_{i, j}$, for all $V_i$ and $V_j$ $\subset V$ and $i \neq j$.

\subsection*{Virus-host interaction prediction\\}
In the interaction prediction phase, we utilized a classifier to perform a multi-class classification task on the interaction representations to learn from the existing network and suggest potential undiscovered interactions. Specifically, for all $I_{i, j} \not\subset E$, we classified $IE_{i,j}$ into three classes, i.e. PPI/infections/no-interaction. Scikit-learn's Multi-layer Perceptron (MLP) classification model~\cite{sklearn_api} was used for this multi-class interaction classification. 

\ \ \ We conducted cross-validation experiments to fine-tune the model, where 20\% of each interaction type was reserved as a positive test set. In this process, we also balanced the test set by inserting an equal number of negative interactions containing both known negatives and unknown negatives. Specifically, known negative interactions (true negatives) were generated from proven unconnected node pairs, such as the interaction between MERS-CoV's S-protein and mammalian hosts' ACE2 receptors. Unknown negative interactions were randomly sampled from the network, which was assumed as not existing. Then, by using the fine-tuned hyperparameters from cross-validation experiments, all positive interactions with an equal number of negative interactions were used for training the classifier.  IMSP later took all other unused unconnected node pairs as input data and classified them into the class of PPI/infection/no interaction.

\ \ \ During prediction result correction, we defined rules from both computational and biological perspectives to mark and remove unlikely predictions. Computationally, because there exist two representations for $I_{i, j}$, i.e., $IE_{i, j}$ and $IE_{j, i}$, we defined the prediction for $I_{i, j}$ as a strong one if and only if $IE_{i, j}$ and $IE_{j, i}$ were all classified into the same type of interaction other than the no-interaction. Biologically, we assumed that each virus's S-protein only interacts/binds with its target receptor. Thus, all other interactions with S-protein are considered unlikely. For example, the predicted interaction, if any, between SARS-CoV-2's S-protein and DPP4 (the target receptor for MERS-CoV's S-protein) would be eliminated. Thirdly, we also defined that an infection interaction was likely only when the corresponding interaction between the virus' S-protein and an appropriate host receptor was predicted. In this way, we ensured that all the remaining predictions are both computationally and biologically meaningful and practically useful to guide further research.

\subsection*{Resource Availability\\}
All data and codes are available at Github repositories: \\
IMSP model, its predictions and performance evaluations can be found at \\
\href{IMSP}{https://github.com/SupremeEthan/IMSP}; \\
Data and data parsing code can be found at \\ \href{IMSP-Płrser}{https://github.com/SupremeEthan/IMSP-Parser}.

%\section*{Future work}
%The first possible limitation in IMSP is that we did not include the gene set enrichment analysis as part of the model. The model could potentially be improved by adding the proteins sub-cellular locations into the embedding result. Another thing that may influence the prediction performance is that IMSP did not consider the protein's motif similarity as the similarity data we used is the overall similarity of proteins. Besides, the motif may also be taken into account for further research. Other than that, the prediction accuracy needs to be determined experimentally. In our classification model, over 85\% of samples in the negative set are randomly selected from unconnected node pairs, which have the possibility of being undetected true positives. However, since these highly possible interactions reside in the negative set, they will be unlikely to be predicted as connected.

\section*{Acknowledgement}
This work was partially supported NSF OAC 1920147.

\section*{Authors' Contributions}
Conceptualization, P.H. and H.L.; Methodology, H.D., F.C., and H.L.; Software, H.D.; Formal Analysis: H.D.; Investigation, F.C. and H.D.; Writing – Original Draft, F.C. and H.D.; Writing – Review \& Editing, all authors; Visualization: F.C. and H.D.; Supervision, H.L. and P.H.

\section*{Lead Contact}

\section*{Declaration of Interests}
The authors declare no competing interests.

\section*{References}
\bibliography{ref}

% Cell Press credit taxonomy: https://els-jbs-prod-cdn.jbs.elsevierhealth.com/pb/assets/raw/shared/guidelines/CRediT-taxonomy.pdf


% \onecolumn
% \newpage

%%%%%%%%%%%%%%%%%%%%%%%%%%%%%
% Supplementary Information %
%%%%%%%%%%%%%%%%%%%%%%%%%%%%%
% \section{Full Map of Binding Interactions between Coronaviruses' S-protein and mammalian hosts' Receptor}

% \begin{figure}
%     \centering
%     \includegraphics[width=1.0\textwidth]{Figures/spike_binding 10.10.jpeg}
%     \caption{\textbf{Virus entry: binding relationships between the S-proteins of human coronaviruses and the ACE2 receptors of mammalian hosts} \textit{This figure of network is visualized by Cytoscape.Virus Protein Layer nodes are represented in green ellipses, and Host Protein Layer nodes are represented in blue ellipses. The original interactions are represented in grey lines, including similarity among nodes in the same type and the known receptor bindings between viruses spike and mammalian hosts ACE2. Predicted receptor bindings are represented in light purple lines. As the infection relations have been checked for likelihood in the IMSP model, all predicted interactions are strong predicted interactions. All host names, except Homo sapiens, are displayed in their short form: M.M. is Mus musculus; F.C. is Felis catus; C.L.F. is Canis lupus familiaris; O.A. is Ovis aries; R.N. is Rattus norvegicus; Ma.M. is Macaca mulatta; R.F. is Rhinolophus ferrumequinum; M.A. is Mesocricetus auratus; C.D. is Camelus dromedarius; B.T. is Bos taurus; G.G. is Gallus gallus; I.T. is Ictidomys tridecemlineatus; E.C. is Equus caballus; S.S.D. is Sus scrofa domesticus; P.T. is Pan troglodytes; O.C. is Oryctolagus cuniculus.}}
%     \label{fig:viral entry graph}
% \end{figure}
% \newpage

% \section{Full Map of Protein-Protein Interactions related to IFN signaling Pathway between Coronaviruses and mammalian hosts}
% \begin{figure}[!ht]
%     \centering
%     \begin{turn}{90}
%     \includegraphics[width=1.22\textwidth]{Figures/IFN_pathway 10.7.jpeg}
%     \end{turn}
%     \caption{\textbf{IFN interactions: virus proteins interactions with IFN signaling pathway to suppress the IFN signaling} \textit{Same representations for nodes and interactions as described in figure1. The predicted interactions are represented in light purple lines. The solid lines stand for strong predictions, and dotted lines stand for weak predictions as defined in the IMSP model.}}
%     \label{fig:IFN graph}
% \end{figure}
% \newpage

% \section{Full Map of Infection Relationships between Coronaviruses and mammalian hosts}
% \begin{figure}[!ht]
%     \centering
%     \includegraphics[width=1.0\textwidth]{Figures/infection_final 10.7.jpeg}
%     \caption{\textbf{Coronaviruses and mammalian hosts infection relationships} \textit{Virus Layer nodes are represented in green rhombi, and Host Layer nodes are represented in blue rhombi. The original infection interactions are represented in grey lines. Predicted infection relations are represented in light purple lines.The solid lines stand for strong predictions and dotted lines stand for weak predictions as defined in the IMSP model.}}
%     \label{fig:infection graph}
% \end{figure}


% \newpage
% \section{Node ID associated with Node Full Name in the Network}
% \scriptsize
% \begin{longtable}[c]{|c|c|c|c|}
% \hline
% \textbf{Node ID} & \textbf{Full name representation} & \textbf{Node ID} & \textbf{Full name representation} \\ \hline
% \endfirsthead
% %
% \multicolumn{4}{c}%
% {{\bfseries Table \thetable\ continued from previous page}} \\
% \hline
% \textbf{Node ID} & \textbf{Full name representation} & \textbf{Node ID} & \textbf{Full name representation} \\ \hline
% \endhead
% %
% 0 & nsp15, Severe acute respiratory syndrome coronavirus 2 & 123 & IRF7, Rattus norvegicus \\ \hline
% 1 & nsp15, Severe acute respiratory syndrome-related coronavirus & 124 & IRF7, Mesocricetus auratus \\ \hline
% 2 & nsp15, Human coronavirus HKU1 & 125 & IRF7, Ovis aries \\ \hline
% 3 & STAT1, Homo sapiens & 126 & IRF7, Mus musculus \\ \hline
% 4 & STAT1, Pan troglodytes & 127 & IRF7, Bos taurus \\ \hline
% 5 & STAT1, Macaca mulatta & 128 & IRF7, Gallus gallus \\ \hline
% 6 & STAT1, Felis catus & 129 & MDA5, Homo sapiens \\ \hline
% 7 & STAT1, Camelus dromedarius & 130 & MDA5, Pan troglodytes \\ \hline
% 8 & STAT1, Equus caballus & 131 & MDA5, Macaca mulatta \\ \hline
% 9 & STAT1, Canis lupus familiaris & 132 & MDA5, Equus caballus \\ \hline
% 10 & STAT1, Rhinolophus ferrumequinum & 133 & MDA5, Ictidomys tridecemlineatus \\ \hline
% 11 & STAT1, Bos taurus & 134 & MDA5, Felis catus \\ \hline
% 12 & STAT1, Ovis aries & 135 & MDA5, Bos taurus \\ \hline
% 13 & STAT1, Ictidomys tridecemlineatus & 136 & MDA5, Mus musculus \\ \hline
% 14 & STAT1, Rattus norvegicus & 137 & MDA5, Rattus norvegicus \\ \hline
% 15 & STAT1, Mus musculus & 138 & MDA5, Mesocricetus auratus \\ \hline
% 16 & STAT1, Mesocricetus auratus & 139 & MDA5, Canis lupus familiaris \\ \hline
% 17 & STAT1, Gallus gallus & 140 & MDA5, Gallus gallus \\ \hline
% 18 & IRF9, Homo sapiens & 141 & PRKRA, Homo sapiens \\ \hline
% 19 & IRF9, Macaca mulatta & 142 & PRKRA, Macaca mulatta \\ \hline
% 20 & IRF9, Pan troglodytes & 143 & PRKRA, Equus caballus \\ \hline
% 21 & IRF9, Felis catus & 144 & PRKRA, Canis lupus familiaris \\ \hline
% 22 & IRF9, Camelus dromedarius & 145 & PRKRA, Camelus dromedarius \\ \hline
% 23 & IRF9, Sus scrofa domesticus & 146 & PRKRA, Mesocricetus auratus \\ \hline
% 24 & IRF9, Equus caballus & 147 & PRKRA, Mus musculus \\ \hline
% 25 & IRF9, Rhinolophus ferrumequinum & 148 & PRKRA, Ictidomys tridecemlineatus \\ \hline
% 26 & IRF9, Ovis aries & 149 & PRKRA, Ovis aries \\ \hline
% 27 & IRF9, Canis lupus familiaris & 150 & PRKRA, Bos taurus \\ \hline
% 28 & IRF9, Bos taurus & 151 & PRKRA, Rattus norvegicus \\ \hline
% 29 & IRF9, Ictidomys tridecemlineatus & 152 & PRKRA, Rhinolophus ferrumequinum \\ \hline
% 30 & IRF9, Mus musculus & 153 & PRKRA, Pan troglodytes \\ \hline
% 31 & IRF9, Mesocricetus auratus & 154 & ORF3b, Severe acute respiratory syndrome-related coronavirus \\ \hline
% 32 & IRF9, Rattus norvegicus & 155 & ORF3b, Human coronavirus NL63 \\ \hline
% 33 & IRF9, Gallus gallus & 156 & DPP4, Homo sapiens \\ \hline
% 34 & RIG-I, Homo sapiens & 157 & DPP4, Pan troglodytes \\ \hline
% 35 & RIG-I, Pan troglodytes & 158 & DPP4, Macaca mulatta \\ \hline
% 36 & RIG-I, Macaca mulatta & 159 & DPP4, Ovis aries \\ \hline
% 37 & RIG-I, Rhinolophus ferrumequinum & 160 & DPP4, Ictidomys tridecemlineatus \\ \hline
% 38 & RIG-I, Canis lupus familiaris & 161 & DPP4, Bos taurus \\ \hline
% 39 & RIG-I, Mesocricetus auratus & 162 & DPP4, Felis catus \\ \hline
% 40 & RIG-I, Mus musculus & 163 & DPP4, Equus caballus \\ \hline
% 41 & RIG-I, Gallus gallus & 164 & DPP4, Rhinolophus ferrumequinum \\ \hline
% 42 & ORF4b, Middle East respiratory syndrome-related coronavirus & 165 & DPP4, Mesocricetus auratus \\ \hline
% 43 & ORF4b, Human coronavirus 229E & 166 & DPP4, Rattus norvegicus \\ \hline
% 44 & nsp1, Severe acute respiratory syndrome coronavirus 2 & 167 & DPP4, Mus musculus \\ \hline
% 45 & nsp1, Severe acute respiratory syndrome-related coronavirus & 168 & DPP4, Camelus dromedarius \\ \hline
% 46 & nsp1, Middle East respiratory syndrome-related coronavirus & 169 & DPP4, Gallus gallus \\ \hline
% 47 & nsp1, Human coronavirus HKU1 & 170 & ORF6, Severe acute respiratory syndrome-related coronavirus \\ \hline
% 48 & spike, Human coronavirus OC43 & 171 & ORF6, Severe acute respiratory syndrome coronavirus 2 \\ \hline
% 49 & spike, Human coronavirus HKU1 & 172 & STAT2, Homo sapiens \\ \hline
% 50 & spike, Middle East respiratory syndrome-related coronavirus & 173 & STAT2, Pan troglodytes \\ \hline
% 51 & spike, Severe acute respiratory syndrome coronavirus 2 & 174 & STAT2, Macaca mulatta \\ \hline
% 52 & spike, Severe acute respiratory syndrome-related coronavirus & 175 & STAT2, Felis catus \\ \hline
% 53 & spike, Human coronavirus NL63 & 176 & STAT2, Canis lupus familiaris \\ \hline
% 54 & spike, Human coronavirus 229E & 177 & STAT2, Camelus dromedarius \\ \hline
% 55 & IRF3, Homo sapiens & 178 & STAT2, Rhinolophus ferrumequinum \\ \hline
% 56 & IRF3, Pan troglodytes & 179 & STAT2, Equus caballus \\ \hline
% 57 & IRF3, Macaca mulatta & 180 & STAT2, Ovis aries \\ \hline
% 58 & IRF3, Ictidomys tridecemlineatus & 181 & STAT2, Bos taurus \\ \hline
% 59 & IRF3, Rhinolophus ferrumequinum & 182 & STAT2, Mesocricetus auratus \\ \hline
% 60 & IRF3, Camelus dromedarius & 183 & STAT2, Rattus norvegicus \\ \hline
% 61 & IRF3, Ovis aries & 184 & STAT2, Ictidomys tridecemlineatus \\ \hline
% 62 & IRF3, Bos taurus & 185 & STAT2, Mus musculus \\ \hline
% 63 & IRF3, Equus caballus & 186 & STAT2, Gallus gallus \\ \hline
% 64 & IRF3, Rattus norvegicus & 187 & PLpro, Middle East respiratory syndrome-related coronavirus \\ \hline
% 65 & IRF3, Mesocricetus auratus & 188 & PLpro, Severe acute respiratory syndrome-related coronavirus \\ \hline
% 66 & IRF3, Mus musculus & 189 & PLpro, Severe acute respiratory syndrome coronavirus 2 \\ \hline
% 67 & IRF3, Canis lupus familiaris & 190 & PLpro, Human coronavirus OC43 \\ \hline
% 68 & IRF3, Gallus gallus & 191 & PLpro, Human coronavirus HKU1 \\ \hline
% 69 & N protein, Middle East respiratory syndrome-related coronavirus & 192 & Homo sapiens \\ \hline
% 70 & N protein, Severe acute respiratory syndrome coronavirus 2 & 193 & Mus musculus \\ \hline
% 71 & N protein, Severe acute respiratory syndrome-related coronavirus & 194 & Rattus norvegicus \\ \hline
% 72 & N protein, Human coronavirus HKU1 & 195 & Canis lupus familiaris \\ \hline
% 73 & N protein, Human coronavirus OC43 & 196 & Camelus dromedarius \\ \hline
% 74 & N protein, Human coronavirus 229E & 197 & Felis catus \\ \hline
% 75 & N protein, Human coronavirus NL63 & 198 & Ictidomys tridecemlineatus \\ \hline
% 76 & Human coronavirus OC43 & 199 & Bos taurus \\ \hline
% 77 & Human coronavirus HKU1 & 200 & Pan troglodytes \\ \hline
% 78 & Middle East respiratory syndrome-related coronavirus & 201 & Gallus gallus \\ \hline
% 79 & Severe acute respiratory syndrome coronavirus 2 & 202 & Oryctolagus cuniculus \\ \hline
% 80 & Severe acute respiratory syndrome-related coronavirus & 203 & Equus caballus \\ \hline
% 81 & Human coronavirus NL63 & 204 & Macaca mulatta \\ \hline
% 82 & Human coronavirus 229E & 205 & Ovis aries \\ \hline
% 83 & ACE2, Homo sapiens & 206 & Sus scrofa domesticus \\ \hline
% 84 & ACE2, Pan troglodytes & 207 & Rhinolophus ferrumequinum \\ \hline
% 85 & ACE2, Macaca mulatta & 208 & Mesocricetus auratus \\ \hline
% 86 & ACE2, Ictidomys tridecemlineatus & 209 & M protein, Middle East respiratory syndrome-related coronavirus \\ \hline
% 87 & ACE2, Oryctolagus cuniculus & 210 & M protein, Human coronavirus HKU1 \\ \hline
% 88 & ACE2, Equus caballus & 211 & M protein, Severe acute respiratory syndrome-related coronavirus \\ \hline
% 89 & ACE2, Felis catus & 212 & M protein, Severe acute respiratory syndrome coronavirus 2 \\ \hline
% 90 & ACE2, Camelus dromedarius & 213 & M protein, Human coronavirus OC43 \\ \hline
% 91 & ACE2, Mesocricetus auratus & 214 & M protein, Human coronavirus NL63 \\ \hline
% 92 & ACE2, Sus scrofa domesticus & 215 & M protein, Human coronavirus 229E \\ \hline
% 93 & ACE2, Ovis aries & 216 & TBK1, Homo sapiens \\ \hline
% 94 & ACE2, Mus musculus & 217 & TBK1, Pan troglodytes \\ \hline
% 95 & ACE2, Bos taurus & 218 & TBK1, Macaca mulatta \\ \hline
% 96 & ACE2, Rattus norvegicus & 219 & TBK1, Ictidomys tridecemlineatus \\ \hline
% 97 & ACE2, Rhinolophus ferrumequinum & 220 & TBK1, Felis catus \\ \hline
% 98 & ACE2, Canis lupus familiaris & 221 & TBK1, Bos taurus \\ \hline
% 99 & ACE2, Gallus gallus & 222 & TBK1, Rhinolophus ferrumequinum \\ \hline
% 100 & ORF4a, Middle East respiratory syndrome-related coronavirus & 223 & TBK1, Ovis aries \\ \hline
% 101 & ORF4a, Human coronavirus 229E & 224 & TBK1, Canis lupus familiaris \\ \hline
% 102 & NF-$\kappa$B, Homo sapiens & 225 & TBK1, Equus caballus \\ \hline
% 103 & NF-$\kappa$B, Pan troglodytes & 226 & TBK1, Mesocricetus auratus \\ \hline
% 104 & NF-$\kappa$B, Macaca mulatta & 227 & TBK1, Mus musculus \\ \hline
% 105 & NF-$\kappa$B, Bos taurus & 228 & TBK1, Camelus dromedarius \\ \hline
% 106 & NF-$\kappa$B, Rhinolophus ferrumequinum & 229 & TBK1, Rattus norvegicus \\ \hline
% 107 & NF-$\kappa$B, Ovis aries & 230 & TBK1, Gallus gallus \\ \hline
% 108 & NF-$\kappa$B, Ictidomys tridecemlineatus & 231 & MAVS, Homo sapiens \\ \hline
% 109 & NF-$\kappa$B, Mus musculus & 232 & MAVS, Pan troglodytes \\ \hline
% 110 & NF-$\kappa$B, Mesocricetus auratus & 233 & MAVS, Macaca mulatta \\ \hline
% 111 & NF-$\kappa$B, Rattus norvegicus & 234 & MAVS, Ovis aries \\ \hline
% 112 & NF-$\kappa$B, Equus caballus & 235 & MAVS, Equus caballus \\ \hline
% 113 & NF-$\kappa$B, Gallus gallus & 236 & MAVS, Bos taurus \\ \hline
% 114 & NF-$\kappa$B, Camelus dromedarius & 237 & MAVS, Rhinolophus ferrumequinum \\ \hline
% 115 & IRF7, Homo sapiens & 238 & MAVS, Felis catus \\ \hline
% 116 & IRF7, Pan troglodytes & 239 & MAVS, Camelus dromedarius \\ \hline
% 117 & IRF7, Macaca mulatta & 240 & MAVS, Ictidomys tridecemlineatus \\ \hline
% 118 & IRF7, Ictidomys tridecemlineatus & 241 & MAVS, Canis lupus familiaris \\ \hline
% 119 & IRF7, Equus caballus & 242 & MAVS, Mesocricetus auratus \\ \hline
% 120 & IRF7, Rhinolophus ferrumequinum & 243 & MAVS, Mus musculus \\ \hline
% 121 & IRF7, Felis catus & 244 & MAVS, Rattus norvegicus \\ \hline
% 122 & IRF7, Camelus dromedarius & 245 & MAVS, Gallus gallus \\ \hline
% \end{longtable}
% \label{tab:node table}
% \newpage


% \section{Original Network's Interaction Table with Interaction Types}
% \begin{longtable}[c]{|c|c|c|c|c|}
% \hline
% \textbf{Source ID} & \textbf{Source Name} & \textbf{Target ID} & \textbf{Target Name} & \textbf{Relation} \\ \hline
% \endfirsthead
% %
% \multicolumn{5}{c}%
% {{\bfseries Table \thetable\ continued from previous page}} \\
% \hline
% \textbf{Source ID} & \textbf{Source Name} & \textbf{Target ID} & \textbf{Target Name} & \textbf{Relation} \\ \hline
% \endhead
% %
% 0 & nsp15  Severe acute respiratory syndrome coronavirus 2 & 1 & nsp15  Severe acute respiratory syndrome-related coronavirus & similar \\ \hline
% 0 & nsp15  Severe acute respiratory syndrome coronavirus 2 & 2 & nsp15  Human coronavirus HKU1 & similar \\ \hline
% 0 & nsp15  Severe acute respiratory syndrome coronavirus 2 & 79 & Severe acute respiratory syndrome coronavirus 2 & belongs \\ \hline
% 0 & nsp15  Severe acute respiratory syndrome coronavirus 2 & 55 & IRF3  Homo sapiens & interacts \\ \hline
% 0 & nsp15  Severe acute respiratory syndrome coronavirus 2 & 65 & IRF3  Mesocricetus auratus & interacts \\ \hline
% 0 & nsp15  Severe acute respiratory syndrome coronavirus 2 & 66 & IRF3  Mus musculus & interacts \\ \hline
% 0 & nsp15  Severe acute respiratory syndrome coronavirus 2 & 67 & IRF3  Canis lupus familiaris & interacts \\ \hline
% 0 & nsp15  Severe acute respiratory syndrome coronavirus 2 & 34 & RIG-I  Homo sapiens & interacts \\ \hline
% 0 & nsp15  Severe acute respiratory syndrome coronavirus 2 & 38 & RIG-I  Canis lupus familiaris & interacts \\ \hline
% 0 & nsp15  Severe acute respiratory syndrome coronavirus 2 & 39 & RIG-I  Mesocricetus auratus & interacts \\ \hline
% 0 & nsp15  Severe acute respiratory syndrome coronavirus 2 & 40 & RIG-I  Mus musculus & interacts \\ \hline
% 1 & nsp15  Severe acute respiratory syndrome-related coronavirus & 2 & nsp15  Human coronavirus HKU1 & similar \\ \hline
% 1 & nsp15  Severe acute respiratory syndrome-related coronavirus & 80 & Severe acute respiratory syndrome-related coronavirus & belongs \\ \hline
% 1 & nsp15  Severe acute respiratory syndrome-related coronavirus & 55 & IRF3  Homo sapiens & interacts \\ \hline
% 1 & nsp15  Severe acute respiratory syndrome-related coronavirus & 59 & IRF3  Rhinolophus ferrumequinum & interacts \\ \hline
% 1 & nsp15  Severe acute respiratory syndrome-related coronavirus & 231 & MAVS  Homo sapiens & interacts \\ \hline
% 1 & nsp15  Severe acute respiratory syndrome-related coronavirus & 237 & MAVS  Rhinolophus ferrumequinum & interacts \\ \hline
% 1 & nsp15  Severe acute respiratory syndrome-related coronavirus & 238 & MAVS  Felis catus & interacts \\ \hline
% 2 & nsp15  Human coronavirus HKU1 & 77 & Human coronavirus HKU1 & belongs \\ \hline
% 3 & STAT1  Homo sapiens & 4 & STAT1  Pan troglodytes & similar \\ \hline
% 3 & STAT1  Homo sapiens & 5 & STAT1  Macaca mulatta & similar \\ \hline
% 3 & STAT1  Homo sapiens & 6 & STAT1  Felis catus & similar \\ \hline
% 3 & STAT1  Homo sapiens & 7 & STAT1  Camelus dromedarius & similar \\ \hline
% 3 & STAT1  Homo sapiens & 8 & STAT1  Equus caballus & similar \\ \hline
% 3 & STAT1  Homo sapiens & 9 & STAT1  Canis lupus familiaris & similar \\ \hline
% 3 & STAT1  Homo sapiens & 10 & STAT1  Rhinolophus ferrumequinum & similar \\ \hline
% 3 & STAT1  Homo sapiens & 11 & STAT1  Bos taurus & similar \\ \hline
% 3 & STAT1  Homo sapiens & 12 & STAT1  Ovis aries & similar \\ \hline
% 3 & STAT1  Homo sapiens & 13 & STAT1  Ictidomys tridecemlineatus & similar \\ \hline
% 3 & STAT1  Homo sapiens & 14 & STAT1  Rattus norvegicus & similar \\ \hline
% 3 & STAT1  Homo sapiens & 15 & STAT1  Mus musculus & similar \\ \hline
% 3 & STAT1  Homo sapiens & 16 & STAT1  Mesocricetus auratus & similar \\ \hline
% 3 & STAT1  Homo sapiens & 17 & STAT1  Gallus gallus & similar \\ \hline
% 3 & STAT1  Homo sapiens & 192 & Homo sapiens & belongs \\ \hline
% 3 & STAT1  Homo sapiens & 154 & ORF3b  Severe acute respiratory syndrome-related coronavirus & interacts \\ \hline
% 3 & STAT1  Homo sapiens & 170 & ORF6  Severe acute respiratory syndrome-related coronavirus & interacts \\ \hline
% 3 & STAT1  Homo sapiens & 209 & M protein  Middle East respiratory syndrome-related coronavirus & interacts \\ \hline
% 3 & STAT1  Homo sapiens & 45 & nsp1  Severe acute respiratory syndrome-related coronavirus & interacts \\ \hline
% 3 & STAT1  Homo sapiens & 100 & ORF4a  Middle East respiratory syndrome-related coronavirus & interacts \\ \hline
% 3 & STAT1  Homo sapiens & 42 & ORF4b  Middle East respiratory syndrome-related coronavirus & interacts \\ \hline
% 4 & STAT1  Pan troglodytes & 5 & STAT1  Macaca mulatta & similar \\ \hline
% 4 & STAT1  Pan troglodytes & 6 & STAT1  Felis catus & similar \\ \hline
% 4 & STAT1  Pan troglodytes & 7 & STAT1  Camelus dromedarius & similar \\ \hline
% 4 & STAT1  Pan troglodytes & 8 & STAT1  Equus caballus & similar \\ \hline
% 4 & STAT1  Pan troglodytes & 9 & STAT1  Canis lupus familiaris & similar \\ \hline
% 4 & STAT1  Pan troglodytes & 10 & STAT1  Rhinolophus ferrumequinum & similar \\ \hline
% 4 & STAT1  Pan troglodytes & 11 & STAT1  Bos taurus & similar \\ \hline
% 4 & STAT1  Pan troglodytes & 12 & STAT1  Ovis aries & similar \\ \hline
% 4 & STAT1  Pan troglodytes & 13 & STAT1  Ictidomys tridecemlineatus & similar \\ \hline
% 4 & STAT1  Pan troglodytes & 14 & STAT1  Rattus norvegicus & similar \\ \hline
% 4 & STAT1  Pan troglodytes & 15 & STAT1  Mus musculus & similar \\ \hline
% 4 & STAT1  Pan troglodytes & 16 & STAT1  Mesocricetus auratus & similar \\ \hline
% 4 & STAT1  Pan troglodytes & 17 & STAT1  Gallus gallus & similar \\ \hline
% 4 & STAT1  Pan troglodytes & 200 & Pan troglodytes & belongs \\ \hline
% 5 & STAT1  Macaca mulatta & 6 & STAT1  Felis catus & similar \\ \hline
% 5 & STAT1  Macaca mulatta & 7 & STAT1  Camelus dromedarius & similar \\ \hline
% 5 & STAT1  Macaca mulatta & 8 & STAT1  Equus caballus & similar \\ \hline
% 5 & STAT1  Macaca mulatta & 9 & STAT1  Canis lupus familiaris & similar \\ \hline
% 5 & STAT1  Macaca mulatta & 10 & STAT1  Rhinolophus ferrumequinum & similar \\ \hline
% 5 & STAT1  Macaca mulatta & 11 & STAT1  Bos taurus & similar \\ \hline
% 5 & STAT1  Macaca mulatta & 12 & STAT1  Ovis aries & similar \\ \hline
% 5 & STAT1  Macaca mulatta & 13 & STAT1  Ictidomys tridecemlineatus & similar \\ \hline
% 5 & STAT1  Macaca mulatta & 14 & STAT1  Rattus norvegicus & similar \\ \hline
% 5 & STAT1  Macaca mulatta & 15 & STAT1  Mus musculus & similar \\ \hline
% 5 & STAT1  Macaca mulatta & 16 & STAT1  Mesocricetus auratus & similar \\ \hline
% 5 & STAT1  Macaca mulatta & 17 & STAT1  Gallus gallus & similar \\ \hline
% 5 & STAT1  Macaca mulatta & 204 & Macaca mulatta & belongs \\ \hline
% 6 & STAT1  Felis catus & 7 & STAT1  Camelus dromedarius & similar \\ \hline
% 6 & STAT1  Felis catus & 8 & STAT1  Equus caballus & similar \\ \hline
% 6 & STAT1  Felis catus & 9 & STAT1  Canis lupus familiaris & similar \\ \hline
% 6 & STAT1  Felis catus & 10 & STAT1  Rhinolophus ferrumequinum & similar \\ \hline
% 6 & STAT1  Felis catus & 11 & STAT1  Bos taurus & similar \\ \hline
% 6 & STAT1  Felis catus & 12 & STAT1  Ovis aries & similar \\ \hline
% 6 & STAT1  Felis catus & 13 & STAT1  Ictidomys tridecemlineatus & similar \\ \hline
% 6 & STAT1  Felis catus & 14 & STAT1  Rattus norvegicus & similar \\ \hline
% 6 & STAT1  Felis catus & 15 & STAT1  Mus musculus & similar \\ \hline
% 6 & STAT1  Felis catus & 16 & STAT1  Mesocricetus auratus & similar \\ \hline
% 6 & STAT1  Felis catus & 17 & STAT1  Gallus gallus & similar \\ \hline
% 6 & STAT1  Felis catus & 197 & Felis catus & belongs \\ \hline
% 6 & STAT1  Felis catus & 154 & ORF3b  Severe acute respiratory syndrome-related coronavirus & interacts \\ \hline
% 6 & STAT1  Felis catus & 170 & ORF6  Severe acute respiratory syndrome-related coronavirus & interacts \\ \hline
% 6 & STAT1  Felis catus & 45 & nsp1  Severe acute respiratory syndrome-related coronavirus & interacts \\ \hline
% 7 & STAT1  Camelus dromedarius & 8 & STAT1  Equus caballus & similar \\ \hline
% 7 & STAT1  Camelus dromedarius & 9 & STAT1  Canis lupus familiaris & similar \\ \hline
% 7 & STAT1  Camelus dromedarius & 10 & STAT1  Rhinolophus ferrumequinum & similar \\ \hline
% 7 & STAT1  Camelus dromedarius & 11 & STAT1  Bos taurus & similar \\ \hline
% 7 & STAT1  Camelus dromedarius & 12 & STAT1  Ovis aries & similar \\ \hline
% 7 & STAT1  Camelus dromedarius & 13 & STAT1  Ictidomys tridecemlineatus & similar \\ \hline
% 7 & STAT1  Camelus dromedarius & 14 & STAT1  Rattus norvegicus & similar \\ \hline
% 7 & STAT1  Camelus dromedarius & 15 & STAT1  Mus musculus & similar \\ \hline
% 7 & STAT1  Camelus dromedarius & 16 & STAT1  Mesocricetus auratus & similar \\ \hline
% 7 & STAT1  Camelus dromedarius & 17 & STAT1  Gallus gallus & similar \\ \hline
% 7 & STAT1  Camelus dromedarius & 196 & Camelus dromedarius & belongs \\ \hline
% 7 & STAT1  Camelus dromedarius & 209 & M protein  Middle East respiratory syndrome-related coronavirus & interacts \\ \hline
% 7 & STAT1  Camelus dromedarius & 100 & ORF4a  Middle East respiratory syndrome-related coronavirus & interacts \\ \hline
% 7 & STAT1  Camelus dromedarius & 42 & ORF4b  Middle East respiratory syndrome-related coronavirus & interacts \\ \hline
% 8 & STAT1  Equus caballus & 9 & STAT1  Canis lupus familiaris & similar \\ \hline
% 8 & STAT1  Equus caballus & 10 & STAT1  Rhinolophus ferrumequinum & similar \\ \hline
% 8 & STAT1  Equus caballus & 11 & STAT1  Bos taurus & similar \\ \hline
% 8 & STAT1  Equus caballus & 12 & STAT1  Ovis aries & similar \\ \hline
% 8 & STAT1  Equus caballus & 13 & STAT1  Ictidomys tridecemlineatus & similar \\ \hline
% 8 & STAT1  Equus caballus & 14 & STAT1  Rattus norvegicus & similar \\ \hline
% 8 & STAT1  Equus caballus & 15 & STAT1  Mus musculus & similar \\ \hline
% 8 & STAT1  Equus caballus & 16 & STAT1  Mesocricetus auratus & similar \\ \hline
% 8 & STAT1  Equus caballus & 17 & STAT1  Gallus gallus & similar \\ \hline
% 8 & STAT1  Equus caballus & 203 & Equus caballus & belongs \\ \hline
% 9 & STAT1  Canis lupus familiaris & 10 & STAT1  Rhinolophus ferrumequinum & similar \\ \hline
% 9 & STAT1  Canis lupus familiaris & 11 & STAT1  Bos taurus & similar \\ \hline
% 9 & STAT1  Canis lupus familiaris & 12 & STAT1  Ovis aries & similar \\ \hline
% 9 & STAT1  Canis lupus familiaris & 13 & STAT1  Ictidomys tridecemlineatus & similar \\ \hline
% 9 & STAT1  Canis lupus familiaris & 14 & STAT1  Rattus norvegicus & similar \\ \hline
% 9 & STAT1  Canis lupus familiaris & 15 & STAT1  Mus musculus & similar \\ \hline
% 9 & STAT1  Canis lupus familiaris & 16 & STAT1  Mesocricetus auratus & similar \\ \hline
% 9 & STAT1  Canis lupus familiaris & 17 & STAT1  Gallus gallus & similar \\ \hline
% 9 & STAT1  Canis lupus familiaris & 195 & Canis lupus familiaris & belongs \\ \hline
% 10 & STAT1  Rhinolophus ferrumequinum & 11 & STAT1  Bos taurus & similar \\ \hline
% 10 & STAT1  Rhinolophus ferrumequinum & 12 & STAT1  Ovis aries & similar \\ \hline
% 10 & STAT1  Rhinolophus ferrumequinum & 13 & STAT1  Ictidomys tridecemlineatus & similar \\ \hline
% 10 & STAT1  Rhinolophus ferrumequinum & 14 & STAT1  Rattus norvegicus & similar \\ \hline
% 10 & STAT1  Rhinolophus ferrumequinum & 15 & STAT1  Mus musculus & similar \\ \hline
% 10 & STAT1  Rhinolophus ferrumequinum & 16 & STAT1  Mesocricetus auratus & similar \\ \hline
% 10 & STAT1  Rhinolophus ferrumequinum & 17 & STAT1  Gallus gallus & similar \\ \hline
% 10 & STAT1  Rhinolophus ferrumequinum & 207 & Rhinolophus ferrumequinum & belongs \\ \hline
% 10 & STAT1  Rhinolophus ferrumequinum & 154 & ORF3b  Severe acute respiratory syndrome-related coronavirus & interacts \\ \hline
% 10 & STAT1  Rhinolophus ferrumequinum & 170 & ORF6  Severe acute respiratory syndrome-related coronavirus & interacts \\ \hline
% 10 & STAT1  Rhinolophus ferrumequinum & 45 & nsp1  Severe acute respiratory syndrome-related coronavirus & interacts \\ \hline
% 11 & STAT1  Bos taurus & 12 & STAT1  Ovis aries & similar \\ \hline
% 11 & STAT1  Bos taurus & 13 & STAT1  Ictidomys tridecemlineatus & similar \\ \hline
% 11 & STAT1  Bos taurus & 14 & STAT1  Rattus norvegicus & similar \\ \hline
% 11 & STAT1  Bos taurus & 15 & STAT1  Mus musculus & similar \\ \hline
% 11 & STAT1  Bos taurus & 16 & STAT1  Mesocricetus auratus & similar \\ \hline
% 11 & STAT1  Bos taurus & 17 & STAT1  Gallus gallus & similar \\ \hline
% 11 & STAT1  Bos taurus & 199 & Bos taurus & belongs \\ \hline
% 12 & STAT1  Ovis aries & 13 & STAT1  Ictidomys tridecemlineatus & similar \\ \hline
% 12 & STAT1  Ovis aries & 14 & STAT1  Rattus norvegicus & similar \\ \hline
% 12 & STAT1  Ovis aries & 15 & STAT1  Mus musculus & similar \\ \hline
% 12 & STAT1  Ovis aries & 16 & STAT1  Mesocricetus auratus & similar \\ \hline
% 12 & STAT1  Ovis aries & 17 & STAT1  Gallus gallus & similar \\ \hline
% 12 & STAT1  Ovis aries & 205 & Ovis aries & belongs \\ \hline
% 13 & STAT1  Ictidomys tridecemlineatus & 14 & STAT1  Rattus norvegicus & similar \\ \hline
% 13 & STAT1  Ictidomys tridecemlineatus & 15 & STAT1  Mus musculus & similar \\ \hline
% 13 & STAT1  Ictidomys tridecemlineatus & 16 & STAT1  Mesocricetus auratus & similar \\ \hline
% 13 & STAT1  Ictidomys tridecemlineatus & 17 & STAT1  Gallus gallus & similar \\ \hline
% 13 & STAT1  Ictidomys tridecemlineatus & 198 & Ictidomys tridecemlineatus & belongs \\ \hline
% 14 & STAT1  Rattus norvegicus & 15 & STAT1  Mus musculus & similar \\ \hline
% 14 & STAT1  Rattus norvegicus & 16 & STAT1  Mesocricetus auratus & similar \\ \hline
% 14 & STAT1  Rattus norvegicus & 17 & STAT1  Gallus gallus & similar \\ \hline
% 14 & STAT1  Rattus norvegicus & 194 & Rattus norvegicus & belongs \\ \hline
% 14 & STAT1  Rattus norvegicus & 154 & ORF3b  Severe acute respiratory syndrome-related coronavirus & interacts \\ \hline
% 14 & STAT1  Rattus norvegicus & 170 & ORF6  Severe acute respiratory syndrome-related coronavirus & interacts \\ \hline
% 14 & STAT1  Rattus norvegicus & 209 & M protein  Middle East respiratory syndrome-related coronavirus & interacts \\ \hline
% 14 & STAT1  Rattus norvegicus & 45 & nsp1  Severe acute respiratory syndrome-related coronavirus & interacts \\ \hline
% 14 & STAT1  Rattus norvegicus & 100 & ORF4a  Middle East respiratory syndrome-related coronavirus & interacts \\ \hline
% 14 & STAT1  Rattus norvegicus & 42 & ORF4b  Middle East respiratory syndrome-related coronavirus & interacts \\ \hline
% 15 & STAT1  Mus musculus & 16 & STAT1  Mesocricetus auratus & similar \\ \hline
% 15 & STAT1  Mus musculus & 17 & STAT1  Gallus gallus & similar \\ \hline
% 15 & STAT1  Mus musculus & 193 & Mus musculus & belongs \\ \hline
% 15 & STAT1  Mus musculus & 154 & ORF3b  Severe acute respiratory syndrome-related coronavirus & interacts \\ \hline
% 15 & STAT1  Mus musculus & 170 & ORF6  Severe acute respiratory syndrome-related coronavirus & interacts \\ \hline
% 15 & STAT1  Mus musculus & 209 & M protein  Middle East respiratory syndrome-related coronavirus & interacts \\ \hline
% 15 & STAT1  Mus musculus & 45 & nsp1  Severe acute respiratory syndrome-related coronavirus & interacts \\ \hline
% 15 & STAT1  Mus musculus & 100 & ORF4a  Middle East respiratory syndrome-related coronavirus & interacts \\ \hline
% 15 & STAT1  Mus musculus & 42 & ORF4b  Middle East respiratory syndrome-related coronavirus & interacts \\ \hline
% 16 & STAT1  Mesocricetus auratus & 17 & STAT1  Gallus gallus & similar \\ \hline
% 16 & STAT1  Mesocricetus auratus & 208 & Mesocricetus auratus & belongs \\ \hline
% 17 & STAT1  Gallus gallus & 201 & Gallus gallus & belongs \\ \hline
% 18 & IRF9  Homo sapiens & 19 & IRF9  Macaca mulatta & similar \\ \hline
% 18 & IRF9  Homo sapiens & 20 & IRF9  Pan troglodytes & similar \\ \hline
% 18 & IRF9  Homo sapiens & 21 & IRF9  Felis catus & similar \\ \hline
% 18 & IRF9  Homo sapiens & 22 & IRF9  Camelus dromedarius & similar \\ \hline
% 18 & IRF9  Homo sapiens & 23 & IRF9  Sus scrofa domesticus & similar \\ \hline
% 18 & IRF9  Homo sapiens & 24 & IRF9  Equus caballus & similar \\ \hline
% 18 & IRF9  Homo sapiens & 25 & IRF9  Rhinolophus ferrumequinum & similar \\ \hline
% 18 & IRF9  Homo sapiens & 26 & IRF9  Ovis aries & similar \\ \hline
% 18 & IRF9  Homo sapiens & 27 & IRF9  Canis lupus familiaris & similar \\ \hline
% 18 & IRF9  Homo sapiens & 28 & IRF9  Bos taurus & similar \\ \hline
% 18 & IRF9  Homo sapiens & 29 & IRF9  Ictidomys tridecemlineatus & similar \\ \hline
% 18 & IRF9  Homo sapiens & 30 & IRF9  Mus musculus & similar \\ \hline
% 18 & IRF9  Homo sapiens & 31 & IRF9  Mesocricetus auratus & similar \\ \hline
% 18 & IRF9  Homo sapiens & 32 & IRF9  Rattus norvegicus & similar \\ \hline
% 18 & IRF9  Homo sapiens & 33 & IRF9  Gallus gallus & similar \\ \hline
% 18 & IRF9  Homo sapiens & 192 & Homo sapiens & belongs \\ \hline
% 18 & IRF9  Homo sapiens & 154 & ORF3b  Severe acute respiratory syndrome-related coronavirus & interacts \\ \hline
% 18 & IRF9  Homo sapiens & 170 & ORF6  Severe acute respiratory syndrome-related coronavirus & interacts \\ \hline
% 18 & IRF9  Homo sapiens & 209 & M protein  Middle East respiratory syndrome-related coronavirus & interacts \\ \hline
% 18 & IRF9  Homo sapiens & 100 & ORF4a  Middle East respiratory syndrome-related coronavirus & interacts \\ \hline
% 18 & IRF9  Homo sapiens & 42 & ORF4b  Middle East respiratory syndrome-related coronavirus & interacts \\ \hline
% 19 & IRF9  Macaca mulatta & 20 & IRF9  Pan troglodytes & similar \\ \hline
% 19 & IRF9  Macaca mulatta & 21 & IRF9  Felis catus & similar \\ \hline
% 19 & IRF9  Macaca mulatta & 22 & IRF9  Camelus dromedarius & similar \\ \hline
% 19 & IRF9  Macaca mulatta & 23 & IRF9  Sus scrofa domesticus & similar \\ \hline
% 19 & IRF9  Macaca mulatta & 24 & IRF9  Equus caballus & similar \\ \hline
% 19 & IRF9  Macaca mulatta & 25 & IRF9  Rhinolophus ferrumequinum & similar \\ \hline
% 19 & IRF9  Macaca mulatta & 26 & IRF9  Ovis aries & similar \\ \hline
% 19 & IRF9  Macaca mulatta & 27 & IRF9  Canis lupus familiaris & similar \\ \hline
% 19 & IRF9  Macaca mulatta & 28 & IRF9  Bos taurus & similar \\ \hline
% 19 & IRF9  Macaca mulatta & 29 & IRF9  Ictidomys tridecemlineatus & similar \\ \hline
% 19 & IRF9  Macaca mulatta & 30 & IRF9  Mus musculus & similar \\ \hline
% 19 & IRF9  Macaca mulatta & 31 & IRF9  Mesocricetus auratus & similar \\ \hline
% 19 & IRF9  Macaca mulatta & 32 & IRF9  Rattus norvegicus & similar \\ \hline
% 19 & IRF9  Macaca mulatta & 33 & IRF9  Gallus gallus & similar \\ \hline
% 19 & IRF9  Macaca mulatta & 204 & Macaca mulatta & belongs \\ \hline
% 20 & IRF9  Pan troglodytes & 21 & IRF9  Felis catus & similar \\ \hline
% 20 & IRF9  Pan troglodytes & 22 & IRF9  Camelus dromedarius & similar \\ \hline
% 20 & IRF9  Pan troglodytes & 23 & IRF9  Sus scrofa domesticus & similar \\ \hline
% 20 & IRF9  Pan troglodytes & 24 & IRF9  Equus caballus & similar \\ \hline
% 20 & IRF9  Pan troglodytes & 25 & IRF9  Rhinolophus ferrumequinum & similar \\ \hline
% 20 & IRF9  Pan troglodytes & 26 & IRF9  Ovis aries & similar \\ \hline
% 20 & IRF9  Pan troglodytes & 27 & IRF9  Canis lupus familiaris & similar \\ \hline
% 20 & IRF9  Pan troglodytes & 28 & IRF9  Bos taurus & similar \\ \hline
% 20 & IRF9  Pan troglodytes & 29 & IRF9  Ictidomys tridecemlineatus & similar \\ \hline
% 20 & IRF9  Pan troglodytes & 30 & IRF9  Mus musculus & similar \\ \hline
% 20 & IRF9  Pan troglodytes & 31 & IRF9  Mesocricetus auratus & similar \\ \hline
% 20 & IRF9  Pan troglodytes & 32 & IRF9  Rattus norvegicus & similar \\ \hline
% 20 & IRF9  Pan troglodytes & 33 & IRF9  Gallus gallus & similar \\ \hline
% 20 & IRF9  Pan troglodytes & 200 & Pan troglodytes & belongs \\ \hline
% 21 & IRF9  Felis catus & 22 & IRF9  Camelus dromedarius & similar \\ \hline
% 21 & IRF9  Felis catus & 23 & IRF9  Sus scrofa domesticus & similar \\ \hline
% 21 & IRF9  Felis catus & 24 & IRF9  Equus caballus & similar \\ \hline
% 21 & IRF9  Felis catus & 25 & IRF9  Rhinolophus ferrumequinum & similar \\ \hline
% 21 & IRF9  Felis catus & 26 & IRF9  Ovis aries & similar \\ \hline
% 21 & IRF9  Felis catus & 27 & IRF9  Canis lupus familiaris & similar \\ \hline
% 21 & IRF9  Felis catus & 28 & IRF9  Bos taurus & similar \\ \hline
% 21 & IRF9  Felis catus & 29 & IRF9  Ictidomys tridecemlineatus & similar \\ \hline
% 21 & IRF9  Felis catus & 30 & IRF9  Mus musculus & similar \\ \hline
% 21 & IRF9  Felis catus & 31 & IRF9  Mesocricetus auratus & similar \\ \hline
% 21 & IRF9  Felis catus & 32 & IRF9  Rattus norvegicus & similar \\ \hline
% 21 & IRF9  Felis catus & 33 & IRF9  Gallus gallus & similar \\ \hline
% 21 & IRF9  Felis catus & 197 & Felis catus & belongs \\ \hline
% 21 & IRF9  Felis catus & 154 & ORF3b  Severe acute respiratory syndrome-related coronavirus & interacts \\ \hline
% 21 & IRF9  Felis catus & 170 & ORF6  Severe acute respiratory syndrome-related coronavirus & interacts \\ \hline
% 22 & IRF9  Camelus dromedarius & 23 & IRF9  Sus scrofa domesticus & similar \\ \hline
% 22 & IRF9  Camelus dromedarius & 24 & IRF9  Equus caballus & similar \\ \hline
% 22 & IRF9  Camelus dromedarius & 25 & IRF9  Rhinolophus ferrumequinum & similar \\ \hline
% 22 & IRF9  Camelus dromedarius & 26 & IRF9  Ovis aries & similar \\ \hline
% 22 & IRF9  Camelus dromedarius & 27 & IRF9  Canis lupus familiaris & similar \\ \hline
% 22 & IRF9  Camelus dromedarius & 28 & IRF9  Bos taurus & similar \\ \hline
% 22 & IRF9  Camelus dromedarius & 29 & IRF9  Ictidomys tridecemlineatus & similar \\ \hline
% 22 & IRF9  Camelus dromedarius & 30 & IRF9  Mus musculus & similar \\ \hline
% 22 & IRF9  Camelus dromedarius & 31 & IRF9  Mesocricetus auratus & similar \\ \hline
% 22 & IRF9  Camelus dromedarius & 32 & IRF9  Rattus norvegicus & similar \\ \hline
% 22 & IRF9  Camelus dromedarius & 33 & IRF9  Gallus gallus & similar \\ \hline
% 22 & IRF9  Camelus dromedarius & 196 & Camelus dromedarius & belongs \\ \hline
% 22 & IRF9  Camelus dromedarius & 209 & M protein  Middle East respiratory syndrome-related coronavirus & interacts \\ \hline
% 22 & IRF9  Camelus dromedarius & 100 & ORF4a  Middle East respiratory syndrome-related coronavirus & interacts \\ \hline
% 22 & IRF9  Camelus dromedarius & 42 & ORF4b  Middle East respiratory syndrome-related coronavirus & interacts \\ \hline
% 23 & IRF9  Sus scrofa domesticus & 24 & IRF9  Equus caballus & similar \\ \hline
% 23 & IRF9  Sus scrofa domesticus & 25 & IRF9  Rhinolophus ferrumequinum & similar \\ \hline
% 23 & IRF9  Sus scrofa domesticus & 26 & IRF9  Ovis aries & similar \\ \hline
% 23 & IRF9  Sus scrofa domesticus & 27 & IRF9  Canis lupus familiaris & similar \\ \hline
% 23 & IRF9  Sus scrofa domesticus & 28 & IRF9  Bos taurus & similar \\ \hline
% 23 & IRF9  Sus scrofa domesticus & 29 & IRF9  Ictidomys tridecemlineatus & similar \\ \hline
% 23 & IRF9  Sus scrofa domesticus & 30 & IRF9  Mus musculus & similar \\ \hline
% 23 & IRF9  Sus scrofa domesticus & 31 & IRF9  Mesocricetus auratus & similar \\ \hline
% 23 & IRF9  Sus scrofa domesticus & 32 & IRF9  Rattus norvegicus & similar \\ \hline
% 23 & IRF9  Sus scrofa domesticus & 33 & IRF9  Gallus gallus & similar \\ \hline
% 23 & IRF9  Sus scrofa domesticus & 206 & Sus scrofa domesticus & belongs \\ \hline
% 24 & IRF9  Equus caballus & 25 & IRF9  Rhinolophus ferrumequinum & similar \\ \hline
% 24 & IRF9  Equus caballus & 26 & IRF9  Ovis aries & similar \\ \hline
% 24 & IRF9  Equus caballus & 27 & IRF9  Canis lupus familiaris & similar \\ \hline
% 24 & IRF9  Equus caballus & 28 & IRF9  Bos taurus & similar \\ \hline
% 24 & IRF9  Equus caballus & 29 & IRF9  Ictidomys tridecemlineatus & similar \\ \hline
% 24 & IRF9  Equus caballus & 30 & IRF9  Mus musculus & similar \\ \hline
% 24 & IRF9  Equus caballus & 31 & IRF9  Mesocricetus auratus & similar \\ \hline
% 24 & IRF9  Equus caballus & 32 & IRF9  Rattus norvegicus & similar \\ \hline
% 24 & IRF9  Equus caballus & 33 & IRF9  Gallus gallus & similar \\ \hline
% 24 & IRF9  Equus caballus & 203 & Equus caballus & belongs \\ \hline
% 25 & IRF9  Rhinolophus ferrumequinum & 26 & IRF9  Ovis aries & similar \\ \hline
% 25 & IRF9  Rhinolophus ferrumequinum & 27 & IRF9  Canis lupus familiaris & similar \\ \hline
% 25 & IRF9  Rhinolophus ferrumequinum & 28 & IRF9  Bos taurus & similar \\ \hline
% 25 & IRF9  Rhinolophus ferrumequinum & 29 & IRF9  Ictidomys tridecemlineatus & similar \\ \hline
% 25 & IRF9  Rhinolophus ferrumequinum & 30 & IRF9  Mus musculus & similar \\ \hline
% 25 & IRF9  Rhinolophus ferrumequinum & 31 & IRF9  Mesocricetus auratus & similar \\ \hline
% 25 & IRF9  Rhinolophus ferrumequinum & 32 & IRF9  Rattus norvegicus & similar \\ \hline
% 25 & IRF9  Rhinolophus ferrumequinum & 33 & IRF9  Gallus gallus & similar \\ \hline
% 25 & IRF9  Rhinolophus ferrumequinum & 207 & Rhinolophus ferrumequinum & belongs \\ \hline
% 25 & IRF9  Rhinolophus ferrumequinum & 154 & ORF3b  Severe acute respiratory syndrome-related coronavirus & interacts \\ \hline
% 25 & IRF9  Rhinolophus ferrumequinum & 170 & ORF6  Severe acute respiratory syndrome-related coronavirus & interacts \\ \hline
% 26 & IRF9  Ovis aries & 27 & IRF9  Canis lupus familiaris & similar \\ \hline
% 26 & IRF9  Ovis aries & 28 & IRF9  Bos taurus & similar \\ \hline
% 26 & IRF9  Ovis aries & 29 & IRF9  Ictidomys tridecemlineatus & similar \\ \hline
% 26 & IRF9  Ovis aries & 30 & IRF9  Mus musculus & similar \\ \hline
% 26 & IRF9  Ovis aries & 31 & IRF9  Mesocricetus auratus & similar \\ \hline
% 26 & IRF9  Ovis aries & 32 & IRF9  Rattus norvegicus & similar \\ \hline
% 26 & IRF9  Ovis aries & 33 & IRF9  Gallus gallus & similar \\ \hline
% 26 & IRF9  Ovis aries & 205 & Ovis aries & belongs \\ \hline
% 27 & IRF9  Canis lupus familiaris & 28 & IRF9  Bos taurus & similar \\ \hline
% 27 & IRF9  Canis lupus familiaris & 29 & IRF9  Ictidomys tridecemlineatus & similar \\ \hline
% 27 & IRF9  Canis lupus familiaris & 30 & IRF9  Mus musculus & similar \\ \hline
% 27 & IRF9  Canis lupus familiaris & 31 & IRF9  Mesocricetus auratus & similar \\ \hline
% 27 & IRF9  Canis lupus familiaris & 32 & IRF9  Rattus norvegicus & similar \\ \hline
% 27 & IRF9  Canis lupus familiaris & 33 & IRF9  Gallus gallus & similar \\ \hline
% 27 & IRF9  Canis lupus familiaris & 195 & Canis lupus familiaris & belongs \\ \hline
% 28 & IRF9  Bos taurus & 29 & IRF9  Ictidomys tridecemlineatus & similar \\ \hline
% 28 & IRF9  Bos taurus & 30 & IRF9  Mus musculus & similar \\ \hline
% 28 & IRF9  Bos taurus & 31 & IRF9  Mesocricetus auratus & similar \\ \hline
% 28 & IRF9  Bos taurus & 32 & IRF9  Rattus norvegicus & similar \\ \hline
% 28 & IRF9  Bos taurus & 33 & IRF9  Gallus gallus & similar \\ \hline
% 28 & IRF9  Bos taurus & 199 & Bos taurus & belongs \\ \hline
% 29 & IRF9  Ictidomys tridecemlineatus & 30 & IRF9  Mus musculus & similar \\ \hline
% 29 & IRF9  Ictidomys tridecemlineatus & 31 & IRF9  Mesocricetus auratus & similar \\ \hline
% 29 & IRF9  Ictidomys tridecemlineatus & 32 & IRF9  Rattus norvegicus & similar \\ \hline
% 29 & IRF9  Ictidomys tridecemlineatus & 33 & IRF9  Gallus gallus & similar \\ \hline
% 29 & IRF9  Ictidomys tridecemlineatus & 198 & Ictidomys tridecemlineatus & belongs \\ \hline
% 30 & IRF9  Mus musculus & 31 & IRF9  Mesocricetus auratus & similar \\ \hline
% 30 & IRF9  Mus musculus & 32 & IRF9  Rattus norvegicus & similar \\ \hline
% 30 & IRF9  Mus musculus & 33 & IRF9  Gallus gallus & similar \\ \hline
% 30 & IRF9  Mus musculus & 193 & Mus musculus & belongs \\ \hline
% 30 & IRF9  Mus musculus & 154 & ORF3b  Severe acute respiratory syndrome-related coronavirus & interacts \\ \hline
% 30 & IRF9  Mus musculus & 170 & ORF6  Severe acute respiratory syndrome-related coronavirus & interacts \\ \hline
% 30 & IRF9  Mus musculus & 209 & M protein  Middle East respiratory syndrome-related coronavirus & interacts \\ \hline
% 30 & IRF9  Mus musculus & 100 & ORF4a  Middle East respiratory syndrome-related coronavirus & interacts \\ \hline
% 30 & IRF9  Mus musculus & 42 & ORF4b  Middle East respiratory syndrome-related coronavirus & interacts \\ \hline
% 31 & IRF9  Mesocricetus auratus & 32 & IRF9  Rattus norvegicus & similar \\ \hline
% 31 & IRF9  Mesocricetus auratus & 33 & IRF9  Gallus gallus & similar \\ \hline
% 31 & IRF9  Mesocricetus auratus & 208 & Mesocricetus auratus & belongs \\ \hline
% 32 & IRF9  Rattus norvegicus & 33 & IRF9  Gallus gallus & similar \\ \hline
% 32 & IRF9  Rattus norvegicus & 194 & Rattus norvegicus & belongs \\ \hline
% 32 & IRF9  Rattus norvegicus & 154 & ORF3b  Severe acute respiratory syndrome-related coronavirus & interacts \\ \hline
% 32 & IRF9  Rattus norvegicus & 170 & ORF6  Severe acute respiratory syndrome-related coronavirus & interacts \\ \hline
% 32 & IRF9  Rattus norvegicus & 209 & M protein  Middle East respiratory syndrome-related coronavirus & interacts \\ \hline
% 32 & IRF9  Rattus norvegicus & 100 & ORF4a  Middle East respiratory syndrome-related coronavirus & interacts \\ \hline
% 32 & IRF9  Rattus norvegicus & 42 & ORF4b  Middle East respiratory syndrome-related coronavirus & interacts \\ \hline
% 33 & IRF9  Gallus gallus & 201 & Gallus gallus & belongs \\ \hline
% 34 & RIG-I  Homo sapiens & 35 & RIG-I  Pan troglodytes & similar \\ \hline
% 34 & RIG-I  Homo sapiens & 36 & RIG-I  Macaca mulatta & similar \\ \hline
% 34 & RIG-I  Homo sapiens & 37 & RIG-I  Rhinolophus ferrumequinum & similar \\ \hline
% 34 & RIG-I  Homo sapiens & 38 & RIG-I  Canis lupus familiaris & similar \\ \hline
% 34 & RIG-I  Homo sapiens & 39 & RIG-I  Mesocricetus auratus & similar \\ \hline
% 34 & RIG-I  Homo sapiens & 40 & RIG-I  Mus musculus & similar \\ \hline
% 34 & RIG-I  Homo sapiens & 41 & RIG-I  Gallus gallus & similar \\ \hline
% 34 & RIG-I  Homo sapiens & 192 & Homo sapiens & belongs \\ \hline
% 34 & RIG-I  Homo sapiens & 154 & ORF3b  Severe acute respiratory syndrome-related coronavirus & interacts \\ \hline
% 34 & RIG-I  Homo sapiens & 171 & ORF6  Severe acute respiratory syndrome coronavirus 2 & interacts \\ \hline
% 34 & RIG-I  Homo sapiens & 71 & N protein  Severe acute respiratory syndrome-related coronavirus & interacts \\ \hline
% 35 & RIG-I  Pan troglodytes & 36 & RIG-I  Macaca mulatta & similar \\ \hline
% 35 & RIG-I  Pan troglodytes & 37 & RIG-I  Rhinolophus ferrumequinum & similar \\ \hline
% 35 & RIG-I  Pan troglodytes & 38 & RIG-I  Canis lupus familiaris & similar \\ \hline
% 35 & RIG-I  Pan troglodytes & 39 & RIG-I  Mesocricetus auratus & similar \\ \hline
% 35 & RIG-I  Pan troglodytes & 40 & RIG-I  Mus musculus & similar \\ \hline
% 35 & RIG-I  Pan troglodytes & 41 & RIG-I  Gallus gallus & similar \\ \hline
% 35 & RIG-I  Pan troglodytes & 200 & Pan troglodytes & belongs \\ \hline
% 36 & RIG-I  Macaca mulatta & 37 & RIG-I  Rhinolophus ferrumequinum & similar \\ \hline
% 36 & RIG-I  Macaca mulatta & 38 & RIG-I  Canis lupus familiaris & similar \\ \hline
% 36 & RIG-I  Macaca mulatta & 39 & RIG-I  Mesocricetus auratus & similar \\ \hline
% 36 & RIG-I  Macaca mulatta & 40 & RIG-I  Mus musculus & similar \\ \hline
% 36 & RIG-I  Macaca mulatta & 41 & RIG-I  Gallus gallus & similar \\ \hline
% 36 & RIG-I  Macaca mulatta & 204 & Macaca mulatta & belongs \\ \hline
% 37 & RIG-I  Rhinolophus ferrumequinum & 38 & RIG-I  Canis lupus familiaris & similar \\ \hline
% 37 & RIG-I  Rhinolophus ferrumequinum & 39 & RIG-I  Mesocricetus auratus & similar \\ \hline
% 37 & RIG-I  Rhinolophus ferrumequinum & 40 & RIG-I  Mus musculus & similar \\ \hline
% 37 & RIG-I  Rhinolophus ferrumequinum & 41 & RIG-I  Gallus gallus & similar \\ \hline
% 37 & RIG-I  Rhinolophus ferrumequinum & 207 & Rhinolophus ferrumequinum & belongs \\ \hline
% 37 & RIG-I  Rhinolophus ferrumequinum & 154 & ORF3b  Severe acute respiratory syndrome-related coronavirus & interacts \\ \hline
% 37 & RIG-I  Rhinolophus ferrumequinum & 71 & N protein  Severe acute respiratory syndrome-related coronavirus & interacts \\ \hline
% 38 & RIG-I  Canis lupus familiaris & 39 & RIG-I  Mesocricetus auratus & similar \\ \hline
% 38 & RIG-I  Canis lupus familiaris & 40 & RIG-I  Mus musculus & similar \\ \hline
% 38 & RIG-I  Canis lupus familiaris & 41 & RIG-I  Gallus gallus & similar \\ \hline
% 38 & RIG-I  Canis lupus familiaris & 195 & Canis lupus familiaris & belongs \\ \hline
% 38 & RIG-I  Canis lupus familiaris & 171 & ORF6  Severe acute respiratory syndrome coronavirus 2 & interacts \\ \hline
% 39 & RIG-I  Mesocricetus auratus & 40 & RIG-I  Mus musculus & similar \\ \hline
% 39 & RIG-I  Mesocricetus auratus & 41 & RIG-I  Gallus gallus & similar \\ \hline
% 39 & RIG-I  Mesocricetus auratus & 208 & Mesocricetus auratus & belongs \\ \hline
% 39 & RIG-I  Mesocricetus auratus & 171 & ORF6  Severe acute respiratory syndrome coronavirus 2 & interacts \\ \hline
% 40 & RIG-I  Mus musculus & 41 & RIG-I  Gallus gallus & similar \\ \hline
% 40 & RIG-I  Mus musculus & 193 & Mus musculus & belongs \\ \hline
% 40 & RIG-I  Mus musculus & 154 & ORF3b  Severe acute respiratory syndrome-related coronavirus & interacts \\ \hline
% 40 & RIG-I  Mus musculus & 171 & ORF6  Severe acute respiratory syndrome coronavirus 2 & interacts \\ \hline
% 40 & RIG-I  Mus musculus & 71 & N protein  Severe acute respiratory syndrome-related coronavirus & interacts \\ \hline
% 41 & RIG-I  Gallus gallus & 201 & Gallus gallus & belongs \\ \hline
% 42 & ORF4b  Middle East respiratory syndrome-related coronavirus & 43 & ORF4b  Human coronavirus 229E & similar \\ \hline
% 42 & ORF4b  Middle East respiratory syndrome-related coronavirus & 78 & Middle East respiratory syndrome-related coronavirus & belongs \\ \hline
% 42 & ORF4b  Middle East respiratory syndrome-related coronavirus & 172 & STAT2  Homo sapiens & interacts \\ \hline
% 42 & ORF4b  Middle East respiratory syndrome-related coronavirus & 177 & STAT2  Camelus dromedarius & interacts \\ \hline
% 42 & ORF4b  Middle East respiratory syndrome-related coronavirus & 183 & STAT2  Rattus norvegicus & interacts \\ \hline
% 42 & ORF4b  Middle East respiratory syndrome-related coronavirus & 185 & STAT2  Mus musculus & interacts \\ \hline
% 42 & ORF4b  Middle East respiratory syndrome-related coronavirus & 102 & NF-$\kappa$B  Homo sapiens & interacts \\ \hline
% 42 & ORF4b  Middle East respiratory syndrome-related coronavirus & 109 & NF-$\kappa$B  Mus musculus & interacts \\ \hline
% 42 & ORF4b  Middle East respiratory syndrome-related coronavirus & 111 & NF-$\kappa$B  Rattus norvegicus & interacts \\ \hline
% 42 & ORF4b  Middle East respiratory syndrome-related coronavirus & 114 & NF-$\kappa$B  Camelus dromedarius & interacts \\ \hline
% 42 & ORF4b  Middle East respiratory syndrome-related coronavirus & 55 & IRF3  Homo sapiens & interacts \\ \hline
% 42 & ORF4b  Middle East respiratory syndrome-related coronavirus & 60 & IRF3  Camelus dromedarius & interacts \\ \hline
% 42 & ORF4b  Middle East respiratory syndrome-related coronavirus & 64 & IRF3  Rattus norvegicus & interacts \\ \hline
% 42 & ORF4b  Middle East respiratory syndrome-related coronavirus & 66 & IRF3  Mus musculus & interacts \\ \hline
% 42 & ORF4b  Middle East respiratory syndrome-related coronavirus & 115 & IRF7  Homo sapiens & interacts \\ \hline
% 42 & ORF4b  Middle East respiratory syndrome-related coronavirus & 122 & IRF7  Camelus dromedarius & interacts \\ \hline
% 42 & ORF4b  Middle East respiratory syndrome-related coronavirus & 123 & IRF7  Rattus norvegicus & interacts \\ \hline
% 42 & ORF4b  Middle East respiratory syndrome-related coronavirus & 126 & IRF7  Mus musculus & interacts \\ \hline
% 42 & ORF4b  Middle East respiratory syndrome-related coronavirus & 231 & MAVS  Homo sapiens & interacts \\ \hline
% 42 & ORF4b  Middle East respiratory syndrome-related coronavirus & 239 & MAVS  Camelus dromedarius & interacts \\ \hline
% 42 & ORF4b  Middle East respiratory syndrome-related coronavirus & 243 & MAVS  Mus musculus & interacts \\ \hline
% 42 & ORF4b  Middle East respiratory syndrome-related coronavirus & 244 & MAVS  Rattus norvegicus & interacts \\ \hline
% 42 & ORF4b  Middle East respiratory syndrome-related coronavirus & 216 & TBK1  Homo sapiens & interacts \\ \hline
% 42 & ORF4b  Middle East respiratory syndrome-related coronavirus & 227 & TBK1  Mus musculus & interacts \\ \hline
% 42 & ORF4b  Middle East respiratory syndrome-related coronavirus & 228 & TBK1  Camelus dromedarius & interacts \\ \hline
% 42 & ORF4b  Middle East respiratory syndrome-related coronavirus & 229 & TBK1  Rattus norvegicus & interacts \\ \hline
% 43 & ORF4b  Human coronavirus 229E & 82 & Human coronavirus 229E & belongs \\ \hline
% 44 & nsp1  Severe acute respiratory syndrome coronavirus 2 & 45 & nsp1  Severe acute respiratory syndrome-related coronavirus & similar \\ \hline
% 44 & nsp1  Severe acute respiratory syndrome coronavirus 2 & 46 & nsp1  Middle East respiratory syndrome-related coronavirus & similar \\ \hline
% 44 & nsp1  Severe acute respiratory syndrome coronavirus 2 & 47 & nsp1  Human coronavirus HKU1 & similar \\ \hline
% 44 & nsp1  Severe acute respiratory syndrome coronavirus 2 & 79 & Severe acute respiratory syndrome coronavirus 2 & belongs \\ \hline
% 45 & nsp1  Severe acute respiratory syndrome-related coronavirus & 46 & nsp1  Middle East respiratory syndrome-related coronavirus & similar \\ \hline
% 45 & nsp1  Severe acute respiratory syndrome-related coronavirus & 47 & nsp1  Human coronavirus HKU1 & similar \\ \hline
% 45 & nsp1  Severe acute respiratory syndrome-related coronavirus & 80 & Severe acute respiratory syndrome-related coronavirus & belongs \\ \hline
% 45 & nsp1  Severe acute respiratory syndrome-related coronavirus & 172 & STAT2  Homo sapiens & interacts \\ \hline
% 45 & nsp1  Severe acute respiratory syndrome-related coronavirus & 175 & STAT2  Felis catus & interacts \\ \hline
% 45 & nsp1  Severe acute respiratory syndrome-related coronavirus & 178 & STAT2  Rhinolophus ferrumequinum & interacts \\ \hline
% 45 & nsp1  Severe acute respiratory syndrome-related coronavirus & 183 & STAT2  Rattus norvegicus & interacts \\ \hline
% 45 & nsp1  Severe acute respiratory syndrome-related coronavirus & 185 & STAT2  Mus musculus & interacts \\ \hline
% 46 & nsp1  Middle East respiratory syndrome-related coronavirus & 47 & nsp1  Human coronavirus HKU1 & similar \\ \hline
% 46 & nsp1  Middle East respiratory syndrome-related coronavirus & 78 & Middle East respiratory syndrome-related coronavirus & belongs \\ \hline
% 47 & nsp1  Human coronavirus HKU1 & 77 & Human coronavirus HKU1 & belongs \\ \hline
% 48 & spike  Human coronavirus OC43 & 49 & spike  Human coronavirus HKU1 & similar \\ \hline
% 48 & spike  Human coronavirus OC43 & 50 & spike  Middle East respiratory syndrome-related coronavirus & similar \\ \hline
% 48 & spike  Human coronavirus OC43 & 51 & spike  Severe acute respiratory syndrome coronavirus 2 & similar \\ \hline
% 48 & spike  Human coronavirus OC43 & 52 & spike  Severe acute respiratory syndrome-related coronavirus & similar \\ \hline
% 48 & spike  Human coronavirus OC43 & 53 & spike  Human coronavirus NL63 & similar \\ \hline
% 48 & spike  Human coronavirus OC43 & 54 & spike  Human coronavirus 229E & similar \\ \hline
% 48 & spike  Human coronavirus OC43 & 76 & Human coronavirus OC43 & belongs \\ \hline
% 49 & spike  Human coronavirus HKU1 & 50 & spike  Middle East respiratory syndrome-related coronavirus & similar \\ \hline
% 49 & spike  Human coronavirus HKU1 & 51 & spike  Severe acute respiratory syndrome coronavirus 2 & similar \\ \hline
% 49 & spike  Human coronavirus HKU1 & 52 & spike  Severe acute respiratory syndrome-related coronavirus & similar \\ \hline
% 49 & spike  Human coronavirus HKU1 & 53 & spike  Human coronavirus NL63 & similar \\ \hline
% 49 & spike  Human coronavirus HKU1 & 54 & spike  Human coronavirus 229E & similar \\ \hline
% 49 & spike  Human coronavirus HKU1 & 77 & Human coronavirus HKU1 & belongs \\ \hline
% 50 & spike  Middle East respiratory syndrome-related coronavirus & 51 & spike  Severe acute respiratory syndrome coronavirus 2 & similar \\ \hline
% 50 & spike  Middle East respiratory syndrome-related coronavirus & 52 & spike  Severe acute respiratory syndrome-related coronavirus & similar \\ \hline
% 50 & spike  Middle East respiratory syndrome-related coronavirus & 53 & spike  Human coronavirus NL63 & similar \\ \hline
% 50 & spike  Middle East respiratory syndrome-related coronavirus & 54 & spike  Human coronavirus 229E & similar \\ \hline
% 50 & spike  Middle East respiratory syndrome-related coronavirus & 78 & Middle East respiratory syndrome-related coronavirus & belongs \\ \hline
% 50 & spike  Middle East respiratory syndrome-related coronavirus & 156 & DPP4  Homo sapiens & interacts \\ \hline
% 50 & spike  Middle East respiratory syndrome-related coronavirus & 164 & DPP4  Rhinolophus ferrumequinum & interacts \\ \hline
% 50 & spike  Middle East respiratory syndrome-related coronavirus & 168 & DPP4  Camelus dromedarius & interacts \\ \hline
% 51 & spike  Severe acute respiratory syndrome coronavirus 2 & 52 & spike  Severe acute respiratory syndrome-related coronavirus & similar \\ \hline
% 51 & spike  Severe acute respiratory syndrome coronavirus 2 & 53 & spike  Human coronavirus NL63 & similar \\ \hline
% 51 & spike  Severe acute respiratory syndrome coronavirus 2 & 54 & spike  Human coronavirus 229E & similar \\ \hline
% 51 & spike  Severe acute respiratory syndrome coronavirus 2 & 79 & Severe acute respiratory syndrome coronavirus 2 & belongs \\ \hline
% 51 & spike  Severe acute respiratory syndrome coronavirus 2 & 83 & ACE2  Homo sapiens & interacts \\ \hline
% 51 & spike  Severe acute respiratory syndrome coronavirus 2 & 85 & ACE2  Macaca mulatta & interacts \\ \hline
% 51 & spike  Severe acute respiratory syndrome coronavirus 2 & 89 & ACE2  Felis catus & interacts \\ \hline
% 51 & spike  Severe acute respiratory syndrome coronavirus 2 & 91 & ACE2  Mesocricetus auratus & interacts \\ \hline
% 51 & spike  Severe acute respiratory syndrome coronavirus 2 & 97 & ACE2  Rhinolophus ferrumequinum & interacts \\ \hline
% 51 & spike  Severe acute respiratory syndrome coronavirus 2 & 98 & ACE2  Canis lupus familiaris & interacts \\ \hline
% 52 & spike  Severe acute respiratory syndrome-related coronavirus & 53 & spike  Human coronavirus NL63 & similar \\ \hline
% 52 & spike  Severe acute respiratory syndrome-related coronavirus & 54 & spike  Human coronavirus 229E & similar \\ \hline
% 52 & spike  Severe acute respiratory syndrome-related coronavirus & 80 & Severe acute respiratory syndrome-related coronavirus & belongs \\ \hline
% 52 & spike  Severe acute respiratory syndrome-related coronavirus & 83 & ACE2  Homo sapiens & interacts \\ \hline
% 52 & spike  Severe acute respiratory syndrome-related coronavirus & 89 & ACE2  Felis catus & interacts \\ \hline
% 52 & spike  Severe acute respiratory syndrome-related coronavirus & 97 & ACE2  Rhinolophus ferrumequinum & interacts \\ \hline
% 53 & spike  Human coronavirus NL63 & 54 & spike  Human coronavirus 229E & similar \\ \hline
% 53 & spike  Human coronavirus NL63 & 81 & Human coronavirus NL63 & belongs \\ \hline
% 53 & spike  Human coronavirus NL63 & 83 & ACE2  Homo sapiens & interacts \\ \hline
% 53 & spike  Human coronavirus NL63 & 97 & ACE2  Rhinolophus ferrumequinum & interacts \\ \hline
% 54 & spike  Human coronavirus 229E & 82 & Human coronavirus 229E & belongs \\ \hline
% 55 & IRF3  Homo sapiens & 56 & IRF3  Pan troglodytes & similar \\ \hline
% 55 & IRF3  Homo sapiens & 57 & IRF3  Macaca mulatta & similar \\ \hline
% 55 & IRF3  Homo sapiens & 58 & IRF3  Ictidomys tridecemlineatus & similar \\ \hline
% 55 & IRF3  Homo sapiens & 59 & IRF3  Rhinolophus ferrumequinum & similar \\ \hline
% 55 & IRF3  Homo sapiens & 60 & IRF3  Camelus dromedarius & similar \\ \hline
% 55 & IRF3  Homo sapiens & 61 & IRF3  Ovis aries & similar \\ \hline
% 55 & IRF3  Homo sapiens & 62 & IRF3  Bos taurus & similar \\ \hline
% 55 & IRF3  Homo sapiens & 63 & IRF3  Equus caballus & similar \\ \hline
% 55 & IRF3  Homo sapiens & 64 & IRF3  Rattus norvegicus & similar \\ \hline
% 55 & IRF3  Homo sapiens & 65 & IRF3  Mesocricetus auratus & similar \\ \hline
% 55 & IRF3  Homo sapiens & 66 & IRF3  Mus musculus & similar \\ \hline
% 55 & IRF3  Homo sapiens & 67 & IRF3  Canis lupus familiaris & similar \\ \hline
% 55 & IRF3  Homo sapiens & 68 & IRF3  Gallus gallus & similar \\ \hline
% 55 & IRF3  Homo sapiens & 192 & Homo sapiens & belongs \\ \hline
% 55 & IRF3  Homo sapiens & 154 & ORF3b  Severe acute respiratory syndrome-related coronavirus & interacts \\ \hline
% 55 & IRF3  Homo sapiens & 170 & ORF6  Severe acute respiratory syndrome-related coronavirus & interacts \\ \hline
% 55 & IRF3  Homo sapiens & 171 & ORF6  Severe acute respiratory syndrome coronavirus 2 & interacts \\ \hline
% 55 & IRF3  Homo sapiens & 188 & PLpro  Severe acute respiratory syndrome-related coronavirus & interacts \\ \hline
% 55 & IRF3  Homo sapiens & 187 & PLpro  Middle East respiratory syndrome-related coronavirus & interacts \\ \hline
% 55 & IRF3  Homo sapiens & 71 & N protein  Severe acute respiratory syndrome-related coronavirus & interacts \\ \hline
% 55 & IRF3  Homo sapiens & 211 & M protein  Severe acute respiratory syndrome-related coronavirus & interacts \\ \hline
% 55 & IRF3  Homo sapiens & 209 & M protein  Middle East respiratory syndrome-related coronavirus & interacts \\ \hline
% 55 & IRF3  Homo sapiens & 100 & ORF4a  Middle East respiratory syndrome-related coronavirus & interacts \\ \hline
% 56 & IRF3  Pan troglodytes & 57 & IRF3  Macaca mulatta & similar \\ \hline
% 56 & IRF3  Pan troglodytes & 58 & IRF3  Ictidomys tridecemlineatus & similar \\ \hline
% 56 & IRF3  Pan troglodytes & 59 & IRF3  Rhinolophus ferrumequinum & similar \\ \hline
% 56 & IRF3  Pan troglodytes & 60 & IRF3  Camelus dromedarius & similar \\ \hline
% 56 & IRF3  Pan troglodytes & 61 & IRF3  Ovis aries & similar \\ \hline
% 56 & IRF3  Pan troglodytes & 62 & IRF3  Bos taurus & similar \\ \hline
% 56 & IRF3  Pan troglodytes & 63 & IRF3  Equus caballus & similar \\ \hline
% 56 & IRF3  Pan troglodytes & 64 & IRF3  Rattus norvegicus & similar \\ \hline
% 56 & IRF3  Pan troglodytes & 65 & IRF3  Mesocricetus auratus & similar \\ \hline
% 56 & IRF3  Pan troglodytes & 66 & IRF3  Mus musculus & similar \\ \hline
% 56 & IRF3  Pan troglodytes & 67 & IRF3  Canis lupus familiaris & similar \\ \hline
% 56 & IRF3  Pan troglodytes & 68 & IRF3  Gallus gallus & similar \\ \hline
% 56 & IRF3  Pan troglodytes & 200 & Pan troglodytes & belongs \\ \hline
% 57 & IRF3  Macaca mulatta & 58 & IRF3  Ictidomys tridecemlineatus & similar \\ \hline
% 57 & IRF3  Macaca mulatta & 59 & IRF3  Rhinolophus ferrumequinum & similar \\ \hline
% 57 & IRF3  Macaca mulatta & 60 & IRF3  Camelus dromedarius & similar \\ \hline
% 57 & IRF3  Macaca mulatta & 61 & IRF3  Ovis aries & similar \\ \hline
% 57 & IRF3  Macaca mulatta & 62 & IRF3  Bos taurus & similar \\ \hline
% 57 & IRF3  Macaca mulatta & 63 & IRF3  Equus caballus & similar \\ \hline
% 57 & IRF3  Macaca mulatta & 64 & IRF3  Rattus norvegicus & similar \\ \hline
% 57 & IRF3  Macaca mulatta & 65 & IRF3  Mesocricetus auratus & similar \\ \hline
% 57 & IRF3  Macaca mulatta & 66 & IRF3  Mus musculus & similar \\ \hline
% 57 & IRF3  Macaca mulatta & 67 & IRF3  Canis lupus familiaris & similar \\ \hline
% 57 & IRF3  Macaca mulatta & 68 & IRF3  Gallus gallus & similar \\ \hline
% 57 & IRF3  Macaca mulatta & 204 & Macaca mulatta & belongs \\ \hline
% 58 & IRF3  Ictidomys tridecemlineatus & 59 & IRF3  Rhinolophus ferrumequinum & similar \\ \hline
% 58 & IRF3  Ictidomys tridecemlineatus & 60 & IRF3  Camelus dromedarius & similar \\ \hline
% 58 & IRF3  Ictidomys tridecemlineatus & 61 & IRF3  Ovis aries & similar \\ \hline
% 58 & IRF3  Ictidomys tridecemlineatus & 62 & IRF3  Bos taurus & similar \\ \hline
% 58 & IRF3  Ictidomys tridecemlineatus & 63 & IRF3  Equus caballus & similar \\ \hline
% 58 & IRF3  Ictidomys tridecemlineatus & 64 & IRF3  Rattus norvegicus & similar \\ \hline
% 58 & IRF3  Ictidomys tridecemlineatus & 65 & IRF3  Mesocricetus auratus & similar \\ \hline
% 58 & IRF3  Ictidomys tridecemlineatus & 66 & IRF3  Mus musculus & similar \\ \hline
% 58 & IRF3  Ictidomys tridecemlineatus & 67 & IRF3  Canis lupus familiaris & similar \\ \hline
% 58 & IRF3  Ictidomys tridecemlineatus & 68 & IRF3  Gallus gallus & similar \\ \hline
% 58 & IRF3  Ictidomys tridecemlineatus & 198 & Ictidomys tridecemlineatus & belongs \\ \hline
% 59 & IRF3  Rhinolophus ferrumequinum & 60 & IRF3  Camelus dromedarius & similar \\ \hline
% 59 & IRF3  Rhinolophus ferrumequinum & 61 & IRF3  Ovis aries & similar \\ \hline
% 59 & IRF3  Rhinolophus ferrumequinum & 62 & IRF3  Bos taurus & similar \\ \hline
% 59 & IRF3  Rhinolophus ferrumequinum & 63 & IRF3  Equus caballus & similar \\ \hline
% 59 & IRF3  Rhinolophus ferrumequinum & 64 & IRF3  Rattus norvegicus & similar \\ \hline
% 59 & IRF3  Rhinolophus ferrumequinum & 65 & IRF3  Mesocricetus auratus & similar \\ \hline
% 59 & IRF3  Rhinolophus ferrumequinum & 66 & IRF3  Mus musculus & similar \\ \hline
% 59 & IRF3  Rhinolophus ferrumequinum & 67 & IRF3  Canis lupus familiaris & similar \\ \hline
% 59 & IRF3  Rhinolophus ferrumequinum & 68 & IRF3  Gallus gallus & similar \\ \hline
% 59 & IRF3  Rhinolophus ferrumequinum & 207 & Rhinolophus ferrumequinum & belongs \\ \hline
% 59 & IRF3  Rhinolophus ferrumequinum & 154 & ORF3b  Severe acute respiratory syndrome-related coronavirus & interacts \\ \hline
% 59 & IRF3  Rhinolophus ferrumequinum & 170 & ORF6  Severe acute respiratory syndrome-related coronavirus & interacts \\ \hline
% 59 & IRF3  Rhinolophus ferrumequinum & 188 & PLpro  Severe acute respiratory syndrome-related coronavirus & interacts \\ \hline
% 59 & IRF3  Rhinolophus ferrumequinum & 71 & N protein  Severe acute respiratory syndrome-related coronavirus & interacts \\ \hline
% 59 & IRF3  Rhinolophus ferrumequinum & 211 & M protein  Severe acute respiratory syndrome-related coronavirus & interacts \\ \hline
% 60 & IRF3  Camelus dromedarius & 61 & IRF3  Ovis aries & similar \\ \hline
% 60 & IRF3  Camelus dromedarius & 62 & IRF3  Bos taurus & similar \\ \hline
% 60 & IRF3  Camelus dromedarius & 63 & IRF3  Equus caballus & similar \\ \hline
% 60 & IRF3  Camelus dromedarius & 64 & IRF3  Rattus norvegicus & similar \\ \hline
% 60 & IRF3  Camelus dromedarius & 65 & IRF3  Mesocricetus auratus & similar \\ \hline
% 60 & IRF3  Camelus dromedarius & 66 & IRF3  Mus musculus & similar \\ \hline
% 60 & IRF3  Camelus dromedarius & 67 & IRF3  Canis lupus familiaris & similar \\ \hline
% 60 & IRF3  Camelus dromedarius & 68 & IRF3  Gallus gallus & similar \\ \hline
% 60 & IRF3  Camelus dromedarius & 196 & Camelus dromedarius & belongs \\ \hline
% 60 & IRF3  Camelus dromedarius & 187 & PLpro  Middle East respiratory syndrome-related coronavirus & interacts \\ \hline
% 60 & IRF3  Camelus dromedarius & 209 & M protein  Middle East respiratory syndrome-related coronavirus & interacts \\ \hline
% 60 & IRF3  Camelus dromedarius & 100 & ORF4a  Middle East respiratory syndrome-related coronavirus & interacts \\ \hline
% 61 & IRF3  Ovis aries & 62 & IRF3  Bos taurus & similar \\ \hline
% 61 & IRF3  Ovis aries & 63 & IRF3  Equus caballus & similar \\ \hline
% 61 & IRF3  Ovis aries & 64 & IRF3  Rattus norvegicus & similar \\ \hline
% 61 & IRF3  Ovis aries & 65 & IRF3  Mesocricetus auratus & similar \\ \hline
% 61 & IRF3  Ovis aries & 66 & IRF3  Mus musculus & similar \\ \hline
% 61 & IRF3  Ovis aries & 67 & IRF3  Canis lupus familiaris & similar \\ \hline
% 61 & IRF3  Ovis aries & 68 & IRF3  Gallus gallus & similar \\ \hline
% 61 & IRF3  Ovis aries & 205 & Ovis aries & belongs \\ \hline
% 62 & IRF3  Bos taurus & 63 & IRF3  Equus caballus & similar \\ \hline
% 62 & IRF3  Bos taurus & 64 & IRF3  Rattus norvegicus & similar \\ \hline
% 62 & IRF3  Bos taurus & 65 & IRF3  Mesocricetus auratus & similar \\ \hline
% 62 & IRF3  Bos taurus & 66 & IRF3  Mus musculus & similar \\ \hline
% 62 & IRF3  Bos taurus & 67 & IRF3  Canis lupus familiaris & similar \\ \hline
% 62 & IRF3  Bos taurus & 68 & IRF3  Gallus gallus & similar \\ \hline
% 62 & IRF3  Bos taurus & 199 & Bos taurus & belongs \\ \hline
% 63 & IRF3  Equus caballus & 64 & IRF3  Rattus norvegicus & similar \\ \hline
% 63 & IRF3  Equus caballus & 65 & IRF3  Mesocricetus auratus & similar \\ \hline
% 63 & IRF3  Equus caballus & 66 & IRF3  Mus musculus & similar \\ \hline
% 63 & IRF3  Equus caballus & 67 & IRF3  Canis lupus familiaris & similar \\ \hline
% 63 & IRF3  Equus caballus & 68 & IRF3  Gallus gallus & similar \\ \hline
% 63 & IRF3  Equus caballus & 203 & Equus caballus & belongs \\ \hline
% 64 & IRF3  Rattus norvegicus & 65 & IRF3  Mesocricetus auratus & similar \\ \hline
% 64 & IRF3  Rattus norvegicus & 66 & IRF3  Mus musculus & similar \\ \hline
% 64 & IRF3  Rattus norvegicus & 67 & IRF3  Canis lupus familiaris & similar \\ \hline
% 64 & IRF3  Rattus norvegicus & 68 & IRF3  Gallus gallus & similar \\ \hline
% 64 & IRF3  Rattus norvegicus & 194 & Rattus norvegicus & belongs \\ \hline
% 64 & IRF3  Rattus norvegicus & 154 & ORF3b  Severe acute respiratory syndrome-related coronavirus & interacts \\ \hline
% 64 & IRF3  Rattus norvegicus & 170 & ORF6  Severe acute respiratory syndrome-related coronavirus & interacts \\ \hline
% 64 & IRF3  Rattus norvegicus & 171 & ORF6  Severe acute respiratory syndrome coronavirus 2 & interacts \\ \hline
% 64 & IRF3  Rattus norvegicus & 188 & PLpro  Severe acute respiratory syndrome-related coronavirus & interacts \\ \hline
% 64 & IRF3  Rattus norvegicus & 187 & PLpro  Middle East respiratory syndrome-related coronavirus & interacts \\ \hline
% 64 & IRF3  Rattus norvegicus & 71 & N protein  Severe acute respiratory syndrome-related coronavirus & interacts \\ \hline
% 64 & IRF3  Rattus norvegicus & 211 & M protein  Severe acute respiratory syndrome-related coronavirus & interacts \\ \hline
% 64 & IRF3  Rattus norvegicus & 209 & M protein  Middle East respiratory syndrome-related coronavirus & interacts \\ \hline
% 64 & IRF3  Rattus norvegicus & 100 & ORF4a  Middle East respiratory syndrome-related coronavirus & interacts \\ \hline
% 65 & IRF3  Mesocricetus auratus & 66 & IRF3  Mus musculus & similar \\ \hline
% 65 & IRF3  Mesocricetus auratus & 67 & IRF3  Canis lupus familiaris & similar \\ \hline
% 65 & IRF3  Mesocricetus auratus & 68 & IRF3  Gallus gallus & similar \\ \hline
% 65 & IRF3  Mesocricetus auratus & 208 & Mesocricetus auratus & belongs \\ \hline
% 65 & IRF3  Mesocricetus auratus & 171 & ORF6  Severe acute respiratory syndrome coronavirus 2 & interacts \\ \hline
% 66 & IRF3  Mus musculus & 67 & IRF3  Canis lupus familiaris & similar \\ \hline
% 66 & IRF3  Mus musculus & 68 & IRF3  Gallus gallus & similar \\ \hline
% 66 & IRF3  Mus musculus & 193 & Mus musculus & belongs \\ \hline
% 66 & IRF3  Mus musculus & 154 & ORF3b  Severe acute respiratory syndrome-related coronavirus & interacts \\ \hline
% 66 & IRF3  Mus musculus & 170 & ORF6  Severe acute respiratory syndrome-related coronavirus & interacts \\ \hline
% 66 & IRF3  Mus musculus & 171 & ORF6  Severe acute respiratory syndrome coronavirus 2 & interacts \\ \hline
% 66 & IRF3  Mus musculus & 188 & PLpro  Severe acute respiratory syndrome-related coronavirus & interacts \\ \hline
% 66 & IRF3  Mus musculus & 187 & PLpro  Middle East respiratory syndrome-related coronavirus & interacts \\ \hline
% 66 & IRF3  Mus musculus & 71 & N protein  Severe acute respiratory syndrome-related coronavirus & interacts \\ \hline
% 66 & IRF3  Mus musculus & 211 & M protein  Severe acute respiratory syndrome-related coronavirus & interacts \\ \hline
% 66 & IRF3  Mus musculus & 209 & M protein  Middle East respiratory syndrome-related coronavirus & interacts \\ \hline
% 66 & IRF3  Mus musculus & 100 & ORF4a  Middle East respiratory syndrome-related coronavirus & interacts \\ \hline
% 67 & IRF3  Canis lupus familiaris & 68 & IRF3  Gallus gallus & similar \\ \hline
% 67 & IRF3  Canis lupus familiaris & 195 & Canis lupus familiaris & belongs \\ \hline
% 67 & IRF3  Canis lupus familiaris & 171 & ORF6  Severe acute respiratory syndrome coronavirus 2 & interacts \\ \hline
% 68 & IRF3  Gallus gallus & 201 & Gallus gallus & belongs \\ \hline
% 69 & N protein  Middle East respiratory syndrome-related coronavirus & 70 & N protein  Severe acute respiratory syndrome coronavirus 2 & similar \\ \hline
% 69 & N protein  Middle East respiratory syndrome-related coronavirus & 71 & N protein  Severe acute respiratory syndrome-related coronavirus & similar \\ \hline
% 69 & N protein  Middle East respiratory syndrome-related coronavirus & 72 & N protein  Human coronavirus HKU1 & similar \\ \hline
% 69 & N protein  Middle East respiratory syndrome-related coronavirus & 73 & N protein  Human coronavirus OC43 & similar \\ \hline
% 69 & N protein  Middle East respiratory syndrome-related coronavirus & 74 & N protein  Human coronavirus 229E & similar \\ \hline
% 69 & N protein  Middle East respiratory syndrome-related coronavirus & 75 & N protein  Human coronavirus NL63 & similar \\ \hline
% 69 & N protein  Middle East respiratory syndrome-related coronavirus & 78 & Middle East respiratory syndrome-related coronavirus & belongs \\ \hline
% 70 & N protein  Severe acute respiratory syndrome coronavirus 2 & 71 & N protein  Severe acute respiratory syndrome-related coronavirus & similar \\ \hline
% 70 & N protein  Severe acute respiratory syndrome coronavirus 2 & 72 & N protein  Human coronavirus HKU1 & similar \\ \hline
% 70 & N protein  Severe acute respiratory syndrome coronavirus 2 & 73 & N protein  Human coronavirus OC43 & similar \\ \hline
% 70 & N protein  Severe acute respiratory syndrome coronavirus 2 & 74 & N protein  Human coronavirus 229E & similar \\ \hline
% 70 & N protein  Severe acute respiratory syndrome coronavirus 2 & 75 & N protein  Human coronavirus NL63 & similar \\ \hline
% 70 & N protein  Severe acute respiratory syndrome coronavirus 2 & 79 & Severe acute respiratory syndrome coronavirus 2 & belongs \\ \hline
% 71 & N protein  Severe acute respiratory syndrome-related coronavirus & 72 & N protein  Human coronavirus HKU1 & similar \\ \hline
% 71 & N protein  Severe acute respiratory syndrome-related coronavirus & 73 & N protein  Human coronavirus OC43 & similar \\ \hline
% 71 & N protein  Severe acute respiratory syndrome-related coronavirus & 74 & N protein  Human coronavirus 229E & similar \\ \hline
% 71 & N protein  Severe acute respiratory syndrome-related coronavirus & 75 & N protein  Human coronavirus NL63 & similar \\ \hline
% 71 & N protein  Severe acute respiratory syndrome-related coronavirus & 80 & Severe acute respiratory syndrome-related coronavirus & belongs \\ \hline
% 71 & N protein  Severe acute respiratory syndrome-related coronavirus & 231 & MAVS  Homo sapiens & interacts \\ \hline
% 71 & N protein  Severe acute respiratory syndrome-related coronavirus & 237 & MAVS  Rhinolophus ferrumequinum & interacts \\ \hline
% 71 & N protein  Severe acute respiratory syndrome-related coronavirus & 238 & MAVS  Felis catus & interacts \\ \hline
% 71 & N protein  Severe acute respiratory syndrome-related coronavirus & 243 & MAVS  Mus musculus & interacts \\ \hline
% 71 & N protein  Severe acute respiratory syndrome-related coronavirus & 244 & MAVS  Rattus norvegicus & interacts \\ \hline
% 71 & N protein  Severe acute respiratory syndrome-related coronavirus & 129 & MDA5  Homo sapiens & interacts \\ \hline
% 71 & N protein  Severe acute respiratory syndrome-related coronavirus & 134 & MDA5  Felis catus & interacts \\ \hline
% 71 & N protein  Severe acute respiratory syndrome-related coronavirus & 136 & MDA5  Mus musculus & interacts \\ \hline
% 71 & N protein  Severe acute respiratory syndrome-related coronavirus & 137 & MDA5  Rattus norvegicus & interacts \\ \hline
% 72 & N protein  Human coronavirus HKU1 & 73 & N protein  Human coronavirus OC43 & similar \\ \hline
% 72 & N protein  Human coronavirus HKU1 & 74 & N protein  Human coronavirus 229E & similar \\ \hline
% 72 & N protein  Human coronavirus HKU1 & 75 & N protein  Human coronavirus NL63 & similar \\ \hline
% 72 & N protein  Human coronavirus HKU1 & 77 & Human coronavirus HKU1 & belongs \\ \hline
% 73 & N protein  Human coronavirus OC43 & 74 & N protein  Human coronavirus 229E & similar \\ \hline
% 73 & N protein  Human coronavirus OC43 & 75 & N protein  Human coronavirus NL63 & similar \\ \hline
% 73 & N protein  Human coronavirus OC43 & 76 & Human coronavirus OC43 & belongs \\ \hline
% 74 & N protein  Human coronavirus 229E & 75 & N protein  Human coronavirus NL63 & similar \\ \hline
% 74 & N protein  Human coronavirus 229E & 82 & Human coronavirus 229E & belongs \\ \hline
% 75 & N protein  Human coronavirus NL63 & 81 & Human coronavirus NL63 & belongs \\ \hline
% 76 & Human coronavirus OC43 & 190 & PLpro  Human coronavirus OC43 & belongs \\ \hline
% 76 & Human coronavirus OC43 & 213 & M protein  Human coronavirus OC43 & belongs \\ \hline
% 76 & Human coronavirus OC43 & 192 & Homo sapiens & infects \\ \hline
% 76 & Human coronavirus OC43 & 193 & Mus musculus & infects \\ \hline
% 76 & Human coronavirus OC43 & 194 & Rattus norvegicus & infects \\ \hline
% 76 & Human coronavirus OC43 & 199 & Bos taurus & infects \\ \hline
% 77 & Human coronavirus HKU1 & 191 & PLpro  Human coronavirus HKU1 & belongs \\ \hline
% 77 & Human coronavirus HKU1 & 210 & M protein  Human coronavirus HKU1 & belongs \\ \hline
% 77 & Human coronavirus HKU1 & 192 & Homo sapiens & infects \\ \hline
% 77 & Human coronavirus HKU1 & 193 & Mus musculus & infects \\ \hline
% 77 & Human coronavirus HKU1 & 194 & Rattus norvegicus & infects \\ \hline
% 77 & Human coronavirus HKU1 & 207 & Rhinolophus ferrumequinum & infects \\ \hline
% 78 & Middle East respiratory syndrome-related coronavirus & 100 & ORF4a  Middle East respiratory syndrome-related coronavirus & belongs \\ \hline
% 78 & Middle East respiratory syndrome-related coronavirus & 187 & PLpro  Middle East respiratory syndrome-related coronavirus & belongs \\ \hline
% 78 & Middle East respiratory syndrome-related coronavirus & 209 & M protein  Middle East respiratory syndrome-related coronavirus & belongs \\ \hline
% 78 & Middle East respiratory syndrome-related coronavirus & 192 & Homo sapiens & infects \\ \hline
% 78 & Middle East respiratory syndrome-related coronavirus & 196 & Camelus dromedarius & infects \\ \hline
% 78 & Middle East respiratory syndrome-related coronavirus & 207 & Rhinolophus ferrumequinum & infects \\ \hline
% 79 & Severe acute respiratory syndrome coronavirus 2 & 171 & ORF6  Severe acute respiratory syndrome coronavirus 2 & belongs \\ \hline
% 79 & Severe acute respiratory syndrome coronavirus 2 & 189 & PLpro  Severe acute respiratory syndrome coronavirus 2 & belongs \\ \hline
% 79 & Severe acute respiratory syndrome coronavirus 2 & 212 & M protein  Severe acute respiratory syndrome coronavirus 2 & belongs \\ \hline
% 79 & Severe acute respiratory syndrome coronavirus 2 & 192 & Homo sapiens & infects \\ \hline
% 79 & Severe acute respiratory syndrome coronavirus 2 & 195 & Canis lupus familiaris & infects \\ \hline
% 79 & Severe acute respiratory syndrome coronavirus 2 & 197 & Felis catus & infects \\ \hline
% 79 & Severe acute respiratory syndrome coronavirus 2 & 204 & Macaca mulatta & infects \\ \hline
% 79 & Severe acute respiratory syndrome coronavirus 2 & 207 & Rhinolophus ferrumequinum & infects \\ \hline
% 79 & Severe acute respiratory syndrome coronavirus 2 & 208 & Mesocricetus auratus & infects \\ \hline
% 80 & Severe acute respiratory syndrome-related coronavirus & 154 & ORF3b  Severe acute respiratory syndrome-related coronavirus & belongs \\ \hline
% 80 & Severe acute respiratory syndrome-related coronavirus & 170 & ORF6  Severe acute respiratory syndrome-related coronavirus & belongs \\ \hline
% 80 & Severe acute respiratory syndrome-related coronavirus & 188 & PLpro  Severe acute respiratory syndrome-related coronavirus & belongs \\ \hline
% 80 & Severe acute respiratory syndrome-related coronavirus & 211 & M protein  Severe acute respiratory syndrome-related coronavirus & belongs \\ \hline
% 80 & Severe acute respiratory syndrome-related coronavirus & 192 & Homo sapiens & infects \\ \hline
% 80 & Severe acute respiratory syndrome-related coronavirus & 197 & Felis catus & infects \\ \hline
% 80 & Severe acute respiratory syndrome-related coronavirus & 207 & Rhinolophus ferrumequinum & infects \\ \hline
% 81 & Human coronavirus NL63 & 155 & ORF3b  Human coronavirus NL63 & belongs \\ \hline
% 81 & Human coronavirus NL63 & 214 & M protein  Human coronavirus NL63 & belongs \\ \hline
% 81 & Human coronavirus NL63 & 192 & Homo sapiens & infects \\ \hline
% 81 & Human coronavirus NL63 & 207 & Rhinolophus ferrumequinum & infects \\ \hline
% 82 & Human coronavirus 229E & 101 & ORF4a  Human coronavirus 229E & belongs \\ \hline
% 82 & Human coronavirus 229E & 215 & M protein  Human coronavirus 229E & belongs \\ \hline
% 82 & Human coronavirus 229E & 192 & Homo sapiens & infects \\ \hline
% 82 & Human coronavirus 229E & 207 & Rhinolophus ferrumequinum & infects \\ \hline
% 83 & ACE2  Homo sapiens & 84 & ACE2  Pan troglodytes & similar \\ \hline
% 83 & ACE2  Homo sapiens & 85 & ACE2  Macaca mulatta & similar \\ \hline
% 83 & ACE2  Homo sapiens & 86 & ACE2  Ictidomys tridecemlineatus & similar \\ \hline
% 83 & ACE2  Homo sapiens & 87 & ACE2  Oryctolagus cuniculus & similar \\ \hline
% 83 & ACE2  Homo sapiens & 88 & ACE2  Equus caballus & similar \\ \hline
% 83 & ACE2  Homo sapiens & 89 & ACE2  Felis catus & similar \\ \hline
% 83 & ACE2  Homo sapiens & 90 & ACE2  Camelus dromedarius & similar \\ \hline
% 83 & ACE2  Homo sapiens & 91 & ACE2  Mesocricetus auratus & similar \\ \hline
% 83 & ACE2  Homo sapiens & 92 & ACE2  Sus scrofa domesticus & similar \\ \hline
% 83 & ACE2  Homo sapiens & 93 & ACE2  Ovis aries & similar \\ \hline
% 83 & ACE2  Homo sapiens & 94 & ACE2  Mus musculus & similar \\ \hline
% 83 & ACE2  Homo sapiens & 95 & ACE2  Bos taurus & similar \\ \hline
% 83 & ACE2  Homo sapiens & 96 & ACE2  Rattus norvegicus & similar \\ \hline
% 83 & ACE2  Homo sapiens & 97 & ACE2  Rhinolophus ferrumequinum & similar \\ \hline
% 83 & ACE2  Homo sapiens & 98 & ACE2  Canis lupus familiaris & similar \\ \hline
% 83 & ACE2  Homo sapiens & 99 & ACE2  Gallus gallus & similar \\ \hline
% 83 & ACE2  Homo sapiens & 192 & Homo sapiens & belongs \\ \hline
% 84 & ACE2  Pan troglodytes & 85 & ACE2  Macaca mulatta & similar \\ \hline
% 84 & ACE2  Pan troglodytes & 86 & ACE2  Ictidomys tridecemlineatus & similar \\ \hline
% 84 & ACE2  Pan troglodytes & 87 & ACE2  Oryctolagus cuniculus & similar \\ \hline
% 84 & ACE2  Pan troglodytes & 88 & ACE2  Equus caballus & similar \\ \hline
% 84 & ACE2  Pan troglodytes & 89 & ACE2  Felis catus & similar \\ \hline
% 84 & ACE2  Pan troglodytes & 90 & ACE2  Camelus dromedarius & similar \\ \hline
% 84 & ACE2  Pan troglodytes & 91 & ACE2  Mesocricetus auratus & similar \\ \hline
% 84 & ACE2  Pan troglodytes & 92 & ACE2  Sus scrofa domesticus & similar \\ \hline
% 84 & ACE2  Pan troglodytes & 93 & ACE2  Ovis aries & similar \\ \hline
% 84 & ACE2  Pan troglodytes & 94 & ACE2  Mus musculus & similar \\ \hline
% 84 & ACE2  Pan troglodytes & 95 & ACE2  Bos taurus & similar \\ \hline
% 84 & ACE2  Pan troglodytes & 96 & ACE2  Rattus norvegicus & similar \\ \hline
% 84 & ACE2  Pan troglodytes & 97 & ACE2  Rhinolophus ferrumequinum & similar \\ \hline
% 84 & ACE2  Pan troglodytes & 98 & ACE2  Canis lupus familiaris & similar \\ \hline
% 84 & ACE2  Pan troglodytes & 99 & ACE2  Gallus gallus & similar \\ \hline
% 84 & ACE2  Pan troglodytes & 200 & Pan troglodytes & belongs \\ \hline
% 85 & ACE2  Macaca mulatta & 86 & ACE2  Ictidomys tridecemlineatus & similar \\ \hline
% 85 & ACE2  Macaca mulatta & 87 & ACE2  Oryctolagus cuniculus & similar \\ \hline
% 85 & ACE2  Macaca mulatta & 88 & ACE2  Equus caballus & similar \\ \hline
% 85 & ACE2  Macaca mulatta & 89 & ACE2  Felis catus & similar \\ \hline
% 85 & ACE2  Macaca mulatta & 90 & ACE2  Camelus dromedarius & similar \\ \hline
% 85 & ACE2  Macaca mulatta & 91 & ACE2  Mesocricetus auratus & similar \\ \hline
% 85 & ACE2  Macaca mulatta & 92 & ACE2  Sus scrofa domesticus & similar \\ \hline
% 85 & ACE2  Macaca mulatta & 93 & ACE2  Ovis aries & similar \\ \hline
% 85 & ACE2  Macaca mulatta & 94 & ACE2  Mus musculus & similar \\ \hline
% 85 & ACE2  Macaca mulatta & 95 & ACE2  Bos taurus & similar \\ \hline
% 85 & ACE2  Macaca mulatta & 96 & ACE2  Rattus norvegicus & similar \\ \hline
% 85 & ACE2  Macaca mulatta & 97 & ACE2  Rhinolophus ferrumequinum & similar \\ \hline
% 85 & ACE2  Macaca mulatta & 98 & ACE2  Canis lupus familiaris & similar \\ \hline
% 85 & ACE2  Macaca mulatta & 99 & ACE2  Gallus gallus & similar \\ \hline
% 85 & ACE2  Macaca mulatta & 204 & Macaca mulatta & belongs \\ \hline
% 86 & ACE2  Ictidomys tridecemlineatus & 87 & ACE2  Oryctolagus cuniculus & similar \\ \hline
% 86 & ACE2  Ictidomys tridecemlineatus & 88 & ACE2  Equus caballus & similar \\ \hline
% 86 & ACE2  Ictidomys tridecemlineatus & 89 & ACE2  Felis catus & similar \\ \hline
% 86 & ACE2  Ictidomys tridecemlineatus & 90 & ACE2  Camelus dromedarius & similar \\ \hline
% 86 & ACE2  Ictidomys tridecemlineatus & 91 & ACE2  Mesocricetus auratus & similar \\ \hline
% 86 & ACE2  Ictidomys tridecemlineatus & 92 & ACE2  Sus scrofa domesticus & similar \\ \hline
% 86 & ACE2  Ictidomys tridecemlineatus & 93 & ACE2  Ovis aries & similar \\ \hline
% 86 & ACE2  Ictidomys tridecemlineatus & 94 & ACE2  Mus musculus & similar \\ \hline
% 86 & ACE2  Ictidomys tridecemlineatus & 95 & ACE2  Bos taurus & similar \\ \hline
% 86 & ACE2  Ictidomys tridecemlineatus & 96 & ACE2  Rattus norvegicus & similar \\ \hline
% 86 & ACE2  Ictidomys tridecemlineatus & 97 & ACE2  Rhinolophus ferrumequinum & similar \\ \hline
% 86 & ACE2  Ictidomys tridecemlineatus & 98 & ACE2  Canis lupus familiaris & similar \\ \hline
% 86 & ACE2  Ictidomys tridecemlineatus & 99 & ACE2  Gallus gallus & similar \\ \hline
% 86 & ACE2  Ictidomys tridecemlineatus & 198 & Ictidomys tridecemlineatus & belongs \\ \hline
% 87 & ACE2  Oryctolagus cuniculus & 88 & ACE2  Equus caballus & similar \\ \hline
% 87 & ACE2  Oryctolagus cuniculus & 89 & ACE2  Felis catus & similar \\ \hline
% 87 & ACE2  Oryctolagus cuniculus & 90 & ACE2  Camelus dromedarius & similar \\ \hline
% 87 & ACE2  Oryctolagus cuniculus & 91 & ACE2  Mesocricetus auratus & similar \\ \hline
% 87 & ACE2  Oryctolagus cuniculus & 92 & ACE2  Sus scrofa domesticus & similar \\ \hline
% 87 & ACE2  Oryctolagus cuniculus & 93 & ACE2  Ovis aries & similar \\ \hline
% 87 & ACE2  Oryctolagus cuniculus & 94 & ACE2  Mus musculus & similar \\ \hline
% 87 & ACE2  Oryctolagus cuniculus & 95 & ACE2  Bos taurus & similar \\ \hline
% 87 & ACE2  Oryctolagus cuniculus & 96 & ACE2  Rattus norvegicus & similar \\ \hline
% 87 & ACE2  Oryctolagus cuniculus & 97 & ACE2  Rhinolophus ferrumequinum & similar \\ \hline
% 87 & ACE2  Oryctolagus cuniculus & 98 & ACE2  Canis lupus familiaris & similar \\ \hline
% 87 & ACE2  Oryctolagus cuniculus & 99 & ACE2  Gallus gallus & similar \\ \hline
% 87 & ACE2  Oryctolagus cuniculus & 202 & Oryctolagus cuniculus & belongs \\ \hline
% 88 & ACE2  Equus caballus & 89 & ACE2  Felis catus & similar \\ \hline
% 88 & ACE2  Equus caballus & 90 & ACE2  Camelus dromedarius & similar \\ \hline
% 88 & ACE2  Equus caballus & 91 & ACE2  Mesocricetus auratus & similar \\ \hline
% 88 & ACE2  Equus caballus & 92 & ACE2  Sus scrofa domesticus & similar \\ \hline
% 88 & ACE2  Equus caballus & 93 & ACE2  Ovis aries & similar \\ \hline
% 88 & ACE2  Equus caballus & 94 & ACE2  Mus musculus & similar \\ \hline
% 88 & ACE2  Equus caballus & 95 & ACE2  Bos taurus & similar \\ \hline
% 88 & ACE2  Equus caballus & 96 & ACE2  Rattus norvegicus & similar \\ \hline
% 88 & ACE2  Equus caballus & 97 & ACE2  Rhinolophus ferrumequinum & similar \\ \hline
% 88 & ACE2  Equus caballus & 98 & ACE2  Canis lupus familiaris & similar \\ \hline
% 88 & ACE2  Equus caballus & 99 & ACE2  Gallus gallus & similar \\ \hline
% 88 & ACE2  Equus caballus & 203 & Equus caballus & belongs \\ \hline
% 89 & ACE2  Felis catus & 90 & ACE2  Camelus dromedarius & similar \\ \hline
% 89 & ACE2  Felis catus & 91 & ACE2  Mesocricetus auratus & similar \\ \hline
% 89 & ACE2  Felis catus & 92 & ACE2  Sus scrofa domesticus & similar \\ \hline
% 89 & ACE2  Felis catus & 93 & ACE2  Ovis aries & similar \\ \hline
% 89 & ACE2  Felis catus & 94 & ACE2  Mus musculus & similar \\ \hline
% 89 & ACE2  Felis catus & 95 & ACE2  Bos taurus & similar \\ \hline
% 89 & ACE2  Felis catus & 96 & ACE2  Rattus norvegicus & similar \\ \hline
% 89 & ACE2  Felis catus & 97 & ACE2  Rhinolophus ferrumequinum & similar \\ \hline
% 89 & ACE2  Felis catus & 98 & ACE2  Canis lupus familiaris & similar \\ \hline
% 89 & ACE2  Felis catus & 99 & ACE2  Gallus gallus & similar \\ \hline
% 89 & ACE2  Felis catus & 197 & Felis catus & belongs \\ \hline
% 90 & ACE2  Camelus dromedarius & 91 & ACE2  Mesocricetus auratus & similar \\ \hline
% 90 & ACE2  Camelus dromedarius & 92 & ACE2  Sus scrofa domesticus & similar \\ \hline
% 90 & ACE2  Camelus dromedarius & 93 & ACE2  Ovis aries & similar \\ \hline
% 90 & ACE2  Camelus dromedarius & 94 & ACE2  Mus musculus & similar \\ \hline
% 90 & ACE2  Camelus dromedarius & 95 & ACE2  Bos taurus & similar \\ \hline
% 90 & ACE2  Camelus dromedarius & 96 & ACE2  Rattus norvegicus & similar \\ \hline
% 90 & ACE2  Camelus dromedarius & 97 & ACE2  Rhinolophus ferrumequinum & similar \\ \hline
% 90 & ACE2  Camelus dromedarius & 98 & ACE2  Canis lupus familiaris & similar \\ \hline
% 90 & ACE2  Camelus dromedarius & 99 & ACE2  Gallus gallus & similar \\ \hline
% 90 & ACE2  Camelus dromedarius & 196 & Camelus dromedarius & belongs \\ \hline
% 91 & ACE2  Mesocricetus auratus & 92 & ACE2  Sus scrofa domesticus & similar \\ \hline
% 91 & ACE2  Mesocricetus auratus & 93 & ACE2  Ovis aries & similar \\ \hline
% 91 & ACE2  Mesocricetus auratus & 94 & ACE2  Mus musculus & similar \\ \hline
% 91 & ACE2  Mesocricetus auratus & 95 & ACE2  Bos taurus & similar \\ \hline
% 91 & ACE2  Mesocricetus auratus & 96 & ACE2  Rattus norvegicus & similar \\ \hline
% 91 & ACE2  Mesocricetus auratus & 97 & ACE2  Rhinolophus ferrumequinum & similar \\ \hline
% 91 & ACE2  Mesocricetus auratus & 98 & ACE2  Canis lupus familiaris & similar \\ \hline
% 91 & ACE2  Mesocricetus auratus & 99 & ACE2  Gallus gallus & similar \\ \hline
% 91 & ACE2  Mesocricetus auratus & 208 & Mesocricetus auratus & belongs \\ \hline
% 92 & ACE2  Sus scrofa domesticus & 93 & ACE2  Ovis aries & similar \\ \hline
% 92 & ACE2  Sus scrofa domesticus & 94 & ACE2  Mus musculus & similar \\ \hline
% 92 & ACE2  Sus scrofa domesticus & 95 & ACE2  Bos taurus & similar \\ \hline
% 92 & ACE2  Sus scrofa domesticus & 96 & ACE2  Rattus norvegicus & similar \\ \hline
% 92 & ACE2  Sus scrofa domesticus & 97 & ACE2  Rhinolophus ferrumequinum & similar \\ \hline
% 92 & ACE2  Sus scrofa domesticus & 98 & ACE2  Canis lupus familiaris & similar \\ \hline
% 92 & ACE2  Sus scrofa domesticus & 99 & ACE2  Gallus gallus & similar \\ \hline
% 92 & ACE2  Sus scrofa domesticus & 206 & Sus scrofa domesticus & belongs \\ \hline
% 93 & ACE2  Ovis aries & 94 & ACE2  Mus musculus & similar \\ \hline
% 93 & ACE2  Ovis aries & 95 & ACE2  Bos taurus & similar \\ \hline
% 93 & ACE2  Ovis aries & 96 & ACE2  Rattus norvegicus & similar \\ \hline
% 93 & ACE2  Ovis aries & 97 & ACE2  Rhinolophus ferrumequinum & similar \\ \hline
% 93 & ACE2  Ovis aries & 98 & ACE2  Canis lupus familiaris & similar \\ \hline
% 93 & ACE2  Ovis aries & 99 & ACE2  Gallus gallus & similar \\ \hline
% 93 & ACE2  Ovis aries & 205 & Ovis aries & belongs \\ \hline
% 94 & ACE2  Mus musculus & 95 & ACE2  Bos taurus & similar \\ \hline
% 94 & ACE2  Mus musculus & 96 & ACE2  Rattus norvegicus & similar \\ \hline
% 94 & ACE2  Mus musculus & 97 & ACE2  Rhinolophus ferrumequinum & similar \\ \hline
% 94 & ACE2  Mus musculus & 98 & ACE2  Canis lupus familiaris & similar \\ \hline
% 94 & ACE2  Mus musculus & 99 & ACE2  Gallus gallus & similar \\ \hline
% 94 & ACE2  Mus musculus & 193 & Mus musculus & belongs \\ \hline
% 95 & ACE2  Bos taurus & 96 & ACE2  Rattus norvegicus & similar \\ \hline
% 95 & ACE2  Bos taurus & 97 & ACE2  Rhinolophus ferrumequinum & similar \\ \hline
% 95 & ACE2  Bos taurus & 98 & ACE2  Canis lupus familiaris & similar \\ \hline
% 95 & ACE2  Bos taurus & 99 & ACE2  Gallus gallus & similar \\ \hline
% 95 & ACE2  Bos taurus & 199 & Bos taurus & belongs \\ \hline
% 96 & ACE2  Rattus norvegicus & 97 & ACE2  Rhinolophus ferrumequinum & similar \\ \hline
% 96 & ACE2  Rattus norvegicus & 98 & ACE2  Canis lupus familiaris & similar \\ \hline
% 96 & ACE2  Rattus norvegicus & 99 & ACE2  Gallus gallus & similar \\ \hline
% 96 & ACE2  Rattus norvegicus & 194 & Rattus norvegicus & belongs \\ \hline
% 97 & ACE2  Rhinolophus ferrumequinum & 98 & ACE2  Canis lupus familiaris & similar \\ \hline
% 97 & ACE2  Rhinolophus ferrumequinum & 99 & ACE2  Gallus gallus & similar \\ \hline
% 97 & ACE2  Rhinolophus ferrumequinum & 207 & Rhinolophus ferrumequinum & belongs \\ \hline
% 98 & ACE2  Canis lupus familiaris & 99 & ACE2  Gallus gallus & similar \\ \hline
% 98 & ACE2  Canis lupus familiaris & 195 & Canis lupus familiaris & belongs \\ \hline
% 99 & ACE2  Gallus gallus & 201 & Gallus gallus & belongs \\ \hline
% 100 & ORF4a  Middle East respiratory syndrome-related coronavirus & 101 & ORF4a  Human coronavirus 229E & similar \\ \hline
% 100 & ORF4a  Middle East respiratory syndrome-related coronavirus & 172 & STAT2  Homo sapiens & interacts \\ \hline
% 100 & ORF4a  Middle East respiratory syndrome-related coronavirus & 177 & STAT2  Camelus dromedarius & interacts \\ \hline
% 100 & ORF4a  Middle East respiratory syndrome-related coronavirus & 183 & STAT2  Rattus norvegicus & interacts \\ \hline
% 100 & ORF4a  Middle East respiratory syndrome-related coronavirus & 185 & STAT2  Mus musculus & interacts \\ \hline
% 100 & ORF4a  Middle East respiratory syndrome-related coronavirus & 102 & NF-$\kappa$B  Homo sapiens & interacts \\ \hline
% 100 & ORF4a  Middle East respiratory syndrome-related coronavirus & 109 & NF-$\kappa$B  Mus musculus & interacts \\ \hline
% 100 & ORF4a  Middle East respiratory syndrome-related coronavirus & 111 & NF-$\kappa$B  Rattus norvegicus & interacts \\ \hline
% 100 & ORF4a  Middle East respiratory syndrome-related coronavirus & 114 & NF-$\kappa$B  Camelus dromedarius & interacts \\ \hline
% 100 & ORF4a  Middle East respiratory syndrome-related coronavirus & 141 & PRKRA  Homo sapiens & interacts \\ \hline
% 100 & ORF4a  Middle East respiratory syndrome-related coronavirus & 145 & PRKRA  Camelus dromedarius & interacts \\ \hline
% 100 & ORF4a  Middle East respiratory syndrome-related coronavirus & 147 & PRKRA  Mus musculus & interacts \\ \hline
% 100 & ORF4a  Middle East respiratory syndrome-related coronavirus & 151 & PRKRA  Rattus norvegicus & interacts \\ \hline
% 100 & ORF4a  Middle East respiratory syndrome-related coronavirus & 129 & MDA5  Homo sapiens & interacts \\ \hline
% 100 & ORF4a  Middle East respiratory syndrome-related coronavirus & 136 & MDA5  Mus musculus & interacts \\ \hline
% 100 & ORF4a  Middle East respiratory syndrome-related coronavirus & 137 & MDA5  Rattus norvegicus & interacts \\ \hline
% 102 & NF-$\kappa$B  Homo sapiens & 103 & NF-$\kappa$B  Pan troglodytes & similar \\ \hline
% 102 & NF-$\kappa$B  Homo sapiens & 104 & NF-$\kappa$B  Macaca mulatta & similar \\ \hline
% 102 & NF-$\kappa$B  Homo sapiens & 105 & NF-$\kappa$B  Bos taurus & similar \\ \hline
% 102 & NF-$\kappa$B  Homo sapiens & 106 & NF-$\kappa$B  Rhinolophus ferrumequinum & similar \\ \hline
% 102 & NF-$\kappa$B  Homo sapiens & 107 & NF-$\kappa$B  Ovis aries & similar \\ \hline
% 102 & NF-$\kappa$B  Homo sapiens & 108 & NF-$\kappa$B  Ictidomys tridecemlineatus & similar \\ \hline
% 102 & NF-$\kappa$B  Homo sapiens & 109 & NF-$\kappa$B  Mus musculus & similar \\ \hline
% 102 & NF-$\kappa$B  Homo sapiens & 110 & NF-$\kappa$B  Mesocricetus auratus & similar \\ \hline
% 102 & NF-$\kappa$B  Homo sapiens & 111 & NF-$\kappa$B  Rattus norvegicus & similar \\ \hline
% 102 & NF-$\kappa$B  Homo sapiens & 112 & NF-$\kappa$B  Equus caballus & similar \\ \hline
% 102 & NF-$\kappa$B  Homo sapiens & 113 & NF-$\kappa$B  Gallus gallus & similar \\ \hline
% 102 & NF-$\kappa$B  Homo sapiens & 114 & NF-$\kappa$B  Camelus dromedarius & similar \\ \hline
% 102 & NF-$\kappa$B  Homo sapiens & 192 & Homo sapiens & belongs \\ \hline
% 102 & NF-$\kappa$B  Homo sapiens & 188 & PLpro  Severe acute respiratory syndrome-related coronavirus & interacts \\ \hline
% 102 & NF-$\kappa$B  Homo sapiens & 187 & PLpro  Middle East respiratory syndrome-related coronavirus & interacts \\ \hline
% 102 & NF-$\kappa$B  Homo sapiens & 211 & M protein  Severe acute respiratory syndrome-related coronavirus & interacts \\ \hline
% 103 & NF-$\kappa$B  Pan troglodytes & 104 & NF-$\kappa$B  Macaca mulatta & similar \\ \hline
% 103 & NF-$\kappa$B  Pan troglodytes & 105 & NF-$\kappa$B  Bos taurus & similar \\ \hline
% 103 & NF-$\kappa$B  Pan troglodytes & 106 & NF-$\kappa$B  Rhinolophus ferrumequinum & similar \\ \hline
% 103 & NF-$\kappa$B  Pan troglodytes & 107 & NF-$\kappa$B  Ovis aries & similar \\ \hline
% 103 & NF-$\kappa$B  Pan troglodytes & 108 & NF-$\kappa$B  Ictidomys tridecemlineatus & similar \\ \hline
% 103 & NF-$\kappa$B  Pan troglodytes & 109 & NF-$\kappa$B  Mus musculus & similar \\ \hline
% 103 & NF-$\kappa$B  Pan troglodytes & 110 & NF-$\kappa$B  Mesocricetus auratus & similar \\ \hline
% 103 & NF-$\kappa$B  Pan troglodytes & 111 & NF-$\kappa$B  Rattus norvegicus & similar \\ \hline
% 103 & NF-$\kappa$B  Pan troglodytes & 112 & NF-$\kappa$B  Equus caballus & similar \\ \hline
% 103 & NF-$\kappa$B  Pan troglodytes & 113 & NF-$\kappa$B  Gallus gallus & similar \\ \hline
% 103 & NF-$\kappa$B  Pan troglodytes & 114 & NF-$\kappa$B  Camelus dromedarius & similar \\ \hline
% 103 & NF-$\kappa$B  Pan troglodytes & 200 & Pan troglodytes & belongs \\ \hline
% 104 & NF-$\kappa$B  Macaca mulatta & 105 & NF-$\kappa$B  Bos taurus & similar \\ \hline
% 104 & NF-$\kappa$B  Macaca mulatta & 106 & NF-$\kappa$B  Rhinolophus ferrumequinum & similar \\ \hline
% 104 & NF-$\kappa$B  Macaca mulatta & 107 & NF-$\kappa$B  Ovis aries & similar \\ \hline
% 104 & NF-$\kappa$B  Macaca mulatta & 108 & NF-$\kappa$B  Ictidomys tridecemlineatus & similar \\ \hline
% 104 & NF-$\kappa$B  Macaca mulatta & 109 & NF-$\kappa$B  Mus musculus & similar \\ \hline
% 104 & NF-$\kappa$B  Macaca mulatta & 110 & NF-$\kappa$B  Mesocricetus auratus & similar \\ \hline
% 104 & NF-$\kappa$B  Macaca mulatta & 111 & NF-$\kappa$B  Rattus norvegicus & similar \\ \hline
% 104 & NF-$\kappa$B  Macaca mulatta & 112 & NF-$\kappa$B  Equus caballus & similar \\ \hline
% 104 & NF-$\kappa$B  Macaca mulatta & 113 & NF-$\kappa$B  Gallus gallus & similar \\ \hline
% 104 & NF-$\kappa$B  Macaca mulatta & 114 & NF-$\kappa$B  Camelus dromedarius & similar \\ \hline
% 104 & NF-$\kappa$B  Macaca mulatta & 204 & Macaca mulatta & belongs \\ \hline
% 105 & NF-$\kappa$B  Bos taurus & 106 & NF-$\kappa$B  Rhinolophus ferrumequinum & similar \\ \hline
% 105 & NF-$\kappa$B  Bos taurus & 107 & NF-$\kappa$B  Ovis aries & similar \\ \hline
% 105 & NF-$\kappa$B  Bos taurus & 108 & NF-$\kappa$B  Ictidomys tridecemlineatus & similar \\ \hline
% 105 & NF-$\kappa$B  Bos taurus & 109 & NF-$\kappa$B  Mus musculus & similar \\ \hline
% 105 & NF-$\kappa$B  Bos taurus & 110 & NF-$\kappa$B  Mesocricetus auratus & similar \\ \hline
% 105 & NF-$\kappa$B  Bos taurus & 111 & NF-$\kappa$B  Rattus norvegicus & similar \\ \hline
% 105 & NF-$\kappa$B  Bos taurus & 112 & NF-$\kappa$B  Equus caballus & similar \\ \hline
% 105 & NF-$\kappa$B  Bos taurus & 113 & NF-$\kappa$B  Gallus gallus & similar \\ \hline
% 105 & NF-$\kappa$B  Bos taurus & 114 & NF-$\kappa$B  Camelus dromedarius & similar \\ \hline
% 105 & NF-$\kappa$B  Bos taurus & 199 & Bos taurus & belongs \\ \hline
% 106 & NF-$\kappa$B  Rhinolophus ferrumequinum & 107 & NF-$\kappa$B  Ovis aries & similar \\ \hline
% 106 & NF-$\kappa$B  Rhinolophus ferrumequinum & 108 & NF-$\kappa$B  Ictidomys tridecemlineatus & similar \\ \hline
% 106 & NF-$\kappa$B  Rhinolophus ferrumequinum & 109 & NF-$\kappa$B  Mus musculus & similar \\ \hline
% 106 & NF-$\kappa$B  Rhinolophus ferrumequinum & 110 & NF-$\kappa$B  Mesocricetus auratus & similar \\ \hline
% 106 & NF-$\kappa$B  Rhinolophus ferrumequinum & 111 & NF-$\kappa$B  Rattus norvegicus & similar \\ \hline
% 106 & NF-$\kappa$B  Rhinolophus ferrumequinum & 112 & NF-$\kappa$B  Equus caballus & similar \\ \hline
% 106 & NF-$\kappa$B  Rhinolophus ferrumequinum & 113 & NF-$\kappa$B  Gallus gallus & similar \\ \hline
% 106 & NF-$\kappa$B  Rhinolophus ferrumequinum & 114 & NF-$\kappa$B  Camelus dromedarius & similar \\ \hline
% 106 & NF-$\kappa$B  Rhinolophus ferrumequinum & 207 & Rhinolophus ferrumequinum & belongs \\ \hline
% 106 & NF-$\kappa$B  Rhinolophus ferrumequinum & 188 & PLpro  Severe acute respiratory syndrome-related coronavirus & interacts \\ \hline
% 106 & NF-$\kappa$B  Rhinolophus ferrumequinum & 211 & M protein  Severe acute respiratory syndrome-related coronavirus & interacts \\ \hline
% 107 & NF-$\kappa$B  Ovis aries & 108 & NF-$\kappa$B  Ictidomys tridecemlineatus & similar \\ \hline
% 107 & NF-$\kappa$B  Ovis aries & 109 & NF-$\kappa$B  Mus musculus & similar \\ \hline
% 107 & NF-$\kappa$B  Ovis aries & 110 & NF-$\kappa$B  Mesocricetus auratus & similar \\ \hline
% 107 & NF-$\kappa$B  Ovis aries & 111 & NF-$\kappa$B  Rattus norvegicus & similar \\ \hline
% 107 & NF-$\kappa$B  Ovis aries & 112 & NF-$\kappa$B  Equus caballus & similar \\ \hline
% 107 & NF-$\kappa$B  Ovis aries & 113 & NF-$\kappa$B  Gallus gallus & similar \\ \hline
% 107 & NF-$\kappa$B  Ovis aries & 114 & NF-$\kappa$B  Camelus dromedarius & similar \\ \hline
% 107 & NF-$\kappa$B  Ovis aries & 205 & Ovis aries & belongs \\ \hline
% 108 & NF-$\kappa$B  Ictidomys tridecemlineatus & 109 & NF-$\kappa$B  Mus musculus & similar \\ \hline
% 108 & NF-$\kappa$B  Ictidomys tridecemlineatus & 110 & NF-$\kappa$B  Mesocricetus auratus & similar \\ \hline
% 108 & NF-$\kappa$B  Ictidomys tridecemlineatus & 111 & NF-$\kappa$B  Rattus norvegicus & similar \\ \hline
% 108 & NF-$\kappa$B  Ictidomys tridecemlineatus & 112 & NF-$\kappa$B  Equus caballus & similar \\ \hline
% 108 & NF-$\kappa$B  Ictidomys tridecemlineatus & 113 & NF-$\kappa$B  Gallus gallus & similar \\ \hline
% 108 & NF-$\kappa$B  Ictidomys tridecemlineatus & 114 & NF-$\kappa$B  Camelus dromedarius & similar \\ \hline
% 108 & NF-$\kappa$B  Ictidomys tridecemlineatus & 198 & Ictidomys tridecemlineatus & belongs \\ \hline
% 109 & NF-$\kappa$B  Mus musculus & 110 & NF-$\kappa$B  Mesocricetus auratus & similar \\ \hline
% 109 & NF-$\kappa$B  Mus musculus & 111 & NF-$\kappa$B  Rattus norvegicus & similar \\ \hline
% 109 & NF-$\kappa$B  Mus musculus & 112 & NF-$\kappa$B  Equus caballus & similar \\ \hline
% 109 & NF-$\kappa$B  Mus musculus & 113 & NF-$\kappa$B  Gallus gallus & similar \\ \hline
% 109 & NF-$\kappa$B  Mus musculus & 114 & NF-$\kappa$B  Camelus dromedarius & similar \\ \hline
% 109 & NF-$\kappa$B  Mus musculus & 193 & Mus musculus & belongs \\ \hline
% 109 & NF-$\kappa$B  Mus musculus & 188 & PLpro  Severe acute respiratory syndrome-related coronavirus & interacts \\ \hline
% 109 & NF-$\kappa$B  Mus musculus & 187 & PLpro  Middle East respiratory syndrome-related coronavirus & interacts \\ \hline
% 109 & NF-$\kappa$B  Mus musculus & 211 & M protein  Severe acute respiratory syndrome-related coronavirus & interacts \\ \hline
% 110 & NF-$\kappa$B  Mesocricetus auratus & 111 & NF-$\kappa$B  Rattus norvegicus & similar \\ \hline
% 110 & NF-$\kappa$B  Mesocricetus auratus & 112 & NF-$\kappa$B  Equus caballus & similar \\ \hline
% 110 & NF-$\kappa$B  Mesocricetus auratus & 113 & NF-$\kappa$B  Gallus gallus & similar \\ \hline
% 110 & NF-$\kappa$B  Mesocricetus auratus & 114 & NF-$\kappa$B  Camelus dromedarius & similar \\ \hline
% 110 & NF-$\kappa$B  Mesocricetus auratus & 208 & Mesocricetus auratus & belongs \\ \hline
% 111 & NF-$\kappa$B  Rattus norvegicus & 112 & NF-$\kappa$B  Equus caballus & similar \\ \hline
% 111 & NF-$\kappa$B  Rattus norvegicus & 113 & NF-$\kappa$B  Gallus gallus & similar \\ \hline
% 111 & NF-$\kappa$B  Rattus norvegicus & 114 & NF-$\kappa$B  Camelus dromedarius & similar \\ \hline
% 111 & NF-$\kappa$B  Rattus norvegicus & 194 & Rattus norvegicus & belongs \\ \hline
% 111 & NF-$\kappa$B  Rattus norvegicus & 188 & PLpro  Severe acute respiratory syndrome-related coronavirus & interacts \\ \hline
% 111 & NF-$\kappa$B  Rattus norvegicus & 187 & PLpro  Middle East respiratory syndrome-related coronavirus & interacts \\ \hline
% 111 & NF-$\kappa$B  Rattus norvegicus & 211 & M protein  Severe acute respiratory syndrome-related coronavirus & interacts \\ \hline
% 112 & NF-$\kappa$B  Equus caballus & 113 & NF-$\kappa$B  Gallus gallus & similar \\ \hline
% 112 & NF-$\kappa$B  Equus caballus & 114 & NF-$\kappa$B  Camelus dromedarius & similar \\ \hline
% 112 & NF-$\kappa$B  Equus caballus & 203 & Equus caballus & belongs \\ \hline
% 113 & NF-$\kappa$B  Gallus gallus & 114 & NF-$\kappa$B  Camelus dromedarius & similar \\ \hline
% 113 & NF-$\kappa$B  Gallus gallus & 201 & Gallus gallus & belongs \\ \hline
% 114 & NF-$\kappa$B  Camelus dromedarius & 196 & Camelus dromedarius & belongs \\ \hline
% 114 & NF-$\kappa$B  Camelus dromedarius & 187 & PLpro  Middle East respiratory syndrome-related coronavirus & interacts \\ \hline
% 115 & IRF7  Homo sapiens & 116 & IRF7  Pan troglodytes & similar \\ \hline
% 115 & IRF7  Homo sapiens & 117 & IRF7  Macaca mulatta & similar \\ \hline
% 115 & IRF7  Homo sapiens & 118 & IRF7  Ictidomys tridecemlineatus & similar \\ \hline
% 115 & IRF7  Homo sapiens & 119 & IRF7  Equus caballus & similar \\ \hline
% 115 & IRF7  Homo sapiens & 120 & IRF7  Rhinolophus ferrumequinum & similar \\ \hline
% 115 & IRF7  Homo sapiens & 121 & IRF7  Felis catus & similar \\ \hline
% 115 & IRF7  Homo sapiens & 122 & IRF7  Camelus dromedarius & similar \\ \hline
% 115 & IRF7  Homo sapiens & 123 & IRF7  Rattus norvegicus & similar \\ \hline
% 115 & IRF7  Homo sapiens & 124 & IRF7  Mesocricetus auratus & similar \\ \hline
% 115 & IRF7  Homo sapiens & 125 & IRF7  Ovis aries & similar \\ \hline
% 115 & IRF7  Homo sapiens & 126 & IRF7  Mus musculus & similar \\ \hline
% 115 & IRF7  Homo sapiens & 127 & IRF7  Bos taurus & similar \\ \hline
% 115 & IRF7  Homo sapiens & 128 & IRF7  Gallus gallus & similar \\ \hline
% 115 & IRF7  Homo sapiens & 192 & Homo sapiens & belongs \\ \hline
% 116 & IRF7  Pan troglodytes & 117 & IRF7  Macaca mulatta & similar \\ \hline
% 116 & IRF7  Pan troglodytes & 118 & IRF7  Ictidomys tridecemlineatus & similar \\ \hline
% 116 & IRF7  Pan troglodytes & 119 & IRF7  Equus caballus & similar \\ \hline
% 116 & IRF7  Pan troglodytes & 120 & IRF7  Rhinolophus ferrumequinum & similar \\ \hline
% 116 & IRF7  Pan troglodytes & 121 & IRF7  Felis catus & similar \\ \hline
% 116 & IRF7  Pan troglodytes & 122 & IRF7  Camelus dromedarius & similar \\ \hline
% 116 & IRF7  Pan troglodytes & 123 & IRF7  Rattus norvegicus & similar \\ \hline
% 116 & IRF7  Pan troglodytes & 124 & IRF7  Mesocricetus auratus & similar \\ \hline
% 116 & IRF7  Pan troglodytes & 125 & IRF7  Ovis aries & similar \\ \hline
% 116 & IRF7  Pan troglodytes & 126 & IRF7  Mus musculus & similar \\ \hline
% 116 & IRF7  Pan troglodytes & 127 & IRF7  Bos taurus & similar \\ \hline
% 116 & IRF7  Pan troglodytes & 128 & IRF7  Gallus gallus & similar \\ \hline
% 116 & IRF7  Pan troglodytes & 200 & Pan troglodytes & belongs \\ \hline
% 117 & IRF7  Macaca mulatta & 118 & IRF7  Ictidomys tridecemlineatus & similar \\ \hline
% 117 & IRF7  Macaca mulatta & 119 & IRF7  Equus caballus & similar \\ \hline
% 117 & IRF7  Macaca mulatta & 120 & IRF7  Rhinolophus ferrumequinum & similar \\ \hline
% 117 & IRF7  Macaca mulatta & 121 & IRF7  Felis catus & similar \\ \hline
% 117 & IRF7  Macaca mulatta & 122 & IRF7  Camelus dromedarius & similar \\ \hline
% 117 & IRF7  Macaca mulatta & 123 & IRF7  Rattus norvegicus & similar \\ \hline
% 117 & IRF7  Macaca mulatta & 124 & IRF7  Mesocricetus auratus & similar \\ \hline
% 117 & IRF7  Macaca mulatta & 125 & IRF7  Ovis aries & similar \\ \hline
% 117 & IRF7  Macaca mulatta & 126 & IRF7  Mus musculus & similar \\ \hline
% 117 & IRF7  Macaca mulatta & 127 & IRF7  Bos taurus & similar \\ \hline
% 117 & IRF7  Macaca mulatta & 128 & IRF7  Gallus gallus & similar \\ \hline
% 117 & IRF7  Macaca mulatta & 204 & Macaca mulatta & belongs \\ \hline
% 118 & IRF7  Ictidomys tridecemlineatus & 119 & IRF7  Equus caballus & similar \\ \hline
% 118 & IRF7  Ictidomys tridecemlineatus & 120 & IRF7  Rhinolophus ferrumequinum & similar \\ \hline
% 118 & IRF7  Ictidomys tridecemlineatus & 121 & IRF7  Felis catus & similar \\ \hline
% 118 & IRF7  Ictidomys tridecemlineatus & 122 & IRF7  Camelus dromedarius & similar \\ \hline
% 118 & IRF7  Ictidomys tridecemlineatus & 123 & IRF7  Rattus norvegicus & similar \\ \hline
% 118 & IRF7  Ictidomys tridecemlineatus & 124 & IRF7  Mesocricetus auratus & similar \\ \hline
% 118 & IRF7  Ictidomys tridecemlineatus & 125 & IRF7  Ovis aries & similar \\ \hline
% 118 & IRF7  Ictidomys tridecemlineatus & 126 & IRF7  Mus musculus & similar \\ \hline
% 118 & IRF7  Ictidomys tridecemlineatus & 127 & IRF7  Bos taurus & similar \\ \hline
% 118 & IRF7  Ictidomys tridecemlineatus & 128 & IRF7  Gallus gallus & similar \\ \hline
% 118 & IRF7  Ictidomys tridecemlineatus & 198 & Ictidomys tridecemlineatus & belongs \\ \hline
% 119 & IRF7  Equus caballus & 120 & IRF7  Rhinolophus ferrumequinum & similar \\ \hline
% 119 & IRF7  Equus caballus & 121 & IRF7  Felis catus & similar \\ \hline
% 119 & IRF7  Equus caballus & 122 & IRF7  Camelus dromedarius & similar \\ \hline
% \end{longtable}
% \label{tab:edge table}

% \newpage


% \section{Full Predicted Infections between Coronaviruses and mammalian hosts}

% \begin{longtable}[c]{|c|c|c|c|c|}
% \hline
% \textbf{Source Name} & \textbf{Target Name} & \textbf{Certainty} & \textbf{Confidence} & \textbf{Likelihood} \\ \hline
% \endfirsthead
% %
% \multicolumn{5}{c}%
% {{\bfseries Table \thetable\ continued from previous page}} \\
% \hline
% \textbf{Source Name} & \textbf{Target Name} & \textbf{Certainty} & \textbf{Confidence} & \textbf{Likelihood} \\ \hline
% \endhead
% %
% virus Severe acute respiratory syndrome coronavirus 2 & host Mus musculus & 100.00\% & strong & likely \\ \hline
% virus Severe acute respiratory syndrome-related coronavirus & host Mesocricetus auratus & 99.98\% & strong & likely \\ \hline
% virus Human coronavirus NL63 & host Rattus norvegicus & 99.96\% & strong & likely \\ \hline
% virus Severe acute respiratory syndrome coronavirus 2 & host Rattus norvegicus & 99.95\% & strong & likely \\ \hline
% virus Severe acute respiratory syndrome-related coronavirus & host Macaca mulatta & 99.94\% & strong & likely \\ \hline
% virus Severe acute respiratory syndrome-related coronavirus & host Canis lupus familiaris & 99.94\% & strong & likely \\ \hline
% virus Human coronavirus HKU1 & host Felis catus & 99.90\% & strong & likely \\ \hline
% virus Human coronavirus NL63 & host Mus musculus & 99.87\% & strong & likely \\ \hline
% virus Human coronavirus HKU1 & host Mesocricetus auratus & 99.86\% & strong & likely \\ \hline
% virus Human coronavirus NL63 & host Felis catus & 99.58\% & strong & likely \\ \hline
% virus Human coronavirus HKU1 & host Canis lupus familiaris & 99.51\% & strong & likely \\ \hline
% virus Human coronavirus HKU1 & host Macaca mulatta & 99.44\% & strong & likely \\ \hline
% virus Human coronavirus 229E & host Mesocricetus auratus & 99.34\% & weak & likely \\ \hline
% virus Human coronavirus NL63 & host Canis lupus familiaris & 92.32\% & strong & likely \\ \hline
% virus Human coronavirus NL63 & host Mesocricetus auratus & 78.57\% & strong & likely \\ \hline
% virus Severe acute respiratory syndrome coronavirus 2 & host Ictidomys tridecemlineatus & 74.07\% & weak & likely \\ \hline
% virus Human coronavirus NL63 & host Macaca mulatta & 66.45\% & strong & likely \\ \hline
% virus Severe acute respiratory syndrome coronavirus 2 & host Bos taurus & 61.00\% & weak & likely \\ \hline
% virus Severe acute respiratory syndrome coronavirus 2 & host Ovis aries & 54.31\% & weak & likely \\ \hline
% virus Human coronavirus OC43 & host Rhinolophus ferrumequinum & 99.99\% & strong & unlikely \\ \hline
% virus Human coronavirus OC43 & host Ovis aries & 99.99\% & weak & unlikely \\ \hline
% virus Middle East respiratory syndrome-related coronavirus & host Rhinolophus ferrumequinum & 99.93\% & strong & unlikely \\ \hline
% virus Human coronavirus 229E & host Rattus norvegicus & 99.78\% & strong & unlikely \\ \hline
% virus Human coronavirus 229E & host Macaca mulatta & 99.70\% & weak & unlikely \\ \hline
% virus Human coronavirus NL63 & host Camelus dromedarius & 99.61\% & weak & unlikely \\ \hline
% virus Human coronavirus HKU1 & host Bos taurus & 99.50\% & strong & unlikely \\ \hline
% virus Human coronavirus HKU1 & host Ovis aries & 99.06\% & weak & unlikely \\ \hline
% virus Severe acute respiratory syndrome-related coronavirus & host Camelus dromedarius & 98.94\% & strong & unlikely \\ \hline
% virus Human coronavirus HKU1 & host Camelus dromedarius & 98.80\% & strong & unlikely \\ \hline
% virus Human coronavirus 229E & host Mus musculus & 98.12\% & strong & unlikely \\ \hline
% virus Human coronavirus OC43 & host Felis catus & 97.23\% & strong & unlikely \\ \hline
% virus Human coronavirus 229E & host Ictidomys tridecemlineatus & 96.83\% & weak & unlikely \\ \hline
% virus Human coronavirus OC43 & host Equus caballus & 96.76\% & weak & unlikely \\ \hline
% virus Human coronavirus OC43 & host Mesocricetus auratus & 96.05\% & strong & unlikely \\ \hline
% virus Middle East respiratory syndrome-related coronavirus & host Sus scrofa domesticus & 94.89\% & weak & unlikely \\ \hline
% virus Human coronavirus 229E & host Canis lupus familiaris & 94.03\% & weak & unlikely \\ \hline
% virus Severe acute respiratory syndrome-related coronavirus & host Bos taurus & 89.91\% & strong & unlikely \\ \hline
% virus Middle East respiratory syndrome-related coronavirus & host Ictidomys tridecemlineatus & 88.31\% & weak & unlikely \\ \hline
% virus Human coronavirus OC43 & host Canis lupus familiaris & 87.41\% & strong & unlikely \\ \hline
% virus Middle East respiratory syndrome-related coronavirus & host Felis catus & 85.28\% & strong & unlikely \\ \hline
% virus Human coronavirus OC43 & host Macaca mulatta & 84.02\% & strong & unlikely \\ \hline
% virus Human coronavirus OC43 & host Sus scrofa domesticus & 82.90\% & weak & unlikely \\ \hline
% virus Severe acute respiratory syndrome-related coronavirus & host Gallus gallus & 81.28\% & weak & unlikely \\ \hline
% virus Human coronavirus HKU1 & host Ictidomys tridecemlineatus & 80.05\% & weak & unlikely \\ \hline
% virus Human coronavirus OC43 & host Camelus dromedarius & 77.19\% & strong & unlikely \\ \hline
% virus Human coronavirus 229E & host Ovis aries & 75.78\% & weak & unlikely \\ \hline
% virus Severe acute respiratory syndrome-related coronavirus & host Sus scrofa domesticus & 75.20\% & weak & unlikely \\ \hline
% virus Human coronavirus OC43 & host Ictidomys tridecemlineatus & 74.53\% & weak & unlikely \\ \hline
% virus Human coronavirus 229E & host Felis catus & 74.30\% & weak & unlikely \\ \hline
% virus Human coronavirus 229E & host Bos taurus & 71.14\% & weak & unlikely \\ \hline
% virus Severe acute respiratory syndrome coronavirus 2 & host Sus scrofa domesticus & 71.09\% & weak & unlikely \\ \hline
% virus Severe acute respiratory syndrome-related coronavirus & host Ovis aries & 68.68\% & weak & unlikely \\ \hline
% virus Severe acute respiratory syndrome coronavirus 2 & host Camelus dromedarius & 68.24\% & weak & unlikely \\ \hline
% virus Severe acute respiratory syndrome coronavirus 2 & host Gallus gallus & 66.89\% & weak & unlikely \\ \hline
% virus Middle East respiratory syndrome-related coronavirus & host Mesocricetus auratus & 65.98\% & strong & unlikely \\ \hline
% virus Human coronavirus HKU1 & host Sus scrofa domesticus & 63.44\% & weak & unlikely \\ \hline
% virus Severe acute respiratory syndrome coronavirus 2 & host Oryctolagus cuniculus & 59.07\% & weak & unlikely \\ \hline
% virus Human coronavirus 229E & host Camelus dromedarius & 59.04\% & weak & unlikely \\ \hline
% virus Middle East respiratory syndrome-related coronavirus & host Gallus gallus & 58.63\% & weak & unlikely \\ \hline
% virus Human coronavirus NL63 & host Bos taurus & 57.98\% & weak & unlikely \\ \hline
% virus Middle East respiratory syndrome-related coronavirus & host Macaca mulatta & 57.54\% & strong & unlikely \\ \hline
% virus Middle East respiratory syndrome-related coronavirus & host Bos taurus & 54.50\% & weak & unlikely \\ \hline
% virus Middle East respiratory syndrome-related coronavirus & host Equus caballus & 54.15\% & weak & unlikely \\ \hline
% virus Severe acute respiratory syndrome-related coronavirus & host Ictidomys tridecemlineatus & 50.98\% & weak & unlikely \\ \hline
% virus Middle East respiratory syndrome-related coronavirus & host Ovis aries & 50.69\% & weak & unlikely \\ \hline
% \end{longtable}
% \label{tab: infection prediction}
% \newpage


% \section{Full Predicted Protein-Protein Interactions between Coronaviruses and mammalian hosts}

% \begin{longtable}[c]{|c|c|c|c|c|}
% \hline
% \textbf{Source Name} & \textbf{Target Name} & \textbf{Certainty} & \textbf{Confidence} & \textbf{Likelihood} \\ \hline
% \endfirsthead
% %
% \multicolumn{5}{c}%
% {{\bfseries Table \thetable\ continued from previous page}} \\
% \hline
% \textbf{Source Name} & \textbf{Target Name} & \textbf{Certainty} & \textbf{Confidence} & \textbf{Likelihood} \\ \hline
% \endhead
% %
% STAT1 Homo sapiens & PLpro Severe acute respiratory syndrome-related coronavirus & 99.99\% & strong & likely \\ \hline
% STAT2 Homo sapiens & PLpro Severe acute respiratory syndrome-related coronavirus & 99.99\% & strong & likely \\ \hline
% STAT1 Homo sapiens & M protein Severe acute respiratory syndrome-related coronavirus & 99.99\% & strong & likely \\ \hline
% STAT2 Homo sapiens & M protein Severe acute respiratory syndrome-related coronavirus & 99.99\% & strong & likely \\ \hline
% IRF9 Homo sapiens & nsp1 Severe acute respiratory syndrome-related coronavirus & 99.97\% & strong & likely \\ \hline
% nsp15 Severe acute respiratory syndrome-related coronavirus & RIG-I Rhinolophus ferrumequinum & 99.97\% & strong & likely \\ \hline
% IRF9 Homo sapiens & PLpro Severe acute respiratory syndrome-related coronavirus & 99.97\% & strong & likely \\ \hline
% RIG-I Rhinolophus ferrumequinum & ORF6 Severe acute respiratory syndrome-related coronavirus & 99.97\% & strong & likely \\ \hline
% IRF9 Rattus norvegicus & PLpro Severe acute respiratory syndrome-related coronavirus & 99.97\% & strong & likely \\ \hline
% STAT1 Rattus norvegicus & PLpro Severe acute respiratory syndrome-related coronavirus & 99.97\% & strong & likely \\ \hline
% ORF3b Severe acute respiratory syndrome-related coronavirus & MAVS Macaca mulatta & 99.95\% & strong & likely \\ \hline
% IRF9 Homo sapiens & M protein Severe acute respiratory syndrome-related coronavirus & 99.95\% & strong & likely \\ \hline
% IRF3 Rattus norvegicus & ORF6 Severe acute respiratory syndrome coronavirus 2 & 99.94\% & strong & likely \\ \hline
% nsp15 Severe acute respiratory syndrome-related coronavirus & MDA5 Rattus norvegicus & 99.94\% & strong & likely \\ \hline
% IRF3 Macaca mulatta & ORF6 Severe acute respiratory syndrome-related coronavirus & 99.92\% & strong & likely \\ \hline
% IRF9 Mus musculus & PLpro Severe acute respiratory syndrome-related coronavirus & 99.90\% & strong & likely \\ \hline
% nsp15 Severe acute respiratory syndrome coronavirus 2 & RIG-I Mus musculus & 99.90\% & strong & likely \\ \hline
% nsp1 Severe acute respiratory syndrome-related coronavirus & STAT2 Macaca mulatta & 99.90\% & strong & likely \\ \hline
% ORF4a Middle East respiratory syndrome-related coronavirus & IRF7 Rattus norvegicus & 99.90\% & strong & likely \\ \hline
% nsp15 Severe acute respiratory syndrome-related coronavirus & RIG-I Homo sapiens & 99.88\% & strong & likely \\ \hline
% IRF7 Homo sapiens & ORF6 Severe acute respiratory syndrome-related coronavirus & 99.87\% & strong & likely \\ \hline
% nsp15 Severe acute respiratory syndrome-related coronavirus & STAT2 Homo sapiens & 99.87\% & strong & likely \\ \hline
% STAT2 Rattus norvegicus & PLpro Severe acute respiratory syndrome-related coronavirus & 99.87\% & strong & likely \\ \hline
% STAT1 Mus musculus & PLpro Severe acute respiratory syndrome-related coronavirus & 99.87\% & strong & likely \\ \hline
% IRF3 Macaca mulatta & M protein Severe acute respiratory syndrome-related coronavirus & 99.86\% & strong & likely \\ \hline
% NF-kB Ovis aries & PLpro Severe acute respiratory syndrome-related coronavirus & 99.85\% & strong & likely \\ \hline
% ORF6 Severe acute respiratory syndrome-related coronavirus & TBK1 Homo sapiens & 99.85\% & strong & likely \\ \hline
% M protein Severe acute respiratory syndrome-related coronavirus & TBK1 Rattus norvegicus & 99.83\% & strong & likely \\ \hline
% STAT2 Mus musculus & PLpro Severe acute respiratory syndrome-related coronavirus & 99.82\% & strong & likely \\ \hline
% IRF7 Homo sapiens & ORF3b Severe acute respiratory syndrome-related coronavirus & 99.78\% & strong & likely \\ \hline
% nsp15 Severe acute respiratory syndrome-related coronavirus & MAVS Macaca mulatta & 99.77\% & strong & likely \\ \hline
% ORF3b Severe acute respiratory syndrome-related coronavirus & TBK1 Homo sapiens & 99.76\% & strong & likely \\ \hline
% nsp15 Severe acute respiratory syndrome-related coronavirus & IRF9 Rhinolophus ferrumequinum & 99.74\% & strong & likely \\ \hline
% IRF9 Macaca mulatta & ORF3b Severe acute respiratory syndrome-related coronavirus & 99.73\% & strong & likely \\ \hline
% ORF6 Severe acute respiratory syndrome-related coronavirus & MAVS Macaca mulatta & 99.72\% & strong & likely \\ \hline
% NF-kB Camelus dromedarius & M protein Middle East respiratory syndrome-related coronavirus & 99.72\% & strong & likely \\ \hline
% ORF6 Severe acute respiratory syndrome-related coronavirus & TBK1 Rattus norvegicus & 99.71\% & strong & likely \\ \hline
% nsp15 Severe acute respiratory syndrome-related coronavirus & STAT1 Rhinolophus ferrumequinum & 99.68\% & strong & likely \\ \hline
% STAT1 Rattus norvegicus & M protein Severe acute respiratory syndrome-related coronavirus & 99.67\% & strong & likely \\ \hline
% IRF9 Rattus norvegicus & PLpro Middle East respiratory syndrome-related coronavirus & 99.67\% & strong & likely \\ \hline
% nsp1 Severe acute respiratory syndrome coronavirus 2 & STAT2 Felis catus & 99.64\% & strong & likely \\ \hline
% nsp15 Severe acute respiratory syndrome-related coronavirus & IRF3 Mesocricetus auratus & 99.63\% & strong & likely \\ \hline
% STAT2 Rattus norvegicus & M protein Severe acute respiratory syndrome-related coronavirus & 99.62\% & strong & likely \\ \hline
% nsp15 Severe acute respiratory syndrome-related coronavirus & STAT2 Rhinolophus ferrumequinum & 99.61\% & strong & likely \\ \hline
% nsp15 Severe acute respiratory syndrome coronavirus 2 & MAVS Canis lupus familiaris & 99.61\% & strong & likely \\ \hline
% Spike Human coronavirus NL63 & ACE2 Canis lupus familiaris & 99.59\% & strong & likely \\ \hline
% IRF3 Mus musculus & ORF6 Severe acute respiratory syndrome coronavirus 2 & 99.56\% & strong & likely \\ \hline
% M protein Severe acute respiratory syndrome-related coronavirus & TBK1 Mus musculus & 99.54\% & strong & likely \\ \hline
% nsp1 Severe acute respiratory syndrome-related coronavirus & IRF3 Rhinolophus ferrumequinum & 99.53\% & strong & likely \\ \hline
% NF-kB Bos taurus & M protein Severe acute respiratory syndrome-related coronavirus & 99.53\% & strong & likely \\ \hline
% IRF3 Canis lupus familiaris & N protein Severe acute respiratory syndrome-related coronavirus & 99.53\% & strong & likely \\ \hline
% STAT1 Mus musculus & M protein Severe acute respiratory syndrome-related coronavirus & 99.52\% & strong & likely \\ \hline
% RIG-I Macaca mulatta & ORF3b Severe acute respiratory syndrome-related coronavirus & 99.50\% & strong & likely \\ \hline
% IRF3 Macaca mulatta & N protein Severe acute respiratory syndrome-related coronavirus & 99.49\% & strong & likely \\ \hline
% STAT1 Macaca mulatta & nsp1 Severe acute respiratory syndrome-related coronavirus & 99.49\% & strong & likely \\ \hline
% nsp15 Severe acute respiratory syndrome-related coronavirus & IRF3 Canis lupus familiaris & 99.49\% & strong & likely \\ \hline
% ORF6 Severe acute respiratory syndrome coronavirus 2 & STAT2 Rhinolophus ferrumequinum & 99.44\% & strong & likely \\ \hline
% STAT1 Mesocricetus auratus & ORF6 Severe acute respiratory syndrome-related coronavirus & 99.42\% & strong & likely \\ \hline
% ORF4a Middle East respiratory syndrome-related coronavirus & TBK1 Homo sapiens & 99.42\% & strong & likely \\ \hline
% M protein Severe acute respiratory syndrome-related coronavirus & TBK1 Homo sapiens & 99.42\% & strong & likely \\ \hline
% ORF4a Middle East respiratory syndrome-related coronavirus & STAT2 Felis catus & 99.39\% & strong & likely \\ \hline
% MDA5 Rattus norvegicus & PLpro Severe acute respiratory syndrome-related coronavirus & 99.39\% & strong & likely \\ \hline
% IRF9 Mus musculus & PLpro Middle East respiratory syndrome-related coronavirus & 99.39\% & strong & likely \\ \hline
% ORF4b Middle East respiratory syndrome-related coronavirus & PRKRA Rattus norvegicus & 99.38\% & strong & likely \\ \hline
% IRF9 Rattus norvegicus & nsp1 Severe acute respiratory syndrome-related coronavirus & 99.37\% & strong & likely \\ \hline
% IRF3 Macaca mulatta & ORF3b Severe acute respiratory syndrome-related coronavirus & 99.35\% & strong & likely \\ \hline
% ORF6 Severe acute respiratory syndrome coronavirus 2 & MAVS Canis lupus familiaris & 99.34\% & strong & likely \\ \hline
% IRF9 Rattus norvegicus & M protein Severe acute respiratory syndrome-related coronavirus & 99.34\% & strong & likely \\ \hline
% STAT1 Homo sapiens & N protein Severe acute respiratory syndrome-related coronavirus & 99.34\% & strong & likely \\ \hline
% RIG-I Macaca mulatta & ORF6 Severe acute respiratory syndrome-related coronavirus & 99.33\% & strong & likely \\ \hline
% M protein Severe acute respiratory syndrome-related coronavirus & TBK1 Macaca mulatta & 99.33\% & strong & likely \\ \hline
% STAT1 Rhinolophus ferrumequinum & N protein Severe acute respiratory syndrome-related coronavirus & 99.31\% & strong & likely \\ \hline
% STAT2 Mus musculus & M protein Severe acute respiratory syndrome-related coronavirus & 99.28\% & strong & likely \\ \hline
% ORF4a Middle East respiratory syndrome-related coronavirus & MAVS Camelus dromedarius & 99.27\% & strong & likely \\ \hline
% STAT1 Macaca mulatta & ORF6 Severe acute respiratory syndrome-related coronavirus & 99.17\% & strong & likely \\ \hline
% IRF9 Mus musculus & M protein Severe acute respiratory syndrome-related coronavirus & 99.16\% & strong & likely \\ \hline
% IRF9 Rhinolophus ferrumequinum & nsp1 Severe acute respiratory syndrome-related coronavirus & 99.14\% & strong & likely \\ \hline
% N protein Severe acute respiratory syndrome-related coronavirus & STAT2 Homo sapiens & 99.11\% & strong & likely \\ \hline
% IRF3 Camelus dromedarius & PLpro Severe acute respiratory syndrome-related coronavirus & 99.09\% & strong & likely \\ \hline
% PLpro Severe acute respiratory syndrome-related coronavirus & TBK1 Macaca mulatta & 99.09\% & strong & likely \\ \hline
% ORF6 Severe acute respiratory syndrome-related coronavirus & STAT2 Mesocricetus auratus & 99.09\% & strong & likely \\ \hline
% ORF6 Severe acute respiratory syndrome coronavirus 2 & MAVS Homo sapiens & 99.08\% & strong & likely \\ \hline
% STAT1 Rhinolophus ferrumequinum & ORF4a Middle East respiratory syndrome-related coronavirus & 99.08\% & strong & likely \\ \hline
% STAT1 Felis catus & N protein Severe acute respiratory syndrome-related coronavirus & 99.08\% & strong & likely \\ \hline
% MDA5 Homo sapiens & M protein Severe acute respiratory syndrome-related coronavirus & 99.06\% & strong & likely \\ \hline
% IRF3 Macaca mulatta & PLpro Severe acute respiratory syndrome-related coronavirus & 99.06\% & strong & likely \\ \hline
% ORF3b Severe acute respiratory syndrome-related coronavirus & TBK1 Rattus norvegicus & 99.06\% & strong & likely \\ \hline
% IRF9 Felis catus & nsp1 Severe acute respiratory syndrome-related coronavirus & 99.04\% & strong & likely \\ \hline
% nsp15 Severe acute respiratory syndrome-related coronavirus & STAT1 Homo sapiens & 99.01\% & strong & likely \\ \hline
% STAT2 Mus musculus & PLpro Middle East respiratory syndrome-related coronavirus & 98.97\% & strong & likely \\ \hline
% IRF7 Mus musculus & M protein Middle East respiratory syndrome-related coronavirus & 98.93\% & strong & likely \\ \hline
% MDA5 Homo sapiens & PLpro Severe acute respiratory syndrome-related coronavirus & 98.91\% & strong & likely \\ \hline
% nsp15 Severe acute respiratory syndrome-related coronavirus & STAT2 Rattus norvegicus & 98.89\% & strong & likely \\ \hline
% ORF6 Severe acute respiratory syndrome-related coronavirus & STAT2 Camelus dromedarius & 98.89\% & strong & likely \\ \hline
% M protein Severe acute respiratory syndrome-related coronavirus & MAVS Rattus norvegicus & 98.88\% & strong & likely \\ \hline
% RIG-I Gallus gallus & ORF6 Severe acute respiratory syndrome coronavirus 2 & 98.88\% & strong & likely \\ \hline
% ORF4b Middle East respiratory syndrome-related coronavirus & STAT2 Rhinolophus ferrumequinum & 98.87\% & strong & likely \\ \hline
% MDA5 Macaca mulatta & ORF3b Severe acute respiratory syndrome-related coronavirus & 98.86\% & strong & likely \\ \hline
% STAT2 Homo sapiens & PLpro Middle East respiratory syndrome-related coronavirus & 98.83\% & strong & likely \\ \hline
% STAT1 Camelus dromedarius & ORF6 Severe acute respiratory syndrome-related coronavirus & 98.82\% & strong & likely \\ \hline
% Spike Severe acute respiratory syndrome-related coronavirus & ACE2 Mus musculus & 98.80\% & strong & likely \\ \hline
% nsp15 Severe acute respiratory syndrome-related coronavirus & STAT1 Mus musculus & 98.79\% & strong & likely \\ \hline
% ORF6 Severe acute respiratory syndrome-related coronavirus & STAT2 Macaca mulatta & 98.78\% & strong & likely \\ \hline
% Spike Severe acute respiratory syndrome coronavirus 2 & ACE2 Mus musculus & 98.77\% & strong & likely \\ \hline
% STAT1 Homo sapiens & PLpro Middle East respiratory syndrome-related coronavirus & 98.76\% & strong & likely \\ \hline
% Spike Severe acute respiratory syndrome-related coronavirus & ACE2 Rattus norvegicus & 98.76\% & strong & likely \\ \hline
% nsp15 Severe acute respiratory syndrome-related coronavirus & IRF9 Homo sapiens & 98.73\% & strong & likely \\ \hline
% nsp15 Severe acute respiratory syndrome-related coronavirus & IRF3 Macaca mulatta & 98.67\% & strong & likely \\ \hline
% IRF9 Felis catus & N protein Severe acute respiratory syndrome-related coronavirus & 98.62\% & strong & likely \\ \hline
% ORF4b Middle East respiratory syndrome-related coronavirus & PRKRA Homo sapiens & 98.59\% & strong & likely \\ \hline
% ORF4b Middle East respiratory syndrome-related coronavirus & PRKRA Camelus dromedarius & 98.54\% & strong & likely \\ \hline
% STAT1 Camelus dromedarius & PLpro Middle East respiratory syndrome-related coronavirus & 98.51\% & strong & likely \\ \hline
% Spike Severe acute respiratory syndrome-related coronavirus & ACE2 Canis lupus familiaris & 98.49\% & strong & likely \\ \hline
% Spike Severe acute respiratory syndrome-related coronavirus & ACE2 Mesocricetus auratus & 98.49\% & strong & likely \\ \hline
% nsp15 Severe acute respiratory syndrome-related coronavirus & STAT2 Felis catus & 98.48\% & strong & likely \\ \hline
% ORF4a Middle East respiratory syndrome-related coronavirus & TBK1 Rattus norvegicus & 98.44\% & strong & likely \\ \hline
% IRF3 Ovis aries & ORF6 Severe acute respiratory syndrome coronavirus 2 & 98.44\% & strong & likely \\ \hline
% ORF6 Severe acute respiratory syndrome-related coronavirus & MAVS Rattus norvegicus & 98.42\% & strong & likely \\ \hline
% nsp15 Severe acute respiratory syndrome-related coronavirus & IRF9 Felis catus & 98.41\% & strong & likely \\ \hline
% MDA5 Rattus norvegicus & ORF6 Severe acute respiratory syndrome-related coronavirus & 98.36\% & strong & likely \\ \hline
% STAT1 Canis lupus familiaris & ORF6 Severe acute respiratory syndrome-related coronavirus & 98.36\% & strong & likely \\ \hline
% STAT2 Camelus dromedarius & PLpro Middle East respiratory syndrome-related coronavirus & 98.35\% & strong & likely \\ \hline
% NF-kB Homo sapiens & M protein Middle East respiratory syndrome-related coronavirus & 98.31\% & strong & likely \\ \hline
% ORF6 Severe acute respiratory syndrome coronavirus 2 & MAVS Rhinolophus ferrumequinum & 98.30\% & strong & likely \\ \hline
% IRF9 Homo sapiens & PLpro Middle East respiratory syndrome-related coronavirus & 98.27\% & strong & likely \\ \hline
% PRKRA Camelus dromedarius & M protein Middle East respiratory syndrome-related coronavirus & 98.22\% & strong & likely \\ \hline
% ORF4b Middle East respiratory syndrome-related coronavirus & STAT2 Felis catus & 98.19\% & strong & likely \\ \hline
% nsp1 Severe acute respiratory syndrome-related coronavirus & IRF3 Homo sapiens & 98.16\% & strong & likely \\ \hline
% nsp15 Severe acute respiratory syndrome coronavirus 2 & IRF3 Rattus norvegicus & 98.15\% & strong & likely \\ \hline
% N protein Severe acute respiratory syndrome-related coronavirus & MAVS Macaca mulatta & 98.10\% & strong & likely \\ \hline
% nsp15 Severe acute respiratory syndrome-related coronavirus & STAT1 Rattus norvegicus & 98.10\% & strong & likely \\ \hline
% N protein Severe acute respiratory syndrome-related coronavirus & STAT2 Felis catus & 98.03\% & strong & likely \\ \hline
% ORF4a Middle East respiratory syndrome-related coronavirus & STAT2 Rhinolophus ferrumequinum & 98.03\% & strong & likely \\ \hline
% ORF4a Middle East respiratory syndrome-related coronavirus & MAVS Mus musculus & 98.02\% & strong & likely \\ \hline
% Spike Human coronavirus 229E & ACE2 Rhinolophus ferrumequinum & 97.93\% & strong & likely \\ \hline
% NF-kB Rattus norvegicus & ORF3b Severe acute respiratory syndrome-related coronavirus & 97.90\% & strong & likely \\ \hline
% MDA5 Rattus norvegicus & PLpro Middle East respiratory syndrome-related coronavirus & 97.86\% & strong & likely \\ \hline
% RIG-I Mesocricetus auratus & N protein Severe acute respiratory syndrome-related coronavirus & 97.77\% & strong & likely \\ \hline
% IRF3 Rattus norvegicus & N protein Middle East respiratory syndrome-related coronavirus & 97.77\% & strong & likely \\ \hline
% MDA5 Mus musculus & ORF6 Severe acute respiratory syndrome-related coronavirus & 97.71\% & strong & likely \\ \hline
% nsp15 Severe acute respiratory syndrome-related coronavirus & RIG-I Mesocricetus auratus & 97.71\% & strong & likely \\ \hline
% nsp15 Severe acute respiratory syndrome coronavirus 2 & MDA5 Felis catus & 97.71\% & strong & likely \\ \hline
% N protein Severe acute respiratory syndrome-related coronavirus & MDA5 Macaca mulatta & 97.69\% & strong & likely \\ \hline
% ORF4b Middle East respiratory syndrome-related coronavirus & IRF3 Rhinolophus ferrumequinum & 97.69\% & strong & likely \\ \hline
% ORF4a Middle East respiratory syndrome-related coronavirus & TBK1 Rhinolophus ferrumequinum & 97.62\% & strong & likely \\ \hline
% nsp15 Severe acute respiratory syndrome-related coronavirus & RIG-I Mus musculus & 97.58\% & strong & likely \\ \hline
% IRF7 Homo sapiens & M protein Middle East respiratory syndrome-related coronavirus & 97.47\% & strong & likely \\ \hline
% nsp15 Severe acute respiratory syndrome coronavirus 2 & MAVS Rattus norvegicus & 97.46\% & strong & likely \\ \hline
% STAT1 Mus musculus & PLpro Middle East respiratory syndrome-related coronavirus & 97.46\% & strong & likely \\ \hline
% Spike Human coronavirus NL63 & ACE2 Felis catus & 97.45\% & strong & likely \\ \hline
% ACE2 Rhinolophus ferrumequinum & ORF6 Severe acute respiratory syndrome coronavirus 2 & 97.38\% & strong & likely \\ \hline
% PLpro Severe acute respiratory syndrome-related coronavirus & TBK1 Ovis aries & 97.24\% & strong & likely \\ \hline
% IRF3 Mesocricetus auratus & M protein Severe acute respiratory syndrome-related coronavirus & 97.22\% & strong & likely \\ \hline
% ORF6 Severe acute respiratory syndrome coronavirus 2 & MAVS Felis catus & 97.21\% & strong & likely \\ \hline
% NF-kB Rattus norvegicus & M protein Middle East respiratory syndrome-related coronavirus & 97.18\% & strong & likely \\ \hline
% M protein Severe acute respiratory syndrome-related coronavirus & TBK1 Camelus dromedarius & 97.17\% & strong & likely \\ \hline
% Spike Human coronavirus HKU1 & ACE2 Rhinolophus ferrumequinum & 97.11\% & strong & likely \\ \hline
% IRF9 Camelus dromedarius & PLpro Middle East respiratory syndrome-related coronavirus & 96.99\% & strong & likely \\ \hline
% N protein Severe acute respiratory syndrome-related coronavirus & MAVS Canis lupus familiaris & 96.90\% & strong & likely \\ \hline
% MDA5 Rattus norvegicus & M protein Severe acute respiratory syndrome-related coronavirus & 96.86\% & strong & likely \\ \hline
% ORF4a Middle East respiratory syndrome-related coronavirus & IRF7 Camelus dromedarius & 96.76\% & strong & likely \\ \hline
% MDA5 Felis catus & ORF6 Severe acute respiratory syndrome coronavirus 2 & 96.76\% & strong & likely \\ \hline
% STAT1 Macaca mulatta & M protein Severe acute respiratory syndrome-related coronavirus & 96.74\% & strong & likely \\ \hline
% NF-kB Homo sapiens & ORF6 Severe acute respiratory syndrome-related coronavirus & 96.74\% & strong & likely \\ \hline
% ACE2 Canis lupus familiaris & ORF6 Severe acute respiratory syndrome coronavirus 2 & 96.74\% & strong & likely \\ \hline
% NF-kB Ovis aries & M protein Severe acute respiratory syndrome-related coronavirus & 96.64\% & strong & likely \\ \hline
% M protein Severe acute respiratory syndrome-related coronavirus & TBK1 Rhinolophus ferrumequinum & 96.57\% & strong & likely \\ \hline
% nsp15 Severe acute respiratory syndrome-related coronavirus & RIG-I Canis lupus familiaris & 96.52\% & strong & likely \\ \hline
% nsp15 Severe acute respiratory syndrome coronavirus 2 & ACE2 Rhinolophus ferrumequinum & 96.49\% & strong & likely \\ \hline
% nsp15 Severe acute respiratory syndrome-related coronavirus & STAT2 Macaca mulatta & 96.44\% & strong & likely \\ \hline
% STAT1 Felis catus & nsp1 Severe acute respiratory syndrome coronavirus 2 & 96.43\% & strong & likely \\ \hline
% nsp15 Severe acute respiratory syndrome-related coronavirus & IRF9 Rattus norvegicus & 96.36\% & strong & likely \\ \hline
% ORF3b Severe acute respiratory syndrome-related coronavirus & MAVS Camelus dromedarius & 96.34\% & strong & likely \\ \hline
% STAT1 Rattus norvegicus & N protein Severe acute respiratory syndrome-related coronavirus & 96.34\% & strong & likely \\ \hline
% IRF7 Homo sapiens & PLpro Severe acute respiratory syndrome-related coronavirus & 96.29\% & strong & likely \\ \hline
% Spike Severe acute respiratory syndrome coronavirus 2 & ACE2 Rattus norvegicus & 96.29\% & strong & likely \\ \hline
% RIG-I Mus musculus & ORF6 Severe acute respiratory syndrome coronavirus 2 & 96.09\% & strong & likely \\ \hline
% STAT1 Macaca mulatta & ORF6 Severe acute respiratory syndrome coronavirus 2 & 95.99\% & strong & likely \\ \hline
% nsp15 Severe acute respiratory syndrome coronavirus 2 & IRF3 Ovis aries & 95.97\% & strong & likely \\ \hline
% IRF7 Rattus norvegicus & M protein Middle East respiratory syndrome-related coronavirus & 95.91\% & strong & likely \\ \hline
% RIG-I Homo sapiens & ORF6 Severe acute respiratory syndrome-related coronavirus & 95.87\% & strong & likely \\ \hline
% ORF4a Middle East respiratory syndrome-related coronavirus & TBK1 Mus musculus & 95.81\% & strong & likely \\ \hline
% NF-kB Camelus dromedarius & PLpro Severe acute respiratory syndrome-related coronavirus & 95.77\% & strong & likely \\ \hline
% STAT2 Rattus norvegicus & PLpro Middle East respiratory syndrome-related coronavirus & 95.71\% & strong & likely \\ \hline
% ORF6 Severe acute respiratory syndrome-related coronavirus & TBK1 Rhinolophus ferrumequinum & 95.70\% & strong & likely \\ \hline
% IRF3 Ictidomys tridecemlineatus & M protein Severe acute respiratory syndrome-related coronavirus & 95.55\% & strong & likely \\ \hline
% STAT1 Camelus dromedarius & PLpro Severe acute respiratory syndrome-related coronavirus & 95.48\% & strong & likely \\ \hline
% MDA5 Mus musculus & PLpro Severe acute respiratory syndrome-related coronavirus & 95.43\% & strong & likely \\ \hline
% Spike Human coronavirus 229E & ACE2 Homo sapiens & 95.39\% & strong & likely \\ \hline
% IRF3 Mesocricetus auratus & ORF6 Severe acute respiratory syndrome-related coronavirus & 95.34\% & strong & likely \\ \hline
% ORF4b Middle East respiratory syndrome-related coronavirus & MDA5 Mus musculus & 95.32\% & strong & likely \\ \hline
% STAT1 Rattus norvegicus & PLpro Middle East respiratory syndrome-related coronavirus & 95.18\% & strong & likely \\ \hline
% nsp1 Severe acute respiratory syndrome-related coronavirus & TBK1 Rattus norvegicus & 95.13\% & strong & likely \\ \hline
% ORF4b Middle East respiratory syndrome-related coronavirus & TBK1 Rhinolophus ferrumequinum & 95.10\% & strong & likely \\ \hline
% STAT1 Felis catus & ORF4a Middle East respiratory syndrome-related coronavirus & 95.07\% & strong & likely \\ \hline
% MDA5 Macaca mulatta & ORF6 Severe acute respiratory syndrome-related coronavirus & 94.99\% & strong & likely \\ \hline
% NF-kB Rhinolophus ferrumequinum & PLpro Middle East respiratory syndrome-related coronavirus & 94.98\% & strong & likely \\ \hline
% PLpro Severe acute respiratory syndrome-related coronavirus & TBK1 Mesocricetus auratus & 94.87\% & strong & likely \\ \hline
% ORF6 Severe acute respiratory syndrome-related coronavirus & TBK1 Mus musculus & 94.83\% & strong & likely \\ \hline
% N protein Severe acute respiratory syndrome-related coronavirus & STAT2 Rattus norvegicus & 94.74\% & strong & likely \\ \hline
% Spike Severe acute respiratory syndrome-related coronavirus & ACE2 Macaca mulatta & 94.70\% & strong & likely \\ \hline
% ORF4b Middle East respiratory syndrome-related coronavirus & PRKRA Mus musculus & 94.62\% & strong & likely \\ \hline
% NF-kB Homo sapiens & ORF3b Severe acute respiratory syndrome-related coronavirus & 94.58\% & strong & likely \\ \hline
% ORF3b Severe acute respiratory syndrome-related coronavirus & TBK1 Rhinolophus ferrumequinum & 94.57\% & strong & likely \\ \hline
% NF-kB Macaca mulatta & PLpro Severe acute respiratory syndrome-related coronavirus & 94.49\% & strong & likely \\ \hline
% MDA5 Homo sapiens & ORF6 Severe acute respiratory syndrome-related coronavirus & 94.37\% & strong & likely \\ \hline
% STAT2 Camelus dromedarius & M protein Severe acute respiratory syndrome-related coronavirus & 94.31\% & strong & likely \\ \hline
% IRF9 Rattus norvegicus & N protein Severe acute respiratory syndrome-related coronavirus & 94.03\% & strong & likely \\ \hline
% ORF4a Middle East respiratory syndrome-related coronavirus & TBK1 Camelus dromedarius & 93.77\% & strong & likely \\ \hline
% nsp15 Severe acute respiratory syndrome coronavirus 2 & RIG-I Gallus gallus & 93.75\% & strong & likely \\ \hline
% IRF3 Mesocricetus auratus & ORF3b Severe acute respiratory syndrome-related coronavirus & 93.69\% & strong & likely \\ \hline
% M protein Severe acute respiratory syndrome-related coronavirus & TBK1 Bos taurus & 93.59\% & strong & likely \\ \hline
% RIG-I Canis lupus familiaris & ORF6 Severe acute respiratory syndrome-related coronavirus & 93.59\% & strong & likely \\ \hline
% nsp1 Severe acute respiratory syndrome-related coronavirus & TBK1 Homo sapiens & 93.52\% & strong & likely \\ \hline
% NF-kB Mus musculus & M protein Middle East respiratory syndrome-related coronavirus & 93.42\% & strong & likely \\ \hline
% PRKRA Rattus norvegicus & ORF3b Severe acute respiratory syndrome-related coronavirus & 93.36\% & strong & likely \\ \hline
% NF-kB Mus musculus & ORF6 Severe acute respiratory syndrome-related coronavirus & 93.33\% & strong & likely \\ \hline
% nsp1 Severe acute respiratory syndrome-related coronavirus & IRF3 Rattus norvegicus & 93.22\% & strong & likely \\ \hline
% ORF3b Severe acute respiratory syndrome-related coronavirus & STAT2 Macaca mulatta & 93.20\% & strong & likely \\ \hline
% ORF6 Severe acute respiratory syndrome coronavirus 2 & STAT2 Homo sapiens & 93.19\% & strong & likely \\ \hline
% PLpro Middle East respiratory syndrome-related coronavirus & MAVS Rattus norvegicus & 93.18\% & strong & likely \\ \hline
% IRF7 Rattus norvegicus & ORF6 Severe acute respiratory syndrome-related coronavirus & 93.15\% & strong & likely \\ \hline
% N protein Severe acute respiratory syndrome-related coronavirus & STAT2 Rhinolophus ferrumequinum & 93.11\% & strong & likely \\ \hline
% N protein Middle East respiratory syndrome-related coronavirus & MAVS Homo sapiens & 93.09\% & strong & likely \\ \hline
% nsp15 Severe acute respiratory syndrome-related coronavirus & STAT1 Felis catus & 93.09\% & strong & likely \\ \hline
% nsp1 Severe acute respiratory syndrome-related coronavirus & MAVS Homo sapiens & 93.09\% & strong & likely \\ \hline
% IRF9 Homo sapiens & N protein Severe acute respiratory syndrome-related coronavirus & 92.95\% & strong & likely \\ \hline
% M protein Severe acute respiratory syndrome-related coronavirus & MAVS Mus musculus & 92.92\% & strong & likely \\ \hline
% M protein Severe acute respiratory syndrome-related coronavirus & TBK1 Mesocricetus auratus & 92.88\% & strong & likely \\ \hline
% STAT1 Rhinolophus ferrumequinum & ORF4b Middle East respiratory syndrome-related coronavirus & 92.86\% & strong & likely \\ \hline
% nsp15 Severe acute respiratory syndrome-related coronavirus & NF-kB Homo sapiens & 92.80\% & strong & likely \\ \hline
% NF-kB Ictidomys tridecemlineatus & PLpro Severe acute respiratory syndrome-related coronavirus & 92.79\% & strong & likely \\ \hline
% N protein Middle East respiratory syndrome-related coronavirus & MAVS Rattus norvegicus & 92.77\% & strong & likely \\ \hline
% nsp1 Middle East respiratory syndrome-related coronavirus & STAT2 Rattus norvegicus & 92.60\% & strong & likely \\ \hline
% ORF6 Severe acute respiratory syndrome coronavirus 2 & STAT2 Felis catus & 92.51\% & strong & likely \\ \hline
% ORF3b Severe acute respiratory syndrome-related coronavirus & TBK1 Mus musculus & 92.47\% & strong & likely \\ \hline
% STAT2 Rhinolophus ferrumequinum & PLpro Severe acute respiratory syndrome-related coronavirus & 92.17\% & strong & likely \\ \hline
% IRF9 Macaca mulatta & M protein Severe acute respiratory syndrome-related coronavirus & 92.05\% & strong & likely \\ \hline
% ORF6 Severe acute respiratory syndrome-related coronavirus & MAVS Homo sapiens & 91.99\% & strong & likely \\ \hline
% MDA5 Homo sapiens & ORF6 Severe acute respiratory syndrome coronavirus 2 & 91.89\% & strong & likely \\ \hline
% nsp15 Severe acute respiratory syndrome-related coronavirus & STAT1 Macaca mulatta & 91.67\% & strong & likely \\ \hline
% ORF4a Middle East respiratory syndrome-related coronavirus & NF-kB Rhinolophus ferrumequinum & 91.50\% & strong & likely \\ \hline
% MDA5 Felis catus & ORF6 Severe acute respiratory syndrome-related coronavirus & 91.18\% & strong & likely \\ \hline
% nsp15 Severe acute respiratory syndrome-related coronavirus & RIG-I Macaca mulatta & 91.17\% & strong & likely \\ \hline
% M protein Middle East respiratory syndrome-related coronavirus & MAVS Mus musculus & 90.93\% & strong & likely \\ \hline
% STAT1 Rattus norvegicus & nsp1 Middle East respiratory syndrome-related coronavirus & 90.85\% & strong & likely \\ \hline
% IRF7 Rattus norvegicus & PLpro Severe acute respiratory syndrome-related coronavirus & 90.82\% & strong & likely \\ \hline
% IRF3 Camelus dromedarius & ORF3b Severe acute respiratory syndrome-related coronavirus & 90.75\% & strong & likely \\ \hline
% NF-kB Rattus norvegicus & ORF6 Severe acute respiratory syndrome-related coronavirus & 90.61\% & strong & likely \\ \hline
% PLpro Severe acute respiratory syndrome-related coronavirus & MAVS Rattus norvegicus & 90.48\% & strong & likely \\ \hline
% N protein Middle East respiratory syndrome-related coronavirus & MAVS Mus musculus & 90.32\% & strong & likely \\ \hline
% IRF3 Rhinolophus ferrumequinum & ORF4a Middle East respiratory syndrome-related coronavirus & 90.28\% & strong & likely \\ \hline
% STAT1 Camelus dromedarius & M protein Severe acute respiratory syndrome-related coronavirus & 90.16\% & strong & likely \\ \hline
% Spike Human coronavirus NL63 & ACE2 Rattus norvegicus & 90.11\% & strong & likely \\ \hline
% NF-kB Camelus dromedarius & M protein Severe acute respiratory syndrome-related coronavirus & 90.08\% & strong & likely \\ \hline
% PRKRA Rattus norvegicus & PLpro Middle East respiratory syndrome-related coronavirus & 90.06\% & strong & likely \\ \hline
% PLpro Severe acute respiratory syndrome-related coronavirus & TBK1 Camelus dromedarius & 89.92\% & strong & likely \\ \hline
% Spike Human coronavirus HKU1 & ACE2 Mesocricetus auratus & 89.92\% & strong & likely \\ \hline
% ORF6 Severe acute respiratory syndrome-related coronavirus & MAVS Felis catus & 89.65\% & strong & likely \\ \hline
% nsp15 Severe acute respiratory syndrome coronavirus 2 & MAVS Rhinolophus ferrumequinum & 89.59\% & strong & likely \\ \hline
% STAT2 Camelus dromedarius & PLpro Severe acute respiratory syndrome-related coronavirus & 89.54\% & strong & likely \\ \hline
% nsp15 Severe acute respiratory syndrome coronavirus 2 & ACE2 Canis lupus familiaris & 89.47\% & strong & likely \\ \hline
% IRF3 Camelus dromedarius & M protein Severe acute respiratory syndrome-related coronavirus & 89.41\% & strong & likely \\ \hline
% RIG-I Macaca mulatta & N protein Severe acute respiratory syndrome-related coronavirus & 89.40\% & strong & likely \\ \hline
% ORF6 Severe acute respiratory syndrome coronavirus 2 & MAVS Rattus norvegicus & 89.28\% & strong & likely \\ \hline
% IRF9 Rhinolophus ferrumequinum & ORF4a Middle East respiratory syndrome-related coronavirus & 89.26\% & strong & likely \\ \hline
% STAT1 Rhinolophus ferrumequinum & nsp1 Severe acute respiratory syndrome coronavirus 2 & 89.26\% & strong & likely \\ \hline
% MDA5 Mus musculus & PLpro Middle East respiratory syndrome-related coronavirus & 89.26\% & strong & likely \\ \hline
% IRF9 Rhinolophus ferrumequinum & M protein Severe acute respiratory syndrome-related coronavirus & 89.11\% & strong & likely \\ \hline
% nsp1 Severe acute respiratory syndrome-related coronavirus & MDA5 Rattus norvegicus & 89.00\% & strong & likely \\ \hline
% IRF3 Equus caballus & M protein Severe acute respiratory syndrome-related coronavirus & 88.92\% & strong & likely \\ \hline
% ORF4b Middle East respiratory syndrome-related coronavirus & NF-kB Rhinolophus ferrumequinum & 88.87\% & strong & likely \\ \hline
% ORF4a Middle East respiratory syndrome-related coronavirus & IRF7 Mus musculus & 88.84\% & strong & likely \\ \hline
% NF-kB Ictidomys tridecemlineatus & M protein Severe acute respiratory syndrome-related coronavirus & 88.71\% & strong & likely \\ \hline
% NF-kB Rhinolophus ferrumequinum & M protein Middle East respiratory syndrome-related coronavirus & 88.60\% & strong & likely \\ \hline
% nsp15 Severe acute respiratory syndrome coronavirus 2 & STAT2 Felis catus & 88.42\% & strong & likely \\ \hline
% nsp15 Severe acute respiratory syndrome coronavirus 2 & MAVS Homo sapiens & 88.41\% & strong & likely \\ \hline
% RIG-I Pan troglodytes & ORF6 Severe acute respiratory syndrome coronavirus 2 & 88.34\% & strong & likely \\ \hline
% nsp1 Severe acute respiratory syndrome-related coronavirus & MDA5 Mus musculus & 88.22\% & strong & likely \\ \hline
% N protein Severe acute respiratory syndrome-related coronavirus & IRF7 Homo sapiens & 87.97\% & strong & likely \\ \hline
% STAT2 Felis catus & PLpro Severe acute respiratory syndrome-related coronavirus & 87.92\% & strong & likely \\ \hline
% STAT1 Ovis aries & ORF6 Severe acute respiratory syndrome-related coronavirus & 87.88\% & strong & likely \\ \hline
% NF-kB Bos taurus & PLpro Severe acute respiratory syndrome-related coronavirus & 87.81\% & strong & likely \\ \hline
% nsp1 Severe acute respiratory syndrome-related coronavirus & MDA5 Homo sapiens & 87.59\% & strong & likely \\ \hline
% ORF4b Middle East respiratory syndrome-related coronavirus & MDA5 Rattus norvegicus & 87.55\% & strong & likely \\ \hline
% ORF3b Severe acute respiratory syndrome-related coronavirus & STAT2 Canis lupus familiaris & 87.26\% & strong & likely \\ \hline
% MDA5 Rattus norvegicus & ORF6 Severe acute respiratory syndrome coronavirus 2 & 87.00\% & strong & likely \\ \hline
% IRF7 Camelus dromedarius & M protein Middle East respiratory syndrome-related coronavirus & 86.96\% & strong & likely \\ \hline
% MDA5 Mus musculus & M protein Severe acute respiratory syndrome-related coronavirus & 86.83\% & strong & likely \\ \hline
% IRF7 Rattus norvegicus & ORF3b Severe acute respiratory syndrome-related coronavirus & 86.69\% & strong & likely \\ \hline
% M protein Middle East respiratory syndrome-related coronavirus & TBK1 Felis catus & 86.45\% & strong & likely \\ \hline
% NF-kB Equus caballus & M protein Severe acute respiratory syndrome-related coronavirus & 86.30\% & strong & likely \\ \hline
% STAT2 Rhinolophus ferrumequinum & M protein Severe acute respiratory syndrome-related coronavirus & 86.29\% & strong & likely \\ \hline
% nsp15 Severe acute respiratory syndrome-related coronavirus & NF-kB Rhinolophus ferrumequinum & 86.25\% & strong & likely \\ \hline
% ORF4a Middle East respiratory syndrome-related coronavirus & MAVS Rattus norvegicus & 86.13\% & strong & likely \\ \hline
% ORF3b Severe acute respiratory syndrome-related coronavirus & STAT2 Mesocricetus auratus & 85.76\% & strong & likely \\ \hline
% STAT1 Rhinolophus ferrumequinum & M protein Severe acute respiratory syndrome-related coronavirus & 85.68\% & strong & likely \\ \hline
% IRF9 Felis catus & ORF6 Severe acute respiratory syndrome coronavirus 2 & 85.66\% & strong & likely \\ \hline
% STAT1 Mus musculus & N protein Severe acute respiratory syndrome-related coronavirus & 85.55\% & strong & likely \\ \hline
% M protein Middle East respiratory syndrome-related coronavirus & MAVS Homo sapiens & 85.43\% & strong & likely \\ \hline
% ORF6 Severe acute respiratory syndrome-related coronavirus & STAT2 Canis lupus familiaris & 85.42\% & strong & likely \\ \hline
% N protein Severe acute respiratory syndrome coronavirus 2 & MAVS Felis catus & 85.36\% & strong & likely \\ \hline
% ORF4b Middle East respiratory syndrome-related coronavirus & MDA5 Homo sapiens & 85.33\% & strong & likely \\ \hline
% nsp1 Severe acute respiratory syndrome coronavirus 2 & STAT2 Rhinolophus ferrumequinum & 85.26\% & strong & likely \\ \hline
% STAT2 Macaca mulatta & M protein Severe acute respiratory syndrome-related coronavirus & 85.09\% & strong & likely \\ \hline
% STAT1 Macaca mulatta & nsp1 Severe acute respiratory syndrome coronavirus 2 & 85.05\% & strong & likely \\ \hline
% PLpro Middle East respiratory syndrome-related coronavirus & MAVS Mus musculus & 84.96\% & strong & likely \\ \hline
% NF-kB Ovis aries & PLpro Middle East respiratory syndrome-related coronavirus & 84.74\% & strong & likely \\ \hline
% nsp15 Severe acute respiratory syndrome-related coronavirus & TBK1 Rattus norvegicus & 84.60\% & strong & likely \\ \hline


% NF-kB Mesocricetus auratus & PLpro Severe acute respiratory syndrome-related coronavirus & 84.43\% & strong & likely \\ \hline
% PLpro Middle East respiratory syndrome-related coronavirus & MAVS Homo sapiens & 83.94\% & strong & likely \\ \hline
% M protein Middle East respiratory syndrome-related coronavirus & MAVS Rattus norvegicus & 83.87\% & strong & likely \\ \hline
% nsp15 Severe acute respiratory syndrome-related coronavirus & MDA5 Mus musculus & 83.86\% & strong & likely \\ \hline
% STAT1 Felis catus & ORF6 Severe acute respiratory syndrome coronavirus 2 & 83.79\% & strong & likely \\ \hline
% IRF9 Rhinolophus ferrumequinum & ORF4b Middle East respiratory syndrome-related coronavirus & 83.73\% & strong & likely \\ \hline
% STAT1 Mesocricetus auratus & ORF3b Severe acute respiratory syndrome-related coronavirus & 83.53\% & strong & likely \\ \hline
% STAT2 Felis catus & M protein Severe acute respiratory syndrome-related coronavirus & 83.35\% & strong & likely \\ \hline
% IRF7 Mus musculus & ORF3b Severe acute respiratory syndrome-related coronavirus & 83.33\% & strong & likely \\ \hline
% nsp15 Severe acute respiratory syndrome-related coronavirus & NF-kB Rattus norvegicus & 83.04\% & strong & likely \\ \hline
% IRF3 Rhinolophus ferrumequinum & N protein Severe acute respiratory syndrome coronavirus 2 & 82.93\% & strong & likely \\ \hline
% NF-kB Homo sapiens & PLpro Human coronavirus OC43 & 82.69\% & strong & likely \\ \hline
% MDA5 Mus musculus & ORF6 Severe acute respiratory syndrome coronavirus 2 & 82.39\% & strong & likely \\ \hline
% nsp15 Severe acute respiratory syndrome-related coronavirus & IRF9 Mus musculus & 82.33\% & strong & likely \\ \hline
% ORF4a Middle East respiratory syndrome-related coronavirus & TBK1 Felis catus & 82.26\% & strong & likely \\ \hline
% ORF6 Severe acute respiratory syndrome-related coronavirus & MAVS Rhinolophus ferrumequinum & 82.25\% & strong & likely \\ \hline
% PLpro Severe acute respiratory syndrome-related coronavirus & MAVS Mus musculus & 82.18\% & strong & likely \\ \hline
% STAT1 Rhinolophus ferrumequinum & PLpro Severe acute respiratory syndrome-related coronavirus & 82.07\% & strong & likely \\ \hline
% IRF3 Rhinolophus ferrumequinum & PLpro Middle East respiratory syndrome-related coronavirus & 81.85\% & strong & likely \\ \hline
% STAT1 Rhinolophus ferrumequinum & ORF6 Severe acute respiratory syndrome coronavirus 2 & 81.81\% & strong & likely \\ \hline
% MDA5 Rattus norvegicus & M protein Middle East respiratory syndrome-related coronavirus & 81.59\% & strong & likely \\ \hline
% nsp1 Middle East respiratory syndrome-related coronavirus & IRF3 Rattus norvegicus & 81.50\% & strong & likely \\ \hline
% nsp15 Severe acute respiratory syndrome coronavirus 2 & MDA5 Rattus norvegicus & 81.35\% & strong & likely \\ \hline
% STAT1 Homo sapiens & ORF6 Severe acute respiratory syndrome coronavirus 2 & 81.04\% & strong & likely \\ \hline
% RIG-I Mus musculus & ORF6 Severe acute respiratory syndrome-related coronavirus & 80.83\% & strong & likely \\ \hline
% nsp15 Severe acute respiratory syndrome coronavirus 2 & ACE2 Mesocricetus auratus & 80.63\% & strong & likely \\ \hline
% IRF3 Bos taurus & M protein Severe acute respiratory syndrome-related coronavirus & 80.41\% & strong & likely \\ \hline
% IRF3 Camelus dromedarius & N protein Severe acute respiratory syndrome-related coronavirus & 80.39\% & strong & likely \\ \hline
% RIG-I Rhinolophus ferrumequinum & nsp1 Severe acute respiratory syndrome-related coronavirus & 80.23\% & strong & likely \\ \hline
% IRF3 Rhinolophus ferrumequinum & PLpro Severe acute respiratory syndrome coronavirus 2 & 80.23\% & strong & likely \\ \hline
% Spike Human coronavirus HKU1 & ACE2 Felis catus & 80.17\% & strong & likely \\ \hline
% STAT1 Macaca mulatta & ORF3b Severe acute respiratory syndrome-related coronavirus & 80.07\% & strong & likely \\ \hline
% RIG-I Canis lupus familiaris & ORF3b Severe acute respiratory syndrome-related coronavirus & 79.91\% & strong & likely \\ \hline
% ORF4b Middle East respiratory syndrome-related coronavirus & TBK1 Felis catus & 79.88\% & strong & likely \\ \hline
% nsp15 Severe acute respiratory syndrome-related coronavirus & TBK1 Homo sapiens & 79.67\% & strong & likely \\ \hline
% nsp15 Severe acute respiratory syndrome-related coronavirus & MDA5 Felis catus & 79.58\% & strong & likely \\ \hline
% ORF3b Severe acute respiratory syndrome-related coronavirus & MAVS Canis lupus familiaris & 79.39\% & strong & likely \\ \hline
% nsp15 Severe acute respiratory syndrome-related coronavirus & MDA5 Homo sapiens & 79.26\% & strong & likely \\ \hline
% MDA5 Mus musculus & M protein Middle East respiratory syndrome-related coronavirus & 79.17\% & strong & likely \\ \hline
% nsp1 Severe acute respiratory syndrome-related coronavirus & MAVS Rattus norvegicus & 79.08\% & strong & likely \\ \hline
% IRF7 Homo sapiens & PLpro Middle East respiratory syndrome-related coronavirus & 79.07\% & strong & likely \\ \hline
% ORF4a Middle East respiratory syndrome-related coronavirus & IRF7 Homo sapiens & 78.82\% & strong & likely \\ \hline
% M protein Severe acute respiratory syndrome-related coronavirus & TBK1 Ovis aries & 78.72\% & strong & likely \\ \hline
% IRF7 Mus musculus & PLpro Middle East respiratory syndrome-related coronavirus & 78.66\% & strong & likely \\ \hline
% ORF6 Severe acute respiratory syndrome-related coronavirus & STAT2 Ovis aries & 78.53\% & strong & likely \\ \hline
% M protein Middle East respiratory syndrome-related coronavirus & TBK1 Rhinolophus ferrumequinum & 78.44\% & strong & likely \\ \hline
% ACE2 Felis catus & ORF6 Severe acute respiratory syndrome coronavirus 2 & 78.20\% & strong & likely \\ \hline
% Spike Human coronavirus HKU1 & ACE2 Macaca mulatta & 78.05\% & strong & likely \\ \hline
% IRF3 Rhinolophus ferrumequinum & M protein Middle East respiratory syndrome-related coronavirus & 77.98\% & strong & likely \\ \hline
% IRF9 Camelus dromedarius & PLpro Severe acute respiratory syndrome-related coronavirus & 77.94\% & strong & likely \\ \hline
% nsp15 Severe acute respiratory syndrome coronavirus 2 & ACE2 Felis catus & 77.76\% & strong & likely \\ \hline
% Spike Human coronavirus HKU1 & ACE2 Mus musculus & 77.74\% & strong & likely \\ \hline
% IRF9 Mesocricetus auratus & ORF6 Severe acute respiratory syndrome-related coronavirus & 77.61\% & strong & likely \\ \hline
% nsp15 Severe acute respiratory syndrome-related coronavirus & MAVS Canis lupus familiaris & 77.39\% & strong & likely \\ \hline
% nsp15 Severe acute respiratory syndrome coronavirus 2 & IRF3 Mus musculus & 77.38\% & strong & likely \\ \hline
% nsp1 Severe acute respiratory syndrome coronavirus 2 & STAT2 Macaca mulatta & 77.21\% & strong & likely \\ \hline
% nsp15 Severe acute respiratory syndrome-related coronavirus & STAT2 Mus musculus & 77.17\% & strong & likely \\ \hline
% nsp1 Severe acute respiratory syndrome-related coronavirus & MAVS Rhinolophus ferrumequinum & 77.10\% & strong & likely \\ \hline
% nsp15 Severe acute respiratory syndrome coronavirus 2 & MDA5 Homo sapiens & 77.01\% & strong & likely \\ \hline
% NF-kB Mus musculus & ORF3b Severe acute respiratory syndrome-related coronavirus & 76.93\% & strong & likely \\ \hline
% nsp15 Severe acute respiratory syndrome-related coronavirus & IRF7 Homo sapiens & 76.92\% & strong & likely \\ \hline
% STAT1 Macaca mulatta & PLpro Severe acute respiratory syndrome-related coronavirus & 76.90\% & strong & likely \\ \hline
% nsp1 Severe acute respiratory syndrome-related coronavirus & IRF3 Mus musculus & 76.63\% & strong & likely \\ \hline
% ORF4a Middle East respiratory syndrome-related coronavirus & MAVS Homo sapiens & 76.32\% & strong & likely \\ \hline
% IRF3 Mesocricetus auratus & PLpro Severe acute respiratory syndrome-related coronavirus & 76.30\% & strong & likely \\ \hline
% ORF6 Severe acute respiratory syndrome coronavirus 2 & MAVS Mus musculus & 76.20\% & strong & likely \\ \hline
% PRKRA Homo sapiens & ORF3b Severe acute respiratory syndrome-related coronavirus & 76.12\% & strong & likely \\ \hline
% RIG-I Rhinolophus ferrumequinum & M protein Severe acute respiratory syndrome-related coronavirus & 75.97\% & strong & likely \\ \hline
% NF-kB Mesocricetus auratus & M protein Severe acute respiratory syndrome-related coronavirus & 75.83\% & strong & likely \\ \hline
% PRKRA Rattus norvegicus & M protein Middle East respiratory syndrome-related coronavirus & 75.71\% & strong & likely \\ \hline
% IRF3 Canis lupus familiaris & ORF3b Severe acute respiratory syndrome-related coronavirus & 75.33\% & strong & likely \\ \hline
% PLpro Middle East respiratory syndrome-related coronavirus & TBK1 Felis catus & 75.30\% & strong & likely \\ \hline
% IRF3 Homo sapiens & N protein Middle East respiratory syndrome-related coronavirus & 75.28\% & strong & likely \\ \hline
% IRF7 Mus musculus & PLpro Severe acute respiratory syndrome-related coronavirus & 75.23\% & strong & likely \\ \hline
% IRF9 Mus musculus & nsp1 Severe acute respiratory syndrome-related coronavirus & 75.21\% & strong & likely \\ \hline
% Spike Severe acute respiratory syndrome coronavirus 2 & ACE2 Equus caballus & 75.15\% & strong & likely \\ \hline
% M protein Severe acute respiratory syndrome-related coronavirus & MAVS Homo sapiens & 75.14\% & strong & likely \\ \hline
% ORF4b Middle East respiratory syndrome-related coronavirus & MAVS Rhinolophus ferrumequinum & 74.84\% & strong & likely \\ \hline
% ORF6 Severe acute respiratory syndrome-related coronavirus & MAVS Mus musculus & 74.39\% & strong & likely \\ \hline
% Spike Severe acute respiratory syndrome coronavirus 2 & ACE2 Ictidomys tridecemlineatus & 74.37\% & strong & likely \\ \hline
% MDA5 Homo sapiens & PLpro Middle East respiratory syndrome-related coronavirus & 74.36\% & strong & likely \\ \hline
% nsp1 Severe acute respiratory syndrome-related coronavirus & TBK1 Mus musculus & 74.34\% & strong & likely \\ \hline
% IRF9 Rhinolophus ferrumequinum & ORF6 Severe acute respiratory syndrome coronavirus 2 & 74.30\% & strong & likely \\ \hline
% IRF7 Mus musculus & ORF6 Severe acute respiratory syndrome-related coronavirus & 73.98\% & strong & likely \\ \hline
% Spike Severe acute respiratory syndrome coronavirus 2 & ACE2 Ovis aries & 73.87\% & strong & likely \\ \hline
% PLpro Severe acute respiratory syndrome-related coronavirus & MAVS Homo sapiens & 73.76\% & strong & likely \\ \hline
% nsp15 Severe acute respiratory syndrome-related coronavirus & TBK1 Rhinolophus ferrumequinum & 73.63\% & strong & likely \\ \hline
% Spike Severe acute respiratory syndrome coronavirus 2 & ACE2 Bos taurus & 73.57\% & strong & likely \\ \hline
% nsp1 Middle East respiratory syndrome-related coronavirus & STAT2 Homo sapiens & 73.24\% & strong & likely \\ \hline
% IRF3 Gallus gallus & ORF6 Severe acute respiratory syndrome coronavirus 2 & 73.10\% & strong & likely \\ \hline
% STAT2 Felis catus & M protein Middle East respiratory syndrome-related coronavirus & 72.87\% & strong & likely \\ \hline
% RIG-I Rhinolophus ferrumequinum & N protein Severe acute respiratory syndrome coronavirus 2 & 72.85\% & strong & likely \\ \hline
% nsp1 Severe acute respiratory syndrome-related coronavirus & MDA5 Felis catus & 72.74\% & strong & likely \\ \hline
% MDA5 Macaca mulatta & ORF6 Severe acute respiratory syndrome coronavirus 2 & 72.71\% & strong & likely \\ \hline
% PLpro Severe acute respiratory syndrome-related coronavirus & TBK1 Bos taurus & 72.71\% & strong & likely \\ \hline
% nsp1 Severe acute respiratory syndrome-related coronavirus & STAT2 Mesocricetus auratus & 72.58\% & strong & likely \\ \hline
% NF-kB Homo sapiens & PLpro Severe acute respiratory syndrome coronavirus 2 & 72.37\% & strong & likely \\ \hline
% N protein Severe acute respiratory syndrome-related coronavirus & MAVS Camelus dromedarius & 72.37\% & strong & likely \\ \hline
% nsp15 Severe acute respiratory syndrome coronavirus 2 & MAVS Mesocricetus auratus & 72.27\% & strong & likely \\ \hline
% IRF7 Rattus norvegicus & PLpro Middle East respiratory syndrome-related coronavirus & 72.27\% & strong & likely \\ \hline
% Spike Human coronavirus NL63 & ACE2 Mus musculus & 72.04\% & strong & likely \\ \hline
% STAT1 Canis lupus familiaris & ORF3b Severe acute respiratory syndrome-related coronavirus & 71.87\% & strong & likely \\ \hline
% NF-kB Camelus dromedarius & ORF6 Severe acute respiratory syndrome-related coronavirus & 71.67\% & strong & likely \\ \hline
% Spike Human coronavirus NL63 & ACE2 Macaca mulatta & 71.62\% & strong & likely \\ \hline
% ORF4a Middle East respiratory syndrome-related coronavirus & STAT2 Ovis aries & 70.64\% & strong & likely \\ \hline
% nsp15 Severe acute respiratory syndrome-related coronavirus & TBK1 Mus musculus & 70.57\% & strong & likely \\ \hline
% nsp15 Severe acute respiratory syndrome-related coronavirus & IRF9 Macaca mulatta & 70.52\% & strong & likely \\ \hline
% Spike Severe acute respiratory syndrome coronavirus 2 & ACE2 Pan troglodytes & 70.44\% & strong & likely \\ \hline
% IRF9 Rhinolophus ferrumequinum & PLpro Severe acute respiratory syndrome-related coronavirus & 69.79\% & strong & likely \\ \hline
% STAT1 Camelus dromedarius & ORF3b Severe acute respiratory syndrome-related coronavirus & 69.79\% & strong & likely \\ \hline
% nsp15 Severe acute respiratory syndrome-related coronavirus & NF-kB Mus musculus & 69.20\% & strong & likely \\ \hline
% M protein Severe acute respiratory syndrome-related coronavirus & TBK1 Felis catus & 69.03\% & strong & likely \\ \hline
% IRF9 Camelus dromedarius & ORF6 Severe acute respiratory syndrome-related coronavirus & 68.99\% & strong & likely \\ \hline
% NF-kB Mesocricetus auratus & PLpro Middle East respiratory syndrome-related coronavirus & 68.80\% & strong & likely \\ \hline
% Spike Human coronavirus 229E & ACE2 Mesocricetus auratus & 68.10\% & strong & likely \\ \hline
% nsp15 Severe acute respiratory syndrome coronavirus 2 & MAVS Felis catus & 67.96\% & strong & likely \\ \hline
% nsp15 Severe acute respiratory syndrome-related coronavirus & MAVS Mesocricetus auratus & 67.88\% & strong & likely \\ \hline
% M protein Severe acute respiratory syndrome-related coronavirus & MAVS Rhinolophus ferrumequinum & 67.75\% & strong & likely \\ \hline
% PRKRA Mus musculus & ORF3b Severe acute respiratory syndrome-related coronavirus & 66.96\% & strong & likely \\ \hline
% IRF7 Rhinolophus ferrumequinum & ORF6 Severe acute respiratory syndrome-related coronavirus & 66.71\% & strong & likely \\ \hline
% IRF9 Felis catus & ORF4a Middle East respiratory syndrome-related coronavirus & 66.59\% & strong & likely \\ \hline
% PLpro Middle East respiratory syndrome-related coronavirus & TBK1 Rhinolophus ferrumequinum & 65.96\% & strong & likely \\ \hline
% IRF9 Macaca mulatta & ORF6 Severe acute respiratory syndrome-related coronavirus & 65.85\% & strong & likely \\ \hline
% NF-kB Gallus gallus & M protein Severe acute respiratory syndrome-related coronavirus & 65.68\% & strong & likely \\ \hline
% N protein Severe acute respiratory syndrome-related coronavirus & MDA5 Canis lupus familiaris & 65.52\% & strong & likely \\ \hline
% NF-kB Macaca mulatta & M protein Severe acute respiratory syndrome-related coronavirus & 65.30\% & strong & likely \\ \hline
% IRF9 Ovis aries & ORF4b Middle East respiratory syndrome-related coronavirus & 64.84\% & strong & likely \\ \hline
% PLpro Severe acute respiratory syndrome-related coronavirus & MAVS Rhinolophus ferrumequinum & 64.08\% & strong & likely \\ \hline
% Spike Human coronavirus HKU1 & ACE2 Canis lupus familiaris & 63.55\% & strong & likely \\ \hline
% PRKRA Rattus norvegicus & PLpro Severe acute respiratory syndrome-related coronavirus & 62.76\% & strong & likely \\ \hline
% nsp15 Severe acute respiratory syndrome coronavirus 2 & STAT1 Macaca mulatta & 62.75\% & strong & likely \\ \hline
% PRKRA Camelus dromedarius & PLpro Middle East respiratory syndrome-related coronavirus & 62.50\% & strong & likely \\ \hline
% RIG-I Mesocricetus auratus & ORF3b Severe acute respiratory syndrome-related coronavirus & 61.87\% & strong & likely \\ \hline
% nsp15 Severe acute respiratory syndrome coronavirus 2 & RIG-I Pan troglodytes & 61.48\% & strong & likely \\ \hline
% IRF9 Felis catus & ORF4b Middle East respiratory syndrome-related coronavirus & 60.90\% & strong & likely \\ \hline
% IRF9 Rattus norvegicus & nsp1 Middle East respiratory syndrome-related coronavirus & 60.20\% & strong & likely \\ \hline
% IRF3 Homo sapiens & PLpro Severe acute respiratory syndrome coronavirus 2 & 59.38\% & strong & likely \\ \hline
% IRF3 Mesocricetus auratus & N protein Severe acute respiratory syndrome-related coronavirus & 59.38\% & strong & likely \\ \hline
% Spike Human coronavirus NL63 & ACE2 Mesocricetus auratus & 59.37\% & strong & likely \\ \hline
% PLpro Middle East respiratory syndrome-related coronavirus & MAVS Camelus dromedarius & 58.43\% & strong & likely \\ \hline
% RIG-I Canis lupus familiaris & N protein Severe acute respiratory syndrome-related coronavirus & 56.60\% & strong & likely \\ \hline
% IRF3 Camelus dromedarius & ORF6 Severe acute respiratory syndrome-related coronavirus & 56.57\% & strong & likely \\ \hline
% PRKRA Homo sapiens & ORF6 Severe acute respiratory syndrome-related coronavirus & 56.14\% & strong & likely \\ \hline
% nsp1 Severe acute respiratory syndrome coronavirus 2 & STAT2 Mesocricetus auratus & 55.35\% & strong & likely \\ \hline
% nsp15 Severe acute respiratory syndrome coronavirus 2 & MAVS Mus musculus & 54.79\% & strong & likely \\ \hline
% Spike Severe acute respiratory syndrome coronavirus 2 & IRF3 Rhinolophus ferrumequinum & 89.09\% & strong & unlikely \\ \hline
% Spike Severe acute respiratory syndrome-related coronavirus & IRF3 Rhinolophus ferrumequinum & 81.84\% & strong & unlikely \\ \hline
% Spike Severe acute respiratory syndrome coronavirus 2 & IRF3 Canis lupus familiaris & 72.57\% & strong & unlikely \\ \hline
% \end{longtable}
% \label{tab: PPI prediction}
% % \printbibliography
% %TC:endignore
% %the command above ignores this section for word count

\end{document}
