## Supplementary material for "Network-based Virus-Host Interaction Prediction with Application to SARS-CoV-2": Latex Package: ref.bib

@article{frieman2008sars,
  title={SARS coronavirus and innate immunity},
  author={Frieman, Matthew and Heise, Mark and Baric, Ralph},
  journal={Virus research},
  volume={133},
  number={1},
  pages={101--112},
  year={2008},
  publisher={Elsevier}
}

@article{messina2020covid,
  title={COVID-19: viral--host interactome analyzed by network based-approach model to study pathogenesis of SARS-CoV-2 infection},
  author={Messina, Francesco and Giombini, Emanuela and Agrati, Chiara and Vairo, Francesco and Bartoli, Tommaso Ascoli and Al Moghazi, Samir and Piacentini, Mauro and Locatelli, Franco and Kobinger, Gary and Maeurer, Markus and others},
  journal={Journal of Translational Medicine},
  volume={18},
  number={1},
  pages={1--10},
  year={2020},
  publisher={Springer}
}
    
@article{yuen2020sars,
  title={SARS-CoV-2 nsp13, nsp14, nsp15 and orf6 function as potent interferon antagonists},
  author={Yuen, Chun-Kit and Lam, Joy-Yan and Wong, Wan-Man and Mak, Long-Fung and Wang, Xiaohui and Chu, Hin and Cai, Jian-Piao and Jin, Dong-Yan and To, Kelvin Kai-Wang and Chan, Jasper Fuk-Woo and others},
  journal={Emerging Microbes \& Infections},
  pages={1--29},
  year={2020},
  publisher={Taylor \& Francis}
}

@article{gordon2020sars,
  title={A SARS-CoV-2 protein interaction map reveals targets for drug repurposing},
  author={Gordon, David E and Jang, Gwendolyn M and Bouhaddou, Mehdi and Xu, Jiewei and Obernier, Kirsten and White, Kris M and O’Meara, Matthew J and Rezelj, Veronica V and Guo, Jeffrey Z and Swaney, Danielle L and others},
  journal={Nature},
  pages={1--13},
  year={2020},
  publisher={Nature Publishing Group}
}

@article{ziegler2020sars,
  title={SARS-CoV-2 receptor ACE2 is an interferon-stimulated gene in human airway epithelial cells and is detected in specific cell subsets across tissues},
  author={Ziegler, Carly GK and Allon, Samuel J and Nyquist, Sarah K and Mbano, Ian M and Miao, Vincent N and Tzouanas, Constantine N and Cao, Yuming and Yousif, Ashraf S and Bals, Julia and Hauser, Blake M and others},
  journal={Cell},
  year={2020},
  publisher={Elsevier}
}

@article{matthews2014sars,
  title={The SARS coronavirus papain like protease can inhibit IRF3 at a post activation step that requires deubiquitination activity},
  author={Matthews, Krystal and Sch{\"a}fer, Alexandra and Pham, Alissa and Frieman, Matthew},
  journal={Virology journal},
  volume={11},
  number={1},
  pages={209},
  year={2014},
  publisher={Springer}
}

@article{kopecky2007severe,
  title={Severe acute respiratory syndrome coronavirus open reading frame (ORF) 3b, ORF 6, and nucleocapsid proteins function as interferon antagonists},
  author={Kopecky-Bromberg, Sarah A and Mart{\'\i}nez-Sobrido, Luis and Frieman, Matthew and Baric, Ralph A and Palese, Peter},
  journal={Journal of virology},
  volume={81},
  number={2},
  pages={548--557},
  year={2007},
  publisher={Am Soc Microbiol}
}

@article{totura2012sars,
  title={SARS coronavirus pathogenesis: host innate immune responses and viral antagonism of interferon},
  author={Totura, Allison L and Baric, Ralph S},
  journal={Current opinion in virology},
  volume={2},
  number={3},
  pages={264--275},
  year={2012},
  publisher={Elsevier}
}

@article{hu2017sars,
  title={SARS coronavirus nucleocapsid inhibits type I interferon production by interfering with TRIM25-mediated RIG-I ubiquitination},
  author={Hu, Yong and Li, Wei and Gao, Ting and Cui, Yan and Jin, Yanwen and Li, Ping and Ma, Qingjun and Liu, Xuan and Cao, Cheng},
  journal={Journal of Virology},
  year={2017},
  publisher={Am Soc Microbiol}
}

@article{bailey2014crystal,
  title={Crystal structure of the Middle East respiratory syndrome coronavirus (MERS-CoV) papain-like protease bound to ubiquitin facilitates targeted disruption of deubiquitinating activity to demonstrate its role in innate immune suppression},
  author={Bailey-Elkin, Ben A and Knaap, Robert CM and Johnson, Garrett G and Dalebout, Tim J and Ninaber, Dennis K and Van Kasteren, Puck B and Bredenbeek, Peter J and Snijder, Eric J and Kikkert, Marjolein and Mark, Brian L},
  journal={Journal of Biological Chemistry},
  volume={289},
  number={50},
  pages={34667--34682},
  year={2014},
  publisher={ASBMB}
}

@article{lui2016middle,
  title={Middle East respiratory syndrome coronavirus M protein suppresses type I interferon expression through the inhibition of TBK1-dependent phosphorylation of IRF3},
  author={Lui, Pak-Yin and Wong, Lok-Yin Roy and Fung, Cheuk-Lai and Siu, Kam-Leung and Yeung, Man-Lung and Yuen, Kit-San and Chan, Chi-Ping and Woo, Patrick Chiu-Yat and Yuen, Kwok-Yung and Jin, Dong-Yan},
  journal={Emerging microbes \& infections},
  volume={5},
  number={1},
  pages={1--9},
  year={2016},
  publisher={Taylor \& Francis}
}

@article{tamura2008irf,
  title={The IRF family transcription factors in immunity and oncogenesis},
  author={Tamura, Tomohiko and Yanai, Hideyuki and Savitsky, David and Taniguchi, Tadatsugu},
  journal={Annu. Rev. Immunol.},
  volume={26},
  pages={535--584},
  year={2008},
  publisher={Annual Reviews}
}

@article{sato2000distinct,
  title={Distinct and essential roles of transcription factors IRF-3 and IRF-7 in response to viruses for IFN-$\alpha$/$\beta$ gene induction},
  author={Sato, Mitsuharu and Suemori, Hirofumi and Hata, Naoki and Asagiri, Masataka and Ogasawara, Kouetsu and Nakao, Kazuki and Nakaya, Takeo and Katsuki, Motoya and Noguchi, Shigeru and Tanaka, Nobuyuki and others},
  journal={Immunity},
  volume={13},
  number={4},
  pages={539--548},
  year={2000},
  publisher={Elsevier}
}

@article{siu2009severe,
  title={Severe acute respiratory syndrome coronavirus M protein inhibits type I interferon production by impeding the formation of TRAF3{\textperiodcentered} TANK{\textperiodcentered} TBK1/IKK$\epsilon$ complex},
  author={Siu, Kam-Leung and Kok, Kin-Hang and Ng, Ming-Him James and Poon, Vincent KM and Yuen, Kwok-Yung and Zheng, Bo-Jian and Jin, Dong-Yan},
  journal={Journal of Biological Chemistry},
  volume={284},
  number={24},
  pages={16202--16209},
  year={2009},
  publisher={ASBMB}
}

@article{chen2014sars,
  title={SARS coronavirus papain-like protease inhibits the type I interferon signaling pathway through interaction with the STING-TRAF3-TBK1 complex},
  author={Chen, Xiaojuan and Yang, Xingxing and Zheng, Yang and Yang, Yudong and Xing, Yaling and Chen, Zhongbin},
  journal={Protein \& cell},
  volume={5},
  number={5},
  pages={369--381},
  year={2014},
  publisher={Springer}
}

@article{sun2012coronavirus,
  title={Coronavirus papain-like proteases negatively regulate antiviral innate immune response through disruption of STING-mediated signaling},
  author={Sun, Li and Xing, Yaling and Chen, Xiaojuan and Zheng, Yang and Yang, Yudong and Nichols, Daniel B and Clementz, Mark A and Banach, Bridget S and Li, Kui and Baker, Susan C and others},
  journal={PloS one},
  volume={7},
  number={2},
  pages={e30802},
  year={2012},
  publisher={Public Library of Science}
}

@article{frieman2007severe,
  title={Severe acute respiratory syndrome coronavirus ORF6 antagonizes STAT1 function by sequestering nuclear import factors on the rough endoplasmic reticulum/Golgi membrane},
  author={Frieman, Matthew and Yount, Boyd and Heise, Mark and Kopecky-Bromberg, Sarah A and Palese, Peter and Baric, Ralph S},
  journal={Journal of virology},
  volume={81},
  number={18},
  pages={9812--9824},
  year={2007},
  publisher={Am Soc Microbiol}
}

@article{netland2008severe,
  title={Severe acute respiratory syndrome coronavirus infection causes neuronal death in the absence of encephalitis in mice transgenic for human ACE2},
  author={Netland, Jason and Meyerholz, David K and Moore, Steven and Cassell, Martin and Perlman, Stanley},
  journal={Journal of virology},
  volume={82},
  number={15},
  pages={7264--7275},
  year={2008},
  publisher={Am Soc Microbiol}
}

@article{cockrell2016mouse,
  title={A mouse model for MERS coronavirus-induced acute respiratory distress syndrome},
  author={Cockrell, Adam S and Yount, Boyd L and Scobey, Trevor and Jensen, Kara and Douglas, Madeline and Beall, Anne and Tang, Xian-Chun and Marasco, Wayne A and Heise, Mark T and Baric, Ralph S},
  journal={Nature microbiology},
  volume={2},
  number={2},
  pages={1--11},
  year={2016},
  publisher={Nature Publishing Group}
}

@article{niu2020non,
  title={Non-invasive bioluminescence imaging of HCoV-OC43 infection and therapy in the central nervous system of live mice},
  author={Niu, Junwei and Shen, Liang and Huang, Baoying and Ye, Fei and Zhao, Li and Wang, Huijuan and Deng, Yao and Tan, Wenjie},
  journal={Antiviral research},
  volume={173},
  pages={104646},
  year={2020},
  publisher={Elsevier}
}

@article{gorse2009human,
  title={Human coronavirus and acute respiratory illness in older adults with chronic obstructive pulmonary disease},
  author={Gorse, Geoffrey J and O’Connor, Theresa Z and Hall, Susan L and Vitale, Joseph N and Nichol, Kristin L},
  journal={The Journal of infectious diseases},
  volume={199},
  number={6},
  pages={847--857},
  year={2009},
  publisher={The University of Chicago Press}
}

@article{lim2016human,
  title={Human coronaviruses: a review of virus--host interactions},
  author={Lim, Yvonne Xinyi and Ng, Yan Ling and Tam, James P and Liu, Ding Xiang},
  journal={Diseases},
  volume={4},
  number={3},
  pages={26},
  year={2016},
  publisher={Multidisciplinary Digital Publishing Institute}
}

@article{lu2020genomic,
  title={Genomic characterisation and epidemiology of 2019 novel coronavirus: implications for virus origins and receptor binding},
  author={Lu, Roujian and Zhao, Xiang and Li, Juan and Niu, Peihua and Yang, Bo and Wu, Honglong and Wang, Wenling and Song, Hao and Huang, Baoying and Zhu, Na and others},
  journal={The Lancet},
  volume={395},
  number={10224},
  pages={565--574},
  year={2020},
  publisher={Elsevier}
}

@article{szczepanski2019canine,
  title={Canine respiratory coronavirus, bovine coronavirus, and human coronavirus OC43: receptors and attachment factors},
  author={Szczepanski, Artur and Owczarek, Katarzyna and Bzowska, Monika and Gula, Katarzyna and Drebot, Inga and Ochman, Marek and Maksym, Beata and Rajfur, Zenon and Mitchell, Judy A and Pyrc, Krzysztof},
  journal={Viruses},
  volume={11},
  number={4},
  pages={328},
  year={2019},
  publisher={Multidisciplinary Digital Publishing Institute}
}

@article{mikolov2013efficient,
  title={Efficient estimation of word representations in vector space},
  author={Mikolov, Tomas and Chen, Kai and Corrado, Greg and Dean, Jeffrey},
  journal={arXiv preprint arXiv:1301.3781},
  year={2013}
}

@inproceedings{Perozzi:2014:DOL:2623330.2623732,
 author = {Perozzi, Bryan and Al-Rfou, Rami and Skiena, Steven},
 title = {DeepWalk: Online Learning of Social Representations},
 booktitle = {Proceedings of the 20th ACM SIGKDD International Conference on Knowledge Discovery and Data Mining},
 series = {KDD '14},
 year = {2014},
 isbn = {978-1-4503-2956-9},
 location = {New York, New York, USA},
 pages = {701--710},
 numpages = {10},
 url = {http://doi.acm.org/10.1145/2623330.2623732},
 doi = {10.1145/2623330.2623732},
 acmid = {2623732},
 publisher = {ACM},
 address = {New York, NY, USA},
 keywords = {deep learning, latent representations, learning with partial labels, network classification, online learning, social networks},
}

@inproceedings{kipf2017semi,
  title={Semi-Supervised Classification with Graph Convolutional Networks},
  author={Kipf, Thomas N. and Welling, Max},
  booktitle={International Conference on Learning Representations (ICLR)},
  year={2017}
}

@inproceedings{mikolov2013distributed,
  title={Distributed representations of words and phrases and their compositionality},
  author={Mikolov, Tomas and Sutskever, Ilya and Chen, Kai and Corrado, Greg S and Dean, Jeff},
  booktitle={Advances in neural information processing systems},
  pages={3111--3119},
  year={2013}
}

@inproceedings{grover2016node2vec,
  title={node2vec: Scalable feature learning for networks},
  author={Grover, Aditya and Leskovec, Jure},
  booktitle={Proceedings of the 22nd ACM SIGKDD international conference on Knowledge discovery and data mining},
  pages={855--864},
  year={2016}
}

@inproceedings{tang2015line,
  title={LINE: Large-scale Information Network Embedding.},
  author={Tang, Jian and Qu, Meng and Wang, Mingzhe and Zhang, Ming and Yan, Jun and Mei, Qiaozhu},
  booktitle={WWW},
  year={2015},
  organization={ACM}
}

@article{storey2003positive,
  title={The positive false discovery rate: a Bayesian interpretation and the q-value},
  author={Storey, John D and others},
  journal={The Annals of Statistics},
  volume={31},
  number={6},
  pages={2013--2035},
  year={2003},
  publisher={Institute of Mathematical Statistics}
}

@article{shang2020structural,
  title={Structural basis of receptor recognition by SARS-CoV-2},
  author={Shang, Jian and Ye, Gang and Shi, Ke and Wan, Yushun and Luo, Chuming and Aihara, Hideki and Geng, Qibin and Auerbach, Ashley and Li, Fang},
  journal={Nature},
  volume={581},
  number={7807},
  pages={221--224},
  year={2020},
  publisher={Nature Publishing Group}
}

@article{matsumiya2010function,
  title={Function and regulation of retinoic acid-inducible gene-I},
  author={Matsumiya, Tomoh and Stafforini, Diana M},
  journal={Critical Reviews™ in Immunology},
  volume={30},
  number={6},
  year={2010},
  publisher={Begel House Inc.}
}

@article{ramos2011rig,
  title={RIG-I like receptors and their signaling crosstalk in the regulation of antiviral immunity},
  author={Ramos, Hilario J and Gale Jr, Michael},
  journal={Current opinion in virology},
  volume={1},
  number={3},
  pages={167--176},
  year={2011},
  publisher={Elsevier}
}

@article{ivashkiv2014regulation,
  title={Regulation of type I interferon responses},
  author={Ivashkiv, Lionel B and Donlin, Laura T},
  journal={Nature reviews Immunology},
  volume={14},
  number={1},
  pages={36--49},
  year={2014},
  publisher={Nature Publishing Group}
}

@article{schneider2014interferon,
  title={Interferon-stimulated genes: a complex web of host defenses},
  author={Schneider, William M and Chevillotte, Meike Dittmann and Rice, Charles M},
  journal={Annual review of immunology},
  volume={32},
  pages={513--545},
  year={2014},
  publisher={Annual Reviews}
}

@incollection{li2010host,
  title={Host immune responses to SARS coronavirus in humans},
  author={Li, Chris Ka-fai and Xu, Xiaoning},
  booktitle={Molecular biology of the SARS-coronavirus},
  pages={259--278},
  year={2010},
  publisher={Springer}
}

@article{liang2020highlight,
  title={Highlight of Immune Pathogenic Response and Hematopathologic Effect in SARS-CoV, MERS-CoV, and SARS-Cov-2 Infection},
  author={Liang, Yanwen and Wang, Mong-Lien and Chien, Chian-Shiu and Yarmishyn, Aliaksandr A and Yang, Yi-Ping and Lai, Wei-Yi and Luo, Yung-Hung and Lin, Yi-Tsung and Chen, Yann-Jang and Chang, Pei-Ching and others},
  journal={Frontiers in Immunology},
  volume={11},
  pages={1022},
  year={2020},
  publisher={Frontiers}
}

@article{al2017middle,
  title={Middle east respiratory syndrome corona virus spike glycoprotein suppresses macrophage responses via DPP4-mediated induction of IRAK-M and PPAR$\gamma$},
  author={Al-Qahtani, Ahmed A and Lyroni, Konstantina and Aznaourova, Marina and Tseliou, Melpomeni and Al-Anazi, Mashael R and Al-Ahdal, Mohammed N and Alkahtani, Saad and Sourvinos, George and Tsatsanis, Christos},
  journal={Oncotarget},
  volume={8},
  number={6},
  pages={9053},
  year={2017},
  publisher={Impact Journals, LLC}
}

@article{narayanan2008severe,
  title={Severe acute respiratory syndrome coronavirus nsp1 suppresses host gene expression, including that of type I interferon, in infected cells},
  author={Narayanan, Krishna and Huang, Cheng and Lokugamage, Kumari and Kamitani, Wataru and Ikegami, Tetsuro and Tseng, Chien-Te K and Makino, Shinji},
  journal={Journal of virology},
  volume={82},
  number={9},
  pages={4471--4479},
  year={2008},
  publisher={Am Soc Microbiol}
}

@article{mou2013receptor,
  title={The receptor binding domain of the new Middle East respiratory syndrome coronavirus maps to a 231-residue region in the spike protein that efficiently elicits neutralizing antibodies},
  author={Mou, Huihui and Raj, V Stalin and Van Kuppeveld, Frank JM and Rottier, Peter JM and Haagmans, Bart L and Bosch, Berend Jan},
  journal={Journal of virology},
  volume={87},
  number={16},
  pages={9379--9383},
  year={2013},
  publisher={Am Soc Microbiol}
}

@article{zhang2020angiotensin,
  title={Angiotensin-converting enzyme 2 (ACE2) as a SARS-CoV-2 receptor: molecular mechanisms and potential therapeutic target},
  author={Zhang, Haibo and Penninger, Josef M and Li, Yimin and Zhong, Nanshan and Slutsky, Arthur S},
  journal={Intensive care medicine},
  volume={46},
  number={4},
  pages={586--590},
  year={2020},
  publisher={Springer}
}

@article{li2003angiotensin,
  title={Angiotensin-converting enzyme 2 is a functional receptor for the SARS coronavirus},
  author={Li, Wenhui and Moore, Michael J and Vasilieva, Natalya and Sui, Jianhua and Wong, Swee Kee and Berne, Michael A and Somasundaran, Mohan and Sullivan, John L and Luzuriaga, Katherine and Greenough, Thomas C and others},
  journal={Nature},
  volume={426},
  number={6965},
  pages={450--454},
  year={2003},
  publisher={Nature Publishing Group}
}

@article{hofmann2005human,
  title={Human coronavirus NL63 employs the severe acute respiratory syndrome coronavirus receptor for cellular entry},
  author={Hofmann, Heike and Pyrc, Krzysztof and Van Der Hoek, Lia and Geier, Martina and Berkhout, Ben and P{\"o}hlmann, Stefan},
  journal={Proceedings of the National Academy of Sciences},
  volume={102},
  number={22},
  pages={7988--7993},
  year={2005},
  publisher={National Acad Sciences}
}

@article{hu2018tank,
  title={TANK-binding kinase 1 (TBK1) isoforms negatively regulate type I interferon induction by inhibiting TBK1-IRF3 interaction and IRF3 phosphorylation},
  author={Hu, Yi Wei and Zhang, Jie and Wu, Xiao Man and Cao, Lu and Nie, Pin and Chang, Ming Xian},
  journal={Frontiers in immunology},
  volume={9},
  pages={84},
  year={2018},
  publisher={Frontiers}
}

@article{haagmans2014middle,
  title={Middle East respiratory syndrome coronavirus in dromedary camels: an outbreak investigation},
  author={Haagmans, Bart L and Al Dhahiry, Said HS and Reusken, Chantal BEM and Raj, V Stalin and Galiano, Monica and Myers, Richard and Godeke, Gert-Jan and Jonges, Marcel and Farag, Elmoubasher and Diab, Ayman and others},
  journal={The Lancet infectious diseases},
  volume={14},
  number={2},
  pages={140--145},
  year={2014},
  publisher={Elsevier}
}

@article{kandeil2019middle,
  title={Middle East respiratory syndrome coronavirus infection in non-camelid domestic mammals},
  author={Kandeil, Ahmed and Gomaa, Mokhtar and Shehata, Mahmoud and El-Taweel, Ahmed and Kayed, Ahmed E and Abiadh, Awatef and Jrijer, Jamel and Moatasim, Yassmin and Kutkat, Omnia and Bagato, Ola and others},
  journal={Emerging microbes \& infections},
  volume={8},
  number={1},
  pages={103--108},
  year={2019},
  publisher={Taylor \& Francis}
}

@article{sit2020infection,
  title={Infection of dogs with SARS-CoV-2},
  author={Sit, Thomas HC and Brackman, Christopher J and Ip, Sin Ming and Tam, Karina WS and Law, Pierra YT and To, Esther MW and Yu, Veronica YT and Sims, Leslie D and Tsang, Dominic NC and Chu, Daniel KW and others},
  journal={Nature},
  pages={1--6},
  year={2020},
  publisher={Nature Publishing Group}
}

@article{halfmann2020transmission,
  title={Transmission of SARS-CoV-2 in domestic cats},
  author={Halfmann, Peter J and Hatta, Masato and Chiba, Shiho and Maemura, Tadashi and Fan, Shufang and Takeda, Makoto and Kinoshita, Noriko and Hattori, Shin-ichiro and Sakai-Tagawa, Yuko and Iwatsuki-Horimoto, Kiyoko and others},
  journal={New England Journal of Medicine},
  year={2020},
  publisher={Mass Medical Soc}
}

@article{wang2020complete,
  title={Complete genome sequence of SARS-CoV-2 in a tiger from a US zoological collection},
  author={Wang, Leyi and Mitchell, Patrick K and Calle, Paul P and Bartlett, Susan L and McAloose, Denise and Killian, Mary Lea and Yuan, Fangfeng and Fang, Ying and Goodman, Laura B and Fredrickson, Richard and others},
  journal={Microbiology Resource Announcements},
  volume={9},
  number={22},
  year={2020},
  publisher={American Society for Microbiology (ASM)}
}

@article{sia2020pathogenesis,
  title={Pathogenesis and transmission of SARS-CoV-2 in golden hamsters},
  author={Sia, Sin Fun and Yan, Li-Meng and Chin, Alex WH and Fung, Kevin and Choy, Ka-Tim and Wong, Alvina YL and Kaewpreedee, Prathanporn and Perera, Ranawaka APM and Poon, Leo LM and Nicholls, John M and others},
  journal={Nature},
  volume={583},
  number={7818},
  pages={834--838},
  year={2020},
  publisher={Nature Publishing Group}
}

@article{li2020early,
  title={Early transmission dynamics in Wuhan, China, of novel coronavirus--infected pneumonia},
  author={Li, Qun and Guan, Xuhua and Wu, Peng and Wang, Xiaoye and Zhou, Lei and Tong, Yeqing and Ren, Ruiqi and Leung, Kathy SM and Lau, Eric HY and Wong, Jessica Y and others},
  journal={New England Journal of Medicine},
  year={2020},
  publisher={Mass Medical Soc}
}

@article{world2020coronavirus,
  title={Coronavirus disease (COVID-19) situation dashboard},
  author={World Health Organization and others},
  journal={World Health Organization website. https://who. sprinklr. com/. Accessed April},
  volume={17},
  year={2020}
}

@article{channappanavar2016dysregulated,
  title={Dysregulated type I interferon and inflammatory monocyte-macrophage responses cause lethal pneumonia in SARS-CoV-infected mice},
  author={Channappanavar, Rudragouda and Fehr, Anthony R and Vijay, Rahul and Mack, Matthias and Zhao, Jincun and Meyerholz, David K and Perlman, Stanley},
  journal={Cell host \& microbe},
  volume={19},
  number={2},
  pages={181--193},
  year={2016},
  publisher={Elsevier}
}

@inproceedings{Wang:2016:SDN:2939672.2939753,
 author = {Wang, Daixin and Cui, Peng and Zhu, Wenwu},
 title = {Structural Deep Network Embedding},
 booktitle = {Proceedings of the 22Nd ACM SIGKDD International Conference on Knowledge Discovery and Data Mining},
 series = {KDD '16},
 year = {2016},
 isbn = {978-1-4503-4232-2},
 location = {San Francisco, California, USA},
 pages = {1225--1234},
 numpages = {10},
 url = {http://doi.acm.org/10.1145/2939672.2939753},
 doi = {10.1145/2939672.2939753},
 acmid = {2939753},
 publisher = {ACM},
 address = {New York, NY, USA},
 keywords = {deep learning, network analysis, network embedding},
}

@inproceedings{ahmed2013distributed,
  title                    = {Distributed large-scale natural graph factorization},
  author                   = {Ahmed, Amr and Shervashidze, Nino and Narayanamurthy, Shravan and Josifovski, Vanja and Smola, Alexander J},
  booktitle                = {Proceedings of the 22nd international conference on World Wide Web},
  pages                    = {37--48},
  year                     = {2013},
  organization             = {ACM}
}


@InProceedings{SciPyProceedings_11,
  author =       {Aric A. Hagberg and Daniel A. Schult and Pieter J. Swart},
  title =        {Exploring Network Structure, Dynamics, and Function using NetworkX},
  booktitle =   {Proceedings of the 7th Python in Science Conference},
  pages =     {11 - 15},
  address = {Pasadena, CA USA},
  year =      {2008},
  editor =    {Ga\"el Varoquaux and Travis Vaught and Jarrod Millman},
}

@article{schulz2016viral,
  title={Viral evasion strategies in type I IFN signaling--a summary of recent developments},
  author={Schulz, Katharina S and Mossman, Karen L},
  journal={Frontiers in immunology},
  volume={7},
  pages={498},
  year={2016},
  publisher={Frontiers}
}

@InProceedings {7474366,
    author = {A. Heidarian and M. J. Dinneen},
    booktitle = {2016 IEEE Second International Conference on Big Data Computing Service and Applications (BigDataService)},
    title = {A Hybrid Geometric Approach for Measuring Similarity Level Among Documents and Document Clustering},
    year = {2016},
    volume = {},
    issn = {},
    pages = {142-151},
    keywords = {euclidean distance;semantics;computational modeling;current measurement;frequency measurement;extraterrestrial measurements;correlation},
    doi = {10.1109/BigDataService.2016.14},
    url = {https://doi.ieeecomputersociety.org/10.1109/BigDataService.2016.14},
    publisher = {IEEE Computer Society},
    address = {Los Alamitos, CA, USA},
    month = {apr}
}

@inproceedings{sklearn_api,
  author    = {Lars Buitinck and Gilles Louppe and Mathieu Blondel and
               Fabian Pedregosa and Andreas Mueller and Olivier Grisel and
               Vlad Niculae and Peter Prettenhofer and Alexandre Gramfort
               and Jaques Grobler and Robert Layton and Jake VanderPlas and
               Arnaud Joly and Brian Holt and Ga{\"{e}}l Varoquaux},
  title     = {{API} design for machine learning software: experiences from the scikit-learn
               project},
  booktitle = {ECML PKDD Workshop: Languages for Data Mining and Machine Learning},
  year      = {2013},
  pages = {108--122},
}

@article{frieman2009severe,
  title={Severe acute respiratory syndrome coronavirus papain-like protease ubiquitin-like domain and catalytic domain regulate antagonism of IRF3 and NF-$\kappa$B signaling},
  author={Frieman, Matthew and Ratia, Kiira and Johnston, Robert E and Mesecar, Andrew D and Baric, Ralph S},
  journal={Journal of virology},
  volume={83},
  number={13},
  pages={6689--6705},
  year={2009},
  publisher={Am Soc Microbiol}
}

@article{mielech2014mers,
  title={MERS-CoV papain-like protease has deISGylating and deubiquitinating activities},
  author={Mielech, Anna M and Kilianski, Andy and Baez-Santos, Yahira M and Mesecar, Andrew D and Baker, Susan C},
  journal={Virology},
  volume={450},
  pages={64--70},
  year={2014},
  publisher={Elsevier}
}

@article {PMID:27344959,
	Title = {SARS and MERS: recent insights into emerging coronaviruses},
	Author = {de Wit, Emmie and van Doremalen, Neeltje and Falzarano, Darryl and Munster, Vincent J},
	DOI = {10.1038/nrmicro.2016.81},
	Number = {8},
	Volume = {14},
	Month = {August},
	Year = {2016},
	Journal = {Nature reviews. Microbiology},
	ISSN = {1740-1526},
	Pages = {523—534},
	URL = {https://europepmc.org/articles/PMC7097822},
}

@incollection{kilpatrick2004cytokines,
  title={Cytokines and inflammatory response in the fetus and neonate},
  author={Kilpatrick, Laurie and Harris, Mary Catherine},
  booktitle={Fetal and neonatal physiology},
  pages={1555--1572},
  year={2004},
  publisher={Elsevier}
}

@article{liu2017nf,
  title={NF-$\kappa$B signaling in inflammation},
  author={Liu, Ting and Zhang, Lingyun and Joo, Donghyun and Sun, Shao-Cong},
  journal={Signal transduction and targeted therapy},
  volume={2},
  number={1},
  pages={1--9},
  year={2017},
  publisher={Nature Publishing Group}
}

@article{michalska2018positive,
  title={A positive feedback amplifier circuit that regulates interferon (IFN)-stimulated gene expression and controls type I and type II IFN responses},
  author={Michalska, Agata and Blaszczyk, Katarzyna and Wesoly, Joanna and Bluyssen, Hans AR},
  journal={Frontiers in immunology},
  volume={9},
  pages={1135},
  year={2018},
  publisher={Frontiers}
}

@article{lei2009mavs,
  title={MAVS-mediated apoptosis and its inhibition by viral proteins},
  author={Lei, Yu and Moore, Chris B and Liesman, Rachael M and O'Connor, Brian P and Bergstralh, Daniel T and Chen, Zhijian J and Pickles, Raymond J and Ting, Jenny P-Y},
  journal={PloS one},
  volume={4},
  number={5},
  pages={e5466},
  year={2009},
  publisher={Public Library of Science}
}

@article{wu2015rig,
  title={How RIG-I like receptors activate MAVS},
  author={Wu, Bin and Hur, Sun},
  journal={Current opinion in virology},
  volume={12},
  pages={91--98},
  year={2015},
  publisher={Elsevier}
}

@article{zhai2020comparison,
  title={Comparison of severe acute respiratory syndrome coronavirus 2 spike protein binding to ACE2 receptors from human, pets, farm animals, and putative intermediate hosts},
  author={Zhai, Xiaofeng and Sun, Jiumeng and Yan, Ziqing and Zhang, Jie and Zhao, Jin and Zhao, Zongzheng and Gao, Qi and He, Wan-Ting and Veit, Michael and Su, Shuo},
  journal={Journal of virology},
  volume={94},
  number={15},
  year={2020},
  publisher={Am Soc Microbiol}
}
