## Supplementary material for "Network-based Virus-Host Interaction Prediction with Application to SARS-CoV-2": Latex Package: model.pptx

### Slide 1
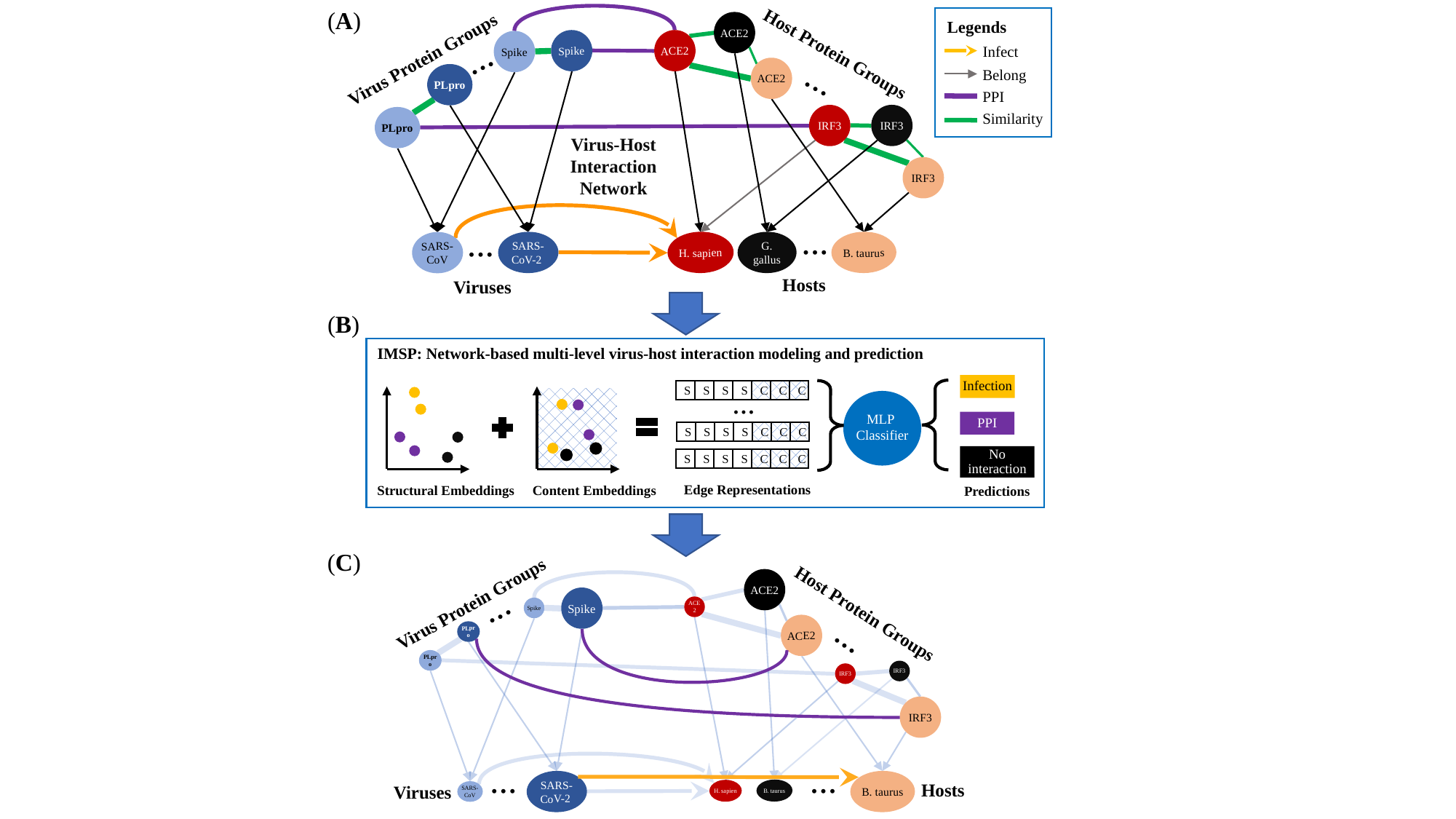

(A)
Legends
Infect
Belong
PPI
Similarity
ACE2
...
Spike
ACE2
Spike
Virus Protein Groups
Host Protein Groups
...
ACE2
PLpro
IRF3
IRF3
PLpro
ACE2
Virus-Host Interaction Network
IRF3
...
...
H. sapien
B. taurus
SARS-CoV-2
G. gallus
SARS-CoV
Hosts
Viruses
(B)
IMSP: Network-based multi-level virus-host interaction modeling and prediction
Infection
C
C
C
S
S
S
S
...
MLP
Classifier
PPI
C
C
C
S
S
S
S
No interaction
C
C
C
S
S
S
S
Edge Representations
Structural Embeddings
Content Embeddings
Predictions
(C)
ACE2
...
Virus Protein Groups
Spike
ACE2
Spike
Host Protein Groups
...
ACE2
PLpro
PLpro
IRF3
IRF3
ACE2
IRF3
...
...
B. taurus
SARS-CoV-2
Hosts
Viruses
B. taurus
H. sapien
SARS-CoV

### Slide 2
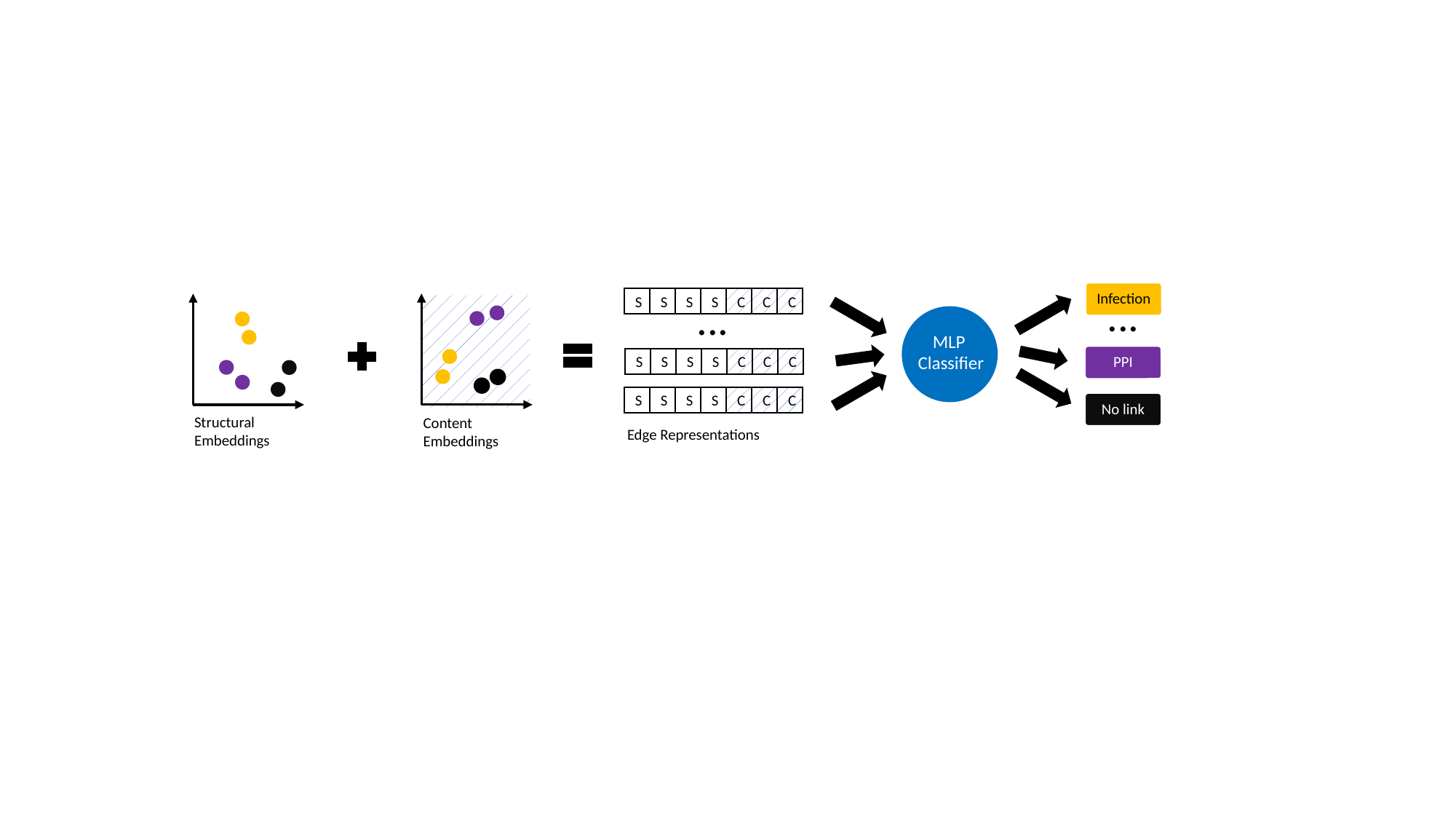

Infection
...
...
C
C
C
S
S
S
S
MLP
Classifier
PPI
C
C
C
S
S
S
S
C
C
C
S
S
S
S
No link
Structural Embeddings
Content Embeddings
Edge Representations

### Slide 3
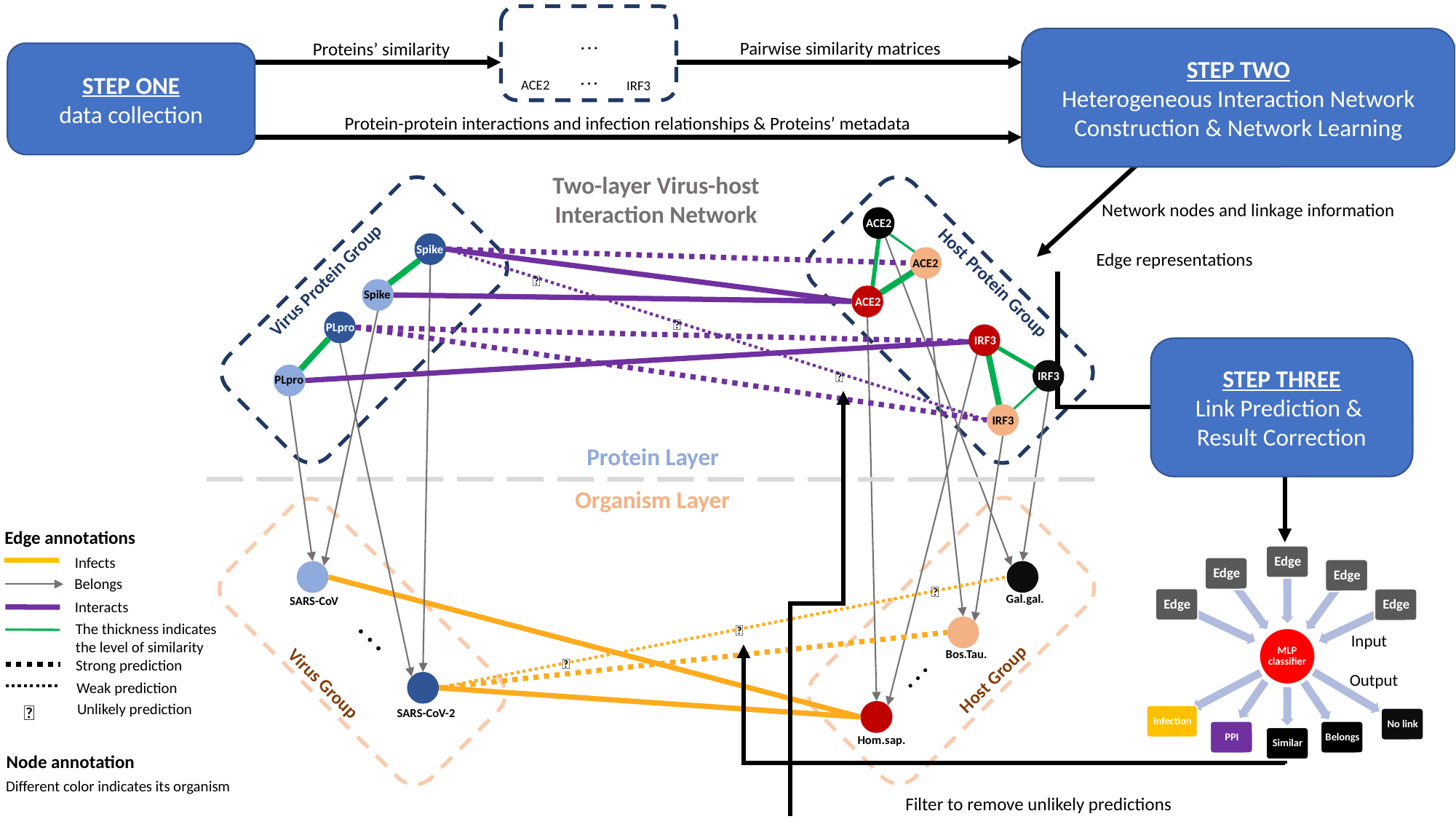

ACE2
IRF3
STEP TWO
Heterogeneous Interaction Network Construction & Network Learning
Pairwise similarity matrices
Proteins’ similarity
STEP ONE
data collection
Protein-protein interactions and infection relationships & Proteins’ metadata
Two-layer Virus-host Interaction Network
Network nodes and linkage information
ACE2
Spike
Edge representations
ACE2
Virus Protein Group
❌
Host Protein Group
Spike
ACE2
❌
PLpro
IRF3
STEP THREE
Link Prediction &
Result Correction
❌
IRF3
PLpro
IRF3
Protein Layer
Organism Layer
Edge annotations
Infects
Belongs
 ❌
Gal.gal.
SARS-CoV
Interacts
The thickness indicates the level of similarity
 ❌
Input
Bos.Tau.
 ❌
Strong prediction
Host Group
Output
Virus Group
Weak prediction
❌
Unlikely prediction
SARS-CoV-2
Hom.sap.
Node annotation
Different color indicates its organism
Filter to remove unlikely predictions

### Slide 4
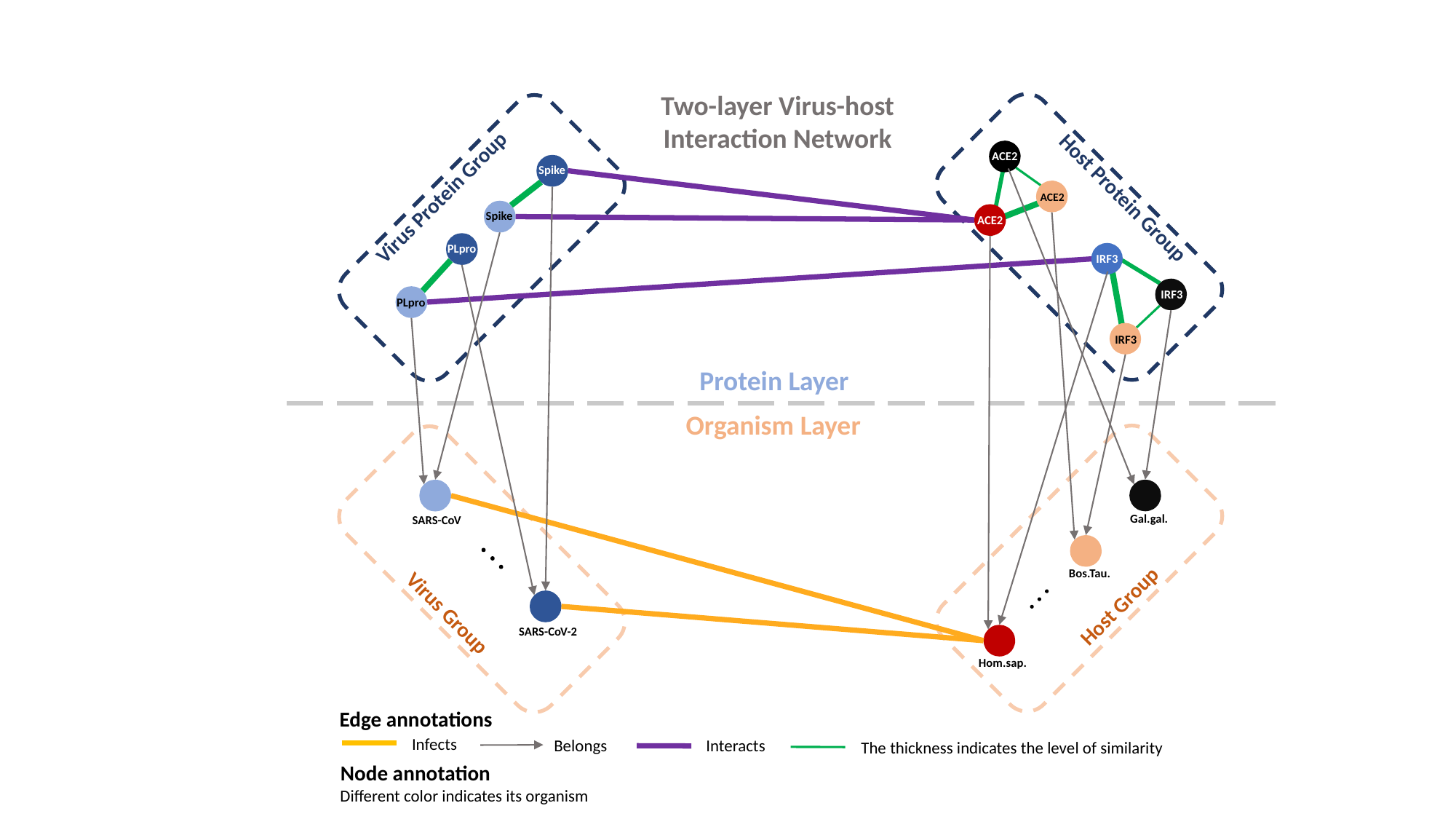

Two-layer Virus-host Interaction Network
ACE2
Spike
Virus Protein Group
Host Protein Group
ACE2
Spike
ACE2
PLpro
IRF3
IRF3
PLpro
IRF3
Protein Layer
Organism Layer
Gal.gal.
SARS-CoV
Bos.Tau.
Host Group
Virus Group
SARS-CoV-2
Hom.sap.
Edge annotations
Infects
Belongs
Interacts
The thickness indicates the level of similarity
Node annotation
Different color indicates its organism

### Slide 5
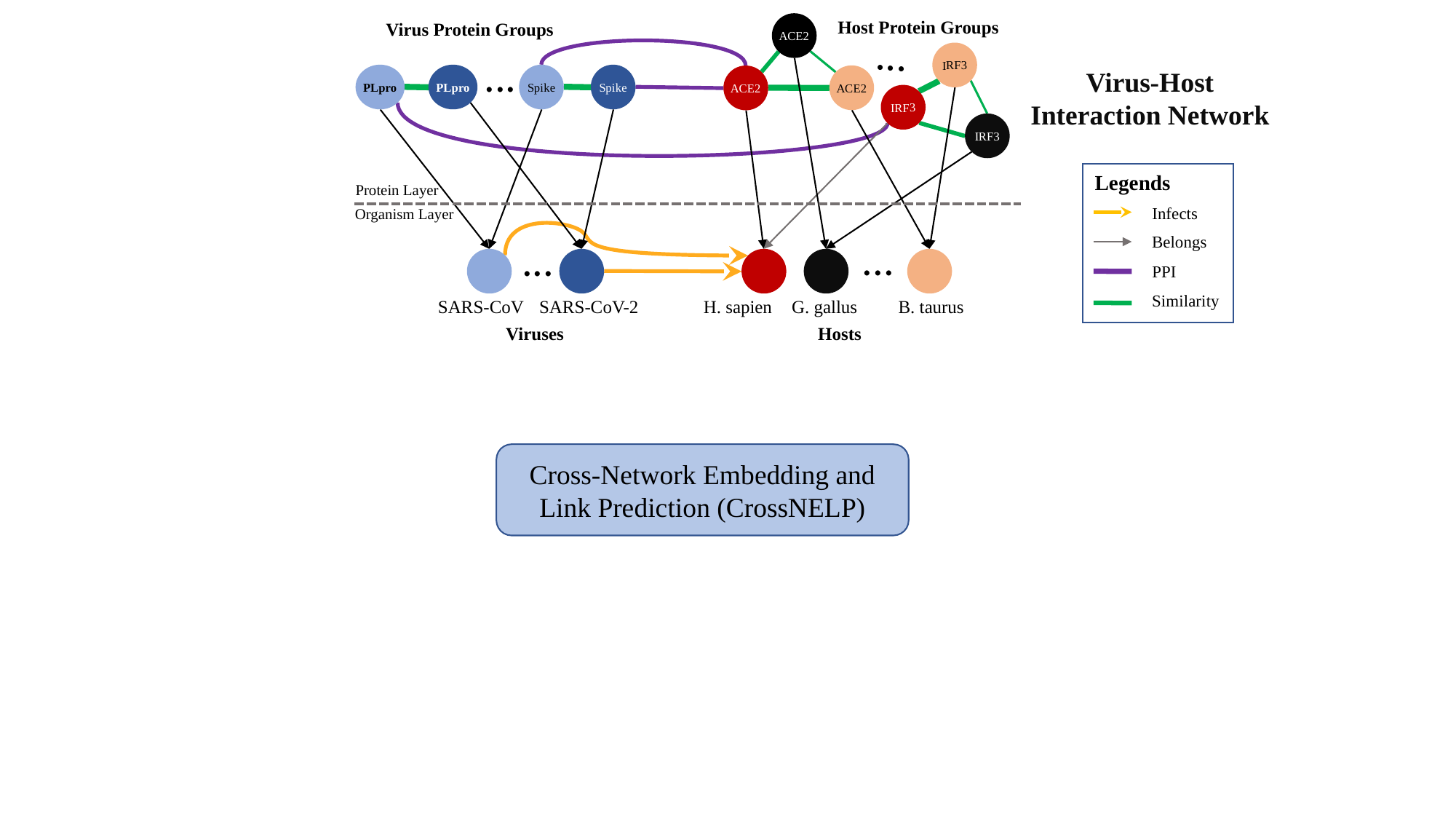

Host Protein Groups
Virus Protein Groups
ACE2
...
IRF3
...
Virus-Host Interaction Network
PLpro
PLpro
Spike
Spike
ACE2
ACE2
IRF3
ACE2
IRF3
Legends
Infects
Belongs
PPI
Similarity
Protein Layer
Organism Layer
...
...
B. taurus
SARS-CoV
SARS-CoV-2
H. sapien
G. gallus
Viruses
Hosts
Cross-Network Embedding and Link Prediction (CrossNELP)

### Slide 6
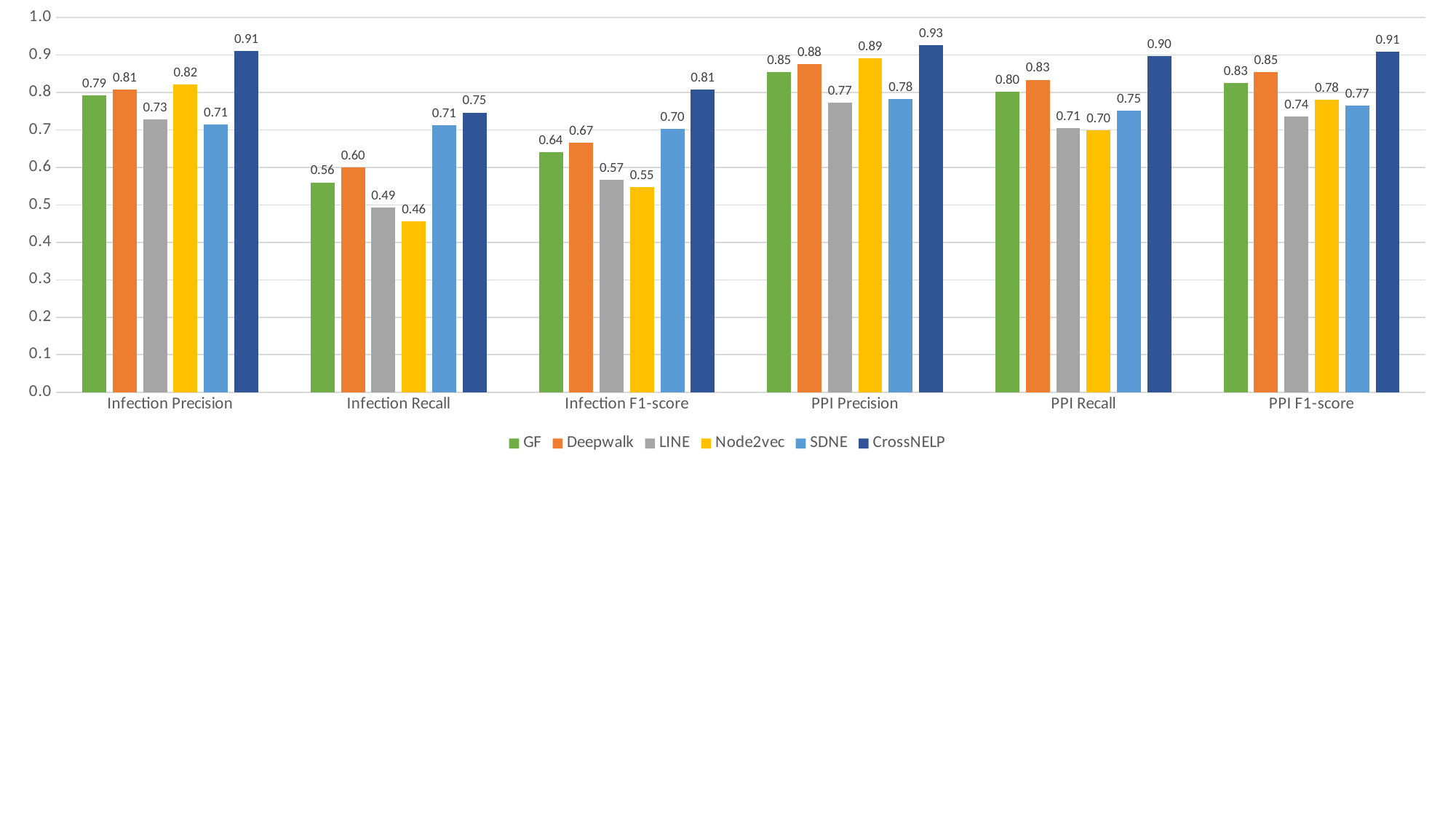

#### Chart
| Category | GF | Deepwalk | LINE | Node2vec | SDNE | CrossNELP |
|---|---|---|---|---|---|---|
| Infection Precision | 0.792 | 0.808 | 0.728649591149591 | 0.821719576719576 | 0.714156352906352 | 0.910882635882635 |
| Infection Recall | 0.56 | 0.6 | 0.493333333333333 | 0.456666666666666 | 0.713333333333333 | 0.746666666666666 |
| Infection F1-score | 0.64 | 0.667 | 0.567178745174101 | 0.547058130208285 | 0.703293827327008 | 0.808096682585846 |
| PPI Precision | 0.854 | 0.876 | 0.773381265650009 | 0.891598111339254 | 0.782020768104096 | 0.926058993489029 |
| PPI Recall | 0.802 | 0.834 | 0.705434782608695 | 0.698188405797101 | 0.751086956521739 | 0.896376811594202 |
| PPI F1-score | 0.826 | 0.854 | 0.736472441960616 | 0.781203312512355 | 0.765055555848165 | 0.909784301057553 |
