## Supplementary figures and images for "Network-based Virus-Host Interaction Prediction with Application to SARS-CoV-2"

### Cartoon infection final.jpg

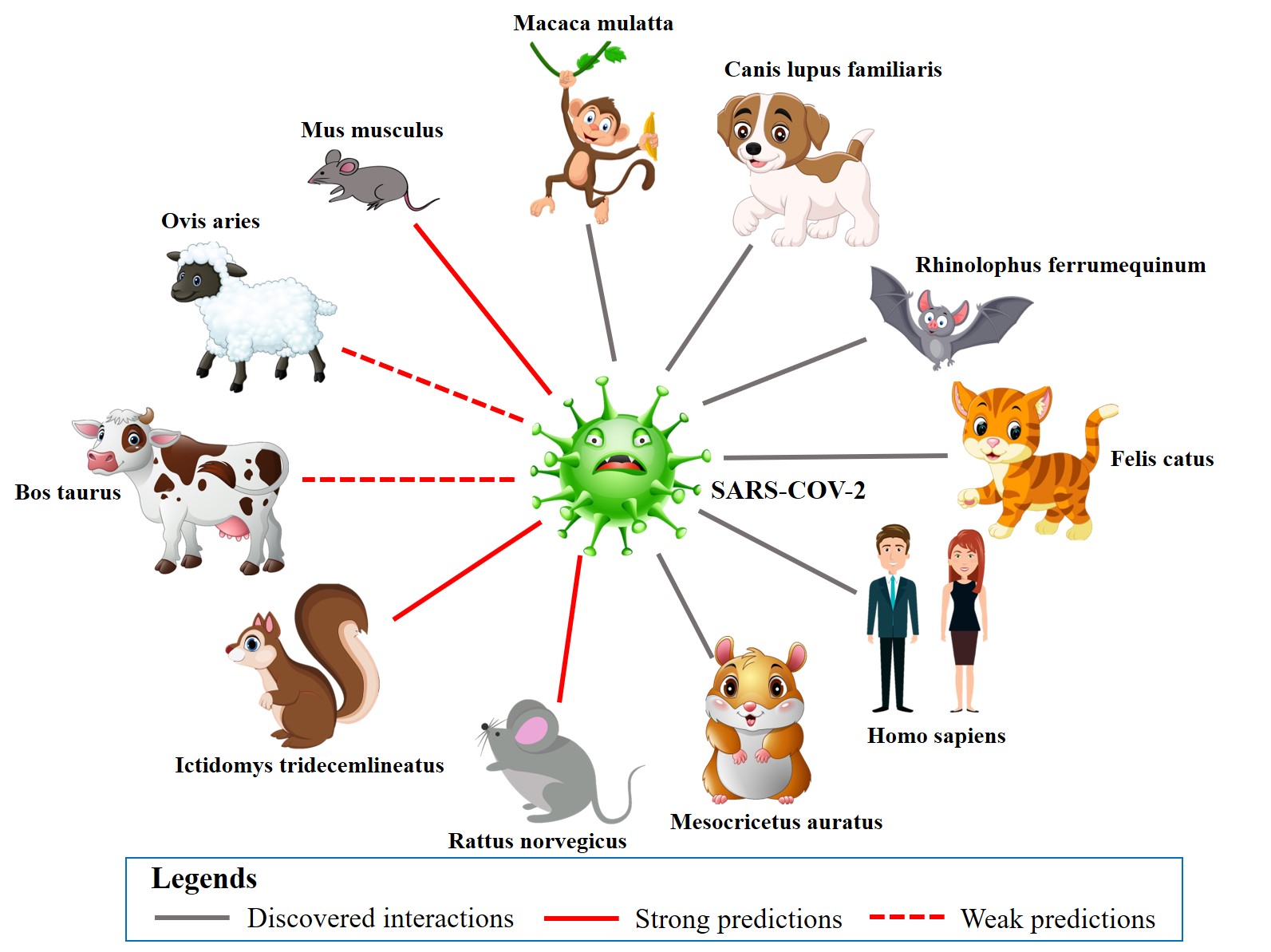

### IFN pathway graph 10.12.jpg

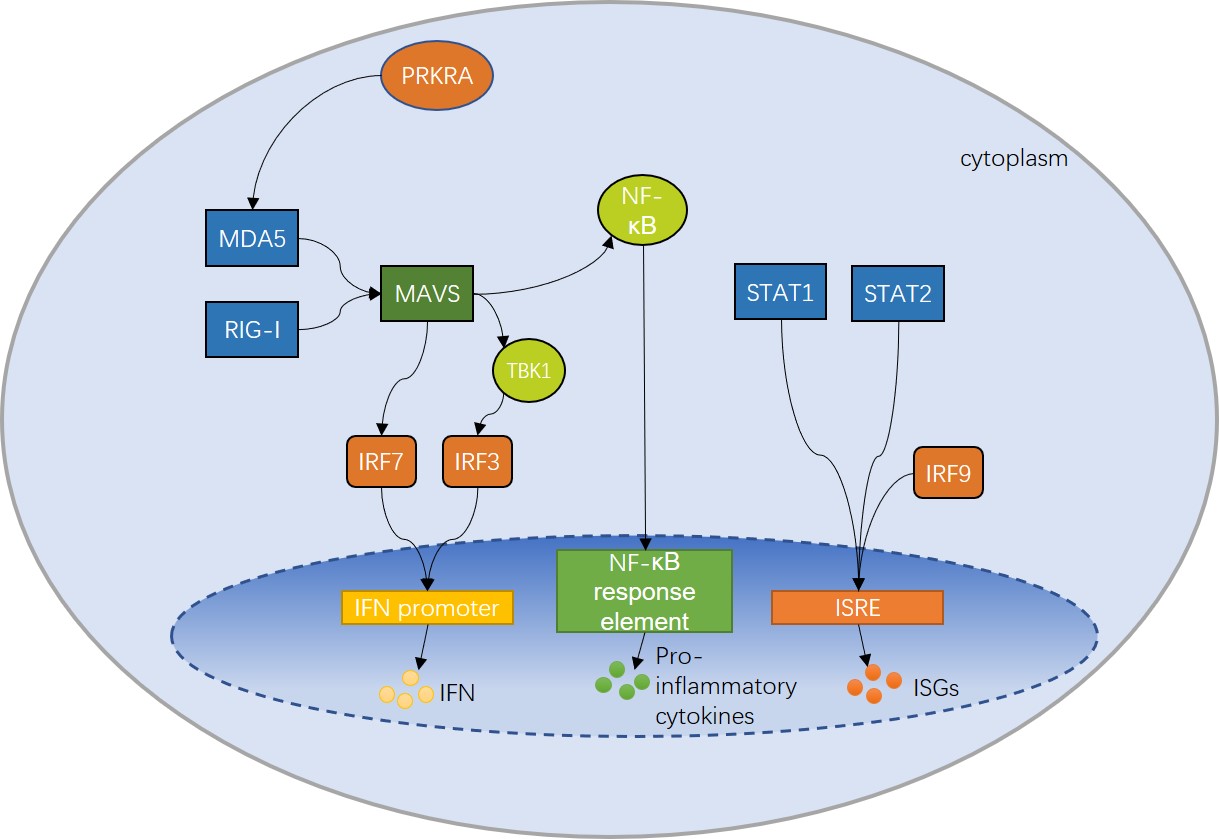

### IFN_pathway 10.24.jpeg

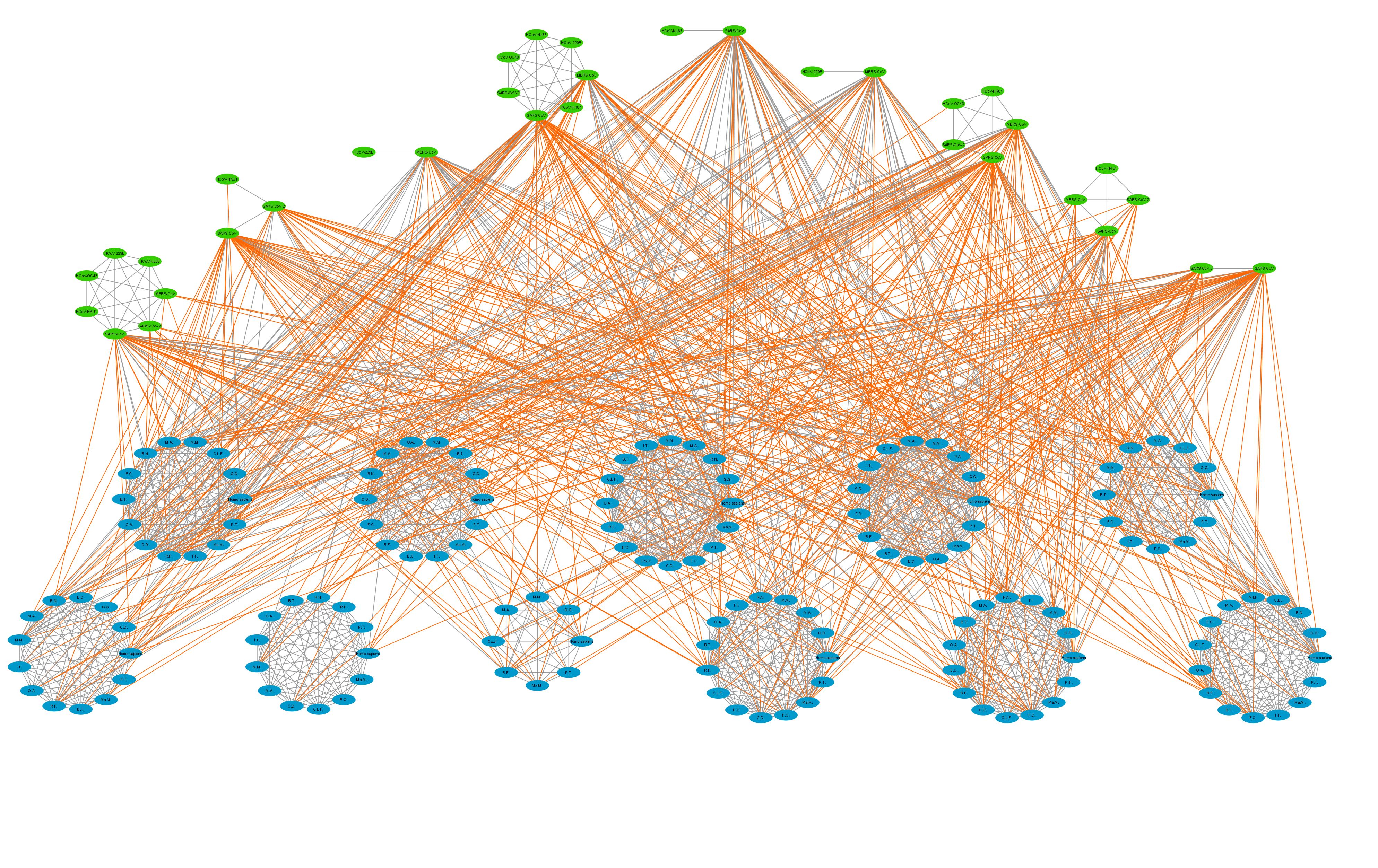

### infection_final 10.24.jpeg

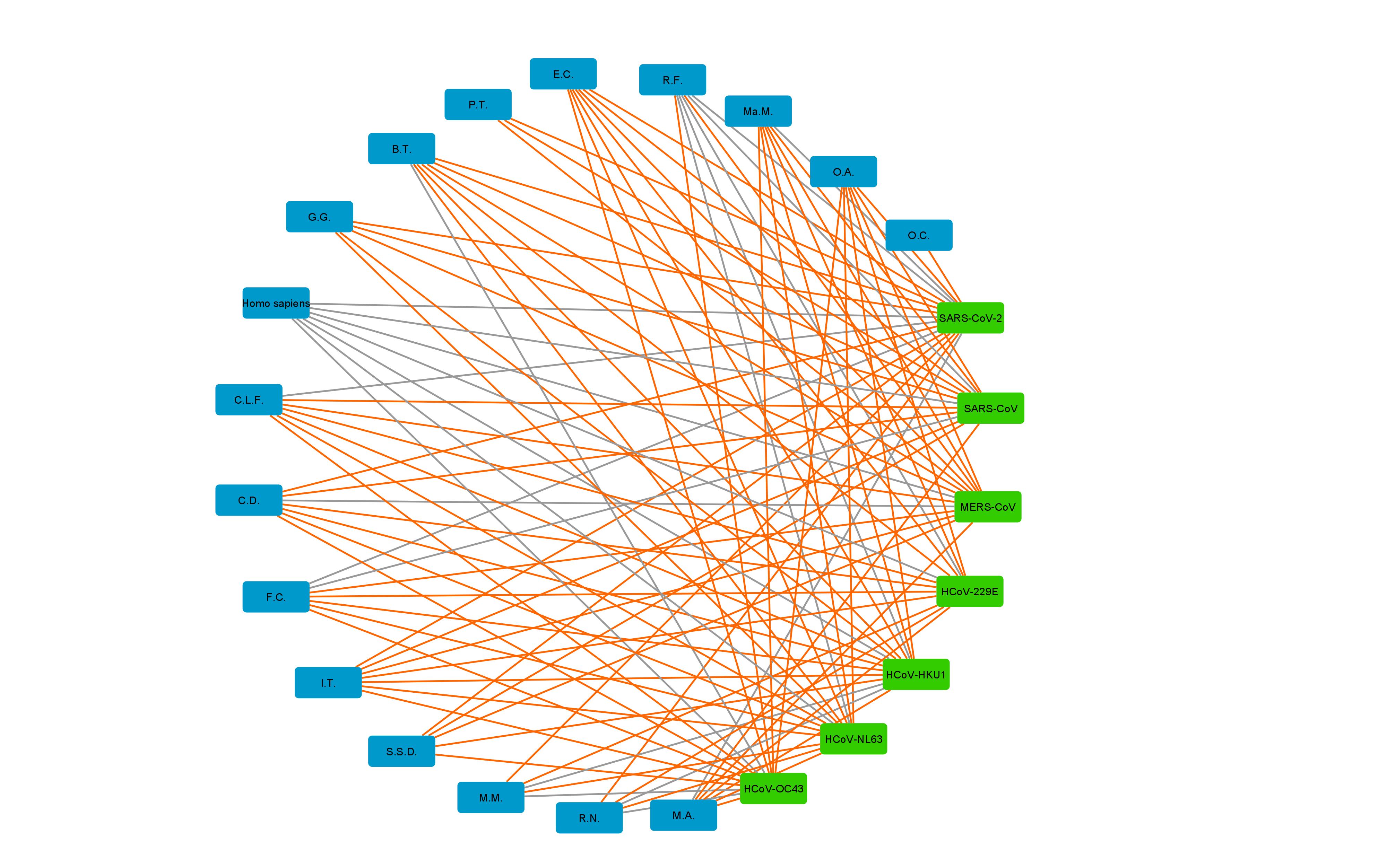

### model.png

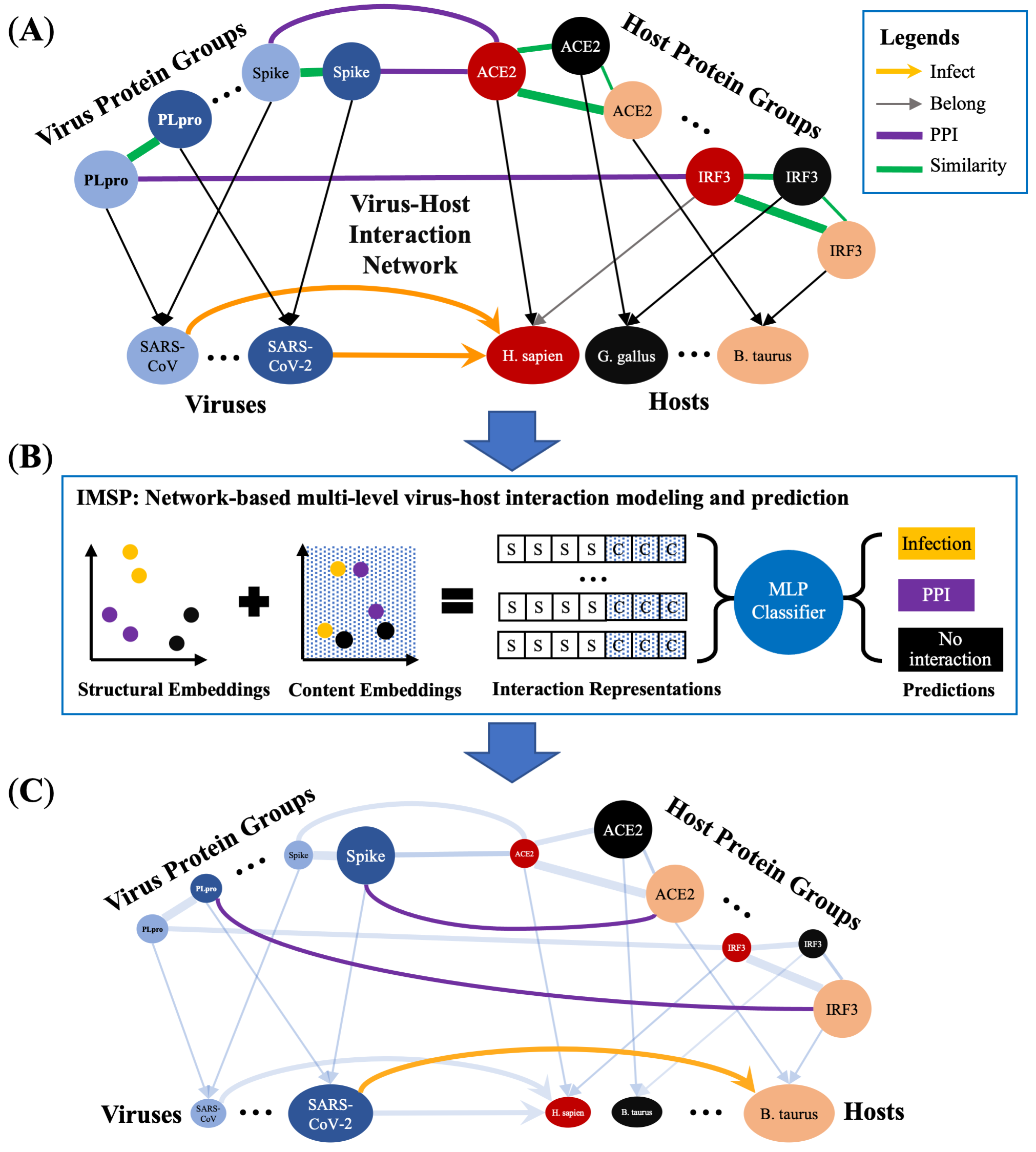

### perf.jpg

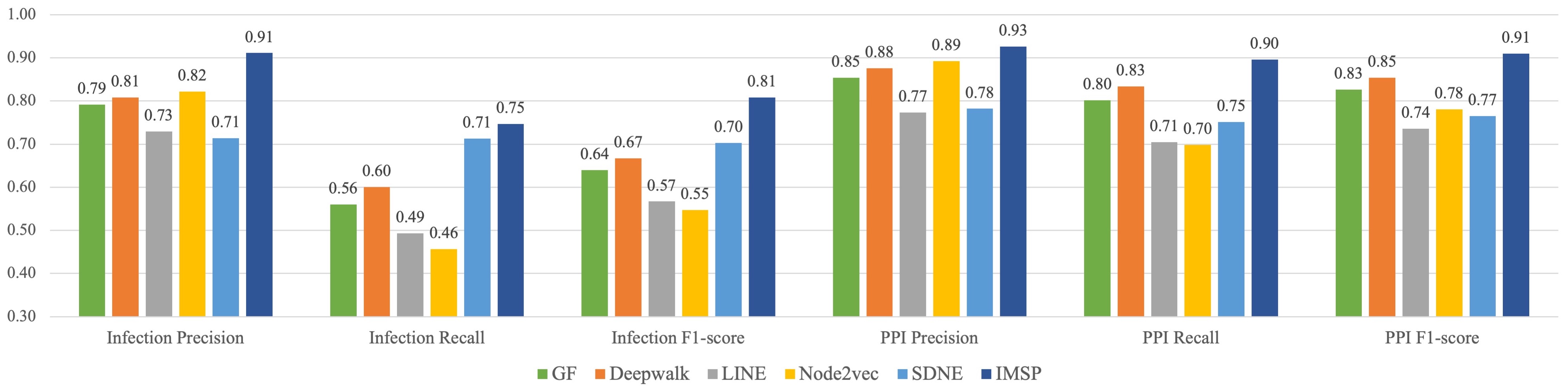

### protein prediction SARS-CoV-2.jpg

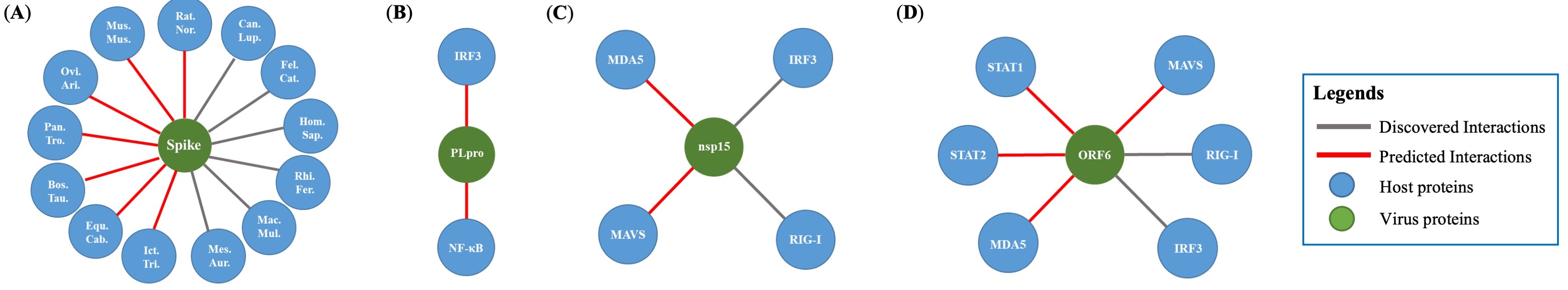

### spike_binding 10.24.jpeg

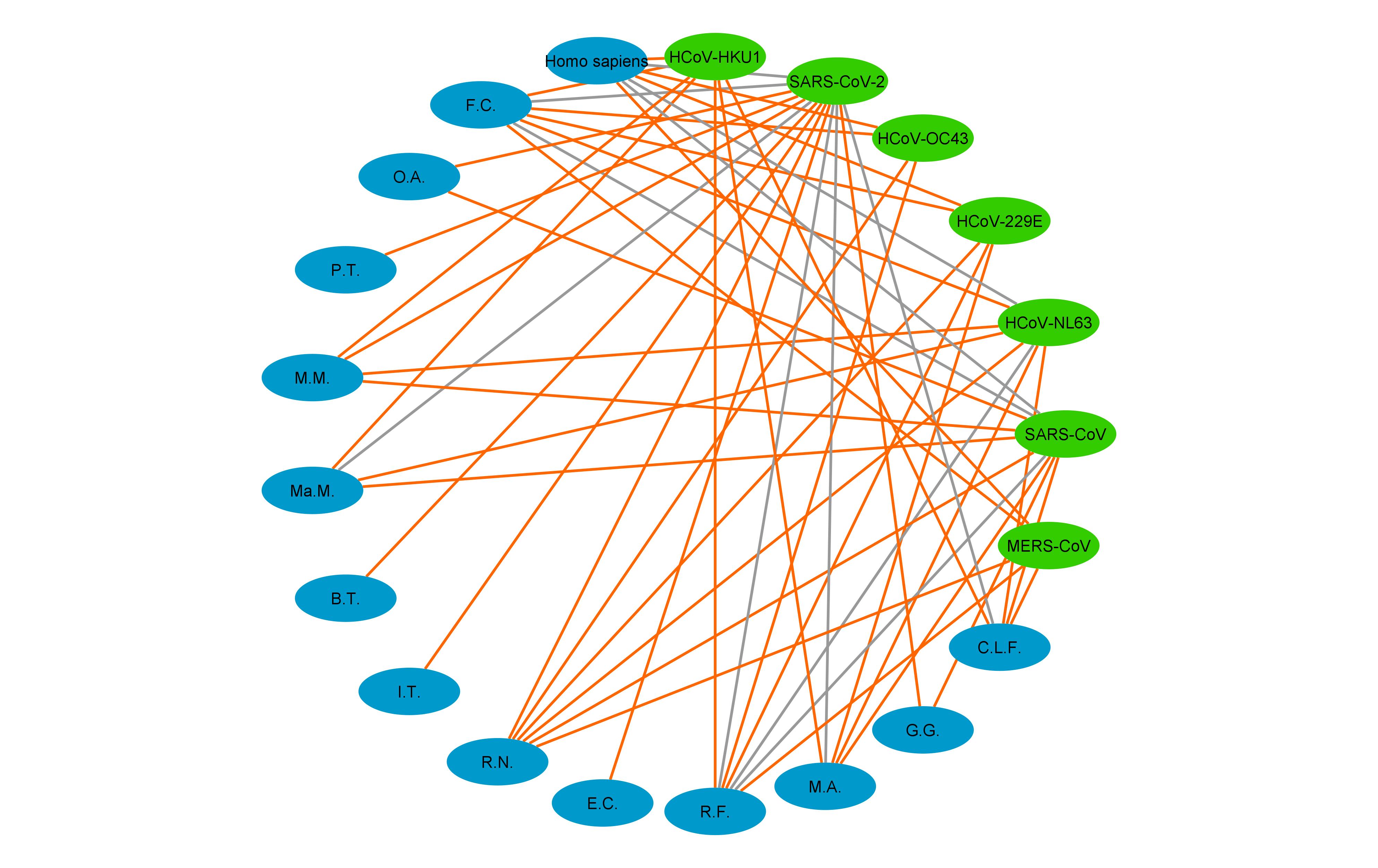

### virus entry graph 10.9.jpg

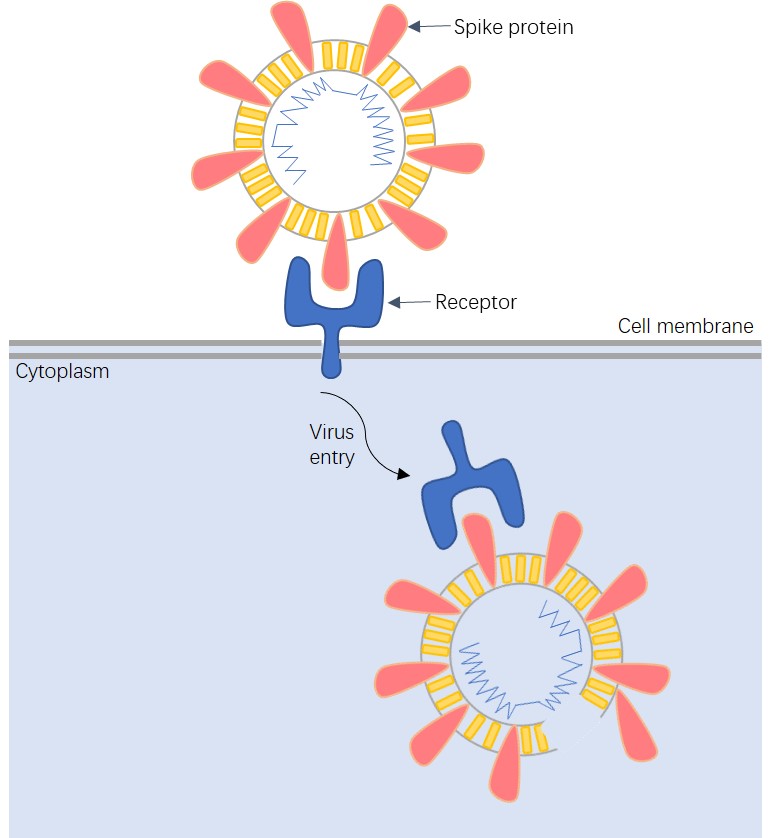
