## Supplementary Information for "Network-based Virus-Host Interaction Prediction with Application to SARS-CoV-2"

The supplementary material contained visualized demonstrations of viral entry and IFN pathway mechanisms. These two parts together contribute to a successful virus invasion. The full network with predictions made by the model was visualized in three figures: Figure S3 for viral entry, Figure S4 for IFN pathway and Figure S5 for host infection. The full nodes and interactions in the network are presented in Table S6 and Table S7. The predicted interactions are presented in TableS8 and Table S9.

### Supplementary Note 1: Virus Entry - receptor binding of S protein

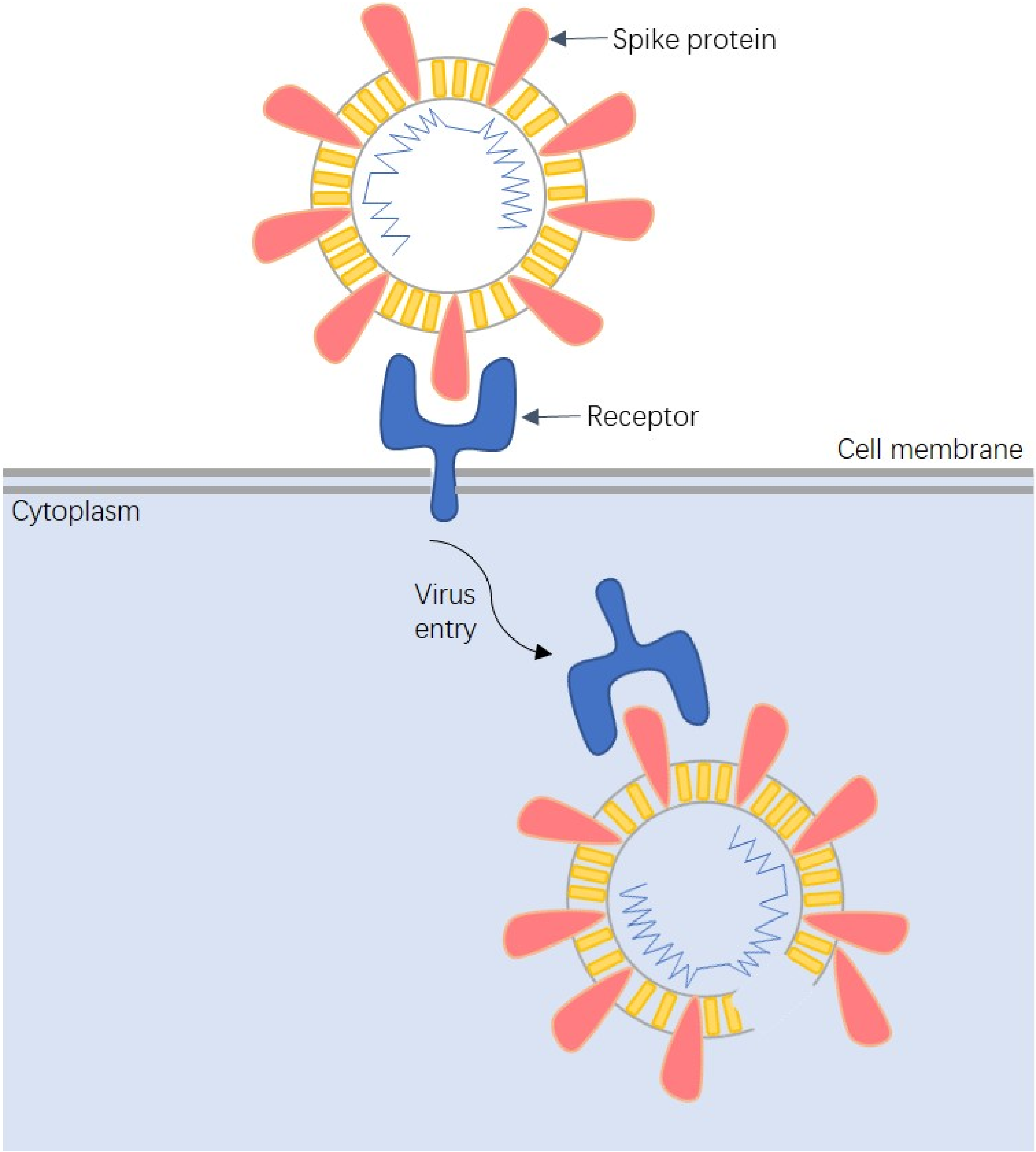


**Fig. 1. The Process for Coronavirus Receptor Binding and Virus Entry.** *The S protein in coronaviruses plays a crucial role in viral entry. It binds with host receptors and facilitates the fusion between the viral envelope and the host cell membrane.*

### Supplementary Note 2: Immune Response - IFN Signaling Pathway

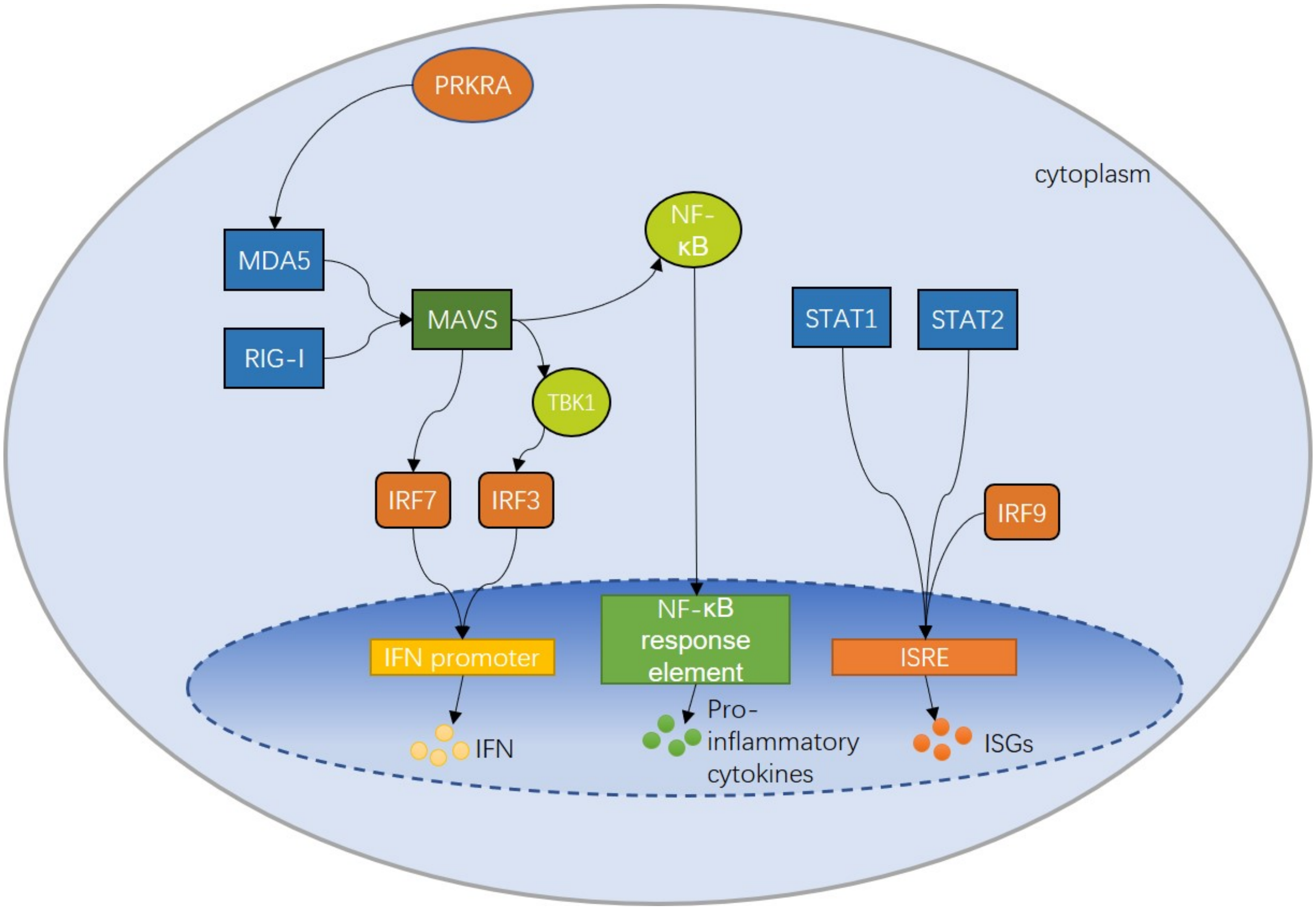


**Fig. 2. Innate Immune Response to Coronaviruses' Viral Infection and IFN signaling Mechanism.** *RIG-I and MDA5 detect the pattern of virus and trigger the production of Interferons (IFNs) (? ) and the activation of the NF-κB (? ). The activated NF-κB induces the Pro-inflammatory cytokines(? ), which play a central role in inflammatory diseases of infectious (? ). STAT1 and STAT2 associate with IRF9 to induce the expression of interferon-stimulated genes (ISGs) (? ) and produce antiviral proteins (? ). In this way, viral interactions with the host innate immune system to suppress immune responses become the critical determinant of the disease outcome and viral infection.*

### Supplementary Note 3: Full Map of Binding Interactions between Coronaviruses' S Protein and Mammalian Hosts' Receptor

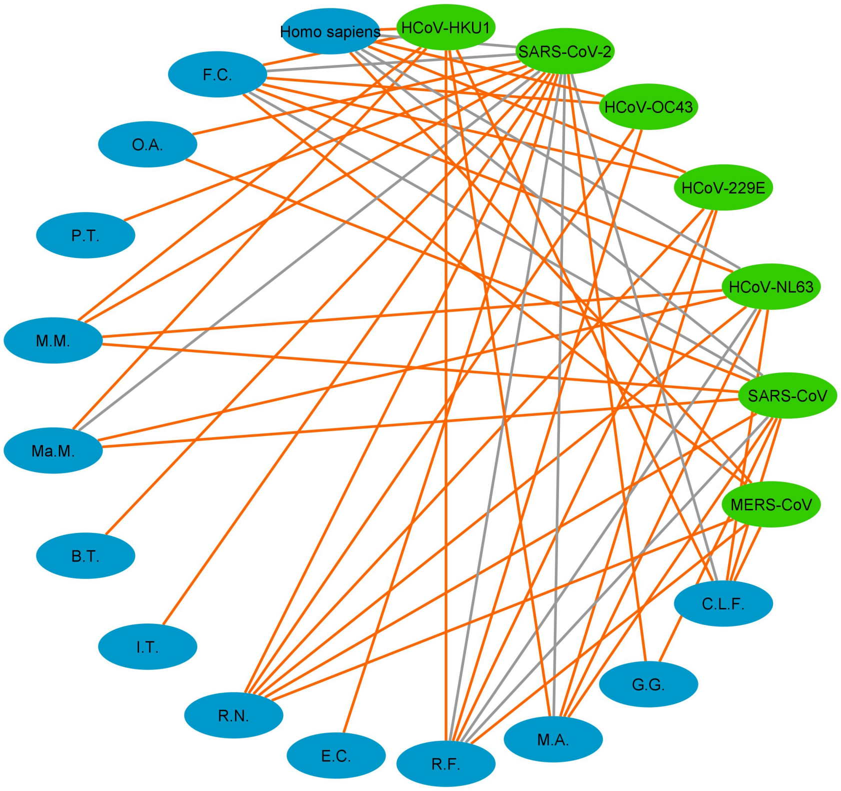


**Fig. 3. Virus entry: binding relationships between the S-proteins of human coronaviruses and the ACE2 receptors of mammalian hosts.** *This figure of the network is visualized by Cytoscape. Virus Protein Layer nodes are represented in green ellipses, and Host Protein Layer nodes are represented in blue ellipses. The original interactions are represented in light grey lines, including the known receptor bindings between viruses spike and mammalian hosts ACE2. Predicted receptor bindings are represented in black lines. As the infection relations have been checked for likelihood in the IMSP model, all predicted interactions are strong predicted interactions. All host names, except Homo sapiens, are displayed in their abbreviation form: M.M. is Mus musculus; F.C. is Felis catus; C.L.F. is Canis lupus familiaris; O.A. is Ovis aries; R.N. is Rattus norvegicus; Ma.M. is Macaca mulatta; R.F. is Rhinolophus ferrumequinum; M.A. is Mesocricetus auratus; C.D. is Camelus dromedarius; B.T. is Bos taurus; G.G. is Gallus gallus; I.T. is Ictidomys tridecemlineatus; E.C. is Equus caballus; S.S.D. is Sus scrofa domesticus; P.T. is Pan troglodytes; O.C. is Oryctolagus cuniculus.*

### Supplementary Note 4: Full Map of Protein-Protein Interactions in IFN Signaling Pathway between Coronaviruses' Proteins and Mammalian Hosts' Proteins

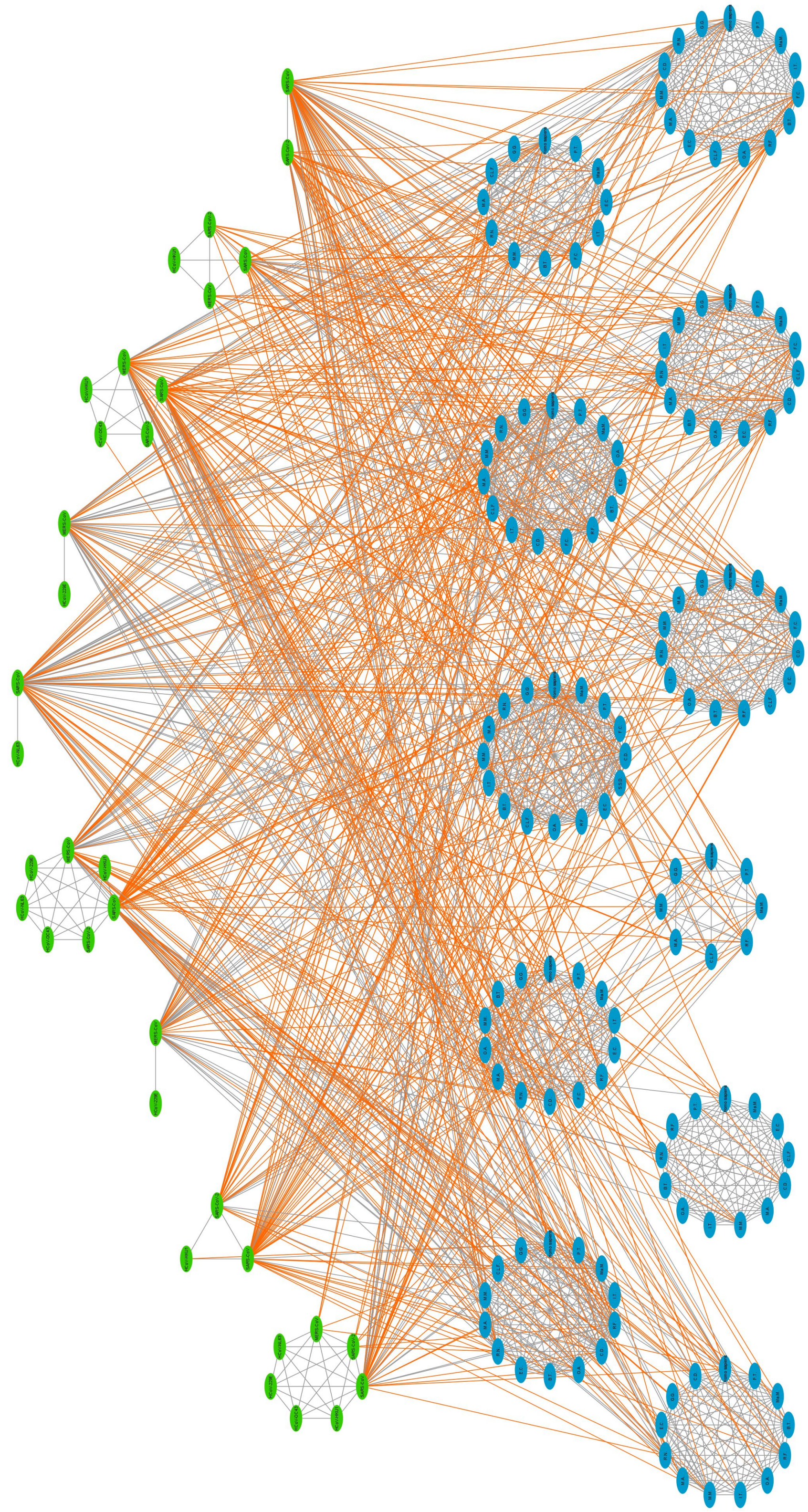

**Fig. 4. IFN interactions: virus proteins interactions with IFN signaling pathway to suppress the IFN signaling.** *Same representations for nodes and interactions as described in figure3. The predicted interactions are represented in black lines: the solid lines stand for strong predictions, and dotted lines stand for weak predictions as defined in the IMSP model.*

### Supplementary Note 5: Full Map of Infection Relationships between Coronaviruses and Mammalian Hosts

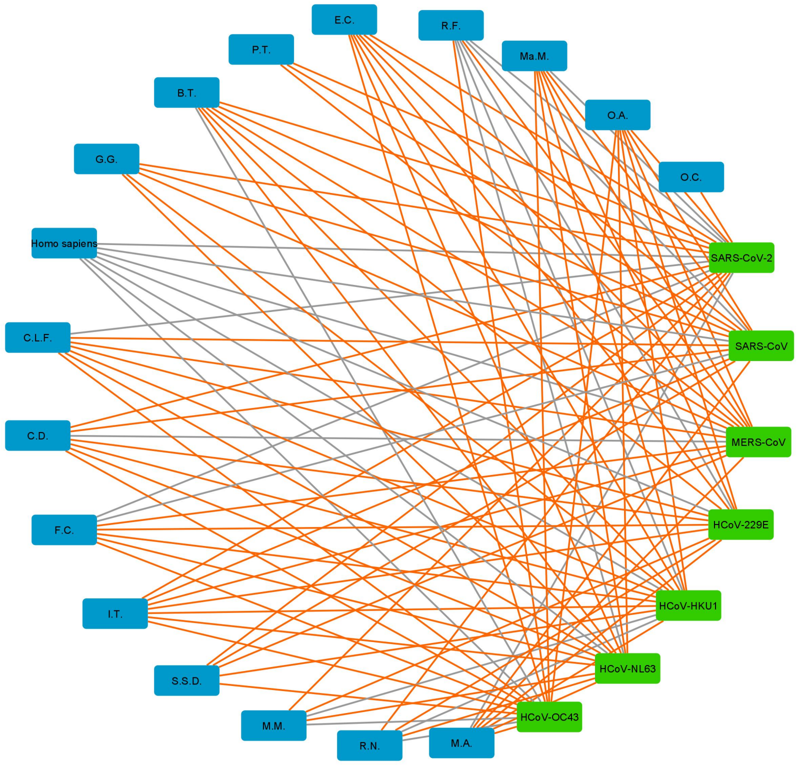


**Fig. 5. Coronaviruses and mammalian hosts infection relationships** *Virus Layer nodes are represented in green rhombi, and Host Layer nodes are represented in blue rhombi. The original infection interactions are represented in grey lines. Predicted infection relations are represented in black lines: the solid lines stand for strong predictions and dotted lines stand for weak predictions as defined in the IMSP model.*

### Supplementary Note 6: Network Node IDs associated with Node Full Names

| Node ID | Full name representation | Node ID | Full name representation |
|---|---|---|---|
| 0 | nsp15, Severe acute respiratory syndrome coronavirus 2 | 123 | IRF7, Rattus norvegicus |
| 1 | nsp15, Severe acute respiratory syndrome-related coronavirus | 124 | IRF7, Mesocricetus auratus |
| 2 | nsp15, Human coronavirus HKU1 | 125 | IRF7, Ovis aries |
| 3 | STAT1, Homo sapiens | 126 | IRF7, Mus musculus |
| 4 | STAT1, Pan troglodytes | 127 | IRF7, Bos taurus |
| 5 | STAT1, Macaca mulatta | 128 | IRF7, Gallus gallus |
| 6 | STAT1, Felis catus | 129 | MDA5, Homo sapiens |
| 7 | STAT1, Camelus dromedarius | 130 | MDA5, Pan troglodytes |
| 8 | STAT1, Equus caballus | 131 | MDA5, Macaca mulatta |
| 9 | STAT1, Canis lupus familiaris | 132 | MDA5, Equus caballus |
| 10 | STAT1, Rhinolophus ferrumequinum | 133 | MDA5, Ictidomys tridecemlineatus |
| 11 | STAT1, Bos taurus | 134 | MDA5, Felis catus |
| 12 | STAT1, Ovis aries | 135 | MDA5, Bos taurus |
| 13 | STAT1, Ictidomys tridecemlineatus | 136 | MDA5, Mus musculus |
| 14 | STAT1, Rattus norvegicus | 137 | MDA5, Rattus norvegicus |
| 15 | STAT1, Mus musculus | 138 | MDA5, Mesocricetus auratus |
| 16 | STAT1, Mesocricetus auratus | 139 | MDA5, Canis lupus familiaris |
| 17 | STAT1, Gallus gallus | 140 | MDA5, Gallus gallus |
| 18 | IRF9, Homo sapiens | 141 | PRKRA, Homo sapiens |
| 19 | IRF9, Macaca mulatta | 142 | PRKRA, Macaca mulatta |
| 20 | IRF9, Pan troglodytes | 143 | PRKRA, Equus caballus |
| 21 | IRF9, Felis catus | 144 | PRKRA, Canis lupus familiaris |
| 22 | IRF9, Camelus dromedarius | 145 | PRKRA, Camelus dromedarius |
| 23 | IRF9, Sus scrofa domesticus | 146 | PRKRA, Mesocricetus auratus |
| 24 | IRF9, Equus caballus | 147 | PRKRA, Mus musculus |
| 25 | IRF9, Rhinolophus ferrumequinum | 148 | PRKRA, Ictidomys tridecemlineatus |
| 26 | IRF9, Ovis aries | 149 | PRKRA, Ovis aries |
| 27 | IRF9, Canis lupus familiaris | 150 | PRKRA, Bos taurus |
| 28 | IRF9, Bos taurus | 151 | PRKRA, Rattus norvegicus |
| 29 | IRF9, Ictidomys tridecemlineatus | 152 | PRKRA, Rhinolophus ferrumequinum |
| 30 | IRF9, Mus musculus | 153 | PRKRA, Pan troglodytes |
| 31 | IRF9, Mesocricetus auratus | 154 | ORF3b, Severe acute respiratory syndrome-related coronavirus |
| 32 | IRF9, Rattus norvegicus | 155 | ORF3b, Human coronavirus NL63 |
| 33 | IRF9, Gallus gallus | 156 | DPP4, Homo sapiens |
| 34 | RIG-I, Homo sapiens | 157 | DPP4, Pan troglodytes |
| 35 | RIG-I, Pan troglodytes | 158 | DPP4, Macaca mulatta |
| 36 | RIG-I, Macaca mulatta | 159 | DPP4, Ovis aries |
| 37 | RIG-I, Rhinolophus ferrumequinum | 160 | DPP4, Ictidomys tridecemlineatus |
| 38 | RIG-I, Canis lupus familiaris | 161 | DPP4, Bos taurus |
| 39 | RIG-I, Mesocricetus auratus | 162 | DPP4, Felis catus |
| 40 | RIG-I, Mus musculus | 163 | DPP4, Equus caballus |
| 41 | RIG-I, Gallus gallus | 164 | DPP4, Rhinolophus ferrumequinum |
| 42 | ORF4b, Middle East respiratory syndrome-related coronavirus | 165 | DPP4, Mesocricetus auratus |
| 43 | ORF4b, Human coronavirus 229E | 166 | DPP4, Rattus norvegicus |
| 44 | nsp1, Severe acute respiratory syndrome coronavirus 2 | 167 | DPP4, Mus musculus |
| 45 | nsp1, Severe acute respiratory syndrome-related coronavirus | 168 | DPP4, Camelus dromedarius |
| 46 | nsp1, Middle East respiratory syndrome-related coronavirus | 169 | DPP4, Gallus gallus |
| 47 | nsp1, Human coronavirus HKU1 | 170 | ORF6, Severe acute respiratory syndrome-related coronavirus |
| 48 | spike, Human coronavirus OC43 | 171 | ORF6, Severe acute respiratory syndrome coronavirus 2 |
| 49 | spike, Human coronavirus HKU1 | 172 | STAT2, Homo sapiens |
| 50 | spike, Middle East respiratory syndrome-related coronavirus | 173 | STAT2, Pan troglodytes |
| 51 | spike, Severe acute respiratory syndrome coronavirus 2 | 174 | STAT2, Macaca mulatta |
| 52 | spike, Severe acute respiratory syndrome-related coronavirus | 175 | STAT2, Felis catus |
| 53 | spike, Human coronavirus NL63 | 176 | STAT2, Canis lupus familiaris |
| 54 | spike, Human coronavirus 229E | 177 | STAT2, Camelus dromedarius |
| 55 | IRF3, Homo sapiens | 178 | STAT2, Rhinolophus ferrumequinum |
| 56 | IRF3, Pan troglodytes | 179 | STAT2, Equus caballus |
| 57 | IRF3, Macaca mulatta | 180 | STAT2, Ovis aries |
| 58 | IRF3, Ictidomys tridecemlineatus | 181 | STAT2, Bos taurus |
| 59 | IRF3, Rhinolophus ferrumequinum | 182 | STAT2, Mesocricetus auratus |
| 60 | IRF3, Camelus dromedarius | 183 | STAT2, Rattus norvegicus |
| 61 | IRF3, Ovis aries | 184 | STAT2, Ictidomys tridecemlineatus |
| 62 | IRF3, Bos taurus | 185 | STAT2, Mus musculus |
| 63 | IRF3, Equus caballus | 186 | STAT2, Gallus gallus |
| 64 | IRF3, Rattus norvegicus | 187 | PLpro, Middle East respiratory syndrome-related coronavirus |
| 65 | IRF3, Mesocricetus auratus | 188 | PLpro, Severe acute respiratory syndrome-related coronavirus |
| 66 | IRF3, Mus musculus | 189 | PLpro, Severe acute respiratory syndrome coronavirus 2 |
| 67 | IRF3, Canis lupus familiaris | 190 | PLpro, Human coronavirus OC43 |
| 68 | IRF3, Gallus gallus | 191 | PLpro, Human coronavirus HKU1 |
| 69 | N protein, Middle East respiratory syndrome-related coronavirus | 192 | Homo sapiens |
| 70 | N protein, Severe acute respiratory syndrome coronavirus 2 | 193 | Mus musculus |
| 71 | N protein, Severe acute respiratory syndrome-related coronavirus | 194 | Rattus norvegicus |
| 72 | N protein, Human coronavirus HKU1 | 195 | Canis lupus familiaris |
| 73 | N protein, Human coronavirus OC43 | 196 | Camelus dromedarius |
| 74 | N protein, Human coronavirus 229E | 197 | Felis catus |
| 75 | N protein, Human coronavirus NL63 | 198 | Ictidomys tridecemlineatus |
| 76 | Human coronavirus OC43 | 199 | Bos taurus |
| 77 | Human coronavirus HKU1 | 200 | Pan troglodytes |

**Table 1 continued from previous page**

| Node ID | Full name representation | Node ID | Full name representation |
|---|---|---|---|
| 78 | Middle East respiratory syndrome-related coronavirus | 201 | Gallus gallus |
| 79 | Severe acute respiratory syndrome coronavirus 2 | 202 | Oryctolagus cuniculus |
| 80 | Severe acute respiratory syndrome-related coronavirus | 203 | Equus caballus |
| 81 | Human coronavirus NL63 | 204 | Macaca mulatta |
| 82 | Human coronavirus 229E | 205 | Ovis aries |
| 83 | ACE2, Homo sapiens | 206 | Sus scrofa domesticus |
| 84 | ACE2, Pan troglodytes | 207 | Rhinolophus ferrumequinum |
| 85 | ACE2, Macaca mulatta | 208 | Mesocricetus auratus |
| 86 | ACE2, Ictidomys tridecemlineatus | 209 | M protein, Middle East respiratory syndrome-related coronavirus |
| 87 | ACE2, Oryctolagus cuniculus | 210 | M protein, Human coronavirus HKU1 |
| 88 | ACE2, Equus caballus | 211 | M protein, Severe acute respiratory syndrome-related coronavirus |
| 89 | ACE2, Felis catus | 212 | M protein, Severe acute respiratory syndrome coronavirus 2 |
| 90 | ACE2, Camelus dromedarius | 213 | M protein, Human coronavirus OC43 |
| 91 | ACE2, Mesocricetus auratus | 214 | M protein, Human coronavirus NL63 |
| 92 | ACE2, Sus scrofa domesticus | 215 | M protein, Human coronavirus 229E |
| 93 | ACE2, Ovis aries | 216 | TBK1, Homo sapiens |
| 94 | ACE2, Mus musculus | 217 | TBK1, Pan troglodytes |
| 95 | ACE2, Bos taurus | 218 | TBK1, Macaca mulatta |
| 96 | ACE2, Rattus norvegicus | 219 | TBK1, Ictidomys tridecemlineatus |
| 97 | ACE2, Rhinolophus ferrumequinum | 220 | TBK1, Felis catus |
| 98 | ACE2, Canis lupus familiaris | 221 | TBK1, Bos taurus |
| 99 | ACE2, Gallus gallus | 222 | TBK1, Rhinolophus ferrumequinum |
| 100 | ORF4a, Middle East respiratory syndrome-related coronavirus | 223 | TBK1, Ovis aries |
| 101 | ORF4a, Human coronavirus 229E | 224 | TBK1, Canis lupus familiaris |
| 102 | NF-$\kappa$B, Homo sapiens | 225 | TBK1, Equus caballus |
| 103 | NF-$\kappa$B, Pan troglodytes | 226 | TBK1, Mesocricetus auratus |
| 104 | NF-$\kappa$B, Macaca mulatta | 227 | TBK1, Mus musculus |
| 105 | NF-$\kappa$B, Bos taurus | 228 | TBK1, Camelus dromedarius |
| 106 | NF-$\kappa$B, Rhinolophus ferrumequinum | 229 | TBK1, Rattus norvegicus |
| 107 | NF-$\kappa$B, Ovis aries | 230 | TBK1, Gallus gallus |
| 108 | NF-$\kappa$B, Ictidomys tridecemlineatus | 231 | MAVS, Homo sapiens |
| 109 | NF-$\kappa$B, Mus musculus | 232 | MAVS, Pan troglodytes |
| 110 | NF-$\kappa$B, Mesocricetus auratus | 233 | MAVS, Macaca mulatta |
| 111 | NF-$\kappa$B, Rattus norvegicus | 234 | MAVS, Ovis aries |
| 112 | NF-$\kappa$B, Equus caballus | 235 | MAVS, Equus caballus |
| 113 | NF-$\kappa$B, Gallus gallus | 236 | MAVS, Bos taurus |
| 114 | NF-$\kappa$B, Camelus dromedarius | 237 | MAVS, Rhinolophus ferrumequinum |
| 115 | IRF7, Homo sapiens | 238 | MAVS, Felis catus |
| 116 | IRF7, Pan troglodytes | 239 | MAVS, Camelus dromedarius |
| 117 | IRF7, Macaca mulatta | 240 | MAVS, Ictidomys tridecemlineatus |
| 118 | IRF7, Ictidomys tridecemlineatus | 241 | MAVS, Canis lupus familiaris |
| 119 | IRF7, Equus caballus | 242 | MAVS, Mesocricetus auratus |
| 120 | IRF7, Rhinolophus ferrumequinum | 243 | MAVS, Mus musculus |
| 121 | IRF7, Felis catus | 244 | MAVS, Rattus norvegicus |
| 122 | IRF7, Camelus dromedarius | 245 | MAVS, Gallus gallus |

### Supplementary Note 7: Original Network's interaction Table with interaction Types

| Source ID | Source Name | Target ID | Target Name | Relation |
|---|---|---|---|---|
| 0 | nsp15 Severe acute respiratory syndrome coronavirus 2 | 1 | nsp15 Severe acute respiratory syndrome-related coronavirus | similar |
| 0 | nsp15 Severe acute respiratory syndrome coronavirus 2 | 2 | nsp15 Human coronavirus HKU1 | similar |
| 0 | nsp15 Severe acute respiratory syndrome coronavirus 2 | 79 | Severe acute respiratory syndrome coronavirus 2 | belongs |
| 0 | nsp15 Severe acute respiratory syndrome coronavirus 2 | 55 | IRF3 Homo sapiens | interacts |
| 0 | nsp15 Severe acute respiratory syndrome coronavirus 2 | 65 | IRF3 Mesocricetus auratus | interacts |
| 0 | nsp15 Severe acute respiratory syndrome coronavirus 2 | 66 | IRF3 Mus musculus | interacts |
| 0 | nsp15 Severe acute respiratory syndrome coronavirus 2 | 67 | IRF3 Canis lupus familiaris | interacts |
| 0 | nsp15 Severe acute respiratory syndrome coronavirus 2 | 34 | RIG-I Homo sapiens | interacts |
| 0 | nsp15 Severe acute respiratory syndrome coronavirus 2 | 38 | RIG-I Canis lupus familiaris | interacts |
| 0 | nsp15 Severe acute respiratory syndrome coronavirus 2 | 39 | RIG-I Mesocricetus auratus | interacts |
| 0 | nsp15 Severe acute respiratory syndrome coronavirus 2 | 40 | RIG-I Mus musculus | interacts |
| 1 | nsp15 Severe acute respiratory syndrome-related coronavirus | 2 | nsp15 Human coronavirus HKU1 | similar |
| 1 | nsp15 Severe acute respiratory syndrome-related coronavirus | 80 | Severe acute respiratory syndrome-related coronavirus | belongs |
| 1 | nsp15 Severe acute respiratory syndrome-related coronavirus | 55 | IRF3 Homo sapiens | interacts |
| 1 | nsp15 Severe acute respiratory syndrome-related coronavirus | 59 | IRF3 Rhinolophus ferrumequinum | interacts |
| 1 | nsp15 Severe acute respiratory syndrome-related coronavirus | 231 | MAVS Homo sapiens | interacts |
| 1 | nsp15 Severe acute respiratory syndrome-related coronavirus | 237 | MAVS Rhinolophus ferrumequinum | interacts |
| 1 | nsp15 Severe acute respiratory syndrome-related coronavirus | 238 | MAVS Felis catus | interacts |
| 2 | nsp15 Human coronavirus HKU1 | 77 | Human coronavirus HKU1 | belongs |
| 3 | STAT1 Homo sapiens | 4 | STAT1 Pan troglodytes | similar |
| 3 | STAT1 Homo sapiens | 5 | STAT1 Macaca mulatta | similar |
| 3 | STAT1 Homo sapiens | 6 | STAT1 Felis catus | similar |
| 3 | STAT1 Homo sapiens | 7 | STAT1 Camelus dromedarius | similar |
| 3 | STAT1 Homo sapiens | 8 | STAT1 Equus caballus | similar |
| 3 | STAT1 Homo sapiens | 9 | STAT1 Canis lupus familiaris | similar |
| 3 | STAT1 Homo sapiens | 10 | STAT1 Rhinolophus ferrumequinum | similar |
| 3 | STAT1 Homo sapiens | 11 | STAT1 Bos taurus | similar |
| 3 | STAT1 Homo sapiens | 12 | STAT1 Ovis aries | similar |
| 3 | STAT1 Homo sapiens | 13 | STAT1 Ictidomys tridecemlineatus | similar |
| 3 | STAT1 Homo sapiens | 14 | STAT1 Rattus norvegicus | similar |
| 3 | STAT1 Homo sapiens | 15 | STAT1 Mus musculus | similar |
| 3 | STAT1 Homo sapiens | 16 | STAT1 Mesocricetus auratus | similar |
| 3 | STAT1 Homo sapiens | 17 | STAT1 Gallus gallus | similar |
| 3 | STAT1 Homo sapiens | 192 | Homo sapiens | belongs |
| 3 | STAT1 Homo sapiens | 154 | ORF3b Severe acute respiratory syndrome-related coronavirus | interacts |
| 3 | STAT1 Homo sapiens | 170 | ORF6 Severe acute respiratory syndrome-related coronavirus | interacts |
| 3 | STAT1 Homo sapiens | 209 | M protein Middle East respiratory syndrome-related coronavirus | interacts |
| 3 | STAT1 Homo sapiens | 45 | nsp1 Severe acute respiratory syndrome-related coronavirus | interacts |
| 3 | STAT1 Homo sapiens | 100 | ORF4a Middle East respiratory syndrome-related coronavirus | interacts |
| 3 | STAT1 Homo sapiens | 42 | ORF4b Middle East respiratory syndrome-related coronavirus | interacts |
| 4 | STAT1 Pan troglodytes | 5 | STAT1 Macaca mulatta | similar |
| 4 | STAT1 Pan troglodytes | 6 | STAT1 Felis catus | similar |
| 4 | STAT1 Pan troglodytes | 7 | STAT1 Camelus dromedarius | similar |
| 4 | STAT1 Pan troglodytes | 8 | STAT1 Equus caballus | similar |
| 4 | STAT1 Pan troglodytes | 9 | STAT1 Canis lupus familiaris | similar |
| 4 | STAT1 Pan troglodytes | 10 | STAT1 Rhinolophus ferrumequinum | similar |
| 4 | STAT1 Pan troglodytes | 11 | STAT1 Bos taurus | similar |
| 4 | STAT1 Pan troglodytes | 12 | STAT1 Ovis aries | similar |
| 4 | STAT1 Pan troglodytes | 13 | STAT1 Ictidomys tridecemlineatus | similar |
| 4 | STAT1 Pan troglodytes | 14 | STAT1 Rattus norvegicus | similar |
| 4 | STAT1 Pan troglodytes | 15 | STAT1 Mus musculus | similar |
| 4 | STAT1 Pan troglodytes | 16 | STAT1 Mesocricetus auratus | similar |
| 4 | STAT1 Pan troglodytes | 17 | STAT1 Gallus gallus | similar |
| 4 | STAT1 Pan troglodytes | 200 | Pan troglodytes | belongs |
| 5 | STAT1 Macaca mulatta | 6 | STAT1 Felis catus | similar |
| 5 | STAT1 Macaca mulatta | 7 | STAT1 Camelus dromedarius | similar |
| 5 | STAT1 Macaca mulatta | 8 | STAT1 Equus caballus | similar |
| 5 | STAT1 Macaca mulatta | 9 | STAT1 Canis lupus familiaris | similar |
| 5 | STAT1 Macaca mulatta | 10 | STAT1 Rhinolophus ferrumequinum | similar |
| 5 | STAT1 Macaca mulatta | 11 | STAT1 Bos taurus | similar |
| 5 | STAT1 Macaca mulatta | 12 | STAT1 Ovis aries | similar |
| 5 | STAT1 Macaca mulatta | 13 | STAT1 Ictidomys tridecemlineatus | similar |
| 5 | STAT1 Macaca mulatta | 14 | STAT1 Rattus norvegicus | similar |
| 5 | STAT1 Macaca mulatta | 15 | STAT1 Mus musculus | similar |
| 5 | STAT1 Macaca mulatta | 16 | STAT1 Mesocricetus auratus | similar |
| 5 | STAT1 Macaca mulatta | 17 | STAT1 Gallus gallus | similar |
| 5 | STAT1 Macaca mulatta | 204 | Macaca mulatta | belongs |
| 6 | STAT1 Felis catus | 7 | STAT1 Camelus dromedarius | similar |
| 6 | STAT1 Felis catus | 8 | STAT1 Equus caballus | similar |
| 6 | STAT1 Felis catus | 9 | STAT1 Canis lupus familiaris | similar |
| 6 | STAT1 Felis catus | 10 | STAT1 Rhinolophus ferrumequinum | similar |
| 6 | STAT1 Felis catus | 11 | STAT1 Bos taurus | similar |
| 6 | STAT1 Felis catus | 12 | STAT1 Ovis aries | similar |
| 6 | STAT1 Felis catus | 13 | STAT1 Ictidomys tridecemlineatus | similar |
| 6 | STAT1 Felis catus | 14 | STAT1 Rattus norvegicus | similar |
| 6 | STAT1 Felis catus | 15 | STAT1 Mus musculus | similar |
| 6 | STAT1 Felis catus | 16 | STAT1 Mesocricetus auratus | similar |
| 6 | STAT1 Felis catus | 17 | STAT1 Gallus gallus | similar |

**Table 2 continued from previous page**

| Source ID | Source Name | Target ID | Target Name | Relation |
|---|---|---|---|---|
| 6 | STAT1 Felis catus | 197 | Felis catus | belongs |
| 6 | STAT1 Felis catus | 154 | ORF3b Severe acute respiratory syndrome-related coronavirus | interacts |
| 6 | STAT1 Felis catus | 170 | ORF6 Severe acute respiratory syndrome-related coronavirus | interacts |
| 6 | STAT1 Felis catus | 45 | nsp1 Severe acute respiratory syndrome-related coronavirus | interacts |
| 7 | STAT1 Camelus dromedarius | 8 | STAT1 Equus caballus | similar |
| 7 | STAT1 Camelus dromedarius | 9 | STAT1 Canis lupus familiaris | similar |
| 7 | STAT1 Camelus dromedarius | 10 | STAT1 Rhinolophus ferrumequinum | similar |
| 7 | STAT1 Camelus dromedarius | 11 | STAT1 Bos taurus | similar |
| 7 | STAT1 Camelus dromedarius | 12 | STAT1 Ovis aries | similar |
| 7 | STAT1 Camelus dromedarius | 13 | STAT1 Ictidomys tridecemlineatus | similar |
| 7 | STAT1 Camelus dromedarius | 14 | STAT1 Rattus norvegicus | similar |
| 7 | STAT1 Camelus dromedarius | 15 | STAT1 Mus musculus | similar |
| 7 | STAT1 Camelus dromedarius | 16 | STAT1 Mesocricetus auratus | similar |
| 7 | STAT1 Camelus dromedarius | 17 | STAT1 Gallus gallus | similar |
| 7 | STAT1 Camelus dromedarius | 196 | Camelus dromedarius | belongs |
| 7 | STAT1 Camelus dromedarius | 209 | M protein Middle East respiratory syndrome-related coronavirus | interacts |
| 7 | STAT1 Camelus dromedarius | 100 | ORF4a Middle East respiratory syndrome-related coronavirus | interacts |
| 7 | STAT1 Camelus dromedarius | 42 | ORF4b Middle East respiratory syndrome-related coronavirus | interacts |
| 8 | STAT1 Equus caballus | 9 | STAT1 Canis lupus familiaris | similar |
| 8 | STAT1 Equus caballus | 10 | STAT1 Rhinolophus ferrumequinum | similar |
| 8 | STAT1 Equus caballus | 11 | STAT1 Bos taurus | similar |
| 8 | STAT1 Equus caballus | 12 | STAT1 Ovis aries | similar |
| 8 | STAT1 Equus caballus | 13 | STAT1 Ictidomys tridecemlineatus | similar |
| 8 | STAT1 Equus caballus | 14 | STAT1 Rattus norvegicus | similar |
| 8 | STAT1 Equus caballus | 15 | STAT1 Mus musculus | similar |
| 8 | STAT1 Equus caballus | 16 | STAT1 Mesocricetus auratus | similar |
| 8 | STAT1 Equus caballus | 17 | STAT1 Gallus gallus | similar |
| 8 | STAT1 Equus caballus | 203 | Equus caballus | belongs |
| 9 | STAT1 Canis lupus familiaris | 10 | STAT1 Rhinolophus ferrumequinum | similar |
| 9 | STAT1 Canis lupus familiaris | 11 | STAT1 Bos taurus | similar |
| 9 | STAT1 Canis lupus familiaris | 12 | STAT1 Ovis aries | similar |
| 9 | STAT1 Canis lupus familiaris | 13 | STAT1 Ictidomys tridecemlineatus | similar |
| 9 | STAT1 Canis lupus familiaris | 14 | STAT1 Rattus norvegicus | similar |
| 9 | STAT1 Canis lupus familiaris | 15 | STAT1 Mus musculus | similar |
| 9 | STAT1 Canis lupus familiaris | 16 | STAT1 Mesocricetus auratus | similar |
| 9 | STAT1 Canis lupus familiaris | 17 | STAT1 Gallus gallus | similar |
| 9 | STAT1 Canis lupus familiaris | 195 | Canis lupus familiaris | belongs |
| 10 | STAT1 Rhinolophus ferrumequinum | 11 | STAT1 Bos taurus | similar |
| 10 | STAT1 Rhinolophus ferrumequinum | 12 | STAT1 Ovis aries | similar |
| 10 | STAT1 Rhinolophus ferrumequinum | 13 | STAT1 Ictidomys tridecemlineatus | similar |
| 10 | STAT1 Rhinolophus ferrumequinum | 14 | STAT1 Rattus norvegicus | similar |
| 10 | STAT1 Rhinolophus ferrumequinum | 15 | STAT1 Mus musculus | similar |
| 10 | STAT1 Rhinolophus ferrumequinum | 16 | STAT1 Mesocricetus auratus | similar |
| 10 | STAT1 Rhinolophus ferrumequinum | 17 | STAT1 Gallus gallus | similar |
| 10 | STAT1 Rhinolophus ferrumequinum | 207 | Rhinolophus ferrumequinum | belongs |
| 10 | STAT1 Rhinolophus ferrumequinum | 154 | ORF3b Severe acute respiratory syndrome-related coronavirus | interacts |
| 10 | STAT1 Rhinolophus ferrumequinum | 170 | ORF6 Severe acute respiratory syndrome-related coronavirus | interacts |
| 10 | STAT1 Rhinolophus ferrumequinum | 45 | nsp1 Severe acute respiratory syndrome-related coronavirus | interacts |
| 11 | STAT1 Bos taurus | 12 | STAT1 Ovis aries | similar |
| 11 | STAT1 Bos taurus | 13 | STAT1 Ictidomys tridecemlineatus | similar |
| 11 | STAT1 Bos taurus | 14 | STAT1 Rattus norvegicus | similar |
| 11 | STAT1 Bos taurus | 15 | STAT1 Mus musculus | similar |
| 11 | STAT1 Bos taurus | 16 | STAT1 Mesocricetus auratus | similar |
| 11 | STAT1 Bos taurus | 17 | STAT1 Gallus gallus | similar |
| 11 | STAT1 Bos taurus | 199 | Bos taurus | belongs |
| 12 | STAT1 Ovis aries | 13 | STAT1 Ictidomys tridecemlineatus | similar |
| 12 | STAT1 Ovis aries | 14 | STAT1 Rattus norvegicus | similar |
| 12 | STAT1 Ovis aries | 15 | STAT1 Mus musculus | similar |
| 12 | STAT1 Ovis aries | 16 | STAT1 Mesocricetus auratus | similar |
| 12 | STAT1 Ovis aries | 17 | STAT1 Gallus gallus | similar |
| 12 | STAT1 Ovis aries | 205 | Ovis aries | belongs |
| 13 | STAT1 Ictidomys tridecemlineatus | 14 | STAT1 Rattus norvegicus | similar |
| 13 | STAT1 Ictidomys tridecemlineatus | 15 | STAT1 Mus musculus | similar |
| 13 | STAT1 Ictidomys tridecemlineatus | 16 | STAT1 Mesocricetus auratus | similar |
| 13 | STAT1 Ictidomys tridecemlineatus | 17 | STAT1 Gallus gallus | similar |
| 13 | STAT1 Ictidomys tridecemlineatus | 198 | Ictidomys tridecemlineatus | belongs |
| 14 | STAT1 Rattus norvegicus | 15 | STAT1 Mus musculus | similar |
| 14 | STAT1 Rattus norvegicus | 16 | STAT1 Mesocricetus auratus | similar |
| 14 | STAT1 Rattus norvegicus | 17 | STAT1 Gallus gallus | similar |
| 14 | STAT1 Rattus norvegicus | 194 | Rattus norvegicus | belongs |
| 14 | STAT1 Rattus norvegicus | 154 | ORF3b Severe acute respiratory syndrome-related coronavirus | interacts |
| 14 | STAT1 Rattus norvegicus | 170 | ORF6 Severe acute respiratory syndrome-related coronavirus | interacts |
| 14 | STAT1 Rattus norvegicus | 209 | M protein Middle East respiratory syndrome-related coronavirus | interacts |
| 14 | STAT1 Rattus norvegicus | 45 | nsp1 Severe acute respiratory syndrome-related coronavirus | interacts |
| 14 | STAT1 Rattus norvegicus | 100 | ORF4a Middle East respiratory syndrome-related coronavirus | interacts |
| 14 | STAT1 Rattus norvegicus | 42 | ORF4b Middle East respiratory syndrome-related coronavirus | interacts |
| 15 | STAT1 Mus musculus | 16 | STAT1 Mesocricetus auratus | similar |
| 15 | STAT1 Mus musculus | 17 | STAT1 Gallus gallus | similar |
| 15 | STAT1 Mus musculus | 193 | Mus musculus | belongs |
| 15 | STAT1 Mus musculus | 154 | ORF3b Severe acute respiratory syndrome-related coronavirus | interacts |

**Table 2 continued from previous page**

| Source ID | Source Name | Target ID | Target Name | Relation |
|---|---|---|---|---|
| 15 | STAT1 Mus musculus | 170 | ORF6 Severe acute respiratory syndrome-related coronavirus | interacts |
| 15 | STAT1 Mus musculus | 209 | M protein Middle East respiratory syndrome-related coronavirus | interacts |
| 15 | STAT1 Mus musculus | 45 | nsp1 Severe acute respiratory syndrome-related coronavirus | interacts |
| 15 | STAT1 Mus musculus | 100 | ORF4a Middle East respiratory syndrome-related coronavirus | interacts |
| 15 | STAT1 Mus musculus | 42 | ORF4b Middle East respiratory syndrome-related coronavirus | interacts |
| 16 | STAT1 Mesocricetus auratus | 17 | STAT1 Gallus gallus | similar |
| 16 | STAT1 Mesocricetus auratus | 208 | Mesocricetus auratus | belongs |
| 17 | STAT1 Gallus gallus | 201 | Gallus gallus | belongs |
| 18 | IRF9 Homo sapiens | 19 | IRF9 Macaca mulatta | similar |
| 18 | IRF9 Homo sapiens | 20 | IRF9 Pan troglodytes | similar |
| 18 | IRF9 Homo sapiens | 21 | IRF9 Felis catus | similar |
| 18 | IRF9 Homo sapiens | 22 | IRF9 Camelus dromedarius | similar |
| 18 | IRF9 Homo sapiens | 23 | IRF9 Sus scrofa domesticus | similar |
| 18 | IRF9 Homo sapiens | 24 | IRF9 Equus caballus | similar |
| 18 | IRF9 Homo sapiens | 25 | IRF9 Rhinolophus ferrumequinum | similar |
| 18 | IRF9 Homo sapiens | 26 | IRF9 Ovis aries | similar |
| 18 | IRF9 Homo sapiens | 27 | IRF9 Canis lupus familiaris | similar |
| 18 | IRF9 Homo sapiens | 28 | IRF9 Bos taurus | similar |
| 18 | IRF9 Homo sapiens | 29 | IRF9 Ictidomys tridecemlineatus | similar |
| 18 | IRF9 Homo sapiens | 30 | IRF9 Mus musculus | similar |
| 18 | IRF9 Homo sapiens | 31 | IRF9 Mesocricetus auratus | similar |
| 18 | IRF9 Homo sapiens | 32 | IRF9 Rattus norvegicus | similar |
| 18 | IRF9 Homo sapiens | 33 | IRF9 Gallus gallus | similar |
| 18 | IRF9 Homo sapiens | 192 | Homo sapiens | belongs |
| 18 | IRF9 Homo sapiens | 154 | ORF3b Severe acute respiratory syndrome-related coronavirus | interacts |
| 18 | IRF9 Homo sapiens | 170 | ORF6 Severe acute respiratory syndrome-related coronavirus | interacts |
| 18 | IRF9 Homo sapiens | 209 | M protein Middle East respiratory syndrome-related coronavirus | interacts |
| 18 | IRF9 Homo sapiens | 100 | ORF4a Middle East respiratory syndrome-related coronavirus | interacts |
| 18 | IRF9 Homo sapiens | 42 | ORF4b Middle East respiratory syndrome-related coronavirus | interacts |
| 19 | IRF9 Macaca mulatta | 20 | IRF9 Pan troglodytes | similar |
| 19 | IRF9 Macaca mulatta | 21 | IRF9 Felis catus | similar |
| 19 | IRF9 Macaca mulatta | 22 | IRF9 Camelus dromedarius | similar |
| 19 | IRF9 Macaca mulatta | 23 | IRF9 Sus scrofa domesticus | similar |
| 19 | IRF9 Macaca mulatta | 24 | IRF9 Equus caballus | similar |
| 19 | IRF9 Macaca mulatta | 25 | IRF9 Rhinolophus ferrumequinum | similar |
| 19 | IRF9 Macaca mulatta | 26 | IRF9 Ovis aries | similar |
| 19 | IRF9 Macaca mulatta | 27 | IRF9 Canis lupus familiaris | similar |
| 19 | IRF9 Macaca mulatta | 28 | IRF9 Bos taurus | similar |
| 19 | IRF9 Macaca mulatta | 29 | IRF9 Ictidomys tridecemlineatus | similar |
| 19 | IRF9 Macaca mulatta | 30 | IRF9 Mus musculus | similar |
| 19 | IRF9 Macaca mulatta | 31 | IRF9 Mesocricetus auratus | similar |
| 19 | IRF9 Macaca mulatta | 32 | IRF9 Rattus norvegicus | similar |
| 19 | IRF9 Macaca mulatta | 33 | IRF9 Gallus gallus | similar |
| 19 | IRF9 Macaca mulatta | 204 | Macaca mulatta | belongs |
| 20 | IRF9 Pan troglodytes | 21 | IRF9 Felis catus | similar |
| 20 | IRF9 Pan troglodytes | 22 | IRF9 Camelus dromedarius | similar |
| 20 | IRF9 Pan troglodytes | 23 | IRF9 Sus scrofa domesticus | similar |
| 20 | IRF9 Pan troglodytes | 24 | IRF9 Equus caballus | similar |
| 20 | IRF9 Pan troglodytes | 25 | IRF9 Rhinolophus ferrumequinum | similar |
| 20 | IRF9 Pan troglodytes | 26 | IRF9 Ovis aries | similar |
| 20 | IRF9 Pan troglodytes | 27 | IRF9 Canis lupus familiaris | similar |
| 20 | IRF9 Pan troglodytes | 28 | IRF9 Bos taurus | similar |
| 20 | IRF9 Pan troglodytes | 29 | IRF9 Ictidomys tridecemlineatus | similar |
| 20 | IRF9 Pan troglodytes | 30 | IRF9 Mus musculus | similar |
| 20 | IRF9 Pan troglodytes | 31 | IRF9 Mesocricetus auratus | similar |
| 20 | IRF9 Pan troglodytes | 32 | IRF9 Rattus norvegicus | similar |
| 20 | IRF9 Pan troglodytes | 33 | IRF9 Gallus gallus | similar |
| 20 | IRF9 Pan troglodytes | 200 | Pan troglodytes | belongs |
| 21 | IRF9 Felis catus | 22 | IRF9 Camelus dromedarius | similar |
| 21 | IRF9 Felis catus | 23 | IRF9 Sus scrofa domesticus | similar |
| 21 | IRF9 Felis catus | 24 | IRF9 Equus caballus | similar |
| 21 | IRF9 Felis catus | 25 | IRF9 Rhinolophus ferrumequinum | similar |
| 21 | IRF9 Felis catus | 26 | IRF9 Ovis aries | similar |
| 21 | IRF9 Felis catus | 27 | IRF9 Canis lupus familiaris | similar |
| 21 | IRF9 Felis catus | 28 | IRF9 Bos taurus | similar |
| 21 | IRF9 Felis catus | 29 | IRF9 Ictidomys tridecemlineatus | similar |
| 21 | IRF9 Felis catus | 30 | IRF9 Mus musculus | similar |
| 21 | IRF9 Felis catus | 31 | IRF9 Mesocricetus auratus | similar |
| 21 | IRF9 Felis catus | 32 | IRF9 Rattus norvegicus | similar |
| 21 | IRF9 Felis catus | 33 | IRF9 Gallus gallus | similar |
| 21 | IRF9 Felis catus | 197 | Felis catus | belongs |
| 21 | IRF9 Felis catus | 154 | ORF3b Severe acute respiratory syndrome-related coronavirus | interacts |
| 21 | IRF9 Felis catus | 170 | ORF6 Severe acute respiratory syndrome-related coronavirus | interacts |
| 22 | IRF9 Camelus dromedarius | 23 | IRF9 Sus scrofa domesticus | similar |
| 22 | IRF9 Camelus dromedarius | 24 | IRF9 Equus caballus | similar |
| 22 | IRF9 Camelus dromedarius | 25 | IRF9 Rhinolophus ferrumequinum | similar |
| 22 | IRF9 Camelus dromedarius | 26 | IRF9 Ovis aries | similar |
| 22 | IRF9 Camelus dromedarius | 27 | IRF9 Canis lupus familiaris | similar |
| 22 | IRF9 Camelus dromedarius | 28 | IRF9 Bos taurus | similar |
| 22 | IRF9 Camelus dromedarius | 29 | IRF9 Ictidomys tridecemlineatus | similar |

**Table 2 continued from previous page**

| Source ID | Source Name | Target ID | Target Name | Relation |
|---|---|---|---|---|
| 22 | IRF9 Camelus dromedarius | 30 | IRF9 Mus musculus | similar |
| 22 | IRF9 Camelus dromedarius | 31 | IRF9 Mesocricetus auratus | similar |
| 22 | IRF9 Camelus dromedarius | 32 | IRF9 Rattus norvegicus | similar |
| 22 | IRF9 Camelus dromedarius | 33 | IRF9 Gallus gallus | similar |
| 22 | IRF9 Camelus dromedarius | 196 | Camelus dromedarius | belongs |
| 22 | IRF9 Camelus dromedarius | 209 | M protein Middle East respiratory syndrome-related coronavirus | interacts |
| 22 | IRF9 Camelus dromedarius | 100 | ORF4a Middle East respiratory syndrome-related coronavirus | interacts |
| 22 | IRF9 Camelus dromedarius | 42 | ORF4b Middle East respiratory syndrome-related coronavirus | interacts |
| 23 | IRF9 Sus scrofa domesticus | 24 | IRF9 Equus caballus | similar |
| 23 | IRF9 Sus scrofa domesticus | 25 | IRF9 Rhinolophus ferrumequinum | similar |
| 23 | IRF9 Sus scrofa domesticus | 26 | IRF9 Ovis aries | similar |
| 23 | IRF9 Sus scrofa domesticus | 27 | IRF9 Canis lupus familiaris | similar |
| 23 | IRF9 Sus scrofa domesticus | 28 | IRF9 Bos taurus | similar |
| 23 | IRF9 Sus scrofa domesticus | 29 | IRF9 Ictidomys tridecemlineatus | similar |
| 23 | IRF9 Sus scrofa domesticus | 30 | IRF9 Mus musculus | similar |
| 23 | IRF9 Sus scrofa domesticus | 31 | IRF9 Mesocricetus auratus | similar |
| 23 | IRF9 Sus scrofa domesticus | 32 | IRF9 Rattus norvegicus | similar |
| 23 | IRF9 Sus scrofa domesticus | 33 | IRF9 Gallus gallus | similar |
| 23 | IRF9 Sus scrofa domesticus | 206 | Sus scrofa domesticus | belongs |
| 24 | IRF9 Equus caballus | 25 | IRF9 Rhinolophus ferrumequinum | similar |
| 24 | IRF9 Equus caballus | 26 | IRF9 Ovis aries | similar |
| 24 | IRF9 Equus caballus | 27 | IRF9 Canis lupus familiaris | similar |
| 24 | IRF9 Equus caballus | 28 | IRF9 Bos taurus | similar |
| 24 | IRF9 Equus caballus | 29 | IRF9 Ictidomys tridecemlineatus | similar |
| 24 | IRF9 Equus caballus | 30 | IRF9 Mus musculus | similar |
| 24 | IRF9 Equus caballus | 31 | IRF9 Mesocricetus auratus | similar |
| 24 | IRF9 Equus caballus | 32 | IRF9 Rattus norvegicus | similar |
| 24 | IRF9 Equus caballus | 33 | IRF9 Gallus gallus | similar |
| 24 | IRF9 Equus caballus | 203 | Equus caballus | belongs |
| 25 | IRF9 Rhinolophus ferrumequinum | 26 | IRF9 Ovis aries | similar |
| 25 | IRF9 Rhinolophus ferrumequinum | 27 | IRF9 Canis lupus familiaris | similar |
| 25 | IRF9 Rhinolophus ferrumequinum | 28 | IRF9 Bos taurus | similar |
| 25 | IRF9 Rhinolophus ferrumequinum | 29 | IRF9 Ictidomys tridecemlineatus | similar |
| 25 | IRF9 Rhinolophus ferrumequinum | 30 | IRF9 Mus musculus | similar |
| 25 | IRF9 Rhinolophus ferrumequinum | 31 | IRF9 Mesocricetus auratus | similar |
| 25 | IRF9 Rhinolophus ferrumequinum | 32 | IRF9 Rattus norvegicus | similar |
| 25 | IRF9 Rhinolophus ferrumequinum | 33 | IRF9 Gallus gallus | similar |
| 25 | IRF9 Rhinolophus ferrumequinum | 207 | Rhinolophus ferrumequinum | belongs |
| 25 | IRF9 Rhinolophus ferrumequinum | 154 | ORF3b Severe acute respiratory syndrome-related coronavirus | interacts |
| 25 | IRF9 Rhinolophus ferrumequinum | 170 | ORF6 Severe acute respiratory syndrome-related coronavirus | interacts |
| 26 | IRF9 Ovis aries | 27 | IRF9 Canis lupus familiaris | similar |
| 26 | IRF9 Ovis aries | 28 | IRF9 Bos taurus | similar |
| 26 | IRF9 Ovis aries | 29 | IRF9 Ictidomys tridecemlineatus | similar |
| 26 | IRF9 Ovis aries | 30 | IRF9 Mus musculus | similar |
| 26 | IRF9 Ovis aries | 31 | IRF9 Mesocricetus auratus | similar |
| 26 | IRF9 Ovis aries | 32 | IRF9 Rattus norvegicus | similar |
| 26 | IRF9 Ovis aries | 33 | IRF9 Gallus gallus | similar |
| 26 | IRF9 Ovis aries | 205 | Ovis aries | belongs |
| 27 | IRF9 Canis lupus familiaris | 28 | IRF9 Bos taurus | similar |
| 27 | IRF9 Canis lupus familiaris | 29 | IRF9 Ictidomys tridecemlineatus | similar |
| 27 | IRF9 Canis lupus familiaris | 30 | IRF9 Mus musculus | similar |
| 27 | IRF9 Canis lupus familiaris | 31 | IRF9 Mesocricetus auratus | similar |
| 27 | IRF9 Canis lupus familiaris | 32 | IRF9 Rattus norvegicus | similar |
| 27 | IRF9 Canis lupus familiaris | 33 | IRF9 Gallus gallus | similar |
| 27 | IRF9 Canis lupus familiaris | 195 | Canis lupus familiaris | belongs |
| 28 | IRF9 Bos taurus | 29 | IRF9 Ictidomys tridecemlineatus | similar |
| 28 | IRF9 Bos taurus | 30 | IRF9 Mus musculus | similar |
| 28 | IRF9 Bos taurus | 31 | IRF9 Mesocricetus auratus | similar |
| 28 | IRF9 Bos taurus | 32 | IRF9 Rattus norvegicus | similar |
| 28 | IRF9 Bos taurus | 33 | IRF9 Gallus gallus | similar |
| 28 | IRF9 Bos taurus | 199 | Bos taurus | belongs |
| 29 | IRF9 Ictidomys tridecemlineatus | 30 | IRF9 Mus musculus | similar |
| 29 | IRF9 Ictidomys tridecemlineatus | 31 | IRF9 Mesocricetus auratus | similar |
| 29 | IRF9 Ictidomys tridecemlineatus | 32 | IRF9 Rattus norvegicus | similar |
| 29 | IRF9 Ictidomys tridecemlineatus | 33 | IRF9 Gallus gallus | similar |
| 29 | IRF9 Ictidomys tridecemlineatus | 198 | Ictidomys tridecemlineatus | belongs |
| 30 | IRF9 Mus musculus | 31 | IRF9 Mesocricetus auratus | similar |
| 30 | IRF9 Mus musculus | 32 | IRF9 Rattus norvegicus | similar |
| 30 | IRF9 Mus musculus | 33 | IRF9 Gallus gallus | similar |
| 30 | IRF9 Mus musculus | 193 | Mus musculus | belongs |
| 30 | IRF9 Mus musculus | 154 | ORF3b Severe acute respiratory syndrome-related coronavirus | interacts |
| 30 | IRF9 Mus musculus | 170 | ORF6 Severe acute respiratory syndrome-related coronavirus | interacts |
| 30 | IRF9 Mus musculus | 209 | M protein Middle East respiratory syndrome-related coronavirus | interacts |
| 30 | IRF9 Mus musculus | 100 | ORF4a Middle East respiratory syndrome-related coronavirus | interacts |
| 30 | IRF9 Mus musculus | 42 | ORF4b Middle East respiratory syndrome-related coronavirus | interacts |
| 31 | IRF9 Mesocricetus auratus | 32 | IRF9 Rattus norvegicus | similar |
| 31 | IRF9 Mesocricetus auratus | 33 | IRF9 Gallus gallus | similar |
| 31 | IRF9 Mesocricetus auratus | 208 | Mesocricetus auratus | belongs |
| 32 | IRF9 Rattus norvegicus | 33 | IRF9 Gallus gallus | similar |
| 32 | IRF9 Rattus norvegicus | 194 | Rattus norvegicus | belongs |

**Table 2 continued from previous page**

| Source ID | Source Name | Target ID | Target Name | Relation |
|---|---|---|---|---|
| 32 | IRF9 Rattus norvegicus | 154 | ORF3b Severe acute respiratory syndrome-related coronavirus | interacts |
| 32 | IRF9 Rattus norvegicus | 170 | ORF6 Severe acute respiratory syndrome-related coronavirus | interacts |
| 32 | IRF9 Rattus norvegicus | 209 | M protein Middle East respiratory syndrome-related coronavirus | interacts |
| 32 | IRF9 Rattus norvegicus | 100 | ORF4a Middle East respiratory syndrome-related coronavirus | interacts |
| 32 | IRF9 Rattus norvegicus | 42 | ORF4b Middle East respiratory syndrome-related coronavirus | interacts |
| 33 | IRF9 Gallus gallus | 201 | Gallus gallus | belongs |
| 34 | RIG-I Homo sapiens | 35 | RIG-I Pan troglodytes | similar |
| 34 | RIG-I Homo sapiens | 36 | RIG-I Macaca mulatta | similar |
| 34 | RIG-I Homo sapiens | 37 | RIG-I Rhinolophus ferrumequinum | similar |
| 34 | RIG-I Homo sapiens | 38 | RIG-I Canis lupus familiaris | similar |
| 34 | RIG-I Homo sapiens | 39 | RIG-I Mesocricetus auratus | similar |
| 34 | RIG-I Homo sapiens | 40 | RIG-I Mus musculus | similar |
| 34 | RIG-I Homo sapiens | 41 | RIG-I Gallus gallus | similar |
| 34 | RIG-I Homo sapiens | 192 | Homo sapiens | belongs |
| 34 | RIG-I Homo sapiens | 154 | ORF3b Severe acute respiratory syndrome-related coronavirus | interacts |
| 34 | RIG-I Homo sapiens | 171 | ORF6 Severe acute respiratory syndrome coronavirus 2 | interacts |
| 34 | RIG-I Homo sapiens | 71 | N protein Severe acute respiratory syndrome-related coronavirus | interacts |
| 35 | RIG-I Pan troglodytes | 36 | RIG-I Macaca mulatta | similar |
| 35 | RIG-I Pan troglodytes | 37 | RIG-I Rhinolophus ferrumequinum | similar |
| 35 | RIG-I Pan troglodytes | 38 | RIG-I Canis lupus familiaris | similar |
| 35 | RIG-I Pan troglodytes | 39 | RIG-I Mesocricetus auratus | similar |
| 35 | RIG-I Pan troglodytes | 40 | RIG-I Mus musculus | similar |
| 35 | RIG-I Pan troglodytes | 41 | RIG-I Gallus gallus | similar |
| 35 | RIG-I Pan troglodytes | 200 | Pan troglodytes | belongs |
| 36 | RIG-I Macaca mulatta | 37 | RIG-I Rhinolophus ferrumequinum | similar |
| 36 | RIG-I Macaca mulatta | 38 | RIG-I Canis lupus familiaris | similar |
| 36 | RIG-I Macaca mulatta | 39 | RIG-I Mesocricetus auratus | similar |
| 36 | RIG-I Macaca mulatta | 40 | RIG-I Mus musculus | similar |
| 36 | RIG-I Macaca mulatta | 41 | RIG-I Gallus gallus | similar |
| 36 | RIG-I Macaca mulatta | 204 | Macaca mulatta | belongs |
| 37 | RIG-I Rhinolophus ferrumequinum | 38 | RIG-I Canis lupus familiaris | similar |
| 37 | RIG-I Rhinolophus ferrumequinum | 39 | RIG-I Mesocricetus auratus | similar |
| 37 | RIG-I Rhinolophus ferrumequinum | 40 | RIG-I Mus musculus | similar |
| 37 | RIG-I Rhinolophus ferrumequinum | 41 | RIG-I Gallus gallus | similar |
| 37 | RIG-I Rhinolophus ferrumequinum | 207 | Rhinolophus ferrumequinum | belongs |
| 37 | RIG-I Rhinolophus ferrumequinum | 154 | ORF3b Severe acute respiratory syndrome-related coronavirus | interacts |
| 37 | RIG-I Rhinolophus ferrumequinum | 71 | N protein Severe acute respiratory syndrome-related coronavirus | interacts |
| 38 | RIG-I Canis lupus familiaris | 39 | RIG-I Mesocricetus auratus | similar |
| 38 | RIG-I Canis lupus familiaris | 40 | RIG-I Mus musculus | similar |
| 38 | RIG-I Canis lupus familiaris | 41 | RIG-I Gallus gallus | similar |
| 38 | RIG-I Canis lupus familiaris | 195 | Canis lupus familiaris | belongs |
| 38 | RIG-I Canis lupus familiaris | 171 | ORF6 Severe acute respiratory syndrome coronavirus 2 | interacts |
| 39 | RIG-I Mesocricetus auratus | 40 | RIG-I Mus musculus | similar |
| 39 | RIG-I Mesocricetus auratus | 41 | RIG-I Gallus gallus | similar |
| 39 | RIG-I Mesocricetus auratus | 208 | Mesocricetus auratus | belongs |
| 39 | RIG-I Mesocricetus auratus | 171 | ORF6 Severe acute respiratory syndrome coronavirus 2 | interacts |
| 40 | RIG-I Mus musculus | 41 | RIG-I Gallus gallus | similar |
| 40 | RIG-I Mus musculus | 193 | Mus musculus | belongs |
| 40 | RIG-I Mus musculus | 154 | ORF3b Severe acute respiratory syndrome-related coronavirus | interacts |
| 40 | RIG-I Mus musculus | 171 | ORF6 Severe acute respiratory syndrome coronavirus 2 | interacts |
| 40 | RIG-I Mus musculus | 71 | N protein Severe acute respiratory syndrome-related coronavirus | interacts |
| 41 | RIG-I Gallus gallus | 201 | Gallus gallus | belongs |
| 42 | ORF4b Middle East respiratory syndrome-related coronavirus | 43 | ORF4b Human coronavirus 229E | similar |
| 42 | ORF4b Middle East respiratory syndrome-related coronavirus | 78 | Middle East respiratory syndrome-related coronavirus | belongs |
| 42 | ORF4b Middle East respiratory syndrome-related coronavirus | 172 | STAT2 Homo sapiens | interacts |
| 42 | ORF4b Middle East respiratory syndrome-related coronavirus | 177 | STAT2 Camelus dromedarius | interacts |
| 42 | ORF4b Middle East respiratory syndrome-related coronavirus | 183 | STAT2 Rattus norvegicus | interacts |
| 42 | ORF4b Middle East respiratory syndrome-related coronavirus | 185 | STAT2 Mus musculus | interacts |
| 42 | ORF4b Middle East respiratory syndrome-related coronavirus | 102 | NF-$\kappa$B Homo sapiens | interacts |
| 42 | ORF4b Middle East respiratory syndrome-related coronavirus | 109 | NF-$\kappa$B Mus musculus | interacts |
| 42 | ORF4b Middle East respiratory syndrome-related coronavirus | 111 | NF-$\kappa$B Rattus norvegicus | interacts |
| 42 | ORF4b Middle East respiratory syndrome-related coronavirus | 114 | NF-$\kappa$B Camelus dromedarius | interacts |
| 42 | ORF4b Middle East respiratory syndrome-related coronavirus | 55 | IRF3 Homo sapiens | interacts |
| 42 | ORF4b Middle East respiratory syndrome-related coronavirus | 60 | IRF3 Camelus dromedarius | interacts |
| 42 | ORF4b Middle East respiratory syndrome-related coronavirus | 64 | IRF3 Rattus norvegicus | interacts |
| 42 | ORF4b Middle East respiratory syndrome-related coronavirus | 66 | IRF3 Mus musculus | interacts |
| 42 | ORF4b Middle East respiratory syndrome-related coronavirus | 115 | IRF7 Homo sapiens | interacts |
| 42 | ORF4b Middle East respiratory syndrome-related coronavirus | 122 | IRF7 Camelus dromedarius | interacts |
| 42 | ORF4b Middle East respiratory syndrome-related coronavirus | 123 | IRF7 Rattus norvegicus | interacts |
| 42 | ORF4b Middle East respiratory syndrome-related coronavirus | 126 | IRF7 Mus musculus | interacts |
| 42 | ORF4b Middle East respiratory syndrome-related coronavirus | 231 | MAVS Homo sapiens | interacts |
| 42 | ORF4b Middle East respiratory syndrome-related coronavirus | 239 | MAVS Camelus dromedarius | interacts |
| 42 | ORF4b Middle East respiratory syndrome-related coronavirus | 243 | MAVS Mus musculus | interacts |
| 42 | ORF4b Middle East respiratory syndrome-related coronavirus | 244 | MAVS Rattus norvegicus | interacts |
| 42 | ORF4b Middle East respiratory syndrome-related coronavirus | 216 | TBK1 Homo sapiens | interacts |
| 42 | ORF4b Middle East respiratory syndrome-related coronavirus | 227 | TBK1 Mus musculus | interacts |
| 42 | ORF4b Middle East respiratory syndrome-related coronavirus | 228 | TBK1 Camelus dromedarius | interacts |
| 42 | ORF4b Middle East respiratory syndrome-related coronavirus | 229 | TBK1 Rattus norvegicus | interacts |
| 43 | ORF4b Human coronavirus 229E | 82 | Human coronavirus 229E | belongs |
| 44 | nsp1 Severe acute respiratory syndrome coronavirus 2 | 45 | nsp1 Severe acute respiratory syndrome-related coronavirus | similar |

**Table 2 continued from previous page**

| Source ID | Source Name | Target ID | Target Name | Relation |
|---|---|---|---|---|
| 44 | nsp1 Severe acute respiratory syndrome coronavirus 2 | 46 | nsp1 Middle East respiratory syndrome-related coronavirus | similar |
| 44 | nsp1 Severe acute respiratory syndrome coronavirus 2 | 47 | nsp1 Human coronavirus HKU1 | similar |
| 44 | nsp1 Severe acute respiratory syndrome coronavirus 2 | 79 | Severe acute respiratory syndrome coronavirus 2 | belongs |
| 45 | nsp1 Severe acute respiratory syndrome-related coronavirus | 46 | nsp1 Middle East respiratory syndrome-related coronavirus | similar |
| 45 | nsp1 Severe acute respiratory syndrome-related coronavirus | 47 | nsp1 Human coronavirus HKU1 | similar |
| 45 | nsp1 Severe acute respiratory syndrome-related coronavirus | 80 | Severe acute respiratory syndrome-related coronavirus | belongs |
| 45 | nsp1 Severe acute respiratory syndrome-related coronavirus | 172 | STAT2 Homo sapiens | interacts |
| 45 | nsp1 Severe acute respiratory syndrome-related coronavirus | 175 | STAT2 Felis catus | interacts |
| 45 | nsp1 Severe acute respiratory syndrome-related coronavirus | 178 | STAT2 Rhinolophus ferrumequinum | interacts |
| 45 | nsp1 Severe acute respiratory syndrome-related coronavirus | 183 | STAT2 Rattus norvegicus | interacts |
| 45 | nsp1 Severe acute respiratory syndrome-related coronavirus | 185 | STAT2 Mus musculus | interacts |
| 46 | nsp1 Middle East respiratory syndrome-related coronavirus | 47 | nsp1 Human coronavirus HKU1 | similar |
| 46 | nsp1 Middle East respiratory syndrome-related coronavirus | 78 | Middle East respiratory syndrome-related coronavirus | belongs |
| 47 | nsp1 Human coronavirus HKU1 | 77 | Human coronavirus HKU1 | belongs |
| 48 | spike Human coronavirus OC43 | 49 | spike Human coronavirus HKU1 | similar |
| 48 | spike Human coronavirus OC43 | 50 | spike Middle East respiratory syndrome-related coronavirus | similar |
| 48 | spike Human coronavirus OC43 | 51 | spike Severe acute respiratory syndrome coronavirus 2 | similar |
| 48 | spike Human coronavirus OC43 | 52 | spike Severe acute respiratory syndrome-related coronavirus | similar |
| 48 | spike Human coronavirus OC43 | 53 | spike Human coronavirus NL63 | similar |
| 48 | spike Human coronavirus OC43 | 54 | spike Human coronavirus 229E | similar |
| 48 | spike Human coronavirus OC43 | 76 | Human coronavirus OC43 | belongs |
| 49 | spike Human coronavirus HKU1 | 50 | spike Middle East respiratory syndrome-related coronavirus | similar |
| 49 | spike Human coronavirus HKU1 | 51 | spike Severe acute respiratory syndrome coronavirus 2 | similar |
| 49 | spike Human coronavirus HKU1 | 52 | spike Severe acute respiratory syndrome-related coronavirus | similar |
| 49 | spike Human coronavirus HKU1 | 53 | spike Human coronavirus NL63 | similar |
| 49 | spike Human coronavirus HKU1 | 54 | spike Human coronavirus 229E | similar |
| 49 | spike Human coronavirus HKU1 | 77 | Human coronavirus HKU1 | belongs |
| 50 | spike Middle East respiratory syndrome-related coronavirus | 51 | spike Severe acute respiratory syndrome coronavirus 2 | similar |
| 50 | spike Middle East respiratory syndrome-related coronavirus | 52 | spike Severe acute respiratory syndrome-related coronavirus | similar |
| 50 | spike Middle East respiratory syndrome-related coronavirus | 53 | spike Human coronavirus NL63 | similar |
| 50 | spike Middle East respiratory syndrome-related coronavirus | 54 | spike Human coronavirus 229E | similar |
| 50 | spike Middle East respiratory syndrome-related coronavirus | 78 | Middle East respiratory syndrome-related coronavirus | belongs |
| 50 | spike Middle East respiratory syndrome-related coronavirus | 156 | DPP4 Homo sapiens | interacts |
| 50 | spike Middle East respiratory syndrome-related coronavirus | 164 | DPP4 Rhinolophus ferrumequinum | interacts |
| 50 | spike Middle East respiratory syndrome-related coronavirus | 168 | DPP4 Camelus dromedarius | interacts |
| 51 | spike Severe acute respiratory syndrome coronavirus 2 | 52 | spike Severe acute respiratory syndrome-related coronavirus | similar |
| 51 | spike Severe acute respiratory syndrome coronavirus 2 | 53 | spike Human coronavirus NL63 | similar |
| 51 | spike Severe acute respiratory syndrome coronavirus 2 | 54 | spike Human coronavirus 229E | similar |
| 51 | spike Severe acute respiratory syndrome coronavirus 2 | 79 | Severe acute respiratory syndrome coronavirus 2 | belongs |
| 51 | spike Severe acute respiratory syndrome coronavirus 2 | 83 | ACE2 Homo sapiens | interacts |
| 51 | spike Severe acute respiratory syndrome coronavirus 2 | 85 | ACE2 Macaca mulatta | interacts |
| 51 | spike Severe acute respiratory syndrome coronavirus 2 | 89 | ACE2 Felis catus | interacts |
| 51 | spike Severe acute respiratory syndrome coronavirus 2 | 91 | ACE2 Mesocricetus auratus | interacts |
| 51 | spike Severe acute respiratory syndrome coronavirus 2 | 97 | ACE2 Rhinolophus ferrumequinum | interacts |
| 51 | spike Severe acute respiratory syndrome coronavirus 2 | 98 | ACE2 Canis lupus familiaris | interacts |
| 52 | spike Severe acute respiratory syndrome-related coronavirus | 53 | spike Human coronavirus NL63 | similar |
| 52 | spike Severe acute respiratory syndrome-related coronavirus | 54 | spike Human coronavirus 229E | similar |
| 52 | spike Severe acute respiratory syndrome-related coronavirus | 80 | Severe acute respiratory syndrome-related coronavirus | belongs |
| 52 | spike Severe acute respiratory syndrome-related coronavirus | 83 | ACE2 Homo sapiens | interacts |
| 52 | spike Severe acute respiratory syndrome-related coronavirus | 89 | ACE2 Felis catus | interacts |
| 52 | spike Severe acute respiratory syndrome-related coronavirus | 97 | ACE2 Rhinolophus ferrumequinum | interacts |
| 53 | spike Human coronavirus NL63 | 54 | spike Human coronavirus 229E | similar |
| 53 | spike Human coronavirus NL63 | 81 | Human coronavirus NL63 | belongs |
| 53 | spike Human coronavirus NL63 | 83 | ACE2 Homo sapiens | interacts |
| 53 | spike Human coronavirus NL63 | 97 | ACE2 Rhinolophus ferrumequinum | interacts |
| 54 | spike Human coronavirus 229E | 82 | Human coronavirus 229E | belongs |
| 55 | IRF3 Homo sapiens | 56 | IRF3 Pan troglodytes | similar |
| 55 | IRF3 Homo sapiens | 57 | IRF3 Macaca mulatta | similar |
| 55 | IRF3 Homo sapiens | 58 | IRF3 Ictidomys tridecemlineatus | similar |
| 55 | IRF3 Homo sapiens | 59 | IRF3 Rhinolophus ferrumequinum | similar |
| 55 | IRF3 Homo sapiens | 60 | IRF3 Camelus dromedarius | similar |
| 55 | IRF3 Homo sapiens | 61 | IRF3 Ovis aries | similar |
| 55 | IRF3 Homo sapiens | 62 | IRF3 Bos taurus | similar |
| 55 | IRF3 Homo sapiens | 63 | IRF3 Equus caballus | similar |
| 55 | IRF3 Homo sapiens | 64 | IRF3 Rattus norvegicus | similar |
| 55 | IRF3 Homo sapiens | 65 | IRF3 Mesocricetus auratus | similar |
| 55 | IRF3 Homo sapiens | 66 | IRF3 Mus musculus | similar |
| 55 | IRF3 Homo sapiens | 67 | IRF3 Canis lupus familiaris | similar |
| 55 | IRF3 Homo sapiens | 68 | IRF3 Gallus gallus | similar |
| 55 | IRF3 Homo sapiens | 192 | Homo sapiens | belongs |
| 55 | IRF3 Homo sapiens | 154 | ORF3b Severe acute respiratory syndrome-related coronavirus | interacts |
| 55 | IRF3 Homo sapiens | 170 | ORF6 Severe acute respiratory syndrome-related coronavirus | interacts |
| 55 | IRF3 Homo sapiens | 171 | ORF6 Severe acute respiratory syndrome coronavirus 2 | interacts |
| 55 | IRF3 Homo sapiens | 188 | PLpro Severe acute respiratory syndrome-related coronavirus | interacts |
| 55 | IRF3 Homo sapiens | 187 | PLpro Middle East respiratory syndrome-related coronavirus | interacts |
| 55 | IRF3 Homo sapiens | 71 | N protein Severe acute respiratory syndrome-related coronavirus | interacts |
| 55 | IRF3 Homo sapiens | 211 | M protein Severe acute respiratory syndrome-related coronavirus | interacts |
| 55 | IRF3 Homo sapiens | 209 | M protein Middle East respiratory syndrome-related coronavirus | interacts |
| 55 | IRF3 Homo sapiens | 100 | ORF4a Middle East respiratory syndrome-related coronavirus | interacts |
| 56 | IRF3 Pan troglodytes | 57 | IRF3 Macaca mulatta | similar |

**Table 2 continued from previous page**

| Source ID | Source Name | Target ID | Target Name | Relation |
|---|---|---|---|---|
| 56 | IRF3 Pan troglodytes | 58 | IRF3 Ictidomys tridecemlineatus | similar |
| 56 | IRF3 Pan troglodytes | 59 | IRF3 Rhinolophus ferrumequinum | similar |
| 56 | IRF3 Pan troglodytes | 60 | IRF3 Camelus dromedarius | similar |
| 56 | IRF3 Pan troglodytes | 61 | IRF3 Ovis aries | similar |
| 56 | IRF3 Pan troglodytes | 62 | IRF3 Bos taurus | similar |
| 56 | IRF3 Pan troglodytes | 63 | IRF3 Equus caballus | similar |
| 56 | IRF3 Pan troglodytes | 64 | IRF3 Rattus norvegicus | similar |
| 56 | IRF3 Pan troglodytes | 65 | IRF3 Mesocricetus auratus | similar |
| 56 | IRF3 Pan troglodytes | 66 | IRF3 Mus musculus | similar |
| 56 | IRF3 Pan troglodytes | 67 | IRF3 Canis lupus familiaris | similar |
| 56 | IRF3 Pan troglodytes | 68 | IRF3 Gallus gallus | similar |
| 56 | IRF3 Pan troglodytes | 200 | Pan troglodytes | belongs |
| 57 | IRF3 Macaca mulatta | 58 | IRF3 Ictidomys tridecemlineatus | similar |
| 57 | IRF3 Macaca mulatta | 59 | IRF3 Rhinolophus ferrumequinum | similar |
| 57 | IRF3 Macaca mulatta | 60 | IRF3 Camelus dromedarius | similar |
| 57 | IRF3 Macaca mulatta | 61 | IRF3 Ovis aries | similar |
| 57 | IRF3 Macaca mulatta | 62 | IRF3 Bos taurus | similar |
| 57 | IRF3 Macaca mulatta | 63 | IRF3 Equus caballus | similar |
| 57 | IRF3 Macaca mulatta | 64 | IRF3 Rattus norvegicus | similar |
| 57 | IRF3 Macaca mulatta | 65 | IRF3 Mesocricetus auratus | similar |
| 57 | IRF3 Macaca mulatta | 66 | IRF3 Mus musculus | similar |
| 57 | IRF3 Macaca mulatta | 67 | IRF3 Canis lupus familiaris | similar |
| 57 | IRF3 Macaca mulatta | 68 | IRF3 Gallus gallus | similar |
| 57 | IRF3 Macaca mulatta | 204 | Macaca mulatta | belongs |
| 58 | IRF3 Ictidomys tridecemlineatus | 59 | IRF3 Rhinolophus ferrumequinum | similar |
| 58 | IRF3 Ictidomys tridecemlineatus | 60 | IRF3 Camelus dromedarius | similar |
| 58 | IRF3 Ictidomys tridecemlineatus | 61 | IRF3 Ovis aries | similar |
| 58 | IRF3 Ictidomys tridecemlineatus | 62 | IRF3 Bos taurus | similar |
| 58 | IRF3 Ictidomys tridecemlineatus | 63 | IRF3 Equus caballus | similar |
| 58 | IRF3 Ictidomys tridecemlineatus | 64 | IRF3 Rattus norvegicus | similar |
| 58 | IRF3 Ictidomys tridecemlineatus | 65 | IRF3 Mesocricetus auratus | similar |
| 58 | IRF3 Ictidomys tridecemlineatus | 66 | IRF3 Mus musculus | similar |
| 58 | IRF3 Ictidomys tridecemlineatus | 67 | IRF3 Canis lupus familiaris | similar |
| 58 | IRF3 Ictidomys tridecemlineatus | 68 | IRF3 Gallus gallus | similar |
| 58 | IRF3 Ictidomys tridecemlineatus | 198 | Ictidomys tridecemlineatus | belongs |
| 59 | IRF3 Rhinolophus ferrumequinum | 60 | IRF3 Camelus dromedarius | similar |
| 59 | IRF3 Rhinolophus ferrumequinum | 61 | IRF3 Ovis aries | similar |
| 59 | IRF3 Rhinolophus ferrumequinum | 62 | IRF3 Bos taurus | similar |
| 59 | IRF3 Rhinolophus ferrumequinum | 63 | IRF3 Equus caballus | similar |
| 59 | IRF3 Rhinolophus ferrumequinum | 64 | IRF3 Rattus norvegicus | similar |
| 59 | IRF3 Rhinolophus ferrumequinum | 65 | IRF3 Mesocricetus auratus | similar |
| 59 | IRF3 Rhinolophus ferrumequinum | 66 | IRF3 Mus musculus | similar |
| 59 | IRF3 Rhinolophus ferrumequinum | 67 | IRF3 Canis lupus familiaris | similar |
| 59 | IRF3 Rhinolophus ferrumequinum | 68 | IRF3 Gallus gallus | similar |
| 59 | IRF3 Rhinolophus ferrumequinum | 207 | Rhinolophus ferrumequinum | belongs |
| 59 | IRF3 Rhinolophus ferrumequinum | 154 | ORF3b Severe acute respiratory syndrome-related coronavirus | interacts |
| 59 | IRF3 Rhinolophus ferrumequinum | 170 | ORF6 Severe acute respiratory syndrome-related coronavirus | interacts |
| 59 | IRF3 Rhinolophus ferrumequinum | 188 | PLpro Severe acute respiratory syndrome-related coronavirus | interacts |
| 59 | IRF3 Rhinolophus ferrumequinum | 71 | N protein Severe acute respiratory syndrome-related coronavirus | interacts |
| 59 | IRF3 Rhinolophus ferrumequinum | 211 | M protein Severe acute respiratory syndrome-related coronavirus | interacts |
| 60 | IRF3 Camelus dromedarius | 61 | IRF3 Ovis aries | similar |
| 60 | IRF3 Camelus dromedarius | 62 | IRF3 Bos taurus | similar |
| 60 | IRF3 Camelus dromedarius | 63 | IRF3 Equus caballus | similar |
| 60 | IRF3 Camelus dromedarius | 64 | IRF3 Rattus norvegicus | similar |
| 60 | IRF3 Camelus dromedarius | 65 | IRF3 Mesocricetus auratus | similar |
| 60 | IRF3 Camelus dromedarius | 66 | IRF3 Mus musculus | similar |
| 60 | IRF3 Camelus dromedarius | 67 | IRF3 Canis lupus familiaris | similar |
| 60 | IRF3 Camelus dromedarius | 68 | IRF3 Gallus gallus | similar |
| 60 | IRF3 Camelus dromedarius | 196 | Camelus dromedarius | belongs |
| 60 | IRF3 Camelus dromedarius | 187 | PLpro Middle East respiratory syndrome-related coronavirus | interacts |
| 60 | IRF3 Camelus dromedarius | 209 | M protein Middle East respiratory syndrome-related coronavirus | interacts |
| 60 | IRF3 Camelus dromedarius | 100 | ORF4a Middle East respiratory syndrome-related coronavirus | interacts |
| 61 | IRF3 Ovis aries | 62 | IRF3 Bos taurus | similar |
| 61 | IRF3 Ovis aries | 63 | IRF3 Equus caballus | similar |
| 61 | IRF3 Ovis aries | 64 | IRF3 Rattus norvegicus | similar |
| 61 | IRF3 Ovis aries | 65 | IRF3 Mesocricetus auratus | similar |
| 61 | IRF3 Ovis aries | 66 | IRF3 Mus musculus | similar |
| 61 | IRF3 Ovis aries | 67 | IRF3 Canis lupus familiaris | similar |
| 61 | IRF3 Ovis aries | 68 | IRF3 Gallus gallus | similar |
| 61 | IRF3 Ovis aries | 205 | Ovis aries | belongs |
| 62 | IRF3 Bos taurus | 63 | IRF3 Equus caballus | similar |
| 62 | IRF3 Bos taurus | 64 | IRF3 Rattus norvegicus | similar |
| 62 | IRF3 Bos taurus | 65 | IRF3 Mesocricetus auratus | similar |
| 62 | IRF3 Bos taurus | 66 | IRF3 Mus musculus | similar |
| 62 | IRF3 Bos taurus | 67 | IRF3 Canis lupus familiaris | similar |
| 62 | IRF3 Bos taurus | 68 | IRF3 Gallus gallus | similar |
| 62 | IRF3 Bos taurus | 199 | Bos taurus | belongs |
| 63 | IRF3 Equus caballus | 64 | IRF3 Rattus norvegicus | similar |
| 63 | IRF3 Equus caballus | 65 | IRF3 Mesocricetus auratus | similar |
| 63 | IRF3 Equus caballus | 66 | IRF3 Mus musculus | similar |

**Table 2 continued from previous page**

| Source ID | Source Name | Target ID | Target Name | Relation |
|---|---|---|---|---|
| 63 | IRF3 Equus caballus | 67 | IRF3 Canis lupus familiaris | similar |
| 63 | IRF3 Equus caballus | 68 | IRF3 Gallus gallus | similar |
| 63 | IRF3 Equus caballus | 203 | Equus caballus | belongs |
| 64 | IRF3 Rattus norvegicus | 65 | IRF3 Mesocricetus auratus | similar |
| 64 | IRF3 Rattus norvegicus | 66 | IRF3 Mus musculus | similar |
| 64 | IRF3 Rattus norvegicus | 67 | IRF3 Canis lupus familiaris | similar |
| 64 | IRF3 Rattus norvegicus | 68 | IRF3 Gallus gallus | similar |
| 64 | IRF3 Rattus norvegicus | 194 | Rattus norvegicus | belongs |
| 64 | IRF3 Rattus norvegicus | 154 | ORF3b Severe acute respiratory syndrome-related coronavirus | interacts |
| 64 | IRF3 Rattus norvegicus | 170 | ORF6 Severe acute respiratory syndrome-related coronavirus | interacts |
| 64 | IRF3 Rattus norvegicus | 171 | ORF6 Severe acute respiratory syndrome coronavirus 2 | interacts |
| 64 | IRF3 Rattus norvegicus | 188 | PLpro Severe acute respiratory syndrome-related coronavirus | interacts |
| 64 | IRF3 Rattus norvegicus | 187 | PLpro Middle East respiratory syndrome-related coronavirus | interacts |
| 64 | IRF3 Rattus norvegicus | 71 | N protein Severe acute respiratory syndrome-related coronavirus | interacts |
| 64 | IRF3 Rattus norvegicus | 211 | M protein Severe acute respiratory syndrome-related coronavirus | interacts |
| 64 | IRF3 Rattus norvegicus | 209 | M protein Middle East respiratory syndrome-related coronavirus | interacts |
| 64 | IRF3 Rattus norvegicus | 100 | ORF4a Middle East respiratory syndrome-related coronavirus | interacts |
| 65 | IRF3 Mesocricetus auratus | 66 | IRF3 Mus musculus | similar |
| 65 | IRF3 Mesocricetus auratus | 67 | IRF3 Canis lupus familiaris | similar |
| 65 | IRF3 Mesocricetus auratus | 68 | IRF3 Gallus gallus | similar |
| 65 | IRF3 Mesocricetus auratus | 208 | Mesocricetus auratus | belongs |
| 65 | IRF3 Mesocricetus auratus | 171 | ORF6 Severe acute respiratory syndrome coronavirus 2 | interacts |
| 66 | IRF3 Mus musculus | 67 | IRF3 Canis lupus familiaris | similar |
| 66 | IRF3 Mus musculus | 68 | IRF3 Gallus gallus | similar |
| 66 | IRF3 Mus musculus | 193 | Mus musculus | belongs |
| 66 | IRF3 Mus musculus | 154 | ORF3b Severe acute respiratory syndrome-related coronavirus | interacts |
| 66 | IRF3 Mus musculus | 170 | ORF6 Severe acute respiratory syndrome-related coronavirus | interacts |
| 66 | IRF3 Mus musculus | 171 | ORF6 Severe acute respiratory syndrome coronavirus 2 | interacts |
| 66 | IRF3 Mus musculus | 188 | PLpro Severe acute respiratory syndrome-related coronavirus | interacts |
| 66 | IRF3 Mus musculus | 187 | PLpro Middle East respiratory syndrome-related coronavirus | interacts |
| 66 | IRF3 Mus musculus | 71 | N protein Severe acute respiratory syndrome-related coronavirus | interacts |
| 66 | IRF3 Mus musculus | 211 | M protein Severe acute respiratory syndrome-related coronavirus | interacts |
| 66 | IRF3 Mus musculus | 209 | M protein Middle East respiratory syndrome-related coronavirus | interacts |
| 66 | IRF3 Mus musculus | 100 | ORF4a Middle East respiratory syndrome-related coronavirus | interacts |
| 67 | IRF3 Canis lupus familiaris | 68 | IRF3 Gallus gallus | similar |
| 67 | IRF3 Canis lupus familiaris | 195 | Canis lupus familiaris | belongs |
| 67 | IRF3 Canis lupus familiaris | 171 | ORF6 Severe acute respiratory syndrome coronavirus 2 | interacts |
| 68 | IRF3 Gallus gallus | 201 | Gallus gallus | belongs |
| 69 | N protein Middle East respiratory syndrome-related coronavirus | 70 | N protein Severe acute respiratory syndrome coronavirus 2 | similar |
| 69 | N protein Middle East respiratory syndrome-related coronavirus | 71 | N protein Severe acute respiratory syndrome-related coronavirus | similar |
| 69 | N protein Middle East respiratory syndrome-related coronavirus | 72 | N protein Human coronavirus HKU1 | similar |
| 69 | N protein Middle East respiratory syndrome-related coronavirus | 73 | N protein Human coronavirus OC43 | similar |
| 69 | N protein Middle East respiratory syndrome-related coronavirus | 74 | N protein Human coronavirus 229E | similar |
| 69 | N protein Middle East respiratory syndrome-related coronavirus | 75 | N protein Human coronavirus NL63 | similar |
| 69 | N protein Middle East respiratory syndrome-related coronavirus | 78 | Middle East respiratory syndrome-related coronavirus | belongs |
| 70 | N protein Severe acute respiratory syndrome coronavirus 2 | 71 | N protein Severe acute respiratory syndrome-related coronavirus | similar |
| 70 | N protein Severe acute respiratory syndrome coronavirus 2 | 72 | N protein Human coronavirus HKU1 | similar |
| 70 | N protein Severe acute respiratory syndrome coronavirus 2 | 73 | N protein Human coronavirus OC43 | similar |
| 70 | N protein Severe acute respiratory syndrome coronavirus 2 | 74 | N protein Human coronavirus 229E | similar |
| 70 | N protein Severe acute respiratory syndrome coronavirus 2 | 75 | N protein Human coronavirus NL63 | similar |
| 70 | N protein Severe acute respiratory syndrome coronavirus 2 | 79 | Severe acute respiratory syndrome coronavirus 2 | belongs |
| 71 | N protein Severe acute respiratory syndrome-related coronavirus | 72 | N protein Human coronavirus HKU1 | similar |
| 71 | N protein Severe acute respiratory syndrome-related coronavirus | 73 | N protein Human coronavirus OC43 | similar |
| 71 | N protein Severe acute respiratory syndrome-related coronavirus | 74 | N protein Human coronavirus 229E | similar |
| 71 | N protein Severe acute respiratory syndrome-related coronavirus | 75 | N protein Human coronavirus NL63 | similar |
| 71 | N protein Severe acute respiratory syndrome-related coronavirus | 80 | Severe acute respiratory syndrome-related coronavirus | belongs |
| 71 | N protein Severe acute respiratory syndrome-related coronavirus | 231 | MAVS Homo sapiens | interacts |
| 71 | N protein Severe acute respiratory syndrome-related coronavirus | 237 | MAVS Rhinolophus ferrumequinum | interacts |
| 71 | N protein Severe acute respiratory syndrome-related coronavirus | 238 | MAVS Felis catus | interacts |
| 71 | N protein Severe acute respiratory syndrome-related coronavirus | 243 | MAVS Mus musculus | interacts |
| 71 | N protein Severe acute respiratory syndrome-related coronavirus | 244 | MAVS Rattus norvegicus | interacts |
| 71 | N protein Severe acute respiratory syndrome-related coronavirus | 129 | MDA5 Homo sapiens | interacts |
| 71 | N protein Severe acute respiratory syndrome-related coronavirus | 134 | MDA5 Felis catus | interacts |
| 71 | N protein Severe acute respiratory syndrome-related coronavirus | 136 | MDA5 Mus musculus | interacts |
| 71 | N protein Severe acute respiratory syndrome-related coronavirus | 137 | MDA5 Rattus norvegicus | interacts |
| 72 | N protein Human coronavirus HKU1 | 73 | N protein Human coronavirus OC43 | similar |
| 72 | N protein Human coronavirus HKU1 | 74 | N protein Human coronavirus 229E | similar |
| 72 | N protein Human coronavirus HKU1 | 75 | N protein Human coronavirus NL63 | similar |
| 72 | N protein Human coronavirus HKU1 | 77 | Human coronavirus HKU1 | belongs |
| 73 | N protein Human coronavirus OC43 | 74 | N protein Human coronavirus 229E | similar |
| 73 | N protein Human coronavirus OC43 | 75 | N protein Human coronavirus NL63 | similar |
| 73 | N protein Human coronavirus OC43 | 76 | Human coronavirus OC43 | belongs |
| 74 | N protein Human coronavirus 229E | 75 | N protein Human coronavirus NL63 | similar |
| 74 | N protein Human coronavirus 229E | 82 | Human coronavirus 229E | belongs |
| 75 | N protein Human coronavirus NL63 | 81 | Human coronavirus NL63 | belongs |
| 76 | Human coronavirus OC43 | 190 | PLpro Human coronavirus OC43 | belongs |
| 76 | Human coronavirus OC43 | 213 | M protein Human coronavirus OC43 | belongs |
| 76 | Human coronavirus OC43 | 192 | Homo sapiens | infects |
| 76 | Human coronavirus OC43 | 193 | Mus musculus | infects |
| 76 | Human coronavirus OC43 | 194 | Rattus norvegicus | infects |

**Table 2 continued from previous page**

| Source ID | Source Name | Target ID | Target Name | Relation |
|---|---|---|---|---|
| 76 | Human coronavirus OC43 | 199 | Bos taurus | infects |
| 77 | Human coronavirus HKU1 | 191 | PLpro Human coronavirus HKU1 | belongs |
| 77 | Human coronavirus HKU1 | 210 | M protein Human coronavirus HKU1 | belongs |
| 77 | Human coronavirus HKU1 | 192 | Homo sapiens | infects |
| 77 | Human coronavirus HKU1 | 193 | Mus musculus | infects |
| 77 | Human coronavirus HKU1 | 194 | Rattus norvegicus | infects |
| 77 | Human coronavirus HKU1 | 207 | Rhinolophus ferrumequinum | infects |
| 78 | Middle East respiratory syndrome-related coronavirus | 100 | ORF4a Middle East respiratory syndrome-related coronavirus | belongs |
| 78 | Middle East respiratory syndrome-related coronavirus | 187 | PLpro Middle East respiratory syndrome-related coronavirus | belongs |
| 78 | Middle East respiratory syndrome-related coronavirus | 209 | M protein Middle East respiratory syndrome-related coronavirus | belongs |
| 78 | Middle East respiratory syndrome-related coronavirus | 192 | Homo sapiens | infects |
| 78 | Middle East respiratory syndrome-related coronavirus | 196 | Camelus dromedarius | infects |
| 78 | Middle East respiratory syndrome-related coronavirus | 207 | Rhinolophus ferrumequinum | infects |
| 79 | Severe acute respiratory syndrome coronavirus 2 | 171 | ORF6 Severe acute respiratory syndrome coronavirus 2 | belongs |
| 79 | Severe acute respiratory syndrome coronavirus 2 | 189 | PLpro Severe acute respiratory syndrome coronavirus 2 | belongs |
| 79 | Severe acute respiratory syndrome coronavirus 2 | 212 | M protein Severe acute respiratory syndrome coronavirus 2 | belongs |
| 79 | Severe acute respiratory syndrome coronavirus 2 | 192 | Homo sapiens | infects |
| 79 | Severe acute respiratory syndrome coronavirus 2 | 195 | Canis lupus familiaris | infects |
| 79 | Severe acute respiratory syndrome coronavirus 2 | 197 | Felis catus | infects |
| 79 | Severe acute respiratory syndrome coronavirus 2 | 204 | Macaca mulatta | infects |
| 79 | Severe acute respiratory syndrome coronavirus 2 | 207 | Rhinolophus ferrumequinum | infects |
| 79 | Severe acute respiratory syndrome coronavirus 2 | 208 | Mesocricetus auratus | infects |
| 80 | Severe acute respiratory syndrome-related coronavirus | 154 | ORF3b Severe acute respiratory syndrome-related coronavirus | belongs |
| 80 | Severe acute respiratory syndrome-related coronavirus | 170 | ORF6 Severe acute respiratory syndrome-related coronavirus | belongs |
| 80 | Severe acute respiratory syndrome-related coronavirus | 188 | PLpro Severe acute respiratory syndrome-related coronavirus | belongs |
| 80 | Severe acute respiratory syndrome-related coronavirus | 211 | M protein Severe acute respiratory syndrome-related coronavirus | belongs |
| 80 | Severe acute respiratory syndrome-related coronavirus | 192 | Homo sapiens | infects |
| 80 | Severe acute respiratory syndrome-related coronavirus | 197 | Felis catus | infects |
| 80 | Severe acute respiratory syndrome-related coronavirus | 207 | Rhinolophus ferrumequinum | infects |
| 81 | Human coronavirus NL63 | 155 | ORF3b Human coronavirus NL63 | belongs |
| 81 | Human coronavirus NL63 | 214 | M protein Human coronavirus NL63 | belongs |
| 81 | Human coronavirus NL63 | 192 | Homo sapiens | infects |
| 81 | Human coronavirus NL63 | 207 | Rhinolophus ferrumequinum | infects |
| 82 | Human coronavirus 229E | 101 | ORF4a Human coronavirus 229E | belongs |
| 82 | Human coronavirus 229E | 215 | M protein Human coronavirus 229E | belongs |
| 82 | Human coronavirus 229E | 192 | Homo sapiens | infects |
| 82 | Human coronavirus 229E | 207 | Rhinolophus ferrumequinum | infects |
| 83 | ACE2 Homo sapiens | 84 | ACE2 Pan troglodytes | similar |
| 83 | ACE2 Homo sapiens | 85 | ACE2 Macaca mulatta | similar |
| 83 | ACE2 Homo sapiens | 86 | ACE2 Ictidomys tridecemlineatus | similar |
| 83 | ACE2 Homo sapiens | 87 | ACE2 Oryctolagus cuniculus | similar |
| 83 | ACE2 Homo sapiens | 88 | ACE2 Equus caballus | similar |
| 83 | ACE2 Homo sapiens | 89 | ACE2 Felis catus | similar |
| 83 | ACE2 Homo sapiens | 90 | ACE2 Camelus dromedarius | similar |
| 83 | ACE2 Homo sapiens | 91 | ACE2 Mesocricetus auratus | similar |
| 83 | ACE2 Homo sapiens | 92 | ACE2 Sus scrofa domesticus | similar |
| 83 | ACE2 Homo sapiens | 93 | ACE2 Ovis aries | similar |
| 83 | ACE2 Homo sapiens | 94 | ACE2 Mus musculus | similar |
| 83 | ACE2 Homo sapiens | 95 | ACE2 Bos taurus | similar |
| 83 | ACE2 Homo sapiens | 96 | ACE2 Rattus norvegicus | similar |
| 83 | ACE2 Homo sapiens | 97 | ACE2 Rhinolophus ferrumequinum | similar |
| 83 | ACE2 Homo sapiens | 98 | ACE2 Canis lupus familiaris | similar |
| 83 | ACE2 Homo sapiens | 99 | ACE2 Gallus gallus | similar |
| 83 | ACE2 Homo sapiens | 192 | Homo sapiens | belongs |
| 84 | ACE2 Pan troglodytes | 85 | ACE2 Macaca mulatta | similar |
| 84 | ACE2 Pan troglodytes | 86 | ACE2 Ictidomys tridecemlineatus | similar |
| 84 | ACE2 Pan troglodytes | 87 | ACE2 Oryctolagus cuniculus | similar |
| 84 | ACE2 Pan troglodytes | 88 | ACE2 Equus caballus | similar |
| 84 | ACE2 Pan troglodytes | 89 | ACE2 Felis catus | similar |
| 84 | ACE2 Pan troglodytes | 90 | ACE2 Camelus dromedarius | similar |
| 84 | ACE2 Pan troglodytes | 91 | ACE2 Mesocricetus auratus | similar |
| 84 | ACE2 Pan troglodytes | 92 | ACE2 Sus scrofa domesticus | similar |
| 84 | ACE2 Pan troglodytes | 93 | ACE2 Ovis aries | similar |
| 84 | ACE2 Pan troglodytes | 94 | ACE2 Mus musculus | similar |
| 84 | ACE2 Pan troglodytes | 95 | ACE2 Bos taurus | similar |
| 84 | ACE2 Pan troglodytes | 96 | ACE2 Rattus norvegicus | similar |
| 84 | ACE2 Pan troglodytes | 97 | ACE2 Rhinolophus ferrumequinum | similar |
| 84 | ACE2 Pan troglodytes | 98 | ACE2 Canis lupus familiaris | similar |
| 84 | ACE2 Pan troglodytes | 99 | ACE2 Gallus gallus | similar |
| 84 | ACE2 Pan troglodytes | 200 | Pan troglodytes | belongs |
| 85 | ACE2 Macaca mulatta | 86 | ACE2 Ictidomys tridecemlineatus | similar |
| 85 | ACE2 Macaca mulatta | 87 | ACE2 Oryctolagus cuniculus | similar |
| 85 | ACE2 Macaca mulatta | 88 | ACE2 Equus caballus | similar |
| 85 | ACE2 Macaca mulatta | 89 | ACE2 Felis catus | similar |
| 85 | ACE2 Macaca mulatta | 90 | ACE2 Camelus dromedarius | similar |
| 85 | ACE2 Macaca mulatta | 91 | ACE2 Mesocricetus auratus | similar |
| 85 | ACE2 Macaca mulatta | 92 | ACE2 Sus scrofa domesticus | similar |
| 85 | ACE2 Macaca mulatta | 93 | ACE2 Ovis aries | similar |
| 85 | ACE2 Macaca mulatta | 94 | ACE2 Mus musculus | similar |
| 85 | ACE2 Macaca mulatta | 95 | ACE2 Bos taurus | similar |

**Table 2 continued from previous page**

| Source ID | Source Name | Target ID | Target Name | Relation |
|---|---|---|---|---|
| 85 | ACE2 Macaca mulatta | 96 | ACE2 Rattus norvegicus | similar |
| 85 | ACE2 Macaca mulatta | 97 | ACE2 Rhinolophus ferrumequinum | similar |
| 85 | ACE2 Macaca mulatta | 98 | ACE2 Canis lupus familiaris | similar |
| 85 | ACE2 Macaca mulatta | 99 | ACE2 Gallus gallus | similar |
| 85 | ACE2 Macaca mulatta | 204 | Macaca mulatta | belongs |
| 86 | ACE2 Ictidomys tridecemlineatus | 87 | ACE2 Oryctolagus cuniculus | similar |
| 86 | ACE2 Ictidomys tridecemlineatus | 88 | ACE2 Equus caballus | similar |
| 86 | ACE2 Ictidomys tridecemlineatus | 89 | ACE2 Felis catus | similar |
| 86 | ACE2 Ictidomys tridecemlineatus | 90 | ACE2 Camelus dromedarius | similar |
| 86 | ACE2 Ictidomys tridecemlineatus | 91 | ACE2 Mesocricetus auratus | similar |
| 86 | ACE2 Ictidomys tridecemlineatus | 92 | ACE2 Sus scrofa domesticus | similar |
| 86 | ACE2 Ictidomys tridecemlineatus | 93 | ACE2 Ovis aries | similar |
| 86 | ACE2 Ictidomys tridecemlineatus | 94 | ACE2 Mus musculus | similar |
| 86 | ACE2 Ictidomys tridecemlineatus | 95 | ACE2 Bos taurus | similar |
| 86 | ACE2 Ictidomys tridecemlineatus | 96 | ACE2 Rattus norvegicus | similar |
| 86 | ACE2 Ictidomys tridecemlineatus | 97 | ACE2 Rhinolophus ferrumequinum | similar |
| 86 | ACE2 Ictidomys tridecemlineatus | 98 | ACE2 Canis lupus familiaris | similar |
| 86 | ACE2 Ictidomys tridecemlineatus | 99 | ACE2 Gallus gallus | similar |
| 86 | ACE2 Ictidomys tridecemlineatus | 198 | Ictidomys tridecemlineatus | belongs |
| 87 | ACE2 Oryctolagus cuniculus | 88 | ACE2 Equus caballus | similar |
| 87 | ACE2 Oryctolagus cuniculus | 89 | ACE2 Felis catus | similar |
| 87 | ACE2 Oryctolagus cuniculus | 90 | ACE2 Camelus dromedarius | similar |
| 87 | ACE2 Oryctolagus cuniculus | 91 | ACE2 Mesocricetus auratus | similar |
| 87 | ACE2 Oryctolagus cuniculus | 92 | ACE2 Sus scrofa domesticus | similar |
| 87 | ACE2 Oryctolagus cuniculus | 93 | ACE2 Ovis aries | similar |
| 87 | ACE2 Oryctolagus cuniculus | 94 | ACE2 Mus musculus | similar |
| 87 | ACE2 Oryctolagus cuniculus | 95 | ACE2 Bos taurus | similar |
| 87 | ACE2 Oryctolagus cuniculus | 96 | ACE2 Rattus norvegicus | similar |
| 87 | ACE2 Oryctolagus cuniculus | 97 | ACE2 Rhinolophus ferrumequinum | similar |
| 87 | ACE2 Oryctolagus cuniculus | 98 | ACE2 Canis lupus familiaris | similar |
| 87 | ACE2 Oryctolagus cuniculus | 99 | ACE2 Gallus gallus | similar |
| 87 | ACE2 Oryctolagus cuniculus | 202 | Oryctolagus cuniculus | belongs |
| 88 | ACE2 Equus caballus | 89 | ACE2 Felis catus | similar |
| 88 | ACE2 Equus caballus | 90 | ACE2 Camelus dromedarius | similar |
| 88 | ACE2 Equus caballus | 91 | ACE2 Mesocricetus auratus | similar |
| 88 | ACE2 Equus caballus | 92 | ACE2 Sus scrofa domesticus | similar |
| 88 | ACE2 Equus caballus | 93 | ACE2 Ovis aries | similar |
| 88 | ACE2 Equus caballus | 94 | ACE2 Mus musculus | similar |
| 88 | ACE2 Equus caballus | 95 | ACE2 Bos taurus | similar |
| 88 | ACE2 Equus caballus | 96 | ACE2 Rattus norvegicus | similar |
| 88 | ACE2 Equus caballus | 97 | ACE2 Rhinolophus ferrumequinum | similar |
| 88 | ACE2 Equus caballus | 98 | ACE2 Canis lupus familiaris | similar |
| 88 | ACE2 Equus caballus | 99 | ACE2 Gallus gallus | similar |
| 88 | ACE2 Equus caballus | 203 | Equus caballus | belongs |
| 89 | ACE2 Felis catus | 90 | ACE2 Camelus dromedarius | similar |
| 89 | ACE2 Felis catus | 91 | ACE2 Mesocricetus auratus | similar |
| 89 | ACE2 Felis catus | 92 | ACE2 Sus scrofa domesticus | similar |
| 89 | ACE2 Felis catus | 93 | ACE2 Ovis aries | similar |
| 89 | ACE2 Felis catus | 94 | ACE2 Mus musculus | similar |
| 89 | ACE2 Felis catus | 95 | ACE2 Bos taurus | similar |
| 89 | ACE2 Felis catus | 96 | ACE2 Rattus norvegicus | similar |
| 89 | ACE2 Felis catus | 97 | ACE2 Rhinolophus ferrumequinum | similar |
| 89 | ACE2 Felis catus | 98 | ACE2 Canis lupus familiaris | similar |
| 89 | ACE2 Felis catus | 99 | ACE2 Gallus gallus | similar |
| 89 | ACE2 Felis catus | 197 | Felis catus | belongs |
| 90 | ACE2 Camelus dromedarius | 91 | ACE2 Mesocricetus auratus | similar |
| 90 | ACE2 Camelus dromedarius | 92 | ACE2 Sus scrofa domesticus | similar |
| 90 | ACE2 Camelus dromedarius | 93 | ACE2 Ovis aries | similar |
| 90 | ACE2 Camelus dromedarius | 94 | ACE2 Mus musculus | similar |
| 90 | ACE2 Camelus dromedarius | 95 | ACE2 Bos taurus | similar |
| 90 | ACE2 Camelus dromedarius | 96 | ACE2 Rattus norvegicus | similar |
| 90 | ACE2 Camelus dromedarius | 97 | ACE2 Rhinolophus ferrumequinum | similar |
| 90 | ACE2 Camelus dromedarius | 98 | ACE2 Canis lupus familiaris | similar |
| 90 | ACE2 Camelus dromedarius | 99 | ACE2 Gallus gallus | similar |
| 90 | ACE2 Camelus dromedarius | 196 | Camelus dromedarius | belongs |
| 91 | ACE2 Mesocricetus auratus | 92 | ACE2 Sus scrofa domesticus | similar |
| 91 | ACE2 Mesocricetus auratus | 93 | ACE2 Ovis aries | similar |
| 91 | ACE2 Mesocricetus auratus | 94 | ACE2 Mus musculus | similar |
| 91 | ACE2 Mesocricetus auratus | 95 | ACE2 Bos taurus | similar |
| 91 | ACE2 Mesocricetus auratus | 96 | ACE2 Rattus norvegicus | similar |
| 91 | ACE2 Mesocricetus auratus | 97 | ACE2 Rhinolophus ferrumequinum | similar |
| 91 | ACE2 Mesocricetus auratus | 98 | ACE2 Canis lupus familiaris | similar |
| 91 | ACE2 Mesocricetus auratus | 99 | ACE2 Gallus gallus | similar |
| 91 | ACE2 Mesocricetus auratus | 208 | Mesocricetus auratus | belongs |
| 92 | ACE2 Sus scrofa domesticus | 93 | ACE2 Ovis aries | similar |
| 92 | ACE2 Sus scrofa domesticus | 94 | ACE2 Mus musculus | similar |
| 92 | ACE2 Sus scrofa domesticus | 95 | ACE2 Bos taurus | similar |
| 92 | ACE2 Sus scrofa domesticus | 96 | ACE2 Rattus norvegicus | similar |
| 92 | ACE2 Sus scrofa domesticus | 97 | ACE2 Rhinolophus ferrumequinum | similar |
| 92 | ACE2 Sus scrofa domesticus | 98 | ACE2 Canis lupus familiaris | similar |

**Table 2 continued from previous page**

| Source ID | Source Name | Target ID | Target Name | Relation |
|---|---|---|---|---|
| 92 | ACE2 Sus scrofa domesticus | 99 | ACE2 Gallus gallus | similar |
| 92 | ACE2 Sus scrofa domesticus | 206 | Sus scrofa domesticus | belongs |
| 93 | ACE2 Ovis aries | 94 | ACE2 Mus musculus | similar |
| 93 | ACE2 Ovis aries | 95 | ACE2 Bos taurus | similar |
| 93 | ACE2 Ovis aries | 96 | ACE2 Rattus norvegicus | similar |
| 93 | ACE2 Ovis aries | 97 | ACE2 Rhinolophus ferrumequinum | similar |
| 93 | ACE2 Ovis aries | 98 | ACE2 Canis lupus familiaris | similar |
| 93 | ACE2 Ovis aries | 99 | ACE2 Gallus gallus | similar |
| 93 | ACE2 Ovis aries | 205 | Ovis aries | belongs |
| 94 | ACE2 Mus musculus | 95 | ACE2 Bos taurus | similar |
| 94 | ACE2 Mus musculus | 96 | ACE2 Rattus norvegicus | similar |
| 94 | ACE2 Mus musculus | 97 | ACE2 Rhinolophus ferrumequinum | similar |
| 94 | ACE2 Mus musculus | 98 | ACE2 Canis lupus familiaris | similar |
| 94 | ACE2 Mus musculus | 99 | ACE2 Gallus gallus | similar |
| 94 | ACE2 Mus musculus | 193 | Mus musculus | belongs |
| 95 | ACE2 Bos taurus | 96 | ACE2 Rattus norvegicus | similar |
| 95 | ACE2 Bos taurus | 97 | ACE2 Rhinolophus ferrumequinum | similar |
| 95 | ACE2 Bos taurus | 98 | ACE2 Canis lupus familiaris | similar |
| 95 | ACE2 Bos taurus | 99 | ACE2 Gallus gallus | similar |
| 95 | ACE2 Bos taurus | 199 | Bos taurus | belongs |
| 96 | ACE2 Rattus norvegicus | 97 | ACE2 Rhinolophus ferrumequinum | similar |
| 96 | ACE2 Rattus norvegicus | 98 | ACE2 Canis lupus familiaris | similar |
| 96 | ACE2 Rattus norvegicus | 99 | ACE2 Gallus gallus | similar |
| 96 | ACE2 Rattus norvegicus | 194 | Rattus norvegicus | belongs |
| 97 | ACE2 Rhinolophus ferrumequinum | 98 | ACE2 Canis lupus familiaris | similar |
| 97 | ACE2 Rhinolophus ferrumequinum | 99 | ACE2 Gallus gallus | similar |
| 97 | ACE2 Rhinolophus ferrumequinum | 207 | Rhinolophus ferrumequinum | belongs |
| 98 | ACE2 Canis lupus familiaris | 99 | ACE2 Gallus gallus | similar |
| 98 | ACE2 Canis lupus familiaris | 195 | Canis lupus familiaris | belongs |
| 99 | ACE2 Gallus gallus | 201 | Gallus gallus | belongs |
| 100 | ORF4a Middle East respiratory syndrome-related coronavirus | 101 | ORF4a Human coronavirus 229E | similar |
| 100 | ORF4a Middle East respiratory syndrome-related coronavirus | 172 | STAT2 Homo sapiens | interacts |
| 100 | ORF4a Middle East respiratory syndrome-related coronavirus | 177 | STAT2 Camelus dromedarius | interacts |
| 100 | ORF4a Middle East respiratory syndrome-related coronavirus | 183 | STAT2 Rattus norvegicus | interacts |
| 100 | ORF4a Middle East respiratory syndrome-related coronavirus | 185 | STAT2 Mus musculus | interacts |
| 100 | ORF4a Middle East respiratory syndrome-related coronavirus | 102 | NF-$\kappa$B Homo sapiens | interacts |
| 100 | ORF4a Middle East respiratory syndrome-related coronavirus | 109 | NF-$\kappa$B Mus musculus | interacts |
| 100 | ORF4a Middle East respiratory syndrome-related coronavirus | 111 | NF-$\kappa$B Rattus norvegicus | interacts |
| 100 | ORF4a Middle East respiratory syndrome-related coronavirus | 114 | NF-$\kappa$B Camelus dromedarius | interacts |
| 100 | ORF4a Middle East respiratory syndrome-related coronavirus | 141 | PRKRA Homo sapiens | interacts |
| 100 | ORF4a Middle East respiratory syndrome-related coronavirus | 145 | PRKRA Camelus dromedarius | interacts |
| 100 | ORF4a Middle East respiratory syndrome-related coronavirus | 147 | PRKRA Mus musculus | interacts |
| 100 | ORF4a Middle East respiratory syndrome-related coronavirus | 151 | PRKRA Rattus norvegicus | interacts |
| 100 | ORF4a Middle East respiratory syndrome-related coronavirus | 129 | MDA5 Homo sapiens | interacts |
| 100 | ORF4a Middle East respiratory syndrome-related coronavirus | 136 | MDA5 Mus musculus | interacts |
| 100 | ORF4a Middle East respiratory syndrome-related coronavirus | 137 | MDA5 Rattus norvegicus | interacts |
| 102 | NF-$\kappa$B Homo sapiens | 103 | NF-$\kappa$B Pan troglodytes | similar |
| 102 | NF-$\kappa$B Homo sapiens | 104 | NF-$\kappa$B Macaca mulatta | similar |
| 102 | NF-$\kappa$B Homo sapiens | 105 | NF-$\kappa$B Bos taurus | similar |
| 102 | NF-$\kappa$B Homo sapiens | 106 | NF-$\kappa$B Rhinolophus ferrumequinum | similar |
| 102 | NF-$\kappa$B Homo sapiens | 107 | NF-$\kappa$B Ovis aries | similar |
| 102 | NF-$\kappa$B Homo sapiens | 108 | NF-$\kappa$B Ictidomys tridecemlineatus | similar |
| 102 | NF-$\kappa$B Homo sapiens | 109 | NF-$\kappa$B Mus musculus | similar |
| 102 | NF-$\kappa$B Homo sapiens | 110 | NF-$\kappa$B Mesocricetus auratus | similar |
| 102 | NF-$\kappa$B Homo sapiens | 111 | NF-$\kappa$B Rattus norvegicus | similar |
| 102 | NF-$\kappa$B Homo sapiens | 112 | NF-$\kappa$B Equus caballus | similar |
| 102 | NF-$\kappa$B Homo sapiens | 113 | NF-$\kappa$B Gallus gallus | similar |
| 102 | NF-$\kappa$B Homo sapiens | 114 | NF-$\kappa$B Camelus dromedarius | similar |
| 102 | NF-$\kappa$B Homo sapiens | 192 | Homo sapiens | belongs |
| 102 | NF-$\kappa$B Homo sapiens | 188 | PLpro Severe acute respiratory syndrome-related coronavirus | interacts |
| 102 | NF-$\kappa$B Homo sapiens | 187 | PLpro Middle East respiratory syndrome-related coronavirus | interacts |
| 102 | NF-$\kappa$B Homo sapiens | 211 | M protein Severe acute respiratory syndrome-related coronavirus | interacts |
| 103 | NF-$\kappa$B Pan troglodytes | 104 | NF-$\kappa$B Macaca mulatta | similar |
| 103 | NF-$\kappa$B Pan troglodytes | 105 | NF-$\kappa$B Bos taurus | similar |
| 103 | NF-$\kappa$B Pan troglodytes | 106 | NF-$\kappa$B Rhinolophus ferrumequinum | similar |
| 103 | NF-$\kappa$B Pan troglodytes | 107 | NF-$\kappa$B Ovis aries | similar |
| 103 | NF-$\kappa$B Pan troglodytes | 108 | NF-$\kappa$B Ictidomys tridecemlineatus | similar |
| 103 | NF-$\kappa$B Pan troglodytes | 109 | NF-$\kappa$B Mus musculus | similar |
| 103 | NF-$\kappa$B Pan troglodytes | 110 | NF-$\kappa$B Mesocricetus auratus | similar |
| 103 | NF-$\kappa$B Pan troglodytes | 111 | NF-$\kappa$B Rattus norvegicus | similar |
| 103 | NF-$\kappa$B Pan troglodytes | 112 | NF-$\kappa$B Equus caballus | similar |
| 103 | NF-$\kappa$B Pan troglodytes | 113 | NF-$\kappa$B Gallus gallus | similar |
| 103 | NF-$\kappa$B Pan troglodytes | 114 | NF-$\kappa$B Camelus dromedarius | similar |
| 103 | NF-$\kappa$B Pan troglodytes | 200 | Pan troglodytes | belongs |
| 104 | NF-$\kappa$B Macaca mulatta | 105 | NF-$\kappa$B Bos taurus | similar |
| 104 | NF-$\kappa$B Macaca mulatta | 106 | NF-$\kappa$B Rhinolophus ferrumequinum | similar |
| 104 | NF-$\kappa$B Macaca mulatta | 107 | NF-$\kappa$B Ovis aries | similar |
| 104 | NF-$\kappa$B Macaca mulatta | 108 | NF-$\kappa$B Ictidomys tridecemlineatus | similar |
| 104 | NF-$\kappa$B Macaca mulatta | 109 | NF-$\kappa$B Mus musculus | similar |
| 104 | NF-$\kappa$B Macaca mulatta | 110 | NF-$\kappa$B Mesocricetus auratus | similar |

**Table 2 continued from previous page**

| Source ID | Source Name | Target ID | Target Name | Relation |
|---|---|---|---|---|
| 104 | NF-$\kappa$B Macaca mulatta | 111 | NF-$\kappa$B Rattus norvegicus | similar |
| 104 | NF-$\kappa$B Macaca mulatta | 112 | NF-$\kappa$B Equus caballus | similar |
| 104 | NF-$\kappa$B Macaca mulatta | 113 | NF-$\kappa$B Gallus gallus | similar |
| 104 | NF-$\kappa$B Macaca mulatta | 114 | NF-$\kappa$B Camelus dromedarius | similar |
| 104 | NF-$\kappa$B Macaca mulatta | 204 | Macaca mulatta | belongs |
| 105 | NF-$\kappa$B Bos taurus | 106 | NF-$\kappa$B Rhinolophus ferrumequinum | similar |
| 105 | NF-$\kappa$B Bos taurus | 107 | NF-$\kappa$B Ovis aries | similar |
| 105 | NF-$\kappa$B Bos taurus | 108 | NF-$\kappa$B Ictidomys tridecemlineatus | similar |
| 105 | NF-$\kappa$B Bos taurus | 109 | NF-$\kappa$B Mus musculus | similar |
| 105 | NF-$\kappa$B Bos taurus | 110 | NF-$\kappa$B Mesocricetus auratus | similar |
| 105 | NF-$\kappa$B Bos taurus | 111 | NF-$\kappa$B Rattus norvegicus | similar |
| 105 | NF-$\kappa$B Bos taurus | 112 | NF-$\kappa$B Equus caballus | similar |
| 105 | NF-$\kappa$B Bos taurus | 113 | NF-$\kappa$B Gallus gallus | similar |
| 105 | NF-$\kappa$B Bos taurus | 114 | NF-$\kappa$B Camelus dromedarius | similar |
| 105 | NF-$\kappa$B Bos taurus | 199 | Bos taurus | belongs |
| 106 | NF-$\kappa$B Rhinolophus ferrumequinum | 107 | NF-$\kappa$B Ovis aries | similar |
| 106 | NF-$\kappa$B Rhinolophus ferrumequinum | 108 | NF-$\kappa$B Ictidomys tridecemlineatus | similar |
| 106 | NF-$\kappa$B Rhinolophus ferrumequinum | 109 | NF-$\kappa$B Mus musculus | similar |
| 106 | NF-$\kappa$B Rhinolophus ferrumequinum | 110 | NF-$\kappa$B Mesocricetus auratus | similar |
| 106 | NF-$\kappa$B Rhinolophus ferrumequinum | 111 | NF-$\kappa$B Rattus norvegicus | similar |
| 106 | NF-$\kappa$B Rhinolophus ferrumequinum | 112 | NF-$\kappa$B Equus caballus | similar |
| 106 | NF-$\kappa$B Rhinolophus ferrumequinum | 113 | NF-$\kappa$B Gallus gallus | similar |
| 106 | NF-$\kappa$B Rhinolophus ferrumequinum | 114 | NF-$\kappa$B Camelus dromedarius | similar |
| 106 | NF-$\kappa$B Rhinolophus ferrumequinum | 207 | Rhinolophus ferrumequinum | belongs |
| 106 | NF-$\kappa$B Rhinolophus ferrumequinum | 188 | PLpro Severe acute respiratory syndrome-related coronavirus | interacts |
| 106 | NF-$\kappa$B Rhinolophus ferrumequinum | 211 | M protein Severe acute respiratory syndrome-related coronavirus | interacts |
| 107 | NF-$\kappa$B Ovis aries | 108 | NF-$\kappa$B Ictidomys tridecemlineatus | similar |
| 107 | NF-$\kappa$B Ovis aries | 109 | NF-$\kappa$B Mus musculus | similar |
| 107 | NF-$\kappa$B Ovis aries | 110 | NF-$\kappa$B Mesocricetus auratus | similar |
| 107 | NF-$\kappa$B Ovis aries | 111 | NF-$\kappa$B Rattus norvegicus | similar |
| 107 | NF-$\kappa$B Ovis aries | 112 | NF-$\kappa$B Equus caballus | similar |
| 107 | NF-$\kappa$B Ovis aries | 113 | NF-$\kappa$B Gallus gallus | similar |
| 107 | NF-$\kappa$B Ovis aries | 114 | NF-$\kappa$B Camelus dromedarius | similar |
| 107 | NF-$\kappa$B Ovis aries | 205 | Ovis aries | belongs |
| 108 | NF-$\kappa$B Ictidomys tridecemlineatus | 109 | NF-$\kappa$B Mus musculus | similar |
| 108 | NF-$\kappa$B Ictidomys tridecemlineatus | 110 | NF-$\kappa$B Mesocricetus auratus | similar |
| 108 | NF-$\kappa$B Ictidomys tridecemlineatus | 111 | NF-$\kappa$B Rattus norvegicus | similar |
| 108 | NF-$\kappa$B Ictidomys tridecemlineatus | 112 | NF-$\kappa$B Equus caballus | similar |
| 108 | NF-$\kappa$B Ictidomys tridecemlineatus | 113 | NF-$\kappa$B Gallus gallus | similar |
| 108 | NF-$\kappa$B Ictidomys tridecemlineatus | 114 | NF-$\kappa$B Camelus dromedarius | similar |
| 108 | NF-$\kappa$B Ictidomys tridecemlineatus | 198 | Ictidomys tridecemlineatus | belongs |
| 109 | NF-$\kappa$B Mus musculus | 110 | NF-$\kappa$B Mesocricetus auratus | similar |
| 109 | NF-$\kappa$B Mus musculus | 111 | NF-$\kappa$B Rattus norvegicus | similar |
| 109 | NF-$\kappa$B Mus musculus | 112 | NF-$\kappa$B Equus caballus | similar |
| 109 | NF-$\kappa$B Mus musculus | 113 | NF-$\kappa$B Gallus gallus | similar |
| 109 | NF-$\kappa$B Mus musculus | 114 | NF-$\kappa$B Camelus dromedarius | similar |
| 109 | NF-$\kappa$B Mus musculus | 193 | Mus musculus | belongs |
| 109 | NF-$\kappa$B Mus musculus | 188 | PLpro Severe acute respiratory syndrome-related coronavirus | interacts |
| 109 | NF-$\kappa$B Mus musculus | 187 | PLpro Middle East respiratory syndrome-related coronavirus | interacts |
| 109 | NF-$\kappa$B Mus musculus | 211 | M protein Severe acute respiratory syndrome-related coronavirus | interacts |
| 110 | NF-$\kappa$B Mesocricetus auratus | 111 | NF-$\kappa$B Rattus norvegicus | similar |
| 110 | NF-$\kappa$B Mesocricetus auratus | 112 | NF-$\kappa$B Equus caballus | similar |
| 110 | NF-$\kappa$B Mesocricetus auratus | 113 | NF-$\kappa$B Gallus gallus | similar |
| 110 | NF-$\kappa$B Mesocricetus auratus | 114 | NF-$\kappa$B Camelus dromedarius | similar |
| 110 | NF-$\kappa$B Mesocricetus auratus | 208 | Mesocricetus auratus | belongs |
| 111 | NF-$\kappa$B Rattus norvegicus | 112 | NF-$\kappa$B Equus caballus | similar |
| 111 | NF-$\kappa$B Rattus norvegicus | 113 | NF-$\kappa$B Gallus gallus | similar |
| 111 | NF-$\kappa$B Rattus norvegicus | 114 | NF-$\kappa$B Camelus dromedarius | similar |
| 111 | NF-$\kappa$B Rattus norvegicus | 194 | Rattus norvegicus | belongs |
| 111 | NF-$\kappa$B Rattus norvegicus | 188 | PLpro Severe acute respiratory syndrome-related coronavirus | interacts |
| 111 | NF-$\kappa$B Rattus norvegicus | 187 | PLpro Middle East respiratory syndrome-related coronavirus | interacts |
| 111 | NF-$\kappa$B Rattus norvegicus | 211 | M protein Severe acute respiratory syndrome-related coronavirus | interacts |
| 112 | NF-$\kappa$B Equus caballus | 113 | NF-$\kappa$B Gallus gallus | similar |
| 112 | NF-$\kappa$B Equus caballus | 114 | NF-$\kappa$B Camelus dromedarius | similar |
| 112 | NF-$\kappa$B Equus caballus | 203 | Equus caballus | belongs |
| 113 | NF-$\kappa$B Gallus gallus | 114 | NF-$\kappa$B Camelus dromedarius | similar |
| 113 | NF-$\kappa$B Gallus gallus | 201 | Gallus gallus | belongs |
| 114 | NF-$\kappa$B Camelus dromedarius | 196 | Camelus dromedarius | belongs |
| 114 | NF-$\kappa$B Camelus dromedarius | 187 | PLpro Middle East respiratory syndrome-related coronavirus | interacts |
| 115 | IRF7 Homo sapiens | 116 | IRF7 Pan troglodytes | similar |
| 115 | IRF7 Homo sapiens | 117 | IRF7 Macaca mulatta | similar |
| 115 | IRF7 Homo sapiens | 118 | IRF7 Ictidomys tridecemlineatus | similar |
| 115 | IRF7 Homo sapiens | 119 | IRF7 Equus caballus | similar |
| 115 | IRF7 Homo sapiens | 120 | IRF7 Rhinolophus ferrumequinum | similar |
| 115 | IRF7 Homo sapiens | 121 | IRF7 Felis catus | similar |
| 115 | IRF7 Homo sapiens | 122 | IRF7 Camelus dromedarius | similar |
| 115 | IRF7 Homo sapiens | 123 | IRF7 Rattus norvegicus | similar |
| 115 | IRF7 Homo sapiens | 124 | IRF7 Mesocricetus auratus | similar |
| 115 | IRF7 Homo sapiens | 125 | IRF7 Ovis aries | similar |
| 115 | IRF7 Homo sapiens | 126 | IRF7 Mus musculus | similar |

**Table 2 continued from previous page**

| Source ID | Source Name | Target ID | Target Name | Relation |
|---|---|---|---|---|
| 115 | IRF7 Homo sapiens | 127 | IRF7 Bos taurus | similar |
| 115 | IRF7 Homo sapiens | 128 | IRF7 Gallus gallus | similar |
| 115 | IRF7 Homo sapiens | 192 | Homo sapiens | belongs |
| 116 | IRF7 Pan troglodytes | 117 | IRF7 Macaca mulatta | similar |
| 116 | IRF7 Pan troglodytes | 118 | IRF7 Ictidomys tridecemlineatus | similar |
| 116 | IRF7 Pan troglodytes | 119 | IRF7 Equus caballus | similar |
| 116 | IRF7 Pan troglodytes | 120 | IRF7 Rhinolophus ferrumequinum | similar |
| 116 | IRF7 Pan troglodytes | 121 | IRF7 Felis catus | similar |
| 116 | IRF7 Pan troglodytes | 122 | IRF7 Camelus dromedarius | similar |
| 116 | IRF7 Pan troglodytes | 123 | IRF7 Rattus norvegicus | similar |
| 116 | IRF7 Pan troglodytes | 124 | IRF7 Mesocricetus auratus | similar |
| 116 | IRF7 Pan troglodytes | 125 | IRF7 Ovis aries | similar |
| 116 | IRF7 Pan troglodytes | 126 | IRF7 Mus musculus | similar |
| 116 | IRF7 Pan troglodytes | 127 | IRF7 Bos taurus | similar |
| 116 | IRF7 Pan troglodytes | 128 | IRF7 Gallus gallus | similar |
| 116 | IRF7 Pan troglodytes | 200 | Pan troglodytes | belongs |
| 117 | IRF7 Macaca mulatta | 118 | IRF7 Ictidomys tridecemlineatus | similar |
| 117 | IRF7 Macaca mulatta | 119 | IRF7 Equus caballus | similar |
| 117 | IRF7 Macaca mulatta | 120 | IRF7 Rhinolophus ferrumequinum | similar |
| 117 | IRF7 Macaca mulatta | 121 | IRF7 Felis catus | similar |
| 117 | IRF7 Macaca mulatta | 122 | IRF7 Camelus dromedarius | similar |
| 117 | IRF7 Macaca mulatta | 123 | IRF7 Rattus norvegicus | similar |
| 117 | IRF7 Macaca mulatta | 124 | IRF7 Mesocricetus auratus | similar |
| 117 | IRF7 Macaca mulatta | 125 | IRF7 Ovis aries | similar |
| 117 | IRF7 Macaca mulatta | 126 | IRF7 Mus musculus | similar |
| 117 | IRF7 Macaca mulatta | 127 | IRF7 Bos taurus | similar |
| 117 | IRF7 Macaca mulatta | 128 | IRF7 Gallus gallus | similar |
| 117 | IRF7 Macaca mulatta | 204 | Macaca mulatta | belongs |
| 118 | IRF7 Ictidomys tridecemlineatus | 119 | IRF7 Equus caballus | similar |
| 118 | IRF7 Ictidomys tridecemlineatus | 120 | IRF7 Rhinolophus ferrumequinum | similar |
| 118 | IRF7 Ictidomys tridecemlineatus | 121 | IRF7 Felis catus | similar |
| 118 | IRF7 Ictidomys tridecemlineatus | 122 | IRF7 Camelus dromedarius | similar |
| 118 | IRF7 Ictidomys tridecemlineatus | 123 | IRF7 Rattus norvegicus | similar |
| 118 | IRF7 Ictidomys tridecemlineatus | 124 | IRF7 Mesocricetus auratus | similar |
| 118 | IRF7 Ictidomys tridecemlineatus | 125 | IRF7 Ovis aries | similar |
| 118 | IRF7 Ictidomys tridecemlineatus | 126 | IRF7 Mus musculus | similar |
| 118 | IRF7 Ictidomys tridecemlineatus | 127 | IRF7 Bos taurus | similar |
| 118 | IRF7 Ictidomys tridecemlineatus | 128 | IRF7 Gallus gallus | similar |
| 118 | IRF7 Ictidomys tridecemlineatus | 198 | Ictidomys tridecemlineatus | belongs |
| 119 | IRF7 Equus caballus | 120 | IRF7 Rhinolophus ferrumequinum | similar |
| 119 | IRF7 Equus caballus | 121 | IRF7 Felis catus | similar |
| 119 | IRF7 Equus caballus | 122 | IRF7 Camelus dromedarius | similar |

### Supplementary Note 8: Predicted Infections Table

Certainty is the probability for the prediction made by IMSP ranging from 0%-100%. Confidence is the computational rule set in IMSP. Strong confidence represents that for a predicted interaction $E_{i,j}$, its two representations $EE_{i,j}$ and $EE_{j,i}$ are all classified into the same class other than the no-interaction class. While for weak confidence interactions, only one representation is classified. Likelihood is the biological rule to validate the predictions. Based on pre-defined filters, the unlikely interactions are predictions that have conflicts with those filters.

| Virus Name | Host Name | Certainty | Confidence | Likelihood |
|---|---|---|---|---|
| Severe acute respiratory syndrome coronavirus 2 | Mus musculus | 100.00% | strong | likely |
| Severe acute respiratory syndrome-related coronavirus | Mesocricetus auratus | 99.98% | strong | likely |
| Human coronavirus NL63 | Rattus norvegicus | 99.96% | strong | likely |
| Severe acute respiratory syndrome coronavirus 2 | Rattus norvegicus | 99.95% | strong | likely |
| Severe acute respiratory syndrome-related coronavirus | Macaca mulatta | 99.94% | strong | likely |
| Severe acute respiratory syndrome-related coronavirus | Canis lupus familiaris | 99.94% | strong | likely |
| Human coronavirus HKU1 | Felis catus | 99.90% | strong | likely |
| Human coronavirus NL63 | Mus musculus | 99.87% | strong | likely |
| Human coronavirus HKU1 | Mesocricetus auratus | 99.86% | strong | likely |
| Human coronavirus NL63 | Felis catus | 99.58% | strong | likely |
| Human coronavirus HKU1 | Canis lupus familiaris | 99.51% | strong | likely |
| Human coronavirus HKU1 | Macaca mulatta | 99.44% | strong | likely |
| Human coronavirus 229E | Mesocricetus auratus | 99.34% | weak | likely |
| Human coronavirus NL63 | Canis lupus familiaris | 92.32% | strong | likely |
| Human coronavirus NL63 | Mesocricetus auratus | 78.57% | strong | likely |
| Severe acute respiratory syndrome coronavirus 2 | Ictidomys tridecemlineatus | 74.07% | weak | likely |
| Human coronavirus NL63 | Macaca mulatta | 66.45% | strong | likely |
| Severe acute respiratory syndrome coronavirus 2 | Bos taurus | 61.00% | weak | likely |
| Severe acute respiratory syndrome coronavirus 2 | Ovis aries | 54.31% | weak | likely |
| Human coronavirus OC43 | Rhinolophus ferrumequinum | 99.99% | strong | unlikely |
| Human coronavirus OC43 | Ovis aries | 99.99% | weak | unlikely |
| Middle East respiratory syndrome-related coronavirus | Rhinolophus ferrumequinum | 99.93% | strong | unlikely |
| Human coronavirus 229E | Rattus norvegicus | 99.78% | strong | unlikely |
| Human coronavirus 229E | Macaca mulatta | 99.70% | weak | unlikely |
| Human coronavirus NL63 | Camelus dromedarius | 99.61% | weak | unlikely |
| Human coronavirus HKU1 | Bos taurus | 99.50% | strong | unlikely |
| Human coronavirus HKU1 | Ovis aries | 99.06% | weak | unlikely |
| Severe acute respiratory syndrome-related coronavirus | Camelus dromedarius | 98.94% | strong | unlikely |
| Human coronavirus HKU1 | Camelus dromedarius | 98.80% | strong | unlikely |
| Human coronavirus 229E | Mus musculus | 98.12% | strong | unlikely |
| Human coronavirus OC43 | Felis catus | 97.23% | strong | unlikely |
| Human coronavirus 229E | Ictidomys tridecemlineatus | 96.83% | weak | unlikely |
| Human coronavirus OC43 | Equus caballus | 96.76% | weak | unlikely |
| Human coronavirus OC43 | Mesocricetus auratus | 96.05% | strong | unlikely |
| Middle East respiratory syndrome-related coronavirus | Sus scrofa domesticus | 94.89% | weak | unlikely |
| Human coronavirus 229E | Canis lupus familiaris | 94.03% | weak | unlikely |
| Severe acute respiratory syndrome-related coronavirus | Bos taurus | 89.91% | strong | unlikely |
| Middle East respiratory syndrome-related coronavirus | Ictidomys tridecemlineatus | 88.31% | weak | unlikely |
| Human coronavirus OC43 | Canis lupus familiaris | 87.41% | strong | unlikely |
| Middle East respiratory syndrome-related coronavirus | Felis catus | 85.28% | strong | unlikely |
| Human coronavirus OC43 | Macaca mulatta | 84.02% | strong | unlikely |
| Human coronavirus OC43 | Sus scrofa domesticus | 82.90% | weak | unlikely |
| Severe acute respiratory syndrome-related coronavirus | Gallus gallus | 81.28% | weak | unlikely |
| Human coronavirus HKU1 | Ictidomys tridecemlineatus | 80.05% | weak | unlikely |
| Human coronavirus OC43 | Camelus dromedarius | 77.19% | strong | unlikely |
| Human coronavirus 229E | Ovis aries | 75.78% | weak | unlikely |
| Severe acute respiratory syndrome-related coronavirus | Sus scrofa domesticus | 75.20% | weak | unlikely |
| Human coronavirus OC43 | Ictidomys tridecemlineatus | 74.53% | weak | unlikely |
| Human coronavirus 229E | Felis catus | 74.30% | weak | unlikely |
| Human coronavirus 229E | Bos taurus | 71.14% | weak | unlikely |
| Severe acute respiratory syndrome coronavirus 2 | Sus scrofa domesticus | 71.09% | weak | unlikely |
| Severe acute respiratory syndrome-related coronavirus | Ovis aries | 68.68% | weak | unlikely |
| Severe acute respiratory syndrome coronavirus 2 | Camelus dromedarius | 68.24% | weak | unlikely |
| Severe acute respiratory syndrome coronavirus 2 | Gallus gallus | 66.89% | weak | unlikely |
| Middle East respiratory syndrome-related coronavirus | Mesocricetus auratus | 65.98% | strong | unlikely |
| Human coronavirus HKU1 | Sus scrofa domesticus | 63.44% | weak | unlikely |
| Severe acute respiratory syndrome coronavirus 2 | Oryctolagus cuniculus | 59.07% | weak | unlikely |
| Human coronavirus 229E | Camelus dromedarius | 59.04% | weak | unlikely |
| Middle East respiratory syndrome-related coronavirus | Gallus gallus | 58.63% | weak | unlikely |
| Human coronavirus NL63 | Bos taurus | 57.98% | weak | unlikely |
| Middle East respiratory syndrome-related coronavirus | Macaca mulatta | 57.54% | strong | unlikely |
| Middle East respiratory syndrome-related coronavirus | Bos taurus | 54.50% | weak | unlikely |
| Middle East respiratory syndrome-related coronavirus | Equus caballus | 54.15% | weak | unlikely |
| Severe acute respiratory syndrome-related coronavirus | Ictidomys tridecemlineatus | 50.98% | weak | unlikely |
| Middle East respiratory syndrome-related coronavirus | Ovis aries | 50.69% | weak | unlikely |

### Supplementary Note 9: Predicted Protein-Protein Interactions Table

| Source Name | Target Name | Certainty | Confidence | Likelihood |
|---|---|---|---|---|
| STAT1 Homo sapiens | PLpro Severe acute respiratory syndrome-related coronavirus | 99.99% | strong | likely |
| STAT2 Homo sapiens | PLpro Severe acute respiratory syndrome-related coronavirus | 99.99% | strong | likely |
| STAT1 Homo sapiens | M protein Severe acute respiratory syndrome-related coronavirus | 99.99% | strong | likely |
| STAT2 Homo sapiens | M protein Severe acute respiratory syndrome-related coronavirus | 99.99% | strong | likely |
| IRF9 Homo sapiens | nsp1 Severe acute respiratory syndrome-related coronavirus | 99.97% | strong | likely |
| nsp15 Severe acute respiratory syndrome-related coronavirus | RIG-I Rhinolophus ferrumequinum | 99.97% | strong | likely |
| IRF9 Homo sapiens | PLpro Severe acute respiratory syndrome-related coronavirus | 99.97% | strong | likely |
| RIG-I Rhinolophus ferrumequinum | ORF6 Severe acute respiratory syndrome-related coronavirus | 99.97% | strong | likely |
| IRF9 Rattus norvegicus | PLpro Severe acute respiratory syndrome-related coronavirus | 99.97% | strong | likely |
| STAT1 Rattus norvegicus | PLpro Severe acute respiratory syndrome-related coronavirus | 99.97% | strong | likely |
| ORF3b Severe acute respiratory syndrome-related coronavirus | MAVS Macaca mulatta | 99.95% | strong | likely |
| IRF9 Homo sapiens | M protein Severe acute respiratory syndrome-related coronavirus | 99.95% | strong | likely |
| IRF3 Rattus norvegicus | ORF6 Severe acute respiratory syndrome coronavirus 2 | 99.94% | strong | likely |
| nsp15 Severe acute respiratory syndrome-related coronavirus | MDA5 Rattus norvegicus | 99.94% | strong | likely |
| IRF3 Macaca mulatta | ORF6 Severe acute respiratory syndrome-related coronavirus | 99.92% | strong | likely |
| IRF9 Mus musculus | PLpro Severe acute respiratory syndrome-related coronavirus | 99.90% | strong | likely |
| nsp15 Severe acute respiratory syndrome coronavirus 2 | RIG-I Mus musculus | 99.90% | strong | likely |
| nsp1 Severe acute respiratory syndrome-related coronavirus | STAT2 Macaca mulatta | 99.90% | strong | likely |
| ORF4a Middle East respiratory syndrome-related coronavirus | IRF7 Rattus norvegicus | 99.90% | strong | likely |
| nsp15 Severe acute respiratory syndrome-related coronavirus | RIG-I Homo sapiens | 99.88% | strong | likely |
| IRF7 Homo sapiens | ORF6 Severe acute respiratory syndrome-related coronavirus | 99.87% | strong | likely |
| nsp15 Severe acute respiratory syndrome-related coronavirus | STAT2 Homo sapiens | 99.87% | strong | likely |
| STAT2 Rattus norvegicus | PLpro Severe acute respiratory syndrome-related coronavirus | 99.87% | strong | likely |
| STAT1 Mus musculus | PLpro Severe acute respiratory syndrome-related coronavirus | 99.87% | strong | likely |
| IRF3 Macaca mulatta | M protein Severe acute respiratory syndrome-related coronavirus | 99.86% | strong | likely |
| NF-kB Ovis aries | PLpro Severe acute respiratory syndrome-related coronavirus | 99.85% | strong | likely |
| ORF6 Severe acute respiratory syndrome-related coronavirus | TBK1 Homo sapiens | 99.85% | strong | likely |
| M protein Severe acute respiratory syndrome-related coronavirus | TBK1 Rattus norvegicus | 99.83% | strong | likely |
| STAT2 Mus musculus | PLpro Severe acute respiratory syndrome-related coronavirus | 99.82% | strong | likely |
| IRF7 Homo sapiens | ORF3b Severe acute respiratory syndrome-related coronavirus | 99.78% | strong | likely |
| nsp15 Severe acute respiratory syndrome-related coronavirus | MAVS Macaca mulatta | 99.77% | strong | likely |
| ORF3b Severe acute respiratory syndrome-related coronavirus | TBK1 Homo sapiens | 99.76% | strong | likely |
| nsp15 Severe acute respiratory syndrome-related coronavirus | IRF9 Rhinolophus ferrumequinum | 99.74% | strong | likely |
| IRF9 Macaca mulatta | ORF3b Severe acute respiratory syndrome-related coronavirus | 99.73% | strong | likely |
| ORF6 Severe acute respiratory syndrome-related coronavirus | MAVS Macaca mulatta | 99.72% | strong | likely |
| NF-kB Camelus dromedarius | M protein Middle East respiratory syndrome-related coronavirus | 99.72% | strong | likely |
| ORF6 Severe acute respiratory syndrome-related coronavirus | TBK1 Rattus norvegicus | 99.71% | strong | likely |
| nsp15 Severe acute respiratory syndrome-related coronavirus | STAT1 Rhinolophus ferrumequinum | 99.68% | strong | likely |
| STAT1 Rattus norvegicus | M protein Severe acute respiratory syndrome-related coronavirus | 99.67% | strong | likely |
| IRF9 Rattus norvegicus | PLpro Middle East respiratory syndrome-related coronavirus | 99.67% | strong | likely |
| nsp1 Severe acute respiratory syndrome coronavirus 2 | STAT2 Felis catus | 99.64% | strong | likely |
| nsp15 Severe acute respiratory syndrome-related coronavirus | IRF3 Mesocricetus auratus | 99.63% | strong | likely |
| STAT2 Rattus norvegicus | M protein Severe acute respiratory syndrome-related coronavirus | 99.62% | strong | likely |
| nsp15 Severe acute respiratory syndrome-related coronavirus | STAT2 Rhinolophus ferrumequinum | 99.61% | strong | likely |
| nsp15 Severe acute respiratory syndrome coronavirus 2 | MAVS Canis lupus familiaris | 99.61% | strong | likely |
| Spike Human coronavirus NL63 | ACE2 Canis lupus familiaris | 99.59% | strong | likely |
| IRF3 Mus musculus | ORF6 Severe acute respiratory syndrome coronavirus 2 | 99.56% | strong | likely |
| M protein Severe acute respiratory syndrome-related coronavirus | TBK1 Mus musculus | 99.54% | strong | likely |
| nsp1 Severe acute respiratory syndrome-related coronavirus | IRF3 Rhinolophus ferrumequinum | 99.53% | strong | likely |
| NF-kB Bos taurus | M protein Severe acute respiratory syndrome-related coronavirus | 99.53% | strong | likely |
| IRF3 Canis lupus familiaris | N protein Severe acute respiratory syndrome-related coronavirus | 99.53% | strong | likely |
| STAT1 Mus musculus | M protein Severe acute respiratory syndrome-related coronavirus | 99.52% | strong | likely |
| RIG-I Macaca mulatta | ORF3b Severe acute respiratory syndrome-related coronavirus | 99.50% | strong | likely |
| IRF3 Macaca mulatta | N protein Severe acute respiratory syndrome-related coronavirus | 99.49% | strong | likely |
| STAT1 Macaca mulatta | nsp1 Severe acute respiratory syndrome-related coronavirus | 99.49% | strong | likely |
| nsp15 Severe acute respiratory syndrome-related coronavirus | IRF3 Canis lupus familiaris | 99.49% | strong | likely |
| ORF6 Severe acute respiratory syndrome coronavirus 2 | STAT2 Rhinolophus ferrumequinum | 99.44% | strong | likely |
| STAT1 Mesocricetus auratus | ORF6 Severe acute respiratory syndrome-related coronavirus | 99.42% | strong | likely |
| ORF4a Middle East respiratory syndrome-related coronavirus | TBK1 Homo sapiens | 99.42% | strong | likely |
| M protein Severe acute respiratory syndrome-related coronavirus | TBK1 Homo sapiens | 99.42% | strong | likely |
| ORF4a Middle East respiratory syndrome-related coronavirus | STAT2 Felis catus | 99.39% | strong | likely |
| MDA5 Rattus norvegicus | PLpro Severe acute respiratory syndrome-related coronavirus | 99.39% | strong | likely |
| IRF9 Mus musculus | PLpro Middle East respiratory syndrome-related coronavirus | 99.39% | strong | likely |
| ORF4b Middle East respiratory syndrome-related coronavirus | PRKRA Rattus norvegicus | 99.38% | strong | likely |
| IRF9 Rattus norvegicus | nsp1 Severe acute respiratory syndrome-related coronavirus | 99.37% | strong | likely |
| IRF3 Macaca mulatta | ORF3b Severe acute respiratory syndrome-related coronavirus | 99.35% | strong | likely |
| ORF6 Severe acute respiratory syndrome coronavirus 2 | MAVS Canis lupus familiaris | 99.34% | strong | likely |
| IRF9 Rattus norvegicus | M protein Severe acute respiratory syndrome-related coronavirus | 99.34% | strong | likely |
| STAT1 Homo sapiens | N protein Severe acute respiratory syndrome-related coronavirus | 99.34% | strong | likely |
| RIG-I Macaca mulatta | ORF6 Severe acute respiratory syndrome-related coronavirus | 99.33% | strong | likely |
| M protein Severe acute respiratory syndrome-related coronavirus | TBK1 Macaca mulatta | 99.33% | strong | likely |
| STAT1 Rhinolophus ferrumequinum | N protein Severe acute respiratory syndrome-related coronavirus | 99.31% | strong | likely |
| STAT2 Mus musculus | M protein Severe acute respiratory syndrome-related coronavirus | 99.28% | strong | likely |
| ORF4a Middle East respiratory syndrome-related coronavirus | MAVS Camelus dromedarius | 99.27% | strong | likely |
| STAT1 Macaca mulatta | ORF6 Severe acute respiratory syndrome-related coronavirus | 99.17% | strong | likely |
| IRF9 Mus musculus | M protein Severe acute respiratory syndrome-related coronavirus | 99.16% | strong | likely |
| IRF9 Rhinolophus ferrumequinum | nsp1 Severe acute respiratory syndrome-related coronavirus | 99.14% | strong | likely |
| N protein Severe acute respiratory syndrome-related coronavirus | STAT2 Homo sapiens | 99.11% | strong | likely |

**Table 4 continued from previous page**

| Source Name | Target Name | Certainty | Confidence | Likelihood |
|---|---|---|---|---|
| IRF3 Camelus dromedarius | PLpro Severe acute respiratory syndrome-related coronavirus | 99.09% | strong | likely |
| PLpro Severe acute respiratory syndrome-related coronavirus | TBK1 Macaca mulatta | 99.09% | strong | likely |
| ORF6 Severe acute respiratory syndrome-related coronavirus | STAT2 Mesocricetus auratus | 99.09% | strong | likely |
| ORF6 Severe acute respiratory syndrome coronavirus 2 | MAVS Homo sapiens | 99.08% | strong | likely |
| STAT1 Rhinolophus ferrumequinum | ORF4a Middle East respiratory syndrome-related coronavirus | 99.08% | strong | likely |
| STAT1 Felis catus | N protein Severe acute respiratory syndrome-related coronavirus | 99.08% | strong | likely |
| MDA5 Homo sapiens | M protein Severe acute respiratory syndrome-related coronavirus | 99.06% | strong | likely |
| IRF3 Macaca mulatta | PLpro Severe acute respiratory syndrome-related coronavirus | 99.06% | strong | likely |
| ORF3b Severe acute respiratory syndrome-related coronavirus | TBK1 Rattus norvegicus | 99.06% | strong | likely |
| IRF9 Felis catus | nsp1 Severe acute respiratory syndrome-related coronavirus | 99.04% | strong | likely |
| nsp15 Severe acute respiratory syndrome-related coronavirus | STAT1 Homo sapiens | 99.01% | strong | likely |
| STAT2 Mus musculus | PLpro Middle East respiratory syndrome-related coronavirus | 98.97% | strong | likely |
| IRF7 Mus musculus | M protein Middle East respiratory syndrome-related coronavirus | 98.93% | strong | likely |
| MDA5 Homo sapiens | PLpro Severe acute respiratory syndrome-related coronavirus | 98.91% | strong | likely |
| nsp15 Severe acute respiratory syndrome-related coronavirus | STAT2 Rattus norvegicus | 98.89% | strong | likely |
| ORF6 Severe acute respiratory syndrome-related coronavirus | STAT2 Camelus dromedarius | 98.89% | strong | likely |
| M protein Severe acute respiratory syndrome-related coronavirus | MAVS Rattus norvegicus | 98.88% | strong | likely |
| RIG-I Gallus gallus | ORF6 Severe acute respiratory syndrome coronavirus 2 | 98.88% | strong | likely |
| ORF4b Middle East respiratory syndrome-related coronavirus | STAT2 Rhinolophus ferrumequinum | 98.87% | strong | likely |
| MDA5 Macaca mulatta | ORF3b Severe acute respiratory syndrome-related coronavirus | 98.86% | strong | likely |
| STAT2 Homo sapiens | PLpro Middle East respiratory syndrome-related coronavirus | 98.83% | strong | likely |
| STAT1 Camelus dromedarius | ORF6 Severe acute respiratory syndrome-related coronavirus | 98.82% | strong | likely |
| Spike Severe acute respiratory syndrome-related coronavirus | ACE2 Mus musculus | 98.80% | strong | likely |
| nsp15 Severe acute respiratory syndrome-related coronavirus | STAT1 Mus musculus | 98.79% | strong | likely |
| ORF6 Severe acute respiratory syndrome-related coronavirus | STAT2 Macaca mulatta | 98.78% | strong | likely |
| Spike Severe acute respiratory syndrome coronavirus 2 | ACE2 Mus musculus | 98.77% | strong | likely |
| STAT1 Homo sapiens | PLpro Middle East respiratory syndrome-related coronavirus | 98.76% | strong | likely |
| Spike Severe acute respiratory syndrome-related coronavirus | ACE2 Rattus norvegicus | 98.76% | strong | likely |
| nsp15 Severe acute respiratory syndrome-related coronavirus | IRF9 Homo sapiens | 98.73% | strong | likely |
| nsp15 Severe acute respiratory syndrome-related coronavirus | IRF3 Macaca mulatta | 98.67% | strong | likely |
| IRF9 Felis catus | N protein Severe acute respiratory syndrome-related coronavirus | 98.62% | strong | likely |
| ORF4b Middle East respiratory syndrome-related coronavirus | PRKRA Homo sapiens | 98.59% | strong | likely |
| ORF4b Middle East respiratory syndrome-related coronavirus | PRKRA Camelus dromedarius | 98.54% | strong | likely |
| STAT1 Camelus dromedarius | PLpro Middle East respiratory syndrome-related coronavirus | 98.51% | strong | likely |
| Spike Severe acute respiratory syndrome-related coronavirus | ACE2 Canis lupus familiaris | 98.49% | strong | likely |
| Spike Severe acute respiratory syndrome-related coronavirus | ACE2 Mesocricetus auratus | 98.49% | strong | likely |
| nsp15 Severe acute respiratory syndrome-related coronavirus | STAT2 Felis catus | 98.48% | strong | likely |
| ORF4a Middle East respiratory syndrome-related coronavirus | TBK1 Rattus norvegicus | 98.44% | strong | likely |
| IRF3 Ovis aries | ORF6 Severe acute respiratory syndrome coronavirus 2 | 98.44% | strong | likely |
| ORF6 Severe acute respiratory syndrome-related coronavirus | MAVS Rattus norvegicus | 98.42% | strong | likely |
| nsp15 Severe acute respiratory syndrome-related coronavirus | IRF9 Felis catus | 98.41% | strong | likely |
| MDA5 Rattus norvegicus | ORF6 Severe acute respiratory syndrome-related coronavirus | 98.36% | strong | likely |
| STAT1 Canis lupus familiaris | ORF6 Severe acute respiratory syndrome-related coronavirus | 98.36% | strong | likely |
| STAT2 Camelus dromedarius | PLpro Middle East respiratory syndrome-related coronavirus | 98.35% | strong | likely |
| NF-kB Homo sapiens | M protein Middle East respiratory syndrome-related coronavirus | 98.31% | strong | likely |
| ORF6 Severe acute respiratory syndrome coronavirus 2 | MAVS Rhinolophus ferrumequinum | 98.30% | strong | likely |
| IRF9 Homo sapiens | PLpro Middle East respiratory syndrome-related coronavirus | 98.27% | strong | likely |
| PRKRA Camelus dromedarius | M protein Middle East respiratory syndrome-related coronavirus | 98.22% | strong | likely |
| ORF4b Middle East respiratory syndrome-related coronavirus | STAT2 Felis catus | 98.19% | strong | likely |
| nsp1 Severe acute respiratory syndrome-related coronavirus | IRF3 Homo sapiens | 98.16% | strong | likely |
| nsp15 Severe acute respiratory syndrome coronavirus 2 | IRF3 Rattus norvegicus | 98.15% | strong | likely |
| N protein Severe acute respiratory syndrome-related coronavirus | MAVS Macaca mulatta | 98.10% | strong | likely |
| nsp15 Severe acute respiratory syndrome-related coronavirus | STAT1 Rattus norvegicus | 98.10% | strong | likely |
| N protein Severe acute respiratory syndrome-related coronavirus | STAT2 Felis catus | 98.03% | strong | likely |
| ORF4a Middle East respiratory syndrome-related coronavirus | STAT2 Rhinolophus ferrumequinum | 98.03% | strong | likely |
| ORF4a Middle East respiratory syndrome-related coronavirus | MAVS Mus musculus | 98.02% | strong | likely |
| Spike Human coronavirus 229E | ACE2 Rhinolophus ferrumequinum | 97.93% | strong | likely |
| NF-kB Rattus norvegicus | ORF3b Severe acute respiratory syndrome-related coronavirus | 97.90% | strong | likely |
| MDA5 Rattus norvegicus | PLpro Middle East respiratory syndrome-related coronavirus | 97.86% | strong | likely |
| RIG-I Mesocricetus auratus | N protein Severe acute respiratory syndrome-related coronavirus | 97.77% | strong | likely |
| IRF3 Rattus norvegicus | N protein Middle East respiratory syndrome-related coronavirus | 97.77% | strong | likely |
| MDA5 Mus musculus | ORF6 Severe acute respiratory syndrome-related coronavirus | 97.71% | strong | likely |
| nsp15 Severe acute respiratory syndrome-related coronavirus | RIG-I Mesocricetus auratus | 97.71% | strong | likely |
| nsp15 Severe acute respiratory syndrome coronavirus 2 | MDA5 Felis catus | 97.71% | strong | likely |
| N protein Severe acute respiratory syndrome-related coronavirus | MDA5 Macaca mulatta | 97.69% | strong | likely |
| ORF4b Middle East respiratory syndrome-related coronavirus | IRF3 Rhinolophus ferrumequinum | 97.69% | strong | likely |
| ORF4a Middle East respiratory syndrome-related coronavirus | TBK1 Rhinolophus ferrumequinum | 97.62% | strong | likely |
| nsp15 Severe acute respiratory syndrome-related coronavirus | RIG-I Mus musculus | 97.58% | strong | likely |
| IRF7 Homo sapiens | M protein Middle East respiratory syndrome-related coronavirus | 97.47% | strong | likely |
| nsp15 Severe acute respiratory syndrome coronavirus 2 | MAVS Rattus norvegicus | 97.46% | strong | likely |
| STAT1 Mus musculus | PLpro Middle East respiratory syndrome-related coronavirus | 97.46% | strong | likely |
| Spike Human coronavirus NL63 | ACE2 Felis catus | 97.45% | strong | likely |
| ACE2 Rhinolophus ferrumequinum | ORF6 Severe acute respiratory syndrome coronavirus 2 | 97.38% | strong | likely |
| PLpro Severe acute respiratory syndrome-related coronavirus | TBK1 Ovis aries | 97.24% | strong | likely |
| IRF3 Mesocricetus auratus | M protein Severe acute respiratory syndrome-related coronavirus | 97.22% | strong | likely |
| ORF6 Severe acute respiratory syndrome coronavirus 2 | MAVS Felis catus | 97.21% | strong | likely |
| NF-kB Rattus norvegicus | M protein Middle East respiratory syndrome-related coronavirus | 97.18% | strong | likely |
| M protein Severe acute respiratory syndrome-related coronavirus | TBK1 Camelus dromedarius | 97.17% | strong | likely |
| Spike Human coronavirus HKU1 | ACE2 Rhinolophus ferrumequinum | 97.11% | strong | likely |
| IRF9 Camelus dromedarius | PLpro Middle East respiratory syndrome-related coronavirus | 96.99% | strong | likely |

**Table 4 continued from previous page**

| Source Name | Target Name | Certainty | Confidence | Likelihood |
|---|---|---|---|---|
| N protein Severe acute respiratory syndrome-related coronavirus | MAVS Canis lupus familiaris | 96.90% | strong | likely |
| MDA5 Rattus norvegicus | M protein Severe acute respiratory syndrome-related coronavirus | 96.86% | strong | likely |
| ORF4a Middle East respiratory syndrome-related coronavirus | IRF7 Camelus dromedarius | 96.76% | strong | likely |
| MDA5 Felis catus | ORF6 Severe acute respiratory syndrome coronavirus 2 | 96.76% | strong | likely |
| STAT1 Macaca mulatta | M protein Severe acute respiratory syndrome-related coronavirus | 96.74% | strong | likely |
| NF-kB Homo sapiens | ORF6 Severe acute respiratory syndrome-related coronavirus | 96.74% | strong | likely |
| ACE2 Canis lupus familiaris | ORF6 Severe acute respiratory syndrome coronavirus 2 | 96.74% | strong | likely |
| NF-kB Ovis aries | M protein Severe acute respiratory syndrome-related coronavirus | 96.64% | strong | likely |
| M protein Severe acute respiratory syndrome-related coronavirus | TBK1 Rhinolophus ferrumequinum | 96.57% | strong | likely |
| nsp15 Severe acute respiratory syndrome-related coronavirus | RIG-I Canis lupus familiaris | 96.52% | strong | likely |
| nsp15 Severe acute respiratory syndrome coronavirus 2 | ACE2 Rhinolophus ferrumequinum | 96.49% | strong | likely |
| nsp15 Severe acute respiratory syndrome-related coronavirus | STAT2 Macaca mulatta | 96.44% | strong | likely |
| STAT1 Felis catus | nsp1 Severe acute respiratory syndrome coronavirus 2 | 96.43% | strong | likely |
| nsp15 Severe acute respiratory syndrome-related coronavirus | IRF9 Rattus norvegicus | 96.36% | strong | likely |
| ORF3b Severe acute respiratory syndrome-related coronavirus | MAVS Camelus dromedarius | 96.34% | strong | likely |
| STAT1 Rattus norvegicus | N protein Severe acute respiratory syndrome-related coronavirus | 96.34% | strong | likely |
| IRF7 Homo sapiens | PLpro Severe acute respiratory syndrome-related coronavirus | 96.29% | strong | likely |
| Spike Severe acute respiratory syndrome coronavirus 2 | ACE2 Rattus norvegicus | 96.29% | strong | likely |
| RIG-I Mus musculus | ORF6 Severe acute respiratory syndrome coronavirus 2 | 96.09% | strong | likely |
| STAT1 Macaca mulatta | ORF6 Severe acute respiratory syndrome coronavirus 2 | 95.99% | strong | likely |
| nsp15 Severe acute respiratory syndrome coronavirus 2 | IRF3 Ovis aries | 95.97% | strong | likely |
| IRF7 Rattus norvegicus | M protein Middle East respiratory syndrome-related coronavirus | 95.91% | strong | likely |
| RIG-I Homo sapiens | ORF6 Severe acute respiratory syndrome-related coronavirus | 95.87% | strong | likely |
| ORF4a Middle East respiratory syndrome-related coronavirus | TBK1 Mus musculus | 95.81% | strong | likely |
| NF-kB Camelus dromedarius | PLpro Severe acute respiratory syndrome-related coronavirus | 95.77% | strong | likely |
| STAT2 Rattus norvegicus | PLpro Middle East respiratory syndrome-related coronavirus | 95.71% | strong | likely |
| ORF6 Severe acute respiratory syndrome-related coronavirus | TBK1 Rhinolophus ferrumequinum | 95.70% | strong | likely |
| IRF3 Ictidomys tridecemlineatus | M protein Severe acute respiratory syndrome-related coronavirus | 95.55% | strong | likely |
| STAT1 Camelus dromedarius | PLpro Severe acute respiratory syndrome-related coronavirus | 95.48% | strong | likely |
| MDA5 Mus musculus | PLpro Severe acute respiratory syndrome-related coronavirus | 95.43% | strong | likely |
| Spike Human coronavirus 229E | ACE2 Homo sapiens | 95.39% | strong | likely |
| IRF3 Mesocricetus auratus | ORF6 Severe acute respiratory syndrome-related coronavirus | 95.34% | strong | likely |
| ORF4b Middle East respiratory syndrome-related coronavirus | MDA5 Mus musculus | 95.32% | strong | likely |
| STAT1 Rattus norvegicus | PLpro Middle East respiratory syndrome-related coronavirus | 95.18% | strong | likely |
| nsp1 Severe acute respiratory syndrome-related coronavirus | TBK1 Rattus norvegicus | 95.13% | strong | likely |
| ORF4b Middle East respiratory syndrome-related coronavirus | TBK1 Rhinolophus ferrumequinum | 95.10% | strong | likely |
| STAT1 Felis catus | ORF4a Middle East respiratory syndrome-related coronavirus | 95.07% | strong | likely |
| MDA5 Macaca mulatta | ORF6 Severe acute respiratory syndrome-related coronavirus | 94.99% | strong | likely |
| NF-kB Rhinolophus ferrumequinum | PLpro Middle East respiratory syndrome-related coronavirus | 94.98% | strong | likely |
| PLpro Severe acute respiratory syndrome-related coronavirus | TBK1 Mesocricetus auratus | 94.87% | strong | likely |
| ORF6 Severe acute respiratory syndrome-related coronavirus | TBK1 Mus musculus | 94.83% | strong | likely |
| N protein Severe acute respiratory syndrome-related coronavirus | STAT2 Rattus norvegicus | 94.74% | strong | likely |
| Spike Severe acute respiratory syndrome-related coronavirus | ACE2 Macaca mulatta | 94.70% | strong | likely |
| ORF4b Middle East respiratory syndrome-related coronavirus | PRKRA Mus musculus | 94.62% | strong | likely |
| NF-kB Homo sapiens | ORF3b Severe acute respiratory syndrome-related coronavirus | 94.58% | strong | likely |
| ORF3b Severe acute respiratory syndrome-related coronavirus | TBK1 Rhinolophus ferrumequinum | 94.57% | strong | likely |
| NF-kB Macaca mulatta | PLpro Severe acute respiratory syndrome-related coronavirus | 94.49% | strong | likely |
| MDA5 Homo sapiens | ORF6 Severe acute respiratory syndrome-related coronavirus | 94.37% | strong | likely |
| STAT2 Camelus dromedarius | M protein Severe acute respiratory syndrome-related coronavirus | 94.31% | strong | likely |
| IRF9 Rattus norvegicus | N protein Severe acute respiratory syndrome-related coronavirus | 94.03% | strong | likely |
| ORF4a Middle East respiratory syndrome-related coronavirus | TBK1 Camelus dromedarius | 93.77% | strong | likely |
| nsp15 Severe acute respiratory syndrome coronavirus 2 | RIG-I Gallus gallus | 93.75% | strong | likely |
| IRF3 Mesocricetus auratus | ORF3b Severe acute respiratory syndrome-related coronavirus | 93.69% | strong | likely |
| M protein Severe acute respiratory syndrome-related coronavirus | TBK1 Bos taurus | 93.59% | strong | likely |
| RIG-I Canis lupus familiaris | ORF6 Severe acute respiratory syndrome-related coronavirus | 93.59% | strong | likely |
| nsp1 Severe acute respiratory syndrome-related coronavirus | TBK1 Homo sapiens | 93.52% | strong | likely |
| NF-kB Mus musculus | M protein Middle East respiratory syndrome-related coronavirus | 93.42% | strong | likely |
| PRKRA Rattus norvegicus | ORF3b Severe acute respiratory syndrome-related coronavirus | 93.36% | strong | likely |
| NF-kB Mus musculus | ORF6 Severe acute respiratory syndrome-related coronavirus | 93.33% | strong | likely |
| nsp1 Severe acute respiratory syndrome-related coronavirus | IRF3 Rattus norvegicus | 93.22% | strong | likely |
| ORF3b Severe acute respiratory syndrome-related coronavirus | STAT2 Macaca mulatta | 93.20% | strong | likely |
| ORF6 Severe acute respiratory syndrome coronavirus 2 | STAT2 Homo sapiens | 93.19% | strong | likely |
| PLpro Middle East respiratory syndrome-related coronavirus | MAVS Rattus norvegicus | 93.18% | strong | likely |
| IRF7 Rattus norvegicus | ORF6 Severe acute respiratory syndrome-related coronavirus | 93.15% | strong | likely |
| N protein Severe acute respiratory syndrome-related coronavirus | STAT2 Rhinolophus ferrumequinum | 93.11% | strong | likely |
| N protein Middle East respiratory syndrome-related coronavirus | MAVS Homo sapiens | 93.09% | strong | likely |
| nsp15 Severe acute respiratory syndrome-related coronavirus | STAT1 Felis catus | 93.09% | strong | likely |
| nsp1 Severe acute respiratory syndrome-related coronavirus | MAVS Homo sapiens | 93.09% | strong | likely |
| IRF9 Homo sapiens | N protein Severe acute respiratory syndrome-related coronavirus | 92.95% | strong | likely |
| M protein Severe acute respiratory syndrome-related coronavirus | MAVS Mus musculus | 92.92% | strong | likely |
| M protein Severe acute respiratory syndrome-related coronavirus | TBK1 Mesocricetus auratus | 92.88% | strong | likely |
| STAT1 Rhinolophus ferrumequinum | ORF4b Middle East respiratory syndrome-related coronavirus | 92.86% | strong | likely |
| nsp15 Severe acute respiratory syndrome-related coronavirus | NF-kB Homo sapiens | 92.80% | strong | likely |
| NF-kB Ictidomys tridecemlineatus | PLpro Severe acute respiratory syndrome-related coronavirus | 92.79% | strong | likely |
| N protein Middle East respiratory syndrome-related coronavirus | MAVS Rattus norvegicus | 92.77% | strong | likely |
| nsp1 Middle East respiratory syndrome-related coronavirus | STAT2 Rattus norvegicus | 92.60% | strong | likely |
| ORF6 Severe acute respiratory syndrome coronavirus 2 | STAT2 Felis catus | 92.51% | strong | likely |
| ORF3b Severe acute respiratory syndrome-related coronavirus | TBK1 Mus musculus | 92.47% | strong | likely |
| STAT2 Rhinolophus ferrumequinum | PLpro Severe acute respiratory syndrome-related coronavirus | 92.17% | strong | likely |
| IRF9 Macaca mulatta | M protein Severe acute respiratory syndrome-related coronavirus | 92.05% | strong | likely |

**Table 4 continued from previous page**

| Source Name | Target Name | Certainty | Confidence | Likelihood |
|---|---|---|---|---|
| ORF6 Severe acute respiratory syndrome-related coronavirus | MAVS Homo sapiens | 91.99% | strong | likely |
| MDA5 Homo sapiens | ORF6 Severe acute respiratory syndrome coronavirus 2 | 91.89% | strong | likely |
| nsp15 Severe acute respiratory syndrome-related coronavirus | STAT1 Macaca mulatta | 91.67% | strong | likely |
| ORF4a Middle East respiratory syndrome-related coronavirus | NF-kB Rhinolophus ferrumequinum | 91.50% | strong | likely |
| MDA5 Felis catus | ORF6 Severe acute respiratory syndrome-related coronavirus | 91.18% | strong | likely |
| nsp15 Severe acute respiratory syndrome-related coronavirus | RIG-I Macaca mulatta | 91.17% | strong | likely |
| M protein Middle East respiratory syndrome-related coronavirus | MAVS Mus musculus | 90.93% | strong | likely |
| STAT1 Rattus norvegicus | nsp1 Middle East respiratory syndrome-related coronavirus | 90.85% | strong | likely |
| IRF7 Rattus norvegicus | PLpro Severe acute respiratory syndrome-related coronavirus | 90.82% | strong | likely |
| IRF3 Camelus dromedarius | ORF3b Severe acute respiratory syndrome-related coronavirus | 90.75% | strong | likely |
| NF-kB Rattus norvegicus | ORF6 Severe acute respiratory syndrome-related coronavirus | 90.61% | strong | likely |
| PLpro Severe acute respiratory syndrome-related coronavirus | MAVS Rattus norvegicus | 90.48% | strong | likely |
| N protein Middle East respiratory syndrome-related coronavirus | MAVS Mus musculus | 90.32% | strong | likely |
| IRF3 Rhinolophus ferrumequinum | ORF4a Middle East respiratory syndrome-related coronavirus | 90.28% | strong | likely |
| STAT1 Camelus dromedarius | M protein Severe acute respiratory syndrome-related coronavirus | 90.16% | strong | likely |
| Spike Human coronavirus NL63 | ACE2 Rattus norvegicus | 90.11% | strong | likely |
| NF-kB Camelus dromedarius | M protein Severe acute respiratory syndrome-related coronavirus | 90.08% | strong | likely |
| PRKRA Rattus norvegicus | PLpro Middle East respiratory syndrome-related coronavirus | 90.06% | strong | likely |
| PLpro Severe acute respiratory syndrome-related coronavirus | TBK1 Camelus dromedarius | 89.92% | strong | likely |
| Spike Human coronavirus HKU1 | ACE2 Mesocricetus auratus | 89.92% | strong | likely |
| ORF6 Severe acute respiratory syndrome-related coronavirus | MAVS Felis catus | 89.65% | strong | likely |
| nsp15 Severe acute respiratory syndrome coronavirus 2 | MAVS Rhinolophus ferrumequinum | 89.59% | strong | likely |
| STAT2 Camelus dromedarius | PLpro Severe acute respiratory syndrome-related coronavirus | 89.54% | strong | likely |
| nsp15 Severe acute respiratory syndrome coronavirus 2 | ACE2 Canis lupus familiaris | 89.47% | strong | likely |
| IRF3 Camelus dromedarius | M protein Severe acute respiratory syndrome-related coronavirus | 89.41% | strong | likely |
| RIG-I Macaca mulatta | N protein Severe acute respiratory syndrome-related coronavirus | 89.40% | strong | likely |
| ORF6 Severe acute respiratory syndrome coronavirus 2 | MAVS Rattus norvegicus | 89.28% | strong | likely |
| IRF9 Rhinolophus ferrumequinum | ORF4a Middle East respiratory syndrome-related coronavirus | 89.26% | strong | likely |
| STAT1 Rhinolophus ferrumequinum | nsp1 Severe acute respiratory syndrome coronavirus 2 | 89.26% | strong | likely |
| MDA5 Mus musculus | PLpro Middle East respiratory syndrome-related coronavirus | 89.26% | strong | likely |
| IRF9 Rhinolophus ferrumequinum | M protein Severe acute respiratory syndrome-related coronavirus | 89.11% | strong | likely |
| nsp1 Severe acute respiratory syndrome-related coronavirus | MDA5 Rattus norvegicus | 89.00% | strong | likely |
| IRF3 Equus caballus | M protein Severe acute respiratory syndrome-related coronavirus | 88.92% | strong | likely |
| ORF4b Middle East respiratory syndrome-related coronavirus | NF-kB Rhinolophus ferrumequinum | 88.87% | strong | likely |
| ORF4a Middle East respiratory syndrome-related coronavirus | IRF7 Mus musculus | 88.84% | strong | likely |
| NF-kB Ictidomys tridecemlineatus | M protein Severe acute respiratory syndrome-related coronavirus | 88.71% | strong | likely |
| NF-kB Rhinolophus ferrumequinum | M protein Middle East respiratory syndrome-related coronavirus | 88.60% | strong | likely |
| nsp15 Severe acute respiratory syndrome coronavirus 2 | STAT2 Felis catus | 88.42% | strong | likely |
| nsp15 Severe acute respiratory syndrome coronavirus 2 | MAVS Homo sapiens | 88.41% | strong | likely |
| RIG-I Pan troglodytes | ORF6 Severe acute respiratory syndrome coronavirus 2 | 88.34% | strong | likely |
| nsp1 Severe acute respiratory syndrome-related coronavirus | MDA5 Mus musculus | 88.22% | strong | likely |
| N protein Severe acute respiratory syndrome-related coronavirus | IRF7 Homo sapiens | 87.97% | strong | likely |
| STAT2 Felis catus | PLpro Severe acute respiratory syndrome-related coronavirus | 87.92% | strong | likely |
| STAT1 Ovis aries | ORF6 Severe acute respiratory syndrome-related coronavirus | 87.88% | strong | likely |
| NF-kB Bos taurus | PLpro Severe acute respiratory syndrome-related coronavirus | 87.81% | strong | likely |
| nsp1 Severe acute respiratory syndrome-related coronavirus | MDA5 Homo sapiens | 87.59% | strong | likely |
| ORF4b Middle East respiratory syndrome-related coronavirus | MDA5 Rattus norvegicus | 87.55% | strong | likely |
| ORF3b Severe acute respiratory syndrome-related coronavirus | STAT2 Canis lupus familiaris | 87.26% | strong | likely |
| MDA5 Rattus norvegicus | ORF6 Severe acute respiratory syndrome coronavirus 2 | 87.00% | strong | likely |
| IRF7 Camelus dromedarius | M protein Middle East respiratory syndrome-related coronavirus | 86.96% | strong | likely |
| MDA5 Mus musculus | M protein Severe acute respiratory syndrome-related coronavirus | 86.83% | strong | likely |
| IRF7 Rattus norvegicus | ORF3b Severe acute respiratory syndrome-related coronavirus | 86.69% | strong | likely |
| M protein Middle East respiratory syndrome-related coronavirus | TBK1 Felis catus | 86.45% | strong | likely |
| NF-kB Equus caballus | M protein Severe acute respiratory syndrome-related coronavirus | 86.30% | strong | likely |
| STAT2 Rhinolophus ferrumequinum | M protein Severe acute respiratory syndrome-related coronavirus | 86.29% | strong | likely |
| nsp15 Severe acute respiratory syndrome-related coronavirus | NF-kB Rhinolophus ferrumequinum | 86.25% | strong | likely |
| ORF4a Middle East respiratory syndrome-related coronavirus | MAVS Rattus norvegicus | 86.13% | strong | likely |
| ORF3b Severe acute respiratory syndrome-related coronavirus | STAT2 Mesocricetus auratus | 85.76% | strong | likely |
| STAT1 Rhinolophus ferrumequinum | M protein Severe acute respiratory syndrome-related coronavirus | 85.68% | strong | likely |
| IRF9 Felis catus | ORF6 Severe acute respiratory syndrome coronavirus 2 | 85.66% | strong | likely |
| STAT1 Mus musculus | N protein Severe acute respiratory syndrome-related coronavirus | 85.55% | strong | likely |
| M protein Middle East respiratory syndrome-related coronavirus | MAVS Homo sapiens | 85.43% | strong | likely |
| ORF6 Severe acute respiratory syndrome-related coronavirus | STAT2 Canis lupus familiaris | 85.42% | strong | likely |
| N protein Severe acute respiratory syndrome coronavirus 2 | MAVS Felis catus | 85.36% | strong | likely |
| ORF4b Middle East respiratory syndrome-related coronavirus | MDA5 Homo sapiens | 85.33% | strong | likely |
| nsp1 Severe acute respiratory syndrome coronavirus 2 | STAT2 Rhinolophus ferrumequinum | 85.26% | strong | likely |
| STAT2 Macaca mulatta | M protein Severe acute respiratory syndrome-related coronavirus | 85.09% | strong | likely |
| STAT1 Macaca mulatta | nsp1 Severe acute respiratory syndrome coronavirus 2 | 85.05% | strong | likely |
| PLpro Middle East respiratory syndrome-related coronavirus | MAVS Mus musculus | 84.96% | strong | likely |
| NF-kB Ovis aries | PLpro Middle East respiratory syndrome-related coronavirus | 84.74% | strong | likely |
| nsp15 Severe acute respiratory syndrome-related coronavirus | TBK1 Rattus norvegicus | 84.60% | strong | likely |
| NF-kB Mesocricetus auratus | PLpro Severe acute respiratory syndrome-related coronavirus | 84.43% | strong | likely |
| PLpro Middle East respiratory syndrome-related coronavirus | MAVS Homo sapiens | 83.94% | strong | likely |
| M protein Middle East respiratory syndrome-related coronavirus | MAVS Rattus norvegicus | 83.87% | strong | likely |
| nsp15 Severe acute respiratory syndrome-related coronavirus | MDA5 Mus musculus | 83.86% | strong | likely |
| STAT1 Felis catus | ORF6 Severe acute respiratory syndrome coronavirus 2 | 83.79% | strong | likely |
| IRF9 Rhinolophus ferrumequinum | ORF4b Middle East respiratory syndrome-related coronavirus | 83.73% | strong | likely |
| STAT1 Mesocricetus auratus | ORF3b Severe acute respiratory syndrome-related coronavirus | 83.53% | strong | likely |
| STAT2 Felis catus | M protein Severe acute respiratory syndrome-related coronavirus | 83.35% | strong | likely |
| IRF7 Mus musculus | ORF3b Severe acute respiratory syndrome-related coronavirus | 83.33% | strong | likely |

**Table 4 continued from previous page**

| Source Name | Target Name | Certainty | Confidence | Likelihood |
|---|---|---|---|---|
| nsp15 Severe acute respiratory syndrome-related coronavirus | NF-kB Rattus norvegicus | 83.04% | strong | likely |
| IRF3 Rhinolophus ferrumequinum | N protein Severe acute respiratory syndrome coronavirus 2 | 82.93% | strong | likely |
| NF-kB Homo sapiens | PLpro Human coronavirus OC43 | 82.69% | strong | likely |
| MDA5 Mus musculus | ORF6 Severe acute respiratory syndrome coronavirus 2 | 82.39% | strong | likely |
| nsp15 Severe acute respiratory syndrome-related coronavirus | IRF9 Mus musculus | 82.33% | strong | likely |
| ORF4a Middle East respiratory syndrome-related coronavirus | TBK1 Felis catus | 82.26% | strong | likely |
| ORF6 Severe acute respiratory syndrome-related coronavirus | MAVS Rhinolophus ferrumequinum | 82.25% | strong | likely |
| PLpro Severe acute respiratory syndrome-related coronavirus | MAVS Mus musculus | 82.18% | strong | likely |
| STAT1 Rhinolophus ferrumequinum | PLpro Severe acute respiratory syndrome-related coronavirus | 82.07% | strong | likely |
| IRF3 Rhinolophus ferrumequinum | PLpro Middle East respiratory syndrome-related coronavirus | 81.85% | strong | likely |
| STAT1 Rhinolophus ferrumequinum | ORF6 Severe acute respiratory syndrome coronavirus 2 | 81.81% | strong | likely |
| MDA5 Rattus norvegicus | M protein Middle East respiratory syndrome-related coronavirus | 81.59% | strong | likely |
| nsp1 Middle East respiratory syndrome-related coronavirus | IRF3 Rattus norvegicus | 81.50% | strong | likely |
| nsp15 Severe acute respiratory syndrome coronavirus 2 | MDA5 Rattus norvegicus | 81.35% | strong | likely |
| STAT1 Homo sapiens | ORF6 Severe acute respiratory syndrome coronavirus 2 | 81.04% | strong | likely |
| RIG-I Mus musculus | ORF6 Severe acute respiratory syndrome-related coronavirus | 80.83% | strong | likely |
| nsp15 Severe acute respiratory syndrome coronavirus 2 | ACE2 Mesocricetus auratus | 80.63% | strong | likely |
| IRF3 Bos taurus | M protein Severe acute respiratory syndrome-related coronavirus | 80.41% | strong | likely |
| IRF3 Camelus dromedarius | N protein Severe acute respiratory syndrome-related coronavirus | 80.39% | strong | likely |
| RIG-I Rhinolophus ferrumequinum | nsp1 Severe acute respiratory syndrome-related coronavirus | 80.23% | strong | likely |
| IRF3 Rhinolophus ferrumequinum | PLpro Severe acute respiratory syndrome coronavirus 2 | 80.23% | strong | likely |
| Spike Human coronavirus HKU1 | ACE2 Felis catus | 80.17% | strong | likely |
| STAT1 Macaca mulatta | ORF3b Severe acute respiratory syndrome-related coronavirus | 80.07% | strong | likely |
| RIG-I Canis lupus familiaris | ORF3b Severe acute respiratory syndrome-related coronavirus | 79.91% | strong | likely |
| ORF4b Middle East respiratory syndrome-related coronavirus | TBK1 Felis catus | 79.88% | strong | likely |
| nsp15 Severe acute respiratory syndrome-related coronavirus | TBK1 Homo sapiens | 79.67% | strong | likely |
| nsp15 Severe acute respiratory syndrome-related coronavirus | MDA5 Felis catus | 79.58% | strong | likely |
| ORF3b Severe acute respiratory syndrome-related coronavirus | MAVS Canis lupus familiaris | 79.39% | strong | likely |
| nsp15 Severe acute respiratory syndrome-related coronavirus | MDA5 Homo sapiens | 79.26% | strong | likely |
| MDA5 Mus musculus | M protein Middle East respiratory syndrome-related coronavirus | 79.17% | strong | likely |
| nsp1 Severe acute respiratory syndrome-related coronavirus | MAVS Rattus norvegicus | 79.08% | strong | likely |
| IRF7 Homo sapiens | PLpro Middle East respiratory syndrome-related coronavirus | 79.07% | strong | likely |
| ORF4a Middle East respiratory syndrome-related coronavirus | IRF7 Homo sapiens | 78.82% | strong | likely |
| M protein Severe acute respiratory syndrome-related coronavirus | TBK1 Ovis aries | 78.72% | strong | likely |
| IRF7 Mus musculus | PLpro Middle East respiratory syndrome-related coronavirus | 78.66% | strong | likely |
| ORF6 Severe acute respiratory syndrome-related coronavirus | STAT2 Ovis aries | 78.53% | strong | likely |
| M protein Middle East respiratory syndrome-related coronavirus | TBK1 Rhinolophus ferrumequinum | 78.44% | strong | likely |
| ACE2 Felis catus | ORF6 Severe acute respiratory syndrome coronavirus 2 | 78.20% | strong | likely |
| Spike Human coronavirus HKU1 | ACE2 Macaca mulatta | 78.05% | strong | likely |
| IRF3 Rhinolophus ferrumequinum | M protein Middle East respiratory syndrome-related coronavirus | 77.98% | strong | likely |
| IRF9 Camelus dromedarius | PLpro Severe acute respiratory syndrome-related coronavirus | 77.94% | strong | likely |
| nsp15 Severe acute respiratory syndrome coronavirus 2 | ACE2 Felis catus | 77.76% | strong | likely |
| Spike Human coronavirus HKU1 | ACE2 Mus musculus | 77.74% | strong | likely |
| IRF9 Mesocricetus auratus | ORF6 Severe acute respiratory syndrome-related coronavirus | 77.61% | strong | likely |
| nsp15 Severe acute respiratory syndrome-related coronavirus | MAVS Canis lupus familiaris | 77.39% | strong | likely |
| nsp15 Severe acute respiratory syndrome coronavirus 2 | IRF3 Mus musculus | 77.38% | strong | likely |
| nsp1 Severe acute respiratory syndrome coronavirus 2 | STAT2 Macaca mulatta | 77.21% | strong | likely |
| nsp15 Severe acute respiratory syndrome-related coronavirus | STAT2 Mus musculus | 77.17% | strong | likely |
| nsp1 Severe acute respiratory syndrome-related coronavirus | MAVS Rhinolophus ferrumequinum | 77.10% | strong | likely |
| nsp15 Severe acute respiratory syndrome coronavirus 2 | MDA5 Homo sapiens | 77.01% | strong | likely |
| NF-kB Mus musculus | ORF3b Severe acute respiratory syndrome-related coronavirus | 76.93% | strong | likely |
| nsp15 Severe acute respiratory syndrome-related coronavirus | IRF7 Homo sapiens | 76.92% | strong | likely |
| STAT1 Macaca mulatta | PLpro Severe acute respiratory syndrome-related coronavirus | 76.90% | strong | likely |
| nsp1 Severe acute respiratory syndrome-related coronavirus | IRF3 Mus musculus | 76.63% | strong | likely |
| ORF4a Middle East respiratory syndrome-related coronavirus | MAVS Homo sapiens | 76.32% | strong | likely |
| IRF3 Mesocricetus auratus | PLpro Severe acute respiratory syndrome-related coronavirus | 76.30% | strong | likely |
| ORF6 Severe acute respiratory syndrome coronavirus 2 | MAVS Mus musculus | 76.20% | strong | likely |
| PRKRA Homo sapiens | ORF3b Severe acute respiratory syndrome-related coronavirus | 76.12% | strong | likely |
| RIG-I Rhinolophus ferrumequinum | M protein Severe acute respiratory syndrome-related coronavirus | 75.97% | strong | likely |
| NF-kB Mesocricetus auratus | M protein Severe acute respiratory syndrome-related coronavirus | 75.83% | strong | likely |
| PRKRA Rattus norvegicus | M protein Middle East respiratory syndrome-related coronavirus | 75.71% | strong | likely |
| IRF3 Canis lupus familiaris | ORF3b Severe acute respiratory syndrome-related coronavirus | 75.33% | strong | likely |
| PLpro Middle East respiratory syndrome-related coronavirus | TBK1 Felis catus | 75.30% | strong | likely |
| IRF3 Homo sapiens | N protein Middle East respiratory syndrome-related coronavirus | 75.28% | strong | likely |
| IRF7 Mus musculus | PLpro Severe acute respiratory syndrome-related coronavirus | 75.23% | strong | likely |
| IRF9 Mus musculus | nsp1 Severe acute respiratory syndrome-related coronavirus | 75.21% | strong | likely |
| Spike Severe acute respiratory syndrome coronavirus 2 | ACE2 Equus caballus | 75.15% | strong | likely |
| M protein Severe acute respiratory syndrome-related coronavirus | MAVS Homo sapiens | 75.14% | strong | likely |
| ORF4b Middle East respiratory syndrome-related coronavirus | MAVS Rhinolophus ferrumequinum | 74.84% | strong | likely |
| ORF6 Severe acute respiratory syndrome-related coronavirus | MAVS Mus musculus | 74.39% | strong | likely |
| Spike Severe acute respiratory syndrome coronavirus 2 | ACE2 Ictidomys tridecemlineatus | 74.37% | strong | likely |
| MDA5 Homo sapiens | PLpro Middle East respiratory syndrome-related coronavirus | 74.36% | strong | likely |
| nsp1 Severe acute respiratory syndrome-related coronavirus | TBK1 Mus musculus | 74.34% | strong | likely |
| IRF9 Rhinolophus ferrumequinum | ORF6 Severe acute respiratory syndrome coronavirus 2 | 74.30% | strong | likely |
| IRF7 Mus musculus | ORF6 Severe acute respiratory syndrome-related coronavirus | 73.98% | strong | likely |
| Spike Severe acute respiratory syndrome coronavirus 2 | ACE2 Ovis aries | 73.87% | strong | likely |
| PLpro Severe acute respiratory syndrome-related coronavirus | MAVS Homo sapiens | 73.76% | strong | likely |
| nsp15 Severe acute respiratory syndrome-related coronavirus | TBK1 Rhinolophus ferrumequinum | 73.63% | strong | likely |
| Spike Severe acute respiratory syndrome coronavirus 2 | ACE2 Bos taurus | 73.57% | strong | likely |
| nsp1 Middle East respiratory syndrome-related coronavirus | STAT2 Homo sapiens | 73.24% | strong | likely |

**Table 4 continued from previous page**

| Source Name | Target Name | Certainty | Confidence | Likelihood |
|---|---|---|---|---|
| IRF3 Gallus gallus | ORF6 Severe acute respiratory syndrome coronavirus 2 | 73.10% | strong | likely |
| STAT2 Felis catus | M protein Middle East respiratory syndrome-related coronavirus | 72.87% | strong | likely |
| RIG-I Rhinolophus ferrumequinum | N protein Severe acute respiratory syndrome coronavirus 2 | 72.85% | strong | likely |
| nsp1 Severe acute respiratory syndrome-related coronavirus | MDA5 Felis catus | 72.74% | strong | likely |
| MDA5 Macaca mulatta | ORF6 Severe acute respiratory syndrome coronavirus 2 | 72.71% | strong | likely |
| PLpro Severe acute respiratory syndrome-related coronavirus | TBK1 Bos taurus | 72.71% | strong | likely |
| nsp1 Severe acute respiratory syndrome-related coronavirus | STAT2 Mesocricetus auratus | 72.58% | strong | likely |
| NF-kB Homo sapiens | PLpro Severe acute respiratory syndrome coronavirus 2 | 72.37% | strong | likely |
| N protein Severe acute respiratory syndrome-related coronavirus | MAVS Camelus dromedarius | 72.37% | strong | likely |
| nsp15 Severe acute respiratory syndrome coronavirus 2 | MAVS Mesocricetus auratus | 72.27% | strong | likely |
| IRF7 Rattus norvegicus | PLpro Middle East respiratory syndrome-related coronavirus | 72.27% | strong | likely |
| Spike Human coronavirus NL63 | ACE2 Mus musculus | 72.04% | strong | likely |
| STAT1 Canis lupus familiaris | ORF3b Severe acute respiratory syndrome-related coronavirus | 71.87% | strong | likely |
| NF-kB Camelus dromedarius | ORF6 Severe acute respiratory syndrome-related coronavirus | 71.67% | strong | likely |
| Spike Human coronavirus NL63 | ACE2 Macaca mulatta | 71.62% | strong | likely |
| ORF4a Middle East respiratory syndrome-related coronavirus | STAT2 Ovis aries | 70.64% | strong | likely |
| nsp15 Severe acute respiratory syndrome-related coronavirus | TBK1 Mus musculus | 70.57% | strong | likely |
| nsp15 Severe acute respiratory syndrome-related coronavirus | IRF9 Macaca mulatta | 70.52% | strong | likely |
| Spike Severe acute respiratory syndrome coronavirus 2 | ACE2 Pan troglodytes | 70.44% | strong | likely |
| IRF9 Rhinolophus ferrumequinum | PLpro Severe acute respiratory syndrome-related coronavirus | 69.79% | strong | likely |
| STAT1 Camelus dromedarius | ORF3b Severe acute respiratory syndrome-related coronavirus | 69.79% | strong | likely |
| nsp15 Severe acute respiratory syndrome-related coronavirus | NF-kB Mus musculus | 69.20% | strong | likely |
| M protein Severe acute respiratory syndrome-related coronavirus | TBK1 Felis catus | 69.03% | strong | likely |
| IRF9 Camelus dromedarius | ORF6 Severe acute respiratory syndrome-related coronavirus | 68.99% | strong | likely |
| NF-kB Mesocricetus auratus | PLpro Middle East respiratory syndrome-related coronavirus | 68.80% | strong | likely |
| Spike Human coronavirus 229E | ACE2 Mesocricetus auratus | 68.10% | strong | likely |
| nsp15 Severe acute respiratory syndrome coronavirus 2 | MAVS Felis catus | 67.96% | strong | likely |
| nsp15 Severe acute respiratory syndrome-related coronavirus | MAVS Mesocricetus auratus | 67.88% | strong | likely |
| M protein Severe acute respiratory syndrome-related coronavirus | MAVS Rhinolophus ferrumequinum | 67.75% | strong | likely |
| PRKRA Mus musculus | ORF3b Severe acute respiratory syndrome-related coronavirus | 66.96% | strong | likely |
| IRF7 Rhinolophus ferrumequinum | ORF6 Severe acute respiratory syndrome-related coronavirus | 66.71% | strong | likely |
| IRF9 Felis catus | ORF4a Middle East respiratory syndrome-related coronavirus | 66.59% | strong | likely |
| PLpro Middle East respiratory syndrome-related coronavirus | TBK1 Rhinolophus ferrumequinum | 65.96% | strong | likely |
| IRF9 Macaca mulatta | ORF6 Severe acute respiratory syndrome-related coronavirus | 65.85% | strong | likely |
| NF-kB Gallus gallus | M protein Severe acute respiratory syndrome-related coronavirus | 65.68% | strong | likely |
| N protein Severe acute respiratory syndrome-related coronavirus | MDA5 Canis lupus familiaris | 65.52% | strong | likely |
| NF-kB Macaca mulatta | M protein Severe acute respiratory syndrome-related coronavirus | 65.30% | strong | likely |
| IRF9 Ovis aries | ORF4b Middle East respiratory syndrome-related coronavirus | 64.84% | strong | likely |
| PLpro Severe acute respiratory syndrome-related coronavirus | MAVS Rhinolophus ferrumequinum | 64.08% | strong | likely |
| Spike Human coronavirus HKU1 | ACE2 Canis lupus familiaris | 63.55% | strong | likely |
| PRKRA Rattus norvegicus | PLpro Severe acute respiratory syndrome-related coronavirus | 62.76% | strong | likely |
| nsp15 Severe acute respiratory syndrome coronavirus 2 | STAT1 Macaca mulatta | 62.75% | strong | likely |
| PRKRA Camelus dromedarius | PLpro Middle East respiratory syndrome-related coronavirus | 62.50% | strong | likely |
| RIG-I Mesocricetus auratus | ORF3b Severe acute respiratory syndrome-related coronavirus | 61.87% | strong | likely |
| nsp15 Severe acute respiratory syndrome coronavirus 2 | RIG-I Pan troglodytes | 61.48% | strong | likely |
| IRF9 Felis catus | ORF4b Middle East respiratory syndrome-related coronavirus | 60.90% | strong | likely |
| IRF9 Rattus norvegicus | nsp1 Middle East respiratory syndrome-related coronavirus | 60.20% | strong | likely |
| IRF3 Homo sapiens | PLpro Severe acute respiratory syndrome coronavirus 2 | 59.38% | strong | likely |
| IRF3 Mesocricetus auratus | N protein Severe acute respiratory syndrome-related coronavirus | 59.38% | strong | likely |
| Spike Human coronavirus NL63 | ACE2 Mesocricetus auratus | 59.37% | strong | likely |
| PLpro Middle East respiratory syndrome-related coronavirus | MAVS Camelus dromedarius | 58.43% | strong | likely |
| RIG-I Canis lupus familiaris | N protein Severe acute respiratory syndrome-related coronavirus | 56.60% | strong | likely |
| IRF3 Camelus dromedarius | ORF6 Severe acute respiratory syndrome-related coronavirus | 56.57% | strong | likely |
| PRKRA Homo sapiens | ORF6 Severe acute respiratory syndrome-related coronavirus | 56.14% | strong | likely |
| nsp1 Severe acute respiratory syndrome coronavirus 2 | STAT2 Mesocricetus auratus | 55.35% | strong | likely |
| nsp15 Severe acute respiratory syndrome coronavirus 2 | MAVS Mus musculus | 54.79% | strong | likely |
| Spike Severe acute respiratory syndrome coronavirus 2 | IRF3 Rhinolophus ferrumequinum | 89.09% | strong | unlikely |
| Spike Severe acute respiratory syndrome-related coronavirus | IRF3 Rhinolophus ferrumequinum | 81.84% | strong | unlikely |
| Spike Severe acute respiratory syndrome coronavirus 2 | IRF3 Canis lupus familiaris | 72.57% | strong | unlikely |
